# Drs2 regulates TRAPPIII in Atg9 transport: exposing the interplay of P4-ATPases and Multisubunit Tethering Complexes

**DOI:** 10.1101/2020.12.25.417865

**Authors:** Irene Pazos, Marta Puig-Tintó, Jorge Cordero, Nereida Jiménez-Menéndez, Marc Abella, Ana G. Duran, Emi Adachi-Fernández, Carla Belmonte-Mateos, Susana Sabido-Bozo, Altair C. Hernández, Sébastien Tosi, Akiko Nezu, Julien Colombelli, Todd R. Graham, Tamotsu Yoshimori, Manuel Muñiz, Maho Hamasaki, Oriol Gallego

## Abstract

Multisubunit Tethering Complexes (MTCs) are a set of conserved protein complexes that tether transported vesicles at the acceptor membrane. Interactions with other components of the trafficking machinery regulate MTCs through mechanisms that are just partially understood. Here we systematically investigate the interactome that regulates the function of MTCs. We found that P4-ATPases, a family of lipid transporters involved in the biogenesis of vesicles, interact with MTCs that participate in the anterograde and retrograde transport at the Golgi, such as TRAPPIII. We used the lipid flippase Drs2 as a model to investigate the mechanism and biological relevance of such interplay during the transport of Atg9 vesicles. Binding to the N-terminal tail of Drs2 stabilizes TRAPPIII on membrane compartments loaded with Atg9 and it is required for the delivery of Atg9 during selective autophagy, a role that is independent of previously reported functions of the P4-ATPase. This mechanism relies on the I(S/R)TTK motif nested in the N-terminal tail cavity of Drs2, a motif that is required for the interaction with MTCs.

---

Multisubunit Tethering Complexes (MTCs) are a group of protein complexes essential for vesicle transport. MTCs mediate the tethering of the vesicle to the acceptor membrane through long-range interactions that precede vesicle fusion and cargo delivery at the destination of the transport pathway^1,2^. Eight MTCs have been described in yeast, all conserved in humans^3^. The conserved oligomeric Golgi (COG) complex is a heterooctamer involved in intra-Golgi retrograde transport^4^. Dsl1, with three subunits, is responsible for tethering COPI vesicles derived from the Golgi to the endoplasmic reticulum (ER)^5^. The class C core vacuole/endosome tethering (CORVET) complex and the homotypic fusion and vacuole protein sorting (HOPS) complex share a common core of four subunits and they have two additional specific subunits each. Both complexes act at the endosomal/vacuolar pathway where CORVET tethers vesicles at early endosomes and HOPS functions in late endosomes and the vacuole^6^. The Golgi-associated retrograde protein (GARP) complex is a heterotetramer that tethers vesicles derived from endosomes to the trans-Golgi network (TGN)^7^. The exocyst is a heterooctamer responsible for tethering secretory vesicles to the plasma membrane^8^. Finally, the transport protein particle (TRAPP) complexes have historically been classified as MTCs despite their major role as guanine nucleotide exchange factors for Rab GTPases^9^. TRAPP comes in two flavors: TRAPPII and TRAPPIII^10^. Both complexes have a heterohexamer core called TRAPPI. TRAPPII, with four more specific subunits^11^, activates Ypt31/32 to regulate intra-Golgi transport^12^. TRAPPIII, with the sole specific subunit Trs85^13^, activates Ypt1 to regulate the transport of ER-derived COPII vesicles to the Golgi^13^. MTCs present similarities in their mechanism of function. For instance, during vesicle transport MTCs interact with other components of the trafficking machinery such as GTPases and SNAREs^1,2^. However, studies have been centered on the mechanism of individual MTCs, which limited the understanding of the molecular bases that play a general role in MTCs function.

To shed light on the molecular mechanisms that control vesicle transport, we have systematically searched for interactions that are functionally relevant for MTCs. Our approach combines genetic interactions and PICT (Protein interactions from Imaging Complexes after Translocation), a method to detect interactions by live-cell imaging^14,15^. We found that MTCs involved in the anterograde and retrograde transport at the Golgi (COG, GARP, TRAPPII and TRAPPIII) present a similar interaction pattern with Type IV P-type ATPases (P4-ATPases), a family of lipid translocases that maintain membrane lipid asymmetry by accumulating phospholipids to the cytosolic leaflet. COG, GARP, TRAPPII and TRAPIII are involved in the transport of the Autophagy-related protein 9 (Atg9)^16,17,18,19^, a transmembrane lipid scramblase^20,21^ essential for selective autophagy. Selective autophagy is a pathway that targets the degradation of cellular components in a specific manner to maintain the cell fitness. In growing cells, Atg9 localizes at Atg9 reservoirs such as endocytic compartments, the Golgi and numerous cytoplasmic vesicular structures^22^. However, for the progression of selective autophagy, it is required that Atg9 is delivered to the PAS (pre-autophagosomal structure). Atg9 transport to the PAS precedes the *de novo* formation of the double membrane that engulfs the cargo of selective autophagy and that ultimately will be internalized in the vacuole/lysosomes^19^. P4-ATPases have not been reported to participate in the transport of Atg9 or to play any critical role in selective autophagy.

We challenged the functional relevance of the network between MTCs and P4-ATPases by studying the role of the interaction between TRAPPIII and Drs2 in the transport of Atg9. Drs2 is a P4-ATPase located at the TGN and early endosomes that flips phosphatidylserine (PS) specifically^23,24^. Here, we show that Drs2 is necessary for the correct function of TRAPPIII in transporting Atg9 during selective autophagy, a role that is independent of Drs2 reported functions. Binding to the N-terminal tail of Drs2 stabilizes TRAPPIII on membranes loaded with Atg9. We show that Drs2 is essential to maintain Atg9 homeostasis through a mechanism that requires the I(S/R)TTK motif located in a cavity of the N-terminal tail of P4-ATPases. This motif is conserved in other flippases and is crucial to sustain the network of interactions between Drs2 and MTCs, including GARP, TRAPPII and TRAPPIII.

## Results

### Protein-protein interactions relevant for MTCs function

In an effort to thoroughly explore the space of protein interactions that are relevant for MTCs function, we conducted a genome-wide search based on reported genetic interactions in yeast^25^. Genes coding for MTCs subunits establish 2587 genetic interactions with 1283 other genes. Based on the probability of these genetic interactions to occur randomly, we selected those gene products that are more likely to be functionally linked with MTCs and that have not been shown to bind MTCs before. Overall, we selected 470 proteins for subsequent protein-protein interaction screening (Supplementary Table 1, see Methods).

We then used the PICT assay to screen protein-protein interactions directly in living cells^14,15^(Fig. 1a). PICT is based on the rapamycin-induced heterodimerization of the FK506-binding protein (FKBP) and the FKBP-rapamycin binding (FRB) domain^14,26^. The addition of rapamycin to the media induces the translocation of proteins tagged with FRB (bait-FRB) to intracellular anchoring platforms tagged with FKBP and RFP (anchor-FKBP-RFP). If a GFP-tagged protein (prey-GFP) interacts with the bait-FRB, it will build-up at the anchor upon addition of rapamycin, which can be quantified by the increment in the co-localization of the GFP and RFP signals (see Methods). In a previous study, we enhanced PICT sensitivity up to 200-fold by engineering yeast cells to express anchoring platforms at the spindle pole body (Tub4-FKBP-FRB). By tagging Tub4 with FKBP and RFP, the resulting cells harbor one or two anchoring platforms only. Thus, even low abundant complexes accumulate enough prey-GFP in the anchoring platforms to be efficiently detected and quantified^15^.

**Fig. 1.**
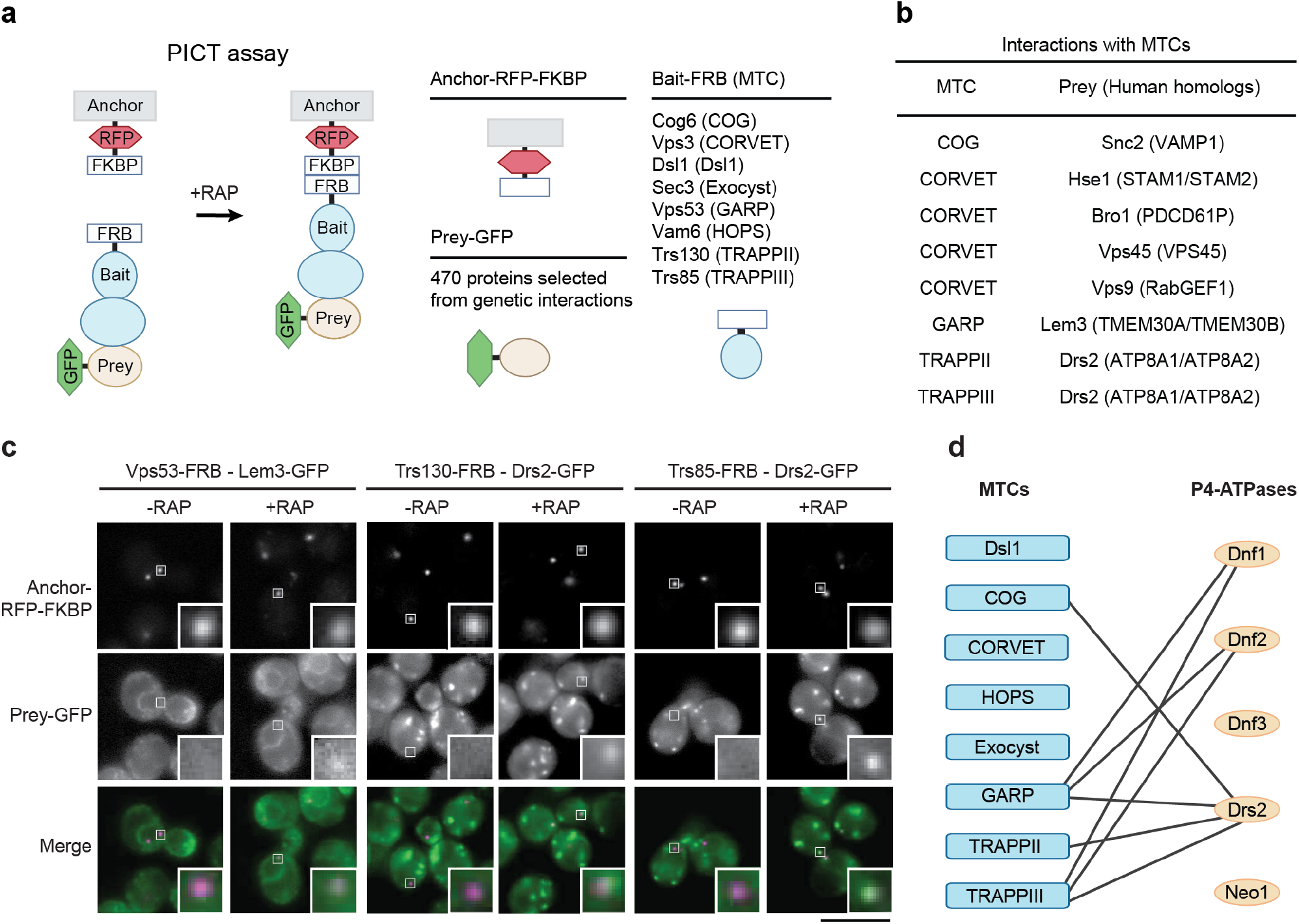
PICT assay and protein-protein interactions relevant for MTCs function. **a** Illustration of the PICT assay (left). Most efficient baits used to anchor each MTC (in parenthesis) are listed (top right) (see Supplementary Table 2). **b** Summary of detected protein-protein interactions. Human homologs are indicated in parenthesis. **c** PICT assay for the interactions between indicated pairs of proteins tagged to FRB and GFP. Representative images before and after adding rapamycin (RAP) of RFP-tagged anchor and GFP-tagged prey are shown in the upper and middle row, respectively. Bottom row shows the merged images. White zoom in boxes, 0.9μm. Scale bar, 5μm. **d** Network of interactions between MTCs and P4-ATPases. MTCs are represented by blue boxes and P4-ATPases by orange oval circles. Black lines show the interactions found with PICT.

This screen identified seven proteins, all conserved in humans, which establish eight interactions with MTCs (Figs. 1b and 1c). The COG complex showed binding to Snc2, an exocytic SNARE known to traffic through the Golgi^27^. CORVET showed binding to Hse1, Bro1, Vps45 and Vps9, all involved in the sorting of vacuolar proteins. These interactions provide molecular bases to the mechanism that underlies the interplay between CORVET and the Vps9-Vps21-Vps45 module^28,29^. We also found that TRAPPII and TRAPPIII bind Drs2, while GARP interacts with Lem3. Drs2 is one of the five yeast P4-ATPases together with Dnf1, Dnf2, Dnf3 and Neo1. Except for the last one, all P4-ATPases act as heterodimers with its ß-subunit: Lem3 for Dnf1 and Dnf2; Cdc50 for Drs2; and Crf1 for Dnf3^30,31^. Thus, the screen identified three MTCs that could bind to P4-ATPases, suggesting a general mechanism underlying these interactions. The interaction between MTCs and P4-ATPases was unexpected given the canonical role of P4-ATPases in early stages of vesicle biogenesis and of MTCs in vesicle tethering at the last steps of vesicle transport. These results prompted us to further investigate the interplay between lipid flippases and MTCs.

### The network of interactions between MTCs and P4-ATPases

To gain a more general view of the relationship between MTCs and P4-ATPases, we expanded the study of protein-protein interactions to all MTCs and P4-ATPases (Fig. 1d). Detected binding events define a network with specific interaction patterns. GARP and TRAPPIII concentrate most of the interactions with the P4-ATPases Drs2, Dnf1 and Dnf2. Note that Dnf1 and Dnf2, which show identical binding specificity with MTCs, are also known to have redundant functions^32^. In addition to GARP, TRAPII and TRAPPIII, Drs2 also interacts with COG, being the flippase that shows the highest number of interactions with MTCs. Overall, MTCs involved in trafficking pathways at the Golgi show preference to bind Drs2, Dnf1 and Dnf2, suggesting that binding to P4-ATPases might be involved in the molecular mechanisms that COG, GARP, TRAPPII and TRAPPIII use to control vesicle trafficking in this organelle. To understand better the molecular mechanism that regulates MTCs function, we further investigated the interactions with Drs2, the only P4-ATPase whose atomic structure was available at the time of this study^33,34^.

### Drs2 regulates the biogenesis of the CVT vesicle

The CVT pathway is used as a model to study selective autophagy in yeast and it initiates with the construction of the PAS. Then, a double membrane elongates from this structure to enwrap the precursor Ape1 (prApe1) aminopeptidase until the double membrane closes generating the CVT vesicle. After fusion and internalization in the vacuole, prApe1 is processed to mature Ape1 (mApe1)^35,36^.

In cells lacking TRAPPIII subunit Trs85 (*trs85*Δ), Atg9 is not delivered to the PAS, the biogenesis of the CVT vesicle is blocked and Ape1-GFP is accumulated in large cytosolic aggregates^19,37^ (Figs. 2a and 2b). Because Drs2 is essential for cell growth below 21°C^38^, we first analyzed the contribution of Drs2 in the CVT pathway at different temperatures. While Ape1 processing is fully blocked in *trs85*Δ cells grown at any of the temperatures tested, cells lacking Drs2 (*drs2*Δ) show a progressive inhibition of Ape1 maturation when decreasing the temperature. Thus, *drs2*Δ cells process prApe1 near to normality at 37°C, while we could only detect traces of mApe1 in cells incubated at 16 °C (Supplementary Fig. 1a). We continued studying the interplay between Drs2 and TRAPPIII at 23°C to avoid undesired effects derived from impaired cell fitness. At this temperature, cells lacking Drs2 mature 11.5% of the aminopeptidase (Fig. 2a). We confirmed that such levels of mApe1 represent the steady-state levels of the CVT pathway in mutant cells grown at 23°C by analyzing Ape1 processing in cells lacking Drs2 and Atg19 (i.e. specific receptor for the CVT pathway), Drs2 and Atg17 (i.e. specific scaffold for non-selective autophagy) and in a time course assay (Supplementary Figs. 1b and 1c). Consistently with a functional relationship, cells lacking either Trs85 or Drs2 accumulate similar Ape1-GFP aggregates (Fig. 2b).

**Fig. 2.**
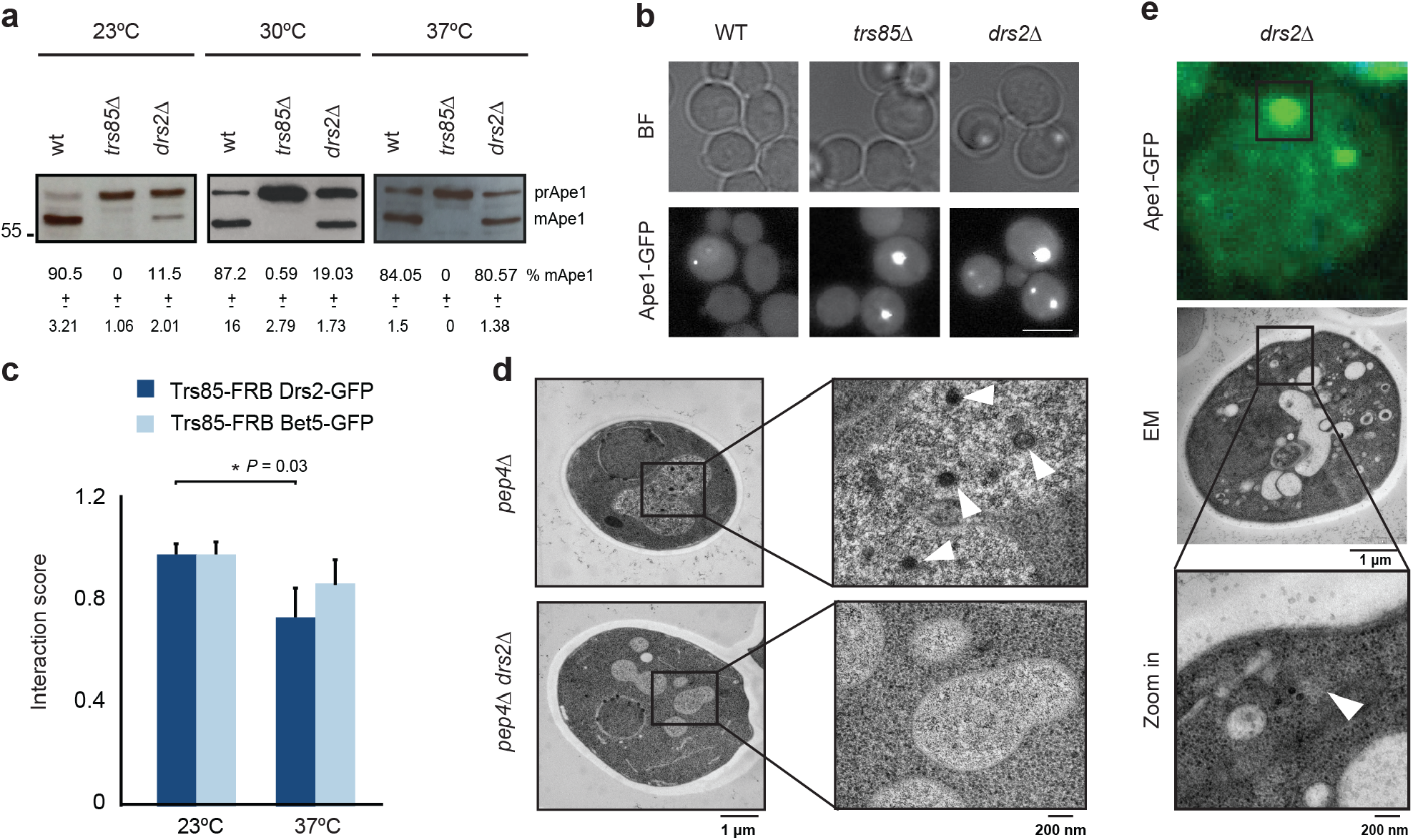
Drs2 is critical for the biogenesis of the CVT vesicle. **a** Processing of Ape1 was analyzed by western blot in the indicated strains and temperatures. Below, percentage of mApe1 ± SD, *n =* 3. **b** Aggregation of Ape1. Representative images of Ape1-GFP in wild-type, *trs85*Δ and *drs2*Δ cells. BF, brightfield images. Scale bar, 5μm. **c** PICT assay for the interaction of Trs85-FRB with Drs2-GFP or Bet5-GFP. Interaction score was normalized to the measurement at 23°C. Error bars: mean ± SD, *n =* 3. Asterisk indicates significant difference as determined by a two-tailed Student’s t-test (P<0.05). **d** Representative EM images of *pep4*Δ and *pep4Δdrs2Δ* strains. Black squares show a zoom-in in the vacuole. White arrowheads point to CVT bodies. **e** Representative CLEM images of *drs2*Δ cells. Ape1-GFP (top) correlates with a membrane-free ribosome exclusion area (middle). Black square (bottom) shows a zoom-in at the position correlating with Ape1-GFP (white arrowhead).

PICT assay showed that, concomitantly to an exacerbation of the Drs2 role in the CVT pathway, the Drs2-TRAPPIII interaction is boosted under colder conditions: at 23°C the Interaction score between Drs2 and Trs85 is 24% higher than at 37°C (Fig. 2c), while no significant difference could be detected for the interaction between Trs85 and Bet5, a subunit of TRAPPI. This observation suggests that the cell regulates the interplay between Drs2 and TRAPPIII in response to temperature shifts. We then used electron microscopy (EM) to study the ultrastructure of the CVT pathway. Pep4 is a protease required for the processing of the CVT vesicle once it is internalized in the vacuole^39^. While 89.3% of the *pep4*Δ cells presented CVT bodies inside the vacuole, we could not detect any CVT body-like structure in the vacuoles of *pep4*Δ*drs2*Δ cells (Fig. 2d). We then analyzed Ape1-GFP aggregates in *drs2*Δ cells with Correlative Light and Electron Microscopy (CLEM)^40^. 91.4% of the Ape1-GFP spots correlated with a ribosome exclusion area of amorphous shape and that was devoid of double membrane or CVT vesicle-like structure (Fig. 2e). Furthermore, co-localization assays show that Ape1-GFP aggregates in the cytosol of *drs2*Δ cells do not reach the vacuole (Supplementary Video 1).

Overall, these results point to a mechanism where Drs2 is necessary for the biogenesis of the CVT vesicle, a role that is further underscored by the enrichment of DRS2 genetic interactions with genes coding for proteins involved in autophagy (*P =* 0.002) (see Methods).

### The role of Drs2 in the CVT pathway is independent from known mechanisms

Drs2 is a lipid flippase with multiple functions in vesicle transport at the interface of endosomes and the Golgi^23,24^. We used yeast genetics to investigate the possibility that defects in the CVT pathway in *drs2*Δ cells resulted indirectly from the perturbation of other processes regulated by the flippase. However, Ape1 processing was not affected in cells lacking Rcy1 (i.e., cells where the transport from early endosomes to TGN is blocked)^24,41,^, Apl4 (i.e., cells where the AP-1 pathway from TGN to early endosomes and the exit of the high-density class of vesicles from the TGN are blocked)^42,43^, Apl5 (i.e., cells where ALP pathway from TGN to the vacuole is blocked)^44^ or both Gga1 and Gga2 (i.e., cells where CPY pathway from TGN to late endosomes is blocked)^45^ (Fig. 3a). Thus, genetic data suggest that the biogenesis of the CVT vesicle does not involve the pathways where Drs2 has been reported to participate. We then explored the molecular mechanism of Drs2 required for the biogenesis of the CVT vesicle. We analyzed the rescue of Ape1 processing in *drs2*Δ cells expressing different constructs of Drs2 (Fig. 3b). All constructs were expressed under the control of DRS2 promoter.

**Fig. 3.**
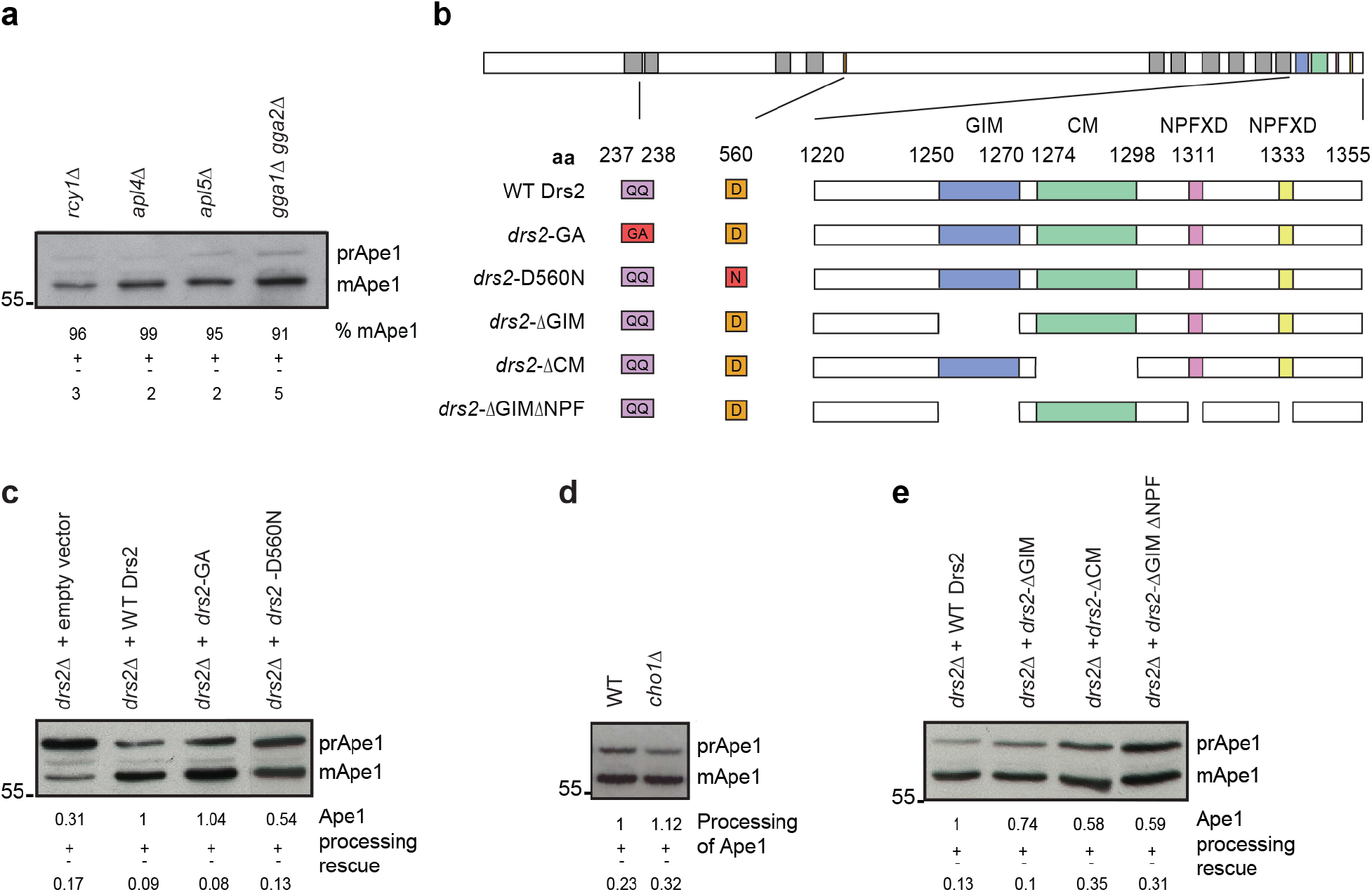
Drs2 role in the CVT pathway is independent from its known mechanisms. **a, c-e** Processing of Ape1 was analyzed by western blot in the indicated strains, *n =* 3. **a** Below, percentage of mApe1 ± SD. **b** Representation of the main structural features of Drs2. Each color in the top bar indicates the location of a feature in the sequence of wild-type Drs2. Grey boxes depict transmembrane domains. Below, a summary of the mutants tested and an enlarged view of their C-terminal tail. QQ>GA and D>N mutations are depicted in red. Numbers indicate the residues position in Drs2 sequence. **c**, **e** Cells harboring an empty plasmid and/or expressing wild-type Drs2 were used as control. Below, mean rescue of Ape1 processing normalized to the rescue achieved in cells expressing wild-type Drs2 ± SD. **d** Below, processing of Ape1 normalized to the processing achieved in wild-type cells ± SD.

We first targeted the PS flipping activity of Drs2, which is required for the bidirectional transport endosome-TGN and for the exit of secretory vesicles from the TGN^46,47,48^. While cells harboring an empty vector do not rescue Ape1 processing in *drs2*Δ cells, cells expressing a mutant unable to flip PS (*drs2*-GA) and cells expressing an ATPase-dead mutant form of Drs2 (*drs2*-D560N) rescue either totally or partially the CVT pathway (Fig. 3c). Cells lacking Cho1, an enzyme essential for the synthesis of PS^49^, show normal processing of Ape1, confirming that flipping of PS is not required for the CVT pathway (Fig. 3d).

The cytosolic C-terminal tail of the flippase is also important for the known functions of Drs2. For instance, the Gea2 interacting motif (GIM) is required for the Drs2-Gea2-Arl1 complex assembly, a mechanism based on protein-protein interactions that regulate a number of trafficking pathways at the TGN and endosomes^50,51^. Adjacent to the GIM, Drs2 presents a conserved motif (CM). CM is fundamental for Drs2 function and to bind to the cytosolic domains of Drs2 that stabilize the autoinhibitory conformation^33,51,^. Drs2 has also two NPFXD motifs (hereafter referred to as NPF motif) that constitute some of its multiple endocytosis signals^52^. However, expression of *drs2-*ΔGIM, *drs2-*ΔCM or *drs2-*ΔGIMΔNPF also rescues more than 50% of Ape1 processing (Fig. 3e). Thus, the mutagenesis experiments indicate that known functional motifs of the Drs2 C-terminal tail are not essential for the role of the flippase in the CVT pathway.

### The I(S/R)TTK motif is required for the function of Drs2 in the CVT pathway

We then investigated the cytosolic N-terminal tail of Drs2. Although P4-ATPases present low homology in this region, multiple sequence alignment identified a conserved stretch of 15 amino acids (residues 198-212 in Drs2; Fig. 4a). The stretch defines a cavity of 18.8 Å in depth located adjacent to the first transmembrane helix in the structure of Drs2 (Fig. 4b). Homology modeling suggests that all yeast P4-ATPases present the N-terminal cavity in their structure (Supplementary Fig. 2), although the function of this cavity is not known. We observed that the bottom of the cavity of those P4-ATPases that bind MTCs (Dnf1, Dnf2 and Drs2) is characterized by an I(S/R)TTK motif that is lacking in the cavity of Dnf3 and Neo1 (Fig. 4a).

**Fig. 4.**
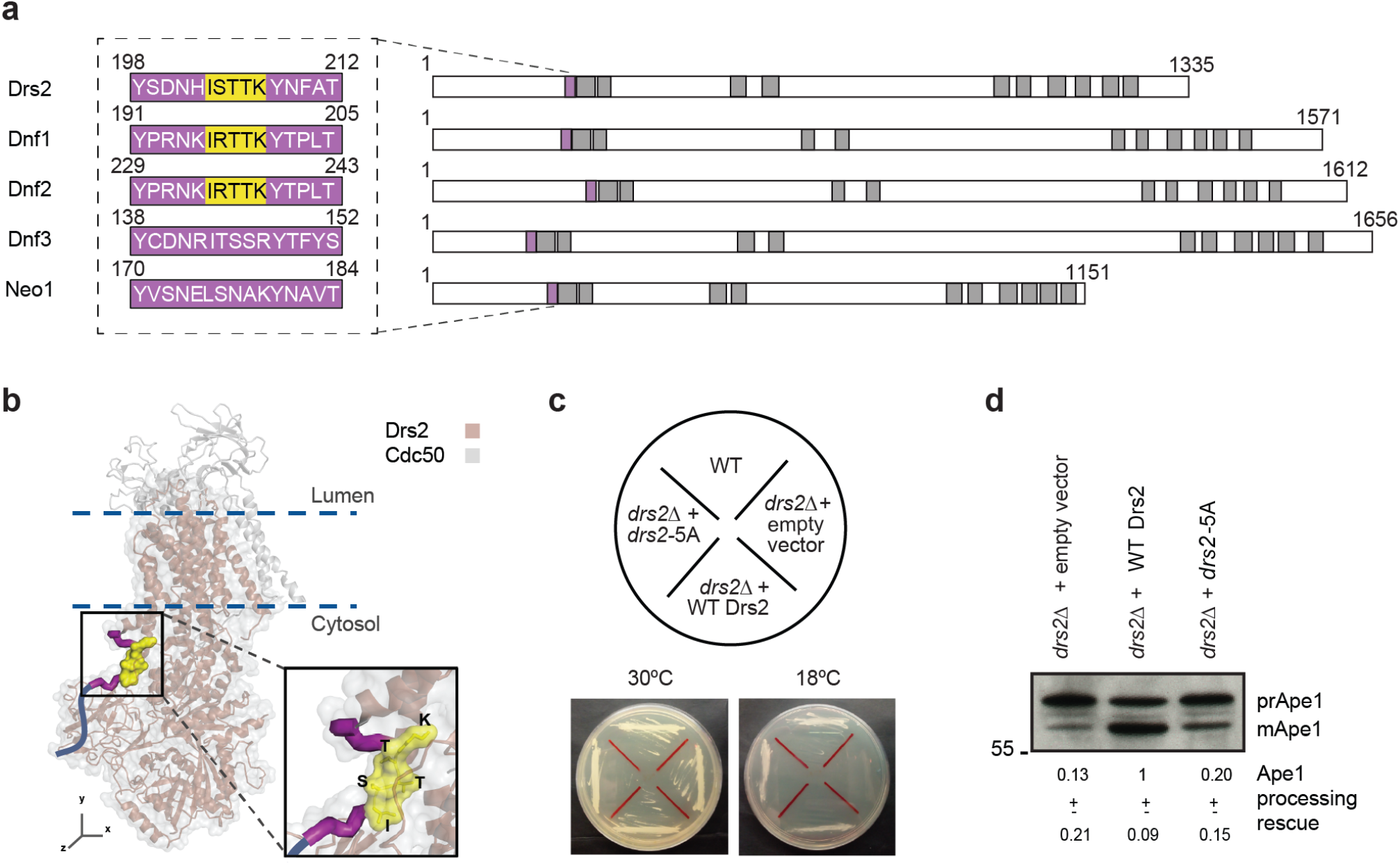
The I(S/R)TTK motif is required for the assembly of the Drs2-TRAPPIII module and its function in the CVT pathway. **a** Representation of the 15 aminoacid (aa) cavity (purple) in the N-terminal tail of P4-ATPases. Grey boxes depict transmembrane domains. Black dashed box shows the sequence coding for the cavity (left). Yellow background highlights the I(S/R)TTK motif in Drs2, Dnf1 and Dnf2. **b** The I(S/R)TTK motif in the Drs2-Cdc50 structure (PDB ID: 6ROH). Black box zooms in the cavity of Drs2 (purple) and the I(S/R)TTK motif (yellow). Dark blue line depicts the unstructured N-terminal region. **c** Cold-sensitive assay for the indicated strains. Cells streaked onto YPD plates were incubated at 30°C and 18°C for three days. **d** Processing of Ape1 was analyzed by western blot in the indicated strains. Below, mean rescue of Ape1 processing normalized to the rescue achieved in cells expressing wild-type Drs2 ± SD, *n* = 3.

The correlation between this structural feature and the P4-ATPases-MTCs interaction pattern motivated us to investigate the relevance of the I(S/R)TTK motif. We first generated cells expressing Drs2 with its I(S/R)TTK motif substituted by five alanines (*drs2*-5A). The expression of *drs2-*5A rescues the cold-sensitive phenotype of *drs2*Δ cells, confirming that the loss of this motif does not induce general protein unfolding (Fig. 4c). However, the expression of *drs2*-5A in *drs2*Δ cells cannot rescue Ape1 processing (Fig. 4d), which accumulates in large cytosolic Ape1-GFP aggregates similar to those seen in *drs2*Δ cells (Supplementary Video 2). These results prompted us to investigate the molecular bases that control the function of the Drs2 I(S/R)TTK motif.

### Drs2 I(S/R)TTK mediates proximal protein interactions with MTCs

To systematically analyze the mechanism mediated by the Drs2 I(S/R)TTK motif, we performed crosslinking-mass spectrometry (XL-MS) immunoprecipitation experiments (see Methods). We performed a partial crosslinking with disuccinimidyl sulfoxide (DSSO), a crosslinker of 10.1Å in length that allowed us to detect proximal peptides in cell extracts^53^. Upon a GFP-specific pulldown, the comparative analysis between the interactome of Drs2-GFP and *drs2-*5A-GFP identified 26 proteins that are likely to bind directly Drs2-GFP and whose binding requires the I(S/R)TTK motif (Supplementary Fig. 3). This set of interactions is enriched with proteins involved in endosomal transport (*P*=3.9 E-06), Golgi vesicle transport (*P*=3.5 E-05) and endosome-to-Golgi transport (*P*=4.9 E-08). On the other side, known protein-protein interactions required for canonical Drs2 functions were not affected (e.g., binding to Cdc50 or Rcy1). Thus, in agreement with the cold-sensitivity rescue assay (Fig. 4c), the interactome data suggest that the mutation of the I(S/R)TTK motif allows us to dissect different mechanisms controlled by Drs2.

Interestingly, *drs2-*5A-GFP could not recapitulate the interaction with Trs85, (the specific subunit of TRAPPIII), which was further confirmed by PICT (Figs. 5a and 5b). The interaction between Drs2 and Trs65, Trs120, Trs130 (specific subunits of TRAPPII), and Vps53 (GARP) were also destabilized upon mutation of the I(S/R)TTK motif. Thus, the interactome data is in agreement with the hypothesis that the I(S/R)TTK motif regulates the network of interactions between P4-ATPases and MTCs, including the interaction Drs2-TRAPPIII (Fig. 5a and Supplementary Fig. 3).

**Fig. 5.**
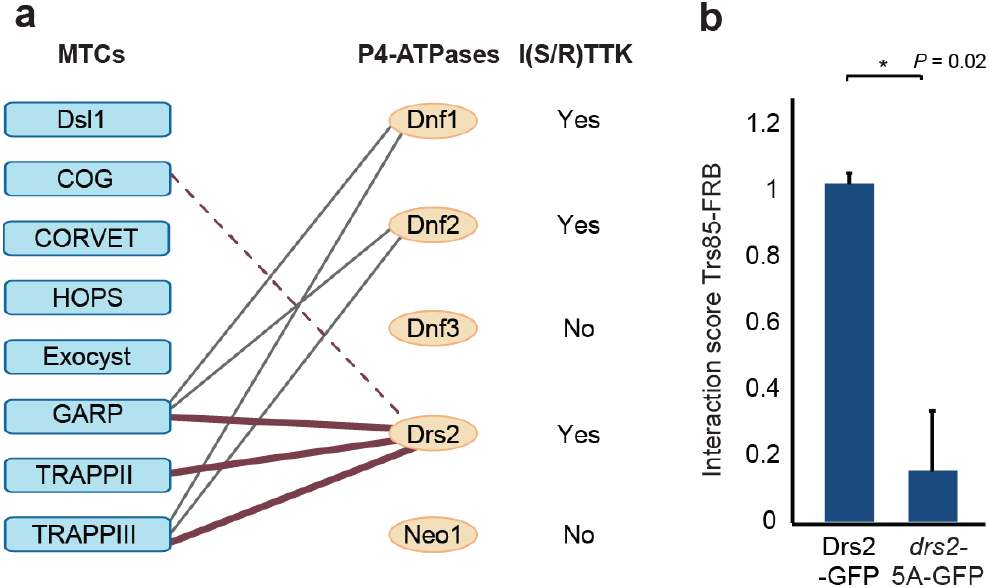
The I(S/R)TTK motif and the interaction between MTCs and P4-ATPases**. a** Network of interactions between MTCs and P4-ATPases (lines) adapted from Fig 1d. XL-MS detected proximal interactions that require the I(S/R)TTK motif (thick maroon lines). The interaction with COG could not be recapitulated with DSSO (dashed line) **b** Interaction between Trs85-FRB and Drs2-GFP or *drs2-*5A-GFP analyzed by PICT. The Interaction score was normalized to the measurement of the Drs2-GFP. Error bars: mean ± SD, *n =* 2. Asterisk indicates significant difference as determined by a two-tailed Student’s t-test (P<0.05).

### Drs2 I(S/R)TTK is fundamental for Atg9 homeostasis

In growing yeast cells, Atg9-GFP can be observed as highly mobile puncta, the vast majority of which correspond to vesicles within the cytoplasm^22^. We implemented a microscopy-based system with high-sensitivity and high-temporal resolution to investigate the role of Drs2 in the trafficking of Atg9 vesicles (see Methods). In agreement with other publications^22^, growing wild-type cells present two populations of Atg9 puncta (Fig. 6a and Supplementary Video 3). The larger population comprises 86% of the puncta and it is highly mobile (mean speed of 2.3 ± 0.7 nm/ms). The smaller population comprises 14% of the puncta and it has slower mobility (mean speed of 0.7 ± 0.2 nm/ms). Interestingly, in *drs2*Δ and *drs2*-5A cells, the low mobility population increases up to 54% (0.6 ± 0.3 nm/ms) and 50% (0.6 ± 0.2 nm/ms) of the puncta, respectively (Fig. 6a and Supplementary Videos 4 and 5). We could not detect Drs2-GFP at the PAS, (Supplementary Fig. 4). Thus, vesicle tracking demonstrates that the I(S/R)TTK motif of Drs2 plays a major role in the trafficking of Atg9 prior to reaching the PAS, most likely in the biogenesis of higher mobility vesicles from lower mobility Atg9 reservoirs.

**Fig. 6.**
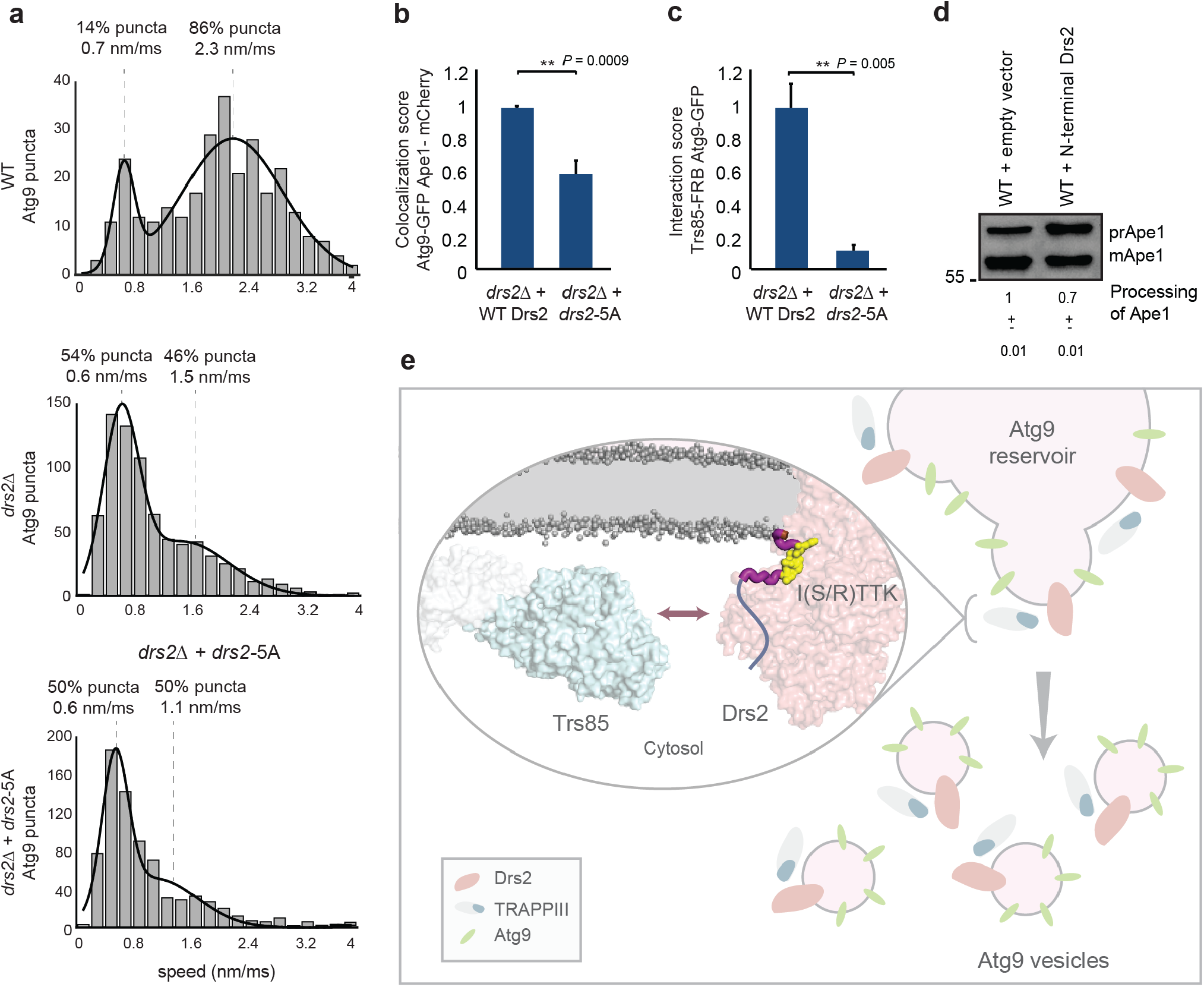
I(S/R)TTK motif of Drs2 plays a central role in the transport of Atg9. **a** Histograms representing the mean speed of Atg9 puncta observed in wild-type*, drs2*Δ and *drs2-*5A cells. The histograms were fitted to a mixture of Gaussian distributions with two components; the percentage of each population and the means of the fitting curves are indicated on top of the plot. **b** Co-localization between Atg9-GFP and Ape1-mCherry in cells expressing wild-type Drs2 or *drs2-*5A mutant. **c** PICT assay for the Trs85-FRB (TRAPPIII) and Atg9-GFP interaction in cells expressing wild-type Drs2 or *drs2-*5A mutant. **b, c** Values were normalized to the measurements in cells expressing wild-type Drs2. Error bars: mean ± SD, *n =* 3. Asterisks indicate significant difference as determined by a two-tailed Student’s t-test (P<0.05). **d** Processing of Ape1 was analyzed by western blot in cells overexpressing N-terminal Drs2 (1-212 aa). Below, processing of Ape1 normalized to the processing achieved in cells with an empty vector ± SD, *n =* 3. **e** Model for the interplay between Drs2 and TRAPPIII. TRAPPIII binds the N-terminal fragment of Drs2 on the surface of a membrane loaded with Atg9. This interaction requires the I(S/R)TTK motif and it is neceessary for the correct biogenesis of Atg9 vesicles from Atg9 reservoirs, becoming essential to sustain the CVT pathway at low temperatures.

We then used live-cell imaging to investigate TRAPPIII function in cells expressing mutated Drs2 (*drs2*-5A). Defects in the TRAPPIII-dependent transport of Atg9 vesicles can be tracked as a decrease of the co-localization between Atg9-GFP and Ape1-mCherry clusters. Although they accumulate Ape1 aggregates in the cytosol (Fig. 2), the presence of Atg9-GFP at the Ape1-mCherry clusters was 38.5% lower in *drs2-*5A cells than in wild-type cells (Fig. 6b). The PICT assay provides a good read-out to measure the capability of proteins to bind membranes *in situ*^54^. We inferred TRAPPIII association to Atg9-loaded membranes by measuring the recruitment of Atg9-GFP to the anchoring platform when using Trs85-FRB as bait. Indeed, the deletion of the I(S/R)TTK motif of Drs2 results in an 84.1% inhibition of the Interaction score between TRAPPIII and Atg9-GFP (Fig. 6c). The residual ability of TRAPPIII to translocate Atg9-loaded membranes to the anchoring platform and partial delivery of Atg9 to the PAS is in agreement with the observation that the CVT pathway remains slightly active in *drs2-*5A cells under our experimental conditions (Fig. 4d).

PICT assay cannot resolve if the detected binding to Atg9-loaded membranes results from protein-protein interactions or binding to the membrane. Either way, this experiment suggests that Trs85 binding to the N-terminal fragment of Drs2 stabilizes TRAPPIII on Atg9-loaded membranes. This mechanism is further supported by the results of a competition assay, where overexpression of the Drs2 N-terminal tail, including the I(S/R)TTK motif of Drs2 (amino acids 1-212), caused a 30% inhibition of Ape1 processing (Fig. 6d). Overall, these results suggest that the assembly of the Drs2-TRAPPIII module is necessary for the normal function of TRAPPIII in the transport of Atg9 and selective autophagy (Fig. 6e).

## Discussion

Understanding the mode of action of MTCs requires an approach dedicated to compare, side-by-side, the mechanistic features of these protein machines. Genetic interactions and the PICT assay allowed us to systematically explore transient and low abundant binding events relevant to the function of MTCs. We found that MTCs localizing at the Golgi recognize P4-ATPases, establishing a network of interactions with unknown biological implications. To delve into the functional relevance and the molecular mechanism of the interplay between MTCs and P4-ATPases, we analyzed, in the context of Atg9 transport and the CVT pathway, the interaction between TRAPIII and the lipid flippase Drs2.

TRAPPIII is a MTC that controls the transport of Atg9 between reservoirs at the endocytic structures and the Golgi. TRAPPIII is essential so that a small fraction of Atg9 can eventually be delivered to the PAS for the formation of the CVT vesicle. Remarkably, both TRAPPIII and Drs2 have been detected on Atg9 vesicles^21,55^. We found that under mild (30°C) and cold (≤23°C) environments, Drs2 is also required for the CVT pathway. Which is the molecular basis of the role of Drs2 in this process and whether this involves the interaction with TRAPPIII was not known.

Recent structural analysis of TRAPPIII showed that the Trs85 subunit binds membranes through a positively charge amphipathic helix, an interaction that is required for TRAPPIII function in non-selective autophagy and the secretory pathway. It has also been postulated that Trs85 might be stabilized on the membrane surface through additional protein-protein interactions that target TRAPPIII to specific compartments^56,57^. The mechanism of TRAPPIII to recognize Atg9-loaded membranes remains to be elucidated.

A possibility would be that Drs2 accumulated negatively charged PS at the cytosolic leaflet of Atg9 compartments and that, in turn, PS stabilized TRAPPIII through interactions with positive charges of the Trs85 amphipathic helix. However, genetics showed that the role of Drs2 in the CVT pathway is independent of previously reported mechanisms of the lipid flippase, including lipid translocation. Electrostatic interactions between Drs2-flipped PS and Trs85 amphipathic helix on the surface of Atg9 vesicles are not necessary for the CVT pathway as cells that cannot synthesize PS process Ape1 normally (Fig. 3).

Combining genetic interactions, live-cell imaging, ultrastructure analysis and XL-MS, we propose that binding to Drs2 stabilizes TRAPPIII on membranes where Atg9 is embedded and we provide insight into the mechanism that regulates the assembly of the Drs2-TRAPPIII module. We demonstrate that the I(S/R)TTK motif is indeed required for the Drs2-TRAPPIII interaction and to stabilize TRAPPIII on Atg9-loaded membranes (Fig. 5 and 6)^57^. The I(S/R)TTK motif is nested at the bottom of the cavity of Drs2 N-terminal tail, linking the first transmembrane helix and the unstructured region of the Drs2 N-terminal^33^. Although the cavity is conserved in the atomic structure of yeast and human P4-ATPases^33,34,58,59,60^, its function is not known. The presence of the I(S/R)TTK motif in yeast Dnf1 and Dnf2 and the related ISTAK motif in the human ATP8A2, a homolog of Drs2, suggests that this structural element is critical for the molecular mechanism of some P4-ATPases (Supplementary Figs. 2 and 5).

Although it is possible that Trs85 directly binds the unstructured region of the Drs2 N-terminal tail, the lack of sequence homology among the P4-ATPases that bind MTCs, hints that this is unlikely to be a conserved mechanism. Alternatively, TRAPPIII subunit Trs85 might bind directly the N-terminal cavity of Drs2. However, the accessibility to this binding pocket is sterically constrained. Recent structural analysis of human ATP8A1 showed that the cavity is momentarily opened in the transition between the E2P and E2Pi-PL states along the lipid translocation cycle. During this rearrangement, the neck of the cavity switches from a close conformation of 14.1Å to an open conformation of 20.6Å (Cys50-to-Asn60)^58^. This allows us to speculate that the Drs2 cavity might undergo similar conformational dynamics during the assembly of the Drs2-TRAPPIII module, which would expose the I(S/R)TTK motif. Although the exact amino acids that mediate the binding between Drs2 and Trs85 remain to be elucidated, the competition assay with the Drs2 N-terminal tail underscores that the interaction between TRAPPIII and the Drs2 N-terminus is central for the CVT pathway.

As we could not detect Drs2-GFP in the PAS, we reasoned that the Drs2-TRAPPIII module is likely not involved in the tethering of Atg9 vesicles at this compartment (Supplementary Fig. 4). Instead, co-localization and vesicle tracking experiments indicate that Drs2 is critical for the release of Atg9-loaded vesicles from the Atg9 reservoirs (Fig. 6). In fact, XL-MS further supports this new role of Drs2, as the mutation of Drs2 I(S/R)TTK motif hinders the binding to six proteins that bind Atg9: Trs85, Vps17, Pep1, Vps45, Akr1 and Vti1, the last two being components of the Atg9 vesicle^21^ (Supplementary Fig. 3). The enhancement of the Drs2-TRAPPIII module assembly in response to temperature decrease indicates that the interaction between Drs2 and TRAPPIII is subjected to regulation in response to environmental cues. Noteworthy, similar to our observations at 37°C, a former study showed that Drs2 is dispensable for the processing of Ape1 at the stationary phase^18^, indicating that there must be a mechanism that regulates Drs2 function accordingly to the cell needs.

Our approach also provided mechanistic insight about the interplay between Drs2 and the other MTCs. We used mutagenesis and XL-MS to investigate the interactome that is in close proximity to Drs2. We could not detect binding of Drs2-GFP to any subunit of the COG complex, suggesting that Drs2 and COG do not interact directly. However, we confirmed that, in addition to TRAPPIII, the Drs2 I(S/R)TTK motif is required for the binding to GARP and TRAPPII, and that the three MTCs are likely to bind the lipid flippase directly. These results indicate that the interplay between Drs2 and MTCs is likely not limited to the action of the Drs2-TRAPPIII module, but it might include additional mechanisms such as the binding to TRAPPII and GARP, two MTCs that also participate in the transport of Atg9 vesicles^16,19^.

MTCs, which are fundamental to sustain vesicle transport in eukaryotic cell, base their mechanism of action on protein-protein interactions with other components of the trafficking machinery, such as GTPases, SNAREs and SM proteins. In this study, we reveal that biding to P4-ATPases is also important for MTCs function, and in particular, for the biogenesis of Atg9 vesicles. Overall, binding to P4-ATPases arises as a critical mechanism to understand MTCs function, with implications that likely go beyond the transport of Atg9 and selective autophagy. For instance, the role of TRAPPIII in the secretory pathway stresses the need to perform further studies to characterize the function of the Drs2-TRAPPIII module in other processes, such as the transport of COPII vesicles.

## Methods

### Yeast strains, plasmids and cultures

*Saccharomyces cerevisiae (S. cerevisiae*) strains are derivatives of BY4741 or BY4742 backgrounds. Strains with genes coding for C-terminal tags, deleted genes, and/or mutated alleles were generated by homologous recombination following standard PCR strategies^64^. Those strains harboring exogenous genes coded in plasmids were generated by transformation. All Drs2 constructs were expressed from a plasmid using the DRS2 promoter except for those plasmids in Fig. 6d, which use GAL1 promoter. All strains are listed in Supplementary Tables 1-4. All plasmids used are listed in Supplementary Table 5.

Strains used for the PICT assay express the anchor Tub4-RFP-FKBP and harbor different combinations of baits (FRB fusions at the C-terminus) and preys (GFP fusions at the C-terminus) (see Supplementary Tables 1, 2, 3)^15^. Parental cells expressing the anchor Tub4-RFP-FKBP and a bait-FRB were constructed on the OGY0307 genetic background. Parental cells expressing the prey-GFP were constructed in the BY4741 genetic background or obtained from the genomic C-terminal GFP fusion collection (ThermoFisher). Excluding those strains harboring plasmids that code for Drs2-GFP and *drs2*-5A-GFP, the PICT experiments were done with yeast strains constructed with the SGA approach^65^.

Yeast cells were grown in YPD liquid medium at 30°C and 220 rpm until saturation (≈16h). Cells harboring plasmids were grown in synthetic minimal liquid medium lacking the amino acid of choice. In experiments with *cho1Δ* cells, the medium was supplemented with 1mM Ethanolamine. After reaching saturation, cells were diluted to an optical density of OD_600_=0.2 in the medium of interest and cultured at 30°C and 220 rpm until they reached an early logarithmic phase (OD_600_≈1) (experiments Fig. 1). Unless indicated, in the rest of experiments cells were then incubated at 23°C or 37°C for two hours before performing the experiment.

### Imaging

After saturation in the YPD culture, cells were diluted and grown in Low Flo medium (Yeast Nitrogen Base Low Fluorescence without Amino acids, Folic Acid and Riboflavin, 2% Synthetic complete Mixture Drop-out (lacking essential amino acids as required) and 2% glucose). When they reached an early logarithmic phase (OD_600_≈1), cells were attached to Concanavalin A (0.1μg/mL)-coated glass bottom plates. When required, 10μM of rapamycin was added 20-30 minutes before imaging. Images in Fig. 1, 2 and 5 and Supplementary Video 1 were acquired on an ScanR (Olympus) microscope based on a IX81 stand, equipped with 100x/1.40 objective lens, an Orca-ER camera (Hamamatsu), a MT20 Xenon arc lamp illumination system holding excitation filter and two complete fluorescence filter cubes from AHF respectively optimized for GFP (ET Bandpass 470/40 + Beamsplitter 500 DVXRUV + ET Bandpass 525/50) and RFP (ET Bandpass 545/30 + Beamsplitter 580 DVXRUV + Bright-Line HC 617/73). Images in Fig. 6b, 6c and Supplementary Video 2, were acquired on a Nikon ECLIPSE Ti2-E inverted microscope equipped with a 100x/1.49 SR TIRF objective (Nikon), a sCMOS Zyla 4.2 Andor camera and a SpectraX Lumencore LED system with two fluorescence filter cubes (Semrock) optimized for GFP (LF488-C) and LED-mCherry. For the PICT experiments shown in Fig. 1b and 1d, automated imaging acquisition was carried out. The acquisition software (ScanR, Olympus) was set up to detect those cells’ planes with the highest RFP signal. This provided a focal plane that captured the highest number of anchoring platforms in focus. Images were obtained sequentially by switching the filter cubes. Strains were imaged in the RFP (200ms) and GFP channels (200 and 1500ms in Fig. 1b; 200 and 1000ms in Fig. 1d) and each strain was imaged in at least 6 fields of view. In PICT experiments in Fig. 1c, 2c and 5b, for the imaging in Fig. 2b and in Supplementary Video 1 cells were imaged using Xcellence (Olympus) as acquisition software. For the co-localization assay in Fig. 6b, in PICT experiments in Fig. 6c, and in Supplementary Video 2 cells were imaged using NIS-Elements as acquisition software. In experiments in Fig. 1c, 2b, 2c, 5b, 6b and 6c strains were imaged in brigthfield and in the RFP (200ms) and/or the GFP (exposition time was optimized manually) channels. For Supplementary Videos 3, 4 and 5, see Vesicle tracking section.

### Protein interactions from Imaging Complexes after Translocation (PICT)

PICT technique allows the quantitative characterization of protein interactions *in vivo*. The addition of rapamycin to culture media induces FRB-FKBP heterodimerization and translocation of the protein tagged to FRB (bait-FRB) to anchoring platforms engineered in the cell by tagging an anchoring molecule to RFP and FKBP (Anchor-RFP-FKBP). GFP-tagged proteins (prey-GFP) that are bound to the bait-FRB are co-translocated to these anchoring platforms. These changes in localization can be subsequently quantified using dual-color live-cell imaging. Recently, we increased PICT sensitivity up to 200-fold. We designed a new anchoring platform by tagging Tub4, a component of the spindle pole body, with RFP and FKBP (Tub4-RFP-FKBP). The resulting cells harbor one or two anchoring platforms only. Thus, even low abundant complexes accumulate enough prey-GFP in each of these anchoring platforms to be efficiently detected and quantified^15^. All PICT experiments were performed in rapamycin-insensitive strains carrying the *tor1-1* mutation in the TOR1 gene and where the endogenous FPR1 gene was deleted^14^. Images were taken prior and after rapamycin addition.

### Selection of proteins for the protein-protein interaction screen

First, we identified the bait-FRBs of each MTC that allows anchoring more efficiently the fully assembled complex (Fig. 1a and Supplementary Table 2). For each MTC we generated a set of strains expressing all possible combinations of subunits tagged to FRB and GFP. Then, we selected the corresponding bait-FRB that showed highest Interaction score for the other subunit of the corresponding complex.

Second, we used genetic interactions to select the prey-GFPs for the screen. *S. cerevisiae* has 8 MTC complexes, each of them with a different protein composition. Genes coding for MTC subunits (MTC Gene or GM) interact genetically with 1283 genes (Candidate Genes or CG) in yeast (SGD September 2013). In total, *S. cerevisiae* has 5820 genes. For each Candidate and MTC Gene pair CG_i_-GM_j_, we calculated its pairwise interaction probability *P_ij_* given by:
*P_ij_ =PC_ij_*PM_ji_*
where *PC_ij_* is the probability that a Candidate Gene *i* interacts with MTC Gene *j*; and *PM_ji_* is the probability that MTC Gene *j* interacts with Candidate Gene *i*.

When a Candidate Gene *i* interacts genetically with a set *J* of subunits of a given MTC, we use a genetic score *S_i_* defined as

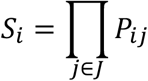

to measure the probability of measuring randomly the genetic interaction profile with Candidate Gene *i.*

676 Candidate Genes with a “Genetic Interaction score” < 10^−6^ were selected for further analysis. We then selected the corresponding 470 prey proteins that are included in the yeast C-terminally GFP-tagged genomic collection^66^ and whose binding to MTC subunits had not been described at the time of the screening (Supplementary Table 1).

### Image analysis

A custom image analysis workflow was implemented as an ImageJ (http://rsb.info.nih.gov/ij/) macro to automate operations and process whole data sets. Essentially, this workflow aimed to estimate the co-localization (overlap) between spots independently segmented in the images from RFP (anchors) and GFP (preys) channels.

To exclude out of focus (or empty) images, a sharpness factor was first computed (standard deviation of the image intensity normalized to its mean intensity in RFP channel). The fields of view with sharpness factor above a user defined threshold were processed by the following sequence of operations aiming to segment bright spots: 1) smoothening, 2) background subtraction, 3) local thresholding, 4) median filtering and 5) discard small particles below a certain area. For each experimental condition, the functional parameters of the workflow were manually optimized to lead to the best results.

The area of overlap between preys and anchors was then estimated by counting the number of foreground pixels common in both segmentation masks (logical AND operation). This area was then normalized to the count of foreground pixels in the RFP segmentation mask (anchors area) to lead to the *fractional anchor to prey overlap* ***F_o_***. Next, the mean intensity ***I_o_*** from the GFP channel was estimated inside the regions of spot overlap and the weighted average of ***F_o_· I_o_*** was computed over the whole field of view (anchors area used for weighting).

### Analysis of PICT experiments

In the primary selection of positive hits (screening of Fig. 1b), a Fold change was estimated for each condition as the ratio between the weighted average of ***F_o_· I_o_*** after and prior to rapamycin addition (see Fold change values in Supplementary Table 1). An arbitrary Fold change of 3 was established as potential interactions. This primary list was manually curated and confirmed in one-to-one PICT experiments.

We defined an Interaction score as the subtraction of the weighted average of ***F_o_·I_o_*** prior to rapamycin addition from the weighted average of ***F_o_· I_o_*** upon rapamycin addition. In PICT experiments conducted to determine the most efficient baits used to anchor each MTC (Fig. 1a and Supplementary Table 2) and to evaluate the network of interactions between P4-ATPases and MTCs (Fig. 1d and Supplementary Table 3), an arbitrary Interaction score of 100 was established to identify positive interactions. This primary list was manually validated. All one-to-one PICT experiments (Fig. 2c, 5b and 6c) were done in at least two biological replicates. For the statistical analysis, the Interaction score of one-to-one experiments was normalized to the measurement of those strains expressing wild-type Drs2 (Fig. 5b and 6c) or to the measurement of those strains after two hours at 23°C (Fig. 2c).

### Co-localization assays

Similar as described above, the co-localization score ***S_o_*** was determined as the area of overlap between GFP and RFP channels estimated by counting the number of foreground pixels common in both segmentation masks (logical AND operation) over all fields of view (RFP area used for weighting) (Fig. 6b). For the statistical analysis, the co-localization score was estimated in cells harboring wild-type Drs2 or *drs2-*5A and normalized to the measurement of those strains expressing wild-type Drs2.

In Supplementary Video 1 and 2, cells grown in YPD were labelled with 8μM FM4-64. Prior to imaging, cells were kept in the dark for 30 minutes to stain the vacuole. Then cells were washed twice with Low Flo liquid medium to remove YPD and free FM4-64 before imaging. Imaris software was used to obtain the 3D reconstruction of yeast cells imaged with Z-stacks of 250nm step in size.

### Vesicle tracking

The protocol used for the tracking of Atg9 vesicles is based on the method reported previously^22^. Atg9 was C-terminally fused to a 3xGFP tag. The best focal plane was selected for imaging and 2D time-lapses were acquired at 20 frames per second with a 488 nm laser excitation providing a power of approximately 50mW at the back aperture of the objective lens, placed on an inverted microscope (IX71 Olympus) optimized for oblique illumination and fast acquisition (Hamamatsu Flash4 v2 sCMOS camera) with a 100x/1.45 objective lens. Oblique illumination reduces background signal and the high-speed camera allows high-temporal resolution. The axial extension of the laser inclined sheet was approximately 1.5 to 2μm and the usable field of view approximately 40×40μm; images were cropped to increase acquisition speed and illumination was kept continuous during the time-lapses. More than 290 trajectories of GFP puncta were collected from several time-lapses for each strain.

The images were compensated for bleaching by applying the correction method “Exponential fit” from ImageJ “Bleach Correction” plugin. Individual vesicles were then identified and tracked by ImageJ Track-Mate plugin^67^. The tracks were linked with “simple LAP” option, a maximal linking distance of 5 pixels and no gap-closing allowed. Finally, only tracks containing a minimum of 5 spots were kept to discard spurious tracks.

For each time-lapse, the average speed of the vesicles was computed within tracks and binned to a 20bin histogram spanning from 0 to 200 nm/frame. The resulting histograms were fitted to a two-component Gaussian mixture model (GMM). The fits were optimized by the nonlinear least-squares regression (MATLAB). From these models, we estimated the mean speed (and standard deviation) for both populations. The fraction of spots belonging to each population was derived as the fractional areas below each Gaussian component normalized to the area below the fitted curve.

### Electron microscopy (EM) and Correlative Light and Electron microscopy (CLEM)

Cells were grown as described before. High-pressure freezing, freeze-substitution and embedding in EM and CLEM experiments were performed as described in the literature^68^. In brief, yeast cells were pelleted and the yeast paste was loaded into 200μm depth planchettes and high-pressure frozen using the High Pressure Freezing Leica EM ICE. The planchette sandwich was disassembled under liquid nitrogen prior to freeze-substitution. Samples were freeze-substituted using a temperature-controlling AFS2 (Leica) with an FPS robot.

EM pictures were then taken at various magnifications in a JEOL JEM 1011 electron microscope equipped with a Veleta 2k x 2k side-mounted TEM CCD camera (Olympus).

For CLEM experiments, samples were stored protected from light after polymerization and processed further within two days. Ultrathin sections were cut with a diamond knife in a microtome (Leica EM UC7) and picked up on carbon-coated 200 mesh copper grids. Fluorescent fiducial markers, 0.02μm Blue FluoSpheres (Molecular Probes) (excitation 365 nm/emission 415nm) were adhered to the section. Then, grids were placed section-face down onto a 15μl drop of FluoSpheres for 10 min, blotted with filter paper and washed with three drops of water with blotting between the washing steps. To minimize bleaching, fluorescence images were taken with an IX83 (Olympus) during the next three days. From 2 to 5 images were taken in different focal planes for both channels (GFP and blue fluorescent fiducials). During imaging, the positions on the grid of the imaged areas of interest were recorded. EM pictures of recorded positions were taken and correlated to the corresponding GFP images. We detected Ape1-GFP aggregates in 2% of the cells of our sections (n=660).

### Immunoblot analysis

In experiments of Fig. 2a, 3a, 3c, 3d, 3e and 4d cells were grown as described before. In experiment of Fig. 6d, cells were grown overnight at 30°C in synthetic minimal liquid medium lacking Uracil and supplemented with 2% galactose to induce overexpression. Next morning, with an optical density (OD) close to 0.5, cultures were left at 23°C for two hours.

An alkaline extraction method described in Kushnirov, V, 2000^69^ was followed to prepare the samples for western blot analysis. The blocking step was done with 5% milk powder in TBS-0.5% Tween 20 for one hour at room temperature. Incubation with primary anti-Ape1 antibody was done overnight at 4°C (1:4000 antibody dilution in blocking solution). Incubation with secondary anti-rabbit antibody was done for one hour at room temperature (1:2500 antibody dilution in blocking solution).

Western Blot quantification was done with the ImageStudio™Lite software. In the exported data, the background of the prApe1 and mApe1 bands’ signal intensities is automatically corrected from a region right under or above the band. To calculate the % mApe1, the signal intensity of mApe1 multiplied by its band area was divided by the result of the addition of the signal intensities of prApe1 and mApe1, each one multiplied by its corresponding band area.

To calculate the processing of Ape1 and the rescue in the processing of Ape1, the % mApe1 for each strain was calculated and divided by the % mApe1 achieved in wild-type cells.

### Cross-linking and immunoprecipitation

Cells were grown as described before. 600 OD_600_ units of yeast cells were washed twice with B88 buffer (19.97mM HEPES pH=6.8, 149.77mM KOAc, 250.32mM sorbitol, 4mM MgOAc). Cell lysis was performed using Freezer/Mill® Cryogenic Grinder (Spex®SamplePrep, model 6775). Cell powder was resuspended in 5ml B88 + PMSF + protein inhibitor and centrifuged at 3000rpm for 10min to eliminate cell debris. The supernatant was taken and DSSO was added to a final concentration of 2.5mM. After 20 minutes, 0.5M glycine was added to stop the reaction. The mix was incubated for 5min. Triton X-100 was added to a final concentration of 1%. The suspension was incubated with rotation for 1h at 4°C before removing the insoluble components by 13000rpm centrifugation at 4°C during 1h. For the immunoprecipitation, the samples were incubated for 1 h with previously equilibrated Bab agarose beads (Chromotek) at 4°C. Then, they were incubated with GFP-Trap_Agarose (Chromotek) overnight at 4°C. The resin was first washed 4 times with a solution containing 8M urea, 2M NaCl and 1% TX-100 in PBS, and it was subsequently washed 4 more times with PBS.

The resin used in immunoprecipitation was cleaned three times with 500μl of 200mM ammonium bicarbonate and 60μl of 6M urea/200mM ammonium bicarbonate were added. Samples were then reduced with dithiothreitol (30nmol, 37°C, 60 min), alkylated in the dark with iodoacetamide (60nmol, 25°C, 30 min) and diluted to 1M urea with 200mM ammonium bicarbonate for trypsin digestion (1μg, 37°C, 8 h, Promega cat # V5113).

After digestion, peptide mix was acidified with formic acid and desalted with a MicroSpin C18 column (The Nest Group, Inc) prior to LC-MS/MS analysis.

### Chromatographic and mass spectrometric analysis

Samples were analyzed using an LTQ-Orbitrap Velos Pro mass spectrometer (Thermo Fisher Scientific, San Jose, CA, USA) coupled to an EASY-nLC 1000 (Thermo Fisher Scientific (Proxeon), Odense, Denmark). Peptides were loaded onto the 2-cm Nano Trap column with an inner diameter of 100μm packed with C18 particles of 5μm particle size (Thermo Fisher Scientific) and were separated by reversed-phase chromatography using a 25-cm column with an inner diameter of 75μm, packed with 1.9μm C18 particles (Nikkyo Technos Co., Ltd. Japan). Chromatographic gradients started at 93% buffer A and 7% buffer B with a flow rate of 250nl/min for 5 minutes and gradually increased 65% buffer A and 35% buffer B in 60 min. After each analysis, the column was washed for 15 min with 10% buffer A and 90% buffer B. Buffer A: 0.1% formic acid in water. Buffer B: 0.1% formic acid in acetonitrile.

The mass spectrometer was operated in positive ionization mode with nanospray voltage set at 2.1 kV and source temperature at 300°C. Ultramark 1621 was used for external calibration of the FT mass analyzer prior the analyses, and an internal calibration was performed using the background polysiloxane ion signal at m/z 445.1200. The acquisition was performed in data-dependent acquisition (DDA) mode and full MS scans with 1 micro scans at resolution of 60,000 were used over a mass range of m/z 350-2000 with detection in the Orbitrap. Auto gain control (AGC) was set to 1E6, dynamic exclusion (60 seconds) and charge state filtering disqualifying singly charged peptides was activated. In each cycle of DDA analysis, following each survey scan, the top twenty most intense ions with multiple charged ions above a threshold ion count of 5000 were selected for fragmentation. Fragment ion spectra were produced via collision-induced dissociation (CID) at normalized collision energy of 35% and they were acquired in the ion trap mass analyzer. AGC was set to 1E4, isolation window of 2.0 m/z, an activation time of 10 ms and a maximum injection time of 100 ms were used. All data were acquired with Xcalibur software v2.2.

Digested bovine serum albumin (New england biolabs cat # P8108S) was analyzed between each sample to avoid sample carryover and to assure stability of the instrument and QCloud^70^ has been used to control instrument longitudinal performance during the project.

### Mass spectrometric data Analysis

Acquired spectra were analyzed using the Proteome Discoverer software suite (v1.4, Thermo Fisher Scientific) and the Mascot search engine (v2.6, Matrix Science)^71^. The data were searched against a *S. cerevisiae* Genome database (as in November 2019, 6080 entries) plus a list^72^ of common contaminants and all the corresponding decoy entries. For peptide identification, a precursor ion mass tolerance of 7 ppm was used for MS1 level, trypsin was chosen as enzyme and up to three missed cleavages were allowed. The fragment ion mass tolerance was set to 0.5 Da for MS2 spectra. Oxidation of methionine and N-terminal protein acetylation were used as variable modifications whereas carbamidomethylation on cysteines was set as a fixed modification. False discovery rate (FDR) in peptide identification was set to a maximum of 5%. PSM were normalized by the median of Drs2 PSM.

SAINTexpress algorithm was used to score protein-protein interactions^73^.

### Structural modeling

Protein flippases Dnf1, Dnf2, Dnf3 and Neo1 were modelled by using Modeller from the HHSearch server^74^, using Drs2 as a template (PDB ID: 6ROH:A). The ATP8A2 homology model was obtained with Modeller from the HHSearch server^74^, using ATP8A1 as template (PDB ID: 6K7N). To model Drs2 within the membrane we used the data from the MemProtMD database^75^. Images were generated using PyMOL^76^.

### Gene Ontology enrichment analysis

The p-value (*P*) for the Gene Ontology enrichment was calculated using the Hypergeometric distribution corrected by the Bonferroni method of multiple test correction.

## Data availability

Data supporting the findings of this manuscript are available from the corresponding authors upon reasonable request. The raw proteomics data will be deposited to the PRIDE^77^ repository once the manuscript is accepted for publication.

## Acknowledgements

The proteomics analyses were performed in the CRG/UPF Proteomics Unit which is part of the of Proteored, PRB3 and is supported by grant PT17/0019, of the PE I+D+i 2013-2016, funded by ISCIII and ERDF. We thank Josep Vilardell, Bernat Crosas, Amy Curwin, Vivek Malhotra, Michael Knop, Marko Kaksonen, Kenji Maeda and Charles Boone for providing devices, reagents, strains and plasmids. We thank the Advanced Digital Microscopy Core Facility (ADMCF), the Biostatistics/Bioinformatics and the Mass Spectrometry and Proteomics facilities of the IRB Barcelona for live-cell imaging, technical support and also sharing devices with us. We thank Simona Barankova, Radovan Dojcilovic and Daniel Castaño-Díez for fruitful discussions. We thank Raúl Méndez and Ignasi Fita for their support and mentoring during these years. OG was funded by research grants from the Spanish funding agency (MINECO; BFU2012-36385, PGC2018-095745-B-I00 and EUR2019-103815) and supported by the “Unidad de Excelencia María de Maeztu”, funded by the MINECO (ref: MDM-2014-0370). IP was supported by FPI awarded by the Spanish Ministry of Economy and Competitiveness (MINECO) (ref: BES-2013-063945) and EMBO Short-Term Fellowship 7076.

## Author contribution

I.P., T.R.G., M.M., M.H. and O.G. designed the research; I.P., A.C.H., A.G., A.N., C.B., E.A., J.C., M.A., M.P, N.J., S.T. and O.G. conducted the experiments and performed the analysis; and O.G. and I.P. discussed results and wrote the manuscript.

## Additional information

**Supplementary Information** accompanies this paper at:

## References

1. Yu, I.-M. & Hughson, F. M. Tethering factors as organizers of intracellular vesicular traffic. Annu Rev Cell Dev Biol 26, 137–156 (2010).

2. Bröcker, C., Engelbrecht-Vandré, S. & Ungermann, C. Multisubunit tethering complexes and their role in membrane fusion. Curr. Biol. 20, 943–952 (2010).

3. Dubuke, M. L. & Munson, M. The Secret Life of Tethers: The Role of Tethering Factors in SNARE Complex Regulation. Front. Cell Dev. Biol. 4, 1–8 (2016).

4. Suvorova, E. S., Duden, R. & Lupashin, V. V. The Sec34/Sec35p complex, a Ypt1p effector required for retrograde intra-Golgi trafficking, interacts with Golgi SNAREs and COPI vesicle coat proteins. J. Cell Biol. 157, 631–643 (2002).

5. Ren, Y. et al. A Structure-Based Mechanism for Vesicle Capture by the Multisubunit Tethering Complex Dsl1. Cell 139, 1119–1129 (2009).

6. Balderhaar, H. J. k. & Ungermann, C. CORVET and HOPS tethering complexes - coordinators of endosome and lysosome fusion. J. Cell Sci. 126, 1307–1316 (2013).

7. Elizabeth Conibear, Jessica N. Cleck, and T. H. S. Vps51p Mediates the Association of the GARP (Vps52/ 53/54) Complex with the Late Golgi t-SNARE Tlg1p. Mol. Biol. Cell 21, 1610–1623 (2002).

8. Heider, M. R. et al. Subunit connectivity, assembly determinants, and architecture of the yeast exocyst complex. Nat Struct Mol Biol 23, 59–66 (2016).

9. Yu, S. & Liang, Y. A trapper keeper for TRAPP, its structures and functions. Cell. Mol. Life Sci. 69, 3933–3944 (2012).

10. Thomas, L. L., Joiner, A. M. N. & Fromme, J. C. The TRAPPIII complex activates the GTPase Ypt1 (Rab1) in the secretory pathway. J. Cell Biol. 217, 283–298 (2018).

11. Yip, C. K., Berscheminski, J. & Walz, T. Molecular architecture of the TRAPPII complex and implications for vesicle tethering. Nat. Struct. Mol. Biol. 17, 1298–1304 (2010).

12. Morozova, N. et al. TRAPPII subunits are required for the specificity switch of a Ypt-Rab GEF. Nat. Cell Biol. 8, 1263–1269 (2006).

13. Tan, D. et al. The EM structure of the TRAPPIII complex leads to the identification of a requirement for COPII vesicles on the macroautophagy pathway. Proc. Natl. Acad. Sci. U. S. A. 110, 19432–19437 (2013).

14. Gallego, O. et al. Detection and Characterization of Protein Interactions In Vivo by a Simple Live-Cell Imaging Method. PLoS One 8, 1–6 (2013).

15. Torreira, E. et al. The dynamic assembly of distinct RNA polymerase i complexes modulates rDNA transcription. eLife 6, (2017).

16. Zou, S. et al. Trs130 participates in autophagy through GTPases Ypt31/32 in Saccharomyces cerevisiae. Traffic 14, 233–246 (2013).

17. Reggiori, F., Wang, C. W., Stromhaug, P. E., Shintani, T. & Klionsky, D. J. Vps51 is part of the yeast Vps fifty-three tethering complex essential for retrograde traffic from the early endosome and Cvt vesicle completion. J. Biol. Chem. 278, 5009–5020 (2003).

18. Wang, I. H., Chen, Y. J., Hsu, J. W. & Lee, F. J. S. The Arl3 and Arl1 GTPases co-operate with Cog8 to regulate selective autophagy via Atg9 trafficking. Traffic 18, 580–589 (2017).

19. Shirahama-Noda, K., Kira, S., Yoshimori, T. & Noda, T. TRAPPIII is responsible for vesicular transport from early endosomes to Golgi, facilitating Atg9 cycling in autophagy. J. Cell Sci. 126, 4963–4973 (2013).

20. Matoba, K. et al. Atg9 is a lipid scramblase that mediates autophagosomal membrane expansion. Nat. Struct. Mol. Biol. 27, 1185–1193 (2020).

21. Sawa-Makarska, J. et al. Reconstitution of autophagosome nucleation defines Atg9 vesicles as seeds for membrane formation. Science. 369, eaaz7714 (2020).

22. Yamamoto, H. et al. Atg9 vesicles are an important membrane source during early steps of autophagosome formation. J. Cell Biol. 198, 219–233 (2012).

23. Natarajan, P., Wang, J., Hua, Z. & Graham, T. R. Drs2p-coupled aminophospholipid translocase activity in yeast Golgi membranes and relationship to in vivo function. Proc. Natl. Acad. Sci. U. S. A. 101, 10614–10619 (2004).

24. Hanamatsu, H., Fujimura-Kamada, K., Yamamoto, T., Furuta, N. & Tanaka, K. Interaction of the phospholipid flippase Drs2p with the F-box protein Rcy1p plays an important role in early endosome to trans-Golgi network vesicle transport in yeast. J. Biochem. 155, 51–62 (2014).

25. Michael Costanzo, Baryshnikova, A., L Myers, C., Andrews, B. & Boone, C. Charting the genetic interaction map of a cell. Curr. Opin. Biotechnol. 22, 66–74 (2011).

26. Chen, J., Zheng, X. F., Brown, E. J. & Schreiber, S. L. Identification of an 11-kDa FKBP12-rapamycin-binding domain within the 289-kDa FKBP12-rapamycin-associated protein and characterization of a critical serine residue. Proc. Natl. Acad. Sci. U. S. A. 92, 4947–4951 (1995).

27. Protopopov, V., Govindan, B., Novick, P. & Gerst, J. E. Homologs of the synaptobrevin/VAMP family of synaptic vesicle proteins function on the late secretory pathway in S. cerevisiae. Cell 74, 855–861 (1993).

28. Balderhaar, H. J. K. et al. The CORVET complex promotes tethering and fusion of Rab5/Vps21-positive membranes. Proc. Natl. Acad. Sci. U. S. A. 110, 3823–3828 (2013).

29. Zhou, F. et al. A Rab5 GTPase module is important for autophagosome clouser. PLoS Genet. 13, 1–24 (2017).

30. Saito, K. et al. Cdc50p, a protein required for polarized growth, associates with the Drs2p P-type ATPase implicated in phospholipid translocation in Saccharomyces cerevisiae. Mol. Biol. Cell 15, 3418–3432 (2004).

31. Sebastian, T. T., D, B. R., Xu, P. & Graham, T. R. Phospholipid flippases: building asymmetric membranes and transport vesicles. Biochim Biophys Acta 1821, 1068–1077 (2012).

32. Hua, Z., Fatheddin, P. & Graham, T. R. An Essential Subfamily of Drs2p-related P-Type ATPases Is Required for Protein Trafficking between Golgi Complex and Endosomal/Vacuolar System. Mol. Biol. Cell 13, 3162–3177 (2002).

33. Timcenko, M. et al. Structure and autoregulation of a P4-ATPase lipid flippase. Nature 571, 366–370 (2019).

34. Bai, L. et al. Autoinhibition and activation mechanisms of the eukaryotic lipid flippase Drs2p-Cdc50p. Nat. Commun. 10, 1–10 (2019).

35. Lynch-Day, M. A. & Klionsky, D. J. The Cvt pathway as a model for selective autophagy. FEBS Lett. 584, 1359–1366 (2010).

36. Guimaraes, R. S., Delorme-Axford, E., Klionsky, D. J. & Reggiori, F. Assays for the biochemical and ultrastructural measurement of selective and nonselective types of autophagy in the yeast Saccharomyces cerevisiae. Methods 75, 141–150 (2015).

37. Meiling-Wesse, K. et al. Trs85 (Gsg1), a component of the TRAPP complexes, is required for the organization of the preautophagosomal structure during selective autophagy via the Cvt pathway. J. Biol. Chem. 280, 33669–33678 (2005).

38. Chen, C.-Y., Ingram, M. F., Rosal, P. H. & Graham, T. R. Role for Drs2p, a P-Type ATPase and Potential Aminophospholipid Translocase, in Yeast Late Golgi Function. J Cell Biol 147, 1223–1236 (1999).

39. Baba, M., Osumi, M., Scott, S. V., Klionsky, D. J. & Ohsumi, Y. Two distinct pathways for targeting proteins from the cytoplasm to the vacuole/lysosome. J. Cell Biol. 139, 1687–1695 (1997).

40. Kukulski, W. et al. Correlated fluorescence and 3D electron microscopy with high sensitivity and spatial precision. J. Cell Biol. 192, 111–119 (2011).

41. Furuta, N., Fujimura-Kamada, K., Saito, K., Yamamoto, T. & Tanaka, K. Endocytic Recycling in Yeast Is Regulated by Putative Phospholipid Translocases and the Ypt31p/32p–Rcy1p Pathway. Mol. Biol. Cell 18, 295–312 (2007).

42. Liu, K., Surendhran, K., Nothwehr, S. F. & Graham, T. R. P4-ATPase Requirement for AP-1/Clathrin Function in Protein Transport from the trans-Golgi Network and Early Endosomes. Mol. Biol. Cell 19, 3526–3535 (2008).

43. Hinners, I. Changing directions: clathrin-mediated transport between the Golgi and endosomes. J. Cell Sci. 116, 763–771 (2003).

44. Odorizzi, G., Cowles, C. R. & Emr, S. D. The AP-3 complex: a coat of many colours. Trends Cell Biol. 8, 282–288 (1998).

45. Sakane, H., Yamamoto, T. & Tanaka, K. The functional relationship between the Cdc50p-Drs2p putative aminophospholipid translocase and the Arf GAP Gcs1p in vesicle formation in the retrieval pathway from yeast early endosomes to the TGN. Cell Struct. Funct. 31, 87–108 (2006).

46. Xu, P., Baldridge, R. D., Chi, R. J., Burd, C. G. & Graham, T. R. Phosphatidylserine flipping enhances membrane curvature and negative charge required for vesicular transport. J. Cell Biol. 202, 875–886 (2013).

47. Liu, K., Surendhran, K., Nothwehr, S. F. & Graham, T. R. P4-ATPase Requirement for AP-1/Clathrin Function in Protein Transport from the trans-Golgi Network and Early Endosomes. Mol. Biol. Cell 19, 3526–3535 (2008).

48. Alder-Baerens, N., Lisman, Q., Luong, L., Pomorski, T. & C.M, H. J. Loss of P4 ATPases Drs2p and Dnf3p Disrupts Aminophospholipid Transport and Asymmetry in Yeast Post-Golgi Secretory Vesicles. Mol. Biol. Cell 17, 1632–1642 (2006).

49. Choi, H. S., Han, G. S. & Carman, G. M. Phosphorylation of yeast phosphatidylserine synthase by protein kinase A: Identification of SER46 and SER47 as major sites of phosphorylation. J. Biol. Chem. 285, 11526–11536 (2010).

50. Tsai, P.-C., Hsu, J.-W., Liu, Y.-W., Chen, K.-Y. & Lee, F.-J. S. Arl1p regulates spatial membrane organization at the trans-Golgi network through interaction with Arf-GEF Gea2p and flippase Drs2p. Proc. Natl. Acad. Sci. U. S. A. 110, E668–E677 (2013).

51. Chantalat, S. The Arf activator Gea2p and the P-type ATPase Drs2p interact at the Golgi in Saccharomyces cerevisiae. J. Cell Sci. 117, 711–722 (2004).

52. Liu, K., Hua, Z., Nepute, J. A. & Graham, T. R. Yeast P4-ATPases Drs2p and Dnf1p Are Essential Cargos of the NPFXD/Sla1p Endocytic Pathway. Mol. Biol. Cell 18, 487–500 (2007).

53. Kao, A. et al. Development of a Novel Cross-linking Strategy for Fast and Accurate Identification of Cross-linked Peptides of Protein Complexes. Mol. Cell. Proteomics 10, M110.002212 (2011).

54. Picco, A. et al. The In Vivo Architecture of the Exocyst Provides Structural Basis for Exocytosis. Cell 168, 400–412.e18 (2017).

55. Kakuta, S. et al. Atg9 vesicles recruit vesicle-tethering proteins Trs85 and Ypt1 to the autophagosome formation site. J. Biol. Chem. 287, 44261–44269 (2012).

56. Galindo, A., Planelles-Herrero, V. J., Degliesposti, G. & Munro, S. Cryo-EM structure of metazoan TRAPPIII, the multi-subunit complex that activates the GTPase Rab1. EMBO J. 40, 1–17 (2021).

57. Joiner, A. M. et al. Structural basis of TRAPPIII-mediated Rab1 activation. EMBO J. 40, 1–19 (2021).

58. Hiraizumi, M., Yamashita, K., Nishizawa, T. & Nureki, O. Cryo-EM structures capture the transport cycle of the P4-ATPase flippase. Science. 365, 1149–1155 (2019).

59. Nakanishi, H. et al. Transport Cycle of Plasma Membrane Flippase ATP11C by Cryo-EM. Cell Rep. 32, 108208 (2020).

60. Nakanishi, H. et al. Crystal structure of a human plasma membrane phospholipid flippase. J. Biol. Chem. 295, 10180–10194 (2020).

61. Gall, W. E. et al. Drs2p-dependent formation of exocytic clathrin-coated vesicles in vivo. Curr. Biol. 12, 1623–1627 (2002).

62. Lee, S. et al. Transport through recycling endosomes requires EHD1 recruitment by a phosphatidylserine translocase. EMBO J. 34, 669–688 (2015).

63. Roland, B. P. & Graham, T. R. Decoding P4-ATPase Substrate Interactions. Crit Rev Biochem Mol Biol 51, 513–527 (2016).

64. Janke, C. et al. A versatile toolbox for PCR-based tagging of yeast genes: New fluorescent proteins, more markers and promoter substitution cassettes. Yeast 21, 947–962 (2004).

65. Tong, Y. & Boone, C. Synthetic Genetic Array Analysis in Saccharomyces cerevisiae. Methods Mol. Biol. 313, 171–192 (2005).

66. Huh et al. Global analysis of protein localization in budding yeast. Nature 425, 686–691 (2003).

67. Tinevez, J. Y. et al. TrackMate: An open and extensible platform for single-particle tracking. Methods 115, 80–90 (2017).

68. Müller-Reichert, M. Verkade P. Correlative Light and Electron Microscopy. 1st edition. Methods in Cell Biol. (2012).

69. Kushnirov, V. V. Rapid and reliable protein extraction from yeast. Yeast 16, 857–860 (2000).

70. Chiva, C. et al. QCloud: A cloud-based quality control system for mass spectrometry-based proteomics laboratories. PLoS One 13, 1–14 (2018).

71. Perkins, D. N., Pappin, D. J., Creasy, D. M. & Cottrell, J. S. Probability-based protein identification by searching sequence databases using mass spectrometry data. Electrophoresis 20, 3551–3567 (1999).

72. Beer, L. A., Liu, P., Ky, B., Barnhart, K. T. & Speicher, D. W. Efficient quantitative comparisons of plasma proteomes using label-free analysis with MaxQuant. Methods Mol Biol 1619, 339–352 (2017).

73. Guoci Teo et al. SAINTexpress: improvements and additional features in Significance Analysis of Interactome software. J Proteomics 100, 37–43 (2014).

74. Zimmermann, L. et al. A Completely Reimplemented MPI Bioinformatics Toolkit with a New HHpred Server at its Core. J. Mol. Biol. 430, 2237–2243 (2018).

75. Newport, T. D., Sansom, M. S. P. & Stansfeld, P. J. The MemProtMD database: A resource for membrane-embedded protein structures and their lipid interactions. Nucleic Acids Res. 47, D390–D397 (2019).

76. The PyMOL Molecular Graphics System, Version 2.3.4 Schrödinger, LLC.

77. Vizcaíno, J. A. et al. 2016 update of the PRIDE database and its related tools. Nucleic Acids Res. 44, D447–D456 (2016).

