## Supplementary material for "Drs2 regulates TRAPPIII in Atg9 transport: exposing the interplay of P4-ATPases and Multisubunit Tethering Complexes": Supplementary Information.pdf

#### Supplementary figures:

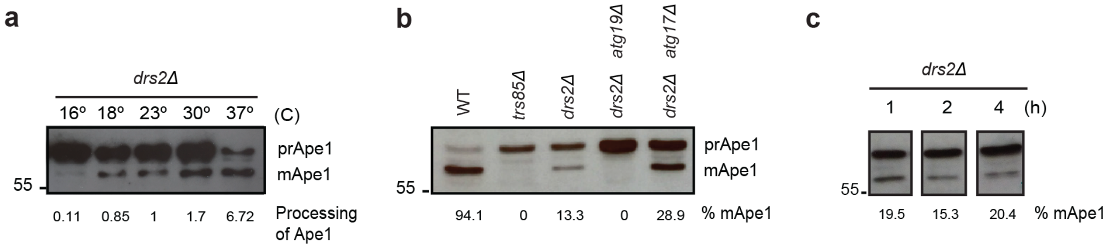


**Supplementary Fig. 1** Drs2 is essential for the CVT pathway as the temperature decreases. **a-c** Analysis of Ape1 processing by western blot in the indicated strains. **a** *drs2Δ* cells were grown at 30ºC and then cultured for 2 hours at the indicated temperatures. Below, processing of Ape1 normalized to the processing achieved at 23ºC. **b** Analysis of Ape1-processing in *drs2Δ* cells where either the CVT pathway (*atg19Δ*) or non-selective autophagy (*atg17Δ*) were blocked. **c** *drs2Δ* cells were grown at 30ºC and then cultured for 1, 2 or 4 hours at 23ºC. **b, c** Below, percentage of mApe1.

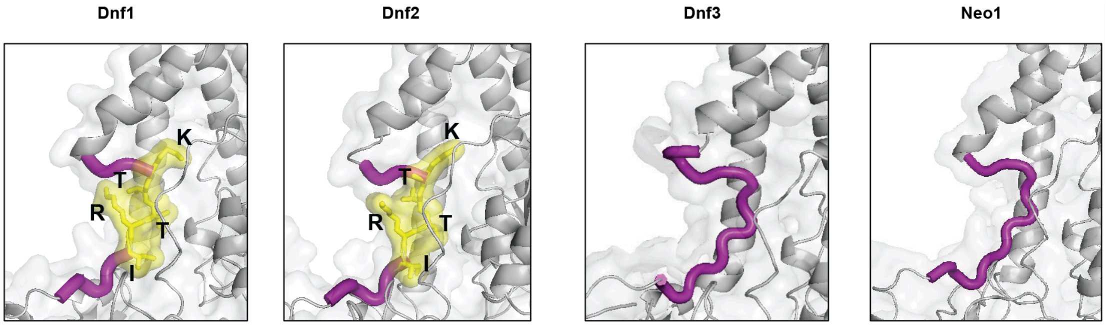


**Supplementary Fig. 2** Modelling of the N-terminal cavity of P4-ATPases. Structural representation of the 15 aa cavity (purple) in the N-terminal tail of Dnf1 (15.5x9.4 Å), Dnf2 (11.6x9.5 Å), Dnf3 (13.5x15.5 Å) and Neo1 (18.1x13.8 Å). Drs2 was used as a template (PDB ID: 6ROH:A). The I(S/R)TTK motif present in Dnf1 and Dnf2 is highlighted in yellow.

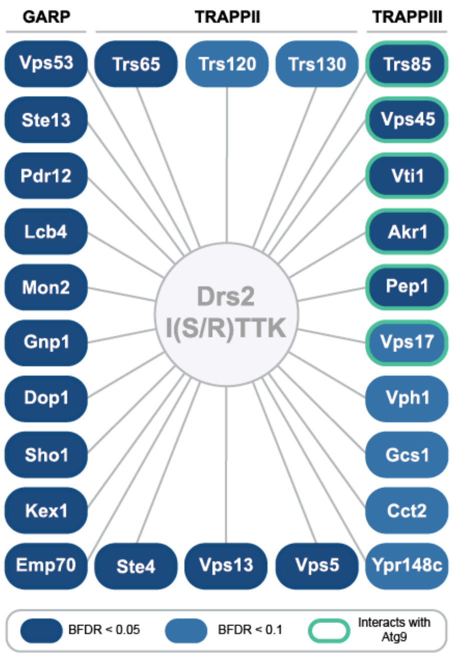


**Supplementary Fig. 3** Graphic representation of the 26 proteins whose interaction with Drs2 is diminished by the mutation of the I(S/R)TTK motif. Binding to Atg9 as reported in https://www.yeastgenome.org in April 2021. BFDR, Bayesian False Discovery Rate.

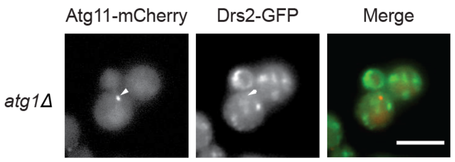


**Supplementary Fig. 4** Co-localization assay of GFP-tagged Drs2 and the PAS marker Atg11 tagged to mCherry in cells lacking the kinase Atg1, required for the CVT vesicle formation. Arrowhead points to Atg11-mCherry spot. Scale bar, 5$\mu$m.

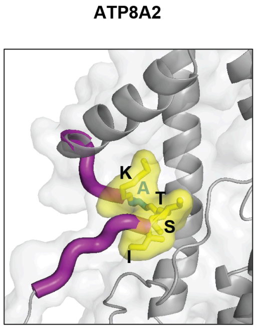


**Supplementary Fig. 5** Modelling of the N-terminal cavity of ATP8A2. Structural representation of the 15 aa cavity (purple) in the N-terminal tail of the human ATP8A2 (17.7x20.5 Å). ATP8A1 was used as a template (PDB ID: 6K7N). The ISTAK motif is highlighted in yellow. The only mismatch with the I(S/R)TTK motif is labeled in blue.

### Supplementary tables:

**Supplementary Table 1.** Yeast strains and PICT assay Fold change to determine protein-protein interactions relevant for the function of MTCs. Strains were constructed with the SGA approach to express the anchor Tub4-RFP-FKBP, MTCs subunits as baits (FRB fusions) and preys selected based on genetic interactions with MTCs (GFP fusions at the C-terminus). Cells were imaged before and after adding rapamycin at 200ms exposure time for the RFP channel, and both 200ms and 1500ms for the GFP channel. A fold change was calculated as the ratio of co-localization between the green and red channels upon and prior to rapamycin addition (see Methods for a more detailed description). Those samples with a fold change higher than 3 were considered as potential interactions and manually validated.

**Supplementary Table 2.** Yeast strains and results of the PICT assay to determine the most efficient bait to anchor each MTC. Strains were constructed with the SGA approach to express the anchor Tub4-RFP-FKBP and all possible pairwise combinations of MTCs subunits as baits (FRB fusions) and preys (GFP fusions at the C-terminus). Cells were imaged before and after adding rapamycin at 200ms exposure time for both RFP and GFP channels. The Interaction score was calculated as described in Methods. Those samples with the highest Interaction score were used to select the bait that recruits the corresponding MTC to the anchoring platform more efficiently.

**Supplementary Table 3.** Yeast strains and results of the PICT assay to determine the network of interactions between MTCs and P4-ATPases. Strains were constructed with the SGA approach to express the anchor Tub4-RFP-FKBP and pairwise combinations of MTCs subunits as baits (FRB fusions) and P4-ATPses as preys (GFP fusions at the C-terminus). Cells were imaged before and after adding rapamycin at 200ms exposure time for the RFP channel and 1500ms exposure time for the GFP channel. An Interaction score was calculated as described in Methods. Those samples with an arbitrary Interaction score of 100 were consider as potential interactions and manually validated.

**Supplementary Table 4.** Yeast strains generated by transformation of plasmids and/or homologous recombination following standard PCR strategies.

**Supplementary Table 5.** List of the plasmids used in this article.

### Supplementary videos:

**Supplementary Video 1.** Analysis of cytosolic Ape1 oligomers. 3D reconstruction of a *drs2*Δ single cell with GFP-tagged Ape1 (green) and the vacuole stained with the lipophilic dye FM4-64 (red). Z-stacks of GFP and RFP pictures were taken in 250nm incremental steps.

**Supplementary Video 2.** Analysis of cytosolic Ape1 oligomers. 3D reconstruction of a *drs2-5A* single cell with GFP-tagged Ape1 (green) and the vacuole stained with the lipophilic dye FM4-64 (red). Z-stacks of GFP and RFP pictures were taken in 250nm incremental steps

**Supplementary Video 3.** Atg9 puncta in wild-type cells. 2D time-lapse sequences of endogenous GFP tagged Atg9 were taken at 20 frames per second with a 488 nm laser excitation in wild-type cells. Number on the left top indicates number of frame. Scale bar, 5µm.

**Supplementary Video 4.** Atg9 puncta in *drs2*Δ cells. 2D time-lapse sequences of endogenous GFP tagged Atg9 were taken at 20 frames per second with a 488 nm laser excitation in *drs2*Δ cells. Number on the left top indicates number of frame. Scale bar, 5µm.

**Supplementary Video 5.** Atg9 puncta in *drs2*Δ cells harboring a *drs2*-5A plasmid. 2D time-lapse sequences of endogenous GFP tagged Atg9 were taken at 20 frames per second with a 488 nm laser excitation in *drs2*Δ cells harboring a *drs2*-5A plasmid. Number on the left top indicates number of frame. Scale bar, 5µm.
