## Supplementary material for "Drs2 regulates TRAPPIII in Atg9 transport: exposing the interplay of P4-ATPases and Multisubunit Tethering Complexes": Supplementary Table 1.pdf

| Yeast strains and PICT assay Fold change to determine protein-protein interactions relevant for the function of MTCs. | | | | |
|---|---|---|---|---|
| **Parental strain** | **Genotype** | **Source** | | |
| OGY0003 | *MATα, his3Δ1 leu2Δ0 ura3Δ0 LYS+, can1Δ::STE2pr-LEU2, lyp1Δ::* | Charlie Boone | | |
| MKY2127 | *MATα, his3Δ1 leu2Δ0 ura3Δ0 LYS+, can1Δ::STE2pr-LEU2, lyp1Δ::, tor1-1* | This study | | |
| MKY2128 | *MATα, his3Δ1 leu2Δ0 ura3Δ0 LYS+, can1Δ::STE2pr-LEU2, lyp1Δ::, tor1-1, fpr1Δ::klURA* | This study | | |
| OGY0307 | *MATα, his3Δ1 leu2Δ0 ura3Δ0 LYS+, can1Δ::STE2pr-LEU2, lyp1Δ::, tor1-1, fpr1Δ::klURA, Tub4-(6)-RFP-(24)-FKBP::natNT2* | This study | | |
| OGY0314 | *MATα, his3Δ1 leu2Δ0 ura3Δ0 LYS+, can1Δ::STE2pr-LEU2, lyp1Δ::, tor1-1, fpr1Δ::klURA, Tub4-(6)-RFP-(24)-FKBP::natNT2, DSL1-FRB Hph* | This study | | |
| OGY0315 | *MATα, his3Δ1 leu2Δ0 ura3Δ0 LYS+, can1Δ::STE2pr-LEU2, lyp1Δ::, tor1-1, fpr1Δ::klURA, Tub4-(6)-RFP-(24)-FKBP::natNT2, COG6-FRB Hph* | This study | | |
| OGY0524 | *MATα, his3Δ1 leu2Δ0 ura3Δ0 LYS+, can1Δ::STE2pr-LEU2, lyp1Δ::, tor1-1, fpr1Δ::klURA, Tub4-(6)-RFP-(24)-FKBP::natNT2, VPS3-FRB Hph* | This study | | |
| OGY0317 | *MATα, his3Δ1 leu2Δ0 ura3Δ0 LYS+, can1Δ::STE2pr-LEU2, lyp1Δ::, tor1-1, fpr1Δ::klURA, Tub4-(6)-RFP-(24)-FKBP::natNT2, VAM6-FRB Hph* | This study | | |
| OGY0318 | *MATα, his3Δ1 leu2Δ0 ura3Δ0 LYS+, can1Δ::STE2pr-LEU2, lyp1Δ::, tor1-1, fpr1Δ::klURA, Tub4-(6)-RFP-(24)-FKBP::natNT2, TRS23-FRB Hph* | This study | | |
| OGY0536 | *MATα, his3Δ1 leu2Δ0 ura3Δ0 LYS+, can1Δ::STE2pr-LEU2, lyp1Δ::, tor1-1, fpr1Δ::klURA, Tub4-(6)-RFP-(24)-FKBP::natNT2, TRS130-FRB Hph* | This study | | |
| OGY0320 | *MATα, his3Δ1 leu2Δ0 ura3Δ0 LYS+, can1Δ::STE2pr-LEU2, lyp1Δ::, tor1-1, fpr1Δ::klURA, Tub4-(6)-RFP-(24)-FKBP::natNT2, TRS85-FRB Hph* | This study | | |
| OGY0321 | *MATα, his3Δ1 leu2Δ0 ura3Δ0 LYS+, can1Δ::STE2pr-LEU2, lyp1Δ::, tor1-1, fpr1Δ::klURA, Tub4-(6)-RFP-(24)-FKBP::natNT2, VPS53-FRB Hph* | This study | | |
| OGY0326 | *MATα, his3Δ1 leu2Δ0 ura3Δ0 LYS+, can1Δ::STE2pr-LEU2, lyp1Δ::, tor1-1, fpr1Δ::klURA, Tub4-(6)-RFP-(24)-FKBP::natNT2, SEC3-FRB Hph* | This study | | |
| **Strain** | **Genotype** | **Source** | **Fold change 200 ms** | **Fold change 1500 ms** |
| OGYSGA1641 | *MATα, his3Δ1, leu2Δ0, ura3Δ0, LYS+, can1Δ::STE2pr-LEU2, lyp1Δ::, tor1-1, fpr1Δ::klURA, Tub4-(6)-RFP-(24)-FKBP::natNT2, DSL1-FRB::hphNT1, YKL033W-GFP::HIS3* | This study | 0 | 0 |
| OGYSGA1642 | *MATα, his3Δ1, leu2Δ0, ura3Δ0, LYS+, can1Δ::STE2pr-LEU2, lyp1Δ::, tor1-1, fpr1Δ::klURA, Tub4-(6)-RFP-(24)-FKBP::natNT2, DSL1-FRB::hphNT1, YNL287W-GFP::HIS3* | This study | 3.25 | 0 |
| OGYSGA1643 | *MATα, his3Δ1, leu2Δ0, ura3Δ0, LYS+, can1Δ::STE2pr-LEU2, lyp1Δ::, tor1-1, fpr1Δ::klURA, Tub4-(6)-RFP-(24)-FKBP::natNT2, DSL1-FRB::hphNT1, YGL054C-GFP::HIS3* | This study | 2.3 | 0.14 |
| OGYSGA1644 | *MATα, his3Δ1, leu2Δ0, ura3Δ0, LYS+, can1Δ::STE2pr-LEU2, lyp1Δ::, tor1-1, fpr1Δ::klURA, Tub4-(6)-RFP-(24)-FKBP::natNT2, DSL1-FRB::hphNT1, YMR224C-GFP::HIS3* | This study | 0 | 0 |
| OGYSGA1645 | *MATα, his3Δ1, leu2Δ0, ura3Δ0, LYS+, can1Δ::STE2pr-LEU2, lyp1Δ::, tor1-1, fpr1Δ::klURA, Tub4-(6)-RFP-(24)-FKBP::natNT2, COG6-FRB::hphNT1, YHL020C-GFP::HIS3* | This study | 1.2 | 1.02 |
| OGYSGA1646 | *MATα, his3Δ1, leu2Δ0, ura3Δ0, LYS+, can1Δ::STE2pr-LEU2, lyp1Δ::, tor1-1, fpr1Δ::klURA, Tub4-(6)-RFP-(24)-FKBP::natNT2, COG6-FRB::hphNT1, YPR051W-GFP::HIS3* | This study | 0 | 0 |
| OGYSGA1647 | *MATα, his3Δ1, leu2Δ0, ura3Δ0, LYS+, can1Δ::STE2pr-LEU2, lyp1Δ::, tor1-1, fpr1Δ::klURA, Tub4-(6)-RFP-(24)-FKBP::natNT2, COG6-FRB::hphNT1, YJL123C-GFP::HIS3* | This study | 1.21 | 2.15 |
| OGYSGA1648 | *MATα, his3Δ1, leu2Δ0, ura3Δ0, LYS+, can1Δ::STE2pr-LEU2, lyp1Δ::, tor1-1, fpr1Δ::klURA, Tub4-(6)-RFP-(24)-FKBP::natNT2, COG6-FRB::hphNT1, YFR051C-GFP::HIS3* | This study | 3.02 | 5.75 |
| OGYSGA1649 | *MATα, his3Δ1, leu2Δ0, ura3Δ0, LYS+, can1Δ::STE2pr-LEU2, lyp1Δ::, tor1-1, fpr1Δ::klURA, Tub4-(6)-RFP-(24)-FKBP::natNT2, COG6-FRB::hphNT1, YDL058W-GFP::HIS3* | This study | 0.84 | 1.7 |
| OGYSGA1650 | *MATα, his3Δ1, leu2Δ0, ura3Δ0, LYS+, can1Δ::STE2pr-LEU2, lyp1Δ::, tor1-1, fpr1Δ::klURA, Tub4-(6)-RFP-(24)-FKBP::natNT2, COG6-FRB::hphNT1, YGL137W-GFP::HIS3* | This study | 2.32 | 0.8 |
| OGYSGA1651 | *MATα, his3Δ1, leu2Δ0, ura3Δ0, LYS+, can1Δ::STE2pr-LEU2, lyp1Δ::, tor1-1, fpr1Δ::klURA, Tub4-(6)-RFP-(24)-FKBP::natNT2, COG6-FRB::hphNT1, YGL020C-GFP::HIS3* | This study | 0.82 | 1.12 |
| OGYSGA1652 | *MATα, his3Δ1, leu2Δ0, ura3Δ0, LYS+, can1Δ::STE2pr-LEU2, lyp1Δ::, tor1-1, fpr1Δ::klURA, Tub4-(6)-RFP-(24)-FKBP::natNT2, COG6-FRB::hphNT1, YJR032W-GFP::HIS3* | This study | 0 | 0 |
| OGYSGA1653 | *MATα, his3Δ1, leu2Δ0, ura3Δ0, LYS+, can1Δ::STE2pr-LEU2, lyp1Δ::, tor1-1, fpr1Δ::klURA, Tub4-(6)-RFP-(24)-FKBP::natNT2, COG6-FRB::hphNT1, YER083C-GFP::HIS3* | This study | 0.66 | 0.8 |
| OGYSGA1654 | *MATα, his3Δ1, leu2Δ0, ura3Δ0, LYS+, can1Δ::STE2pr-LEU2, lyp1Δ::, tor1-1, fpr1Δ::klURA, Tub4-(6)-RFP-(24)-FKBP::natNT2, COG6-FRB::hphNT1, YGR187C-GFP::HIS3* | This study | 0 | 0 |
| OGYSGA1655 | *MATα, his3Δ1, leu2Δ0, ura3Δ0, LYS+, can1Δ::STE2pr-LEU2, lyp1Δ::, tor1-1, fpr1Δ::klURA, Tub4-(6)-RFP-(24)-FKBP::natNT2, COG6-FRB::hphNT1, YIL109C-GFP::HIS3* | This study | 0.52 | 0.56 |
| OGYSGA1656 | *MATα, his3Δ1, leu2Δ0, ura3Δ0, LYS+, can1Δ::STE2pr-LEU2, lyp1Δ::, tor1-1, fpr1Δ::klURA, Tub4-(6)-RFP-(24)-FKBP::natNT2, COG6-FRB::hphNT1, YJL004C-GFP::HIS3* | This study | 1.51 | 6.7 |
| OGYSGA1657 | *MATα, his3Δ1, leu2Δ0, ura3Δ0, LYS+, can1Δ::STE2pr-LEU2, lyp1Δ::, tor1-1, fpr1Δ::klURA, Tub4-(6)-RFP-(24)-FKBP::natNT2, COG6-FRB::hphNT1, YGL054C-GFP::HIS3* | This study | 1.46 | 0 |
| OGYSGA1658 | *MATα, his3Δ1, leu2Δ0, ura3Δ0, LYS+, can1Δ::STE2pr-LEU2, lyp1Δ::, tor1-1, fpr1Δ::klURA, Tub4-(6)-RFP-(24)-FKBP::natNT2, COG6-FRB::hphNT1, YDL100C-GFP::HIS3* | This study | 0 | 0 |
| OGYSGA1659 | *MATα, his3Δ1, leu2Δ0, ura3Δ0, LYS+, can1Δ::STE2pr-LEU2, lyp1Δ::, tor1-1, fpr1Δ::klURA, Tub4-(6)-RFP-(24)-FKBP::natNT2, COG6-FRB::hphNT1, YPL051W-GFP::HIS3* | This study | 19.38 | 11.04 |
| OGYSGA1660 | *MATα, his3Δ1, leu2Δ0, ura3Δ0, LYS+, can1Δ::STE2pr-LEU2, lyp1Δ::, tor1-1, fpr1Δ::klURA, Tub4-(6)-RFP-(24)-FKBP::natNT2, COG6-FRB::hphNT1, YMR292W-GFP::HIS3* | This study | 0.88 | 0.87 |
| OGYSGA1661 | *MATα, his3Δ1, leu2Δ0, ura3Δ0, LYS+, can1Δ::STE2pr-LEU2, lyp1Δ::, tor1-1, fpr1Δ::klURA, Tub4-(6)-RFP-(24)-FKBP::natNT2, COG6-FRB::hphNT1, YOR216C-GFP::HIS3* | This study | 7.75 | 3.18 |
| OGYSGA1662 | *MATα, his3Δ1, leu2Δ0, ura3Δ0, LYS+, can1Δ::STE2pr-LEU2, lyp1Δ::, tor1-1, fpr1Δ::klURA, Tub4-(6)-RFP-(24)-FKBP::natNT2, COG6-FRB::hphNT1, YJR060W-GFP::HIS3* | This study | 0.98 | 0.52 |
| OGYSGA1663 | *MATα, his3Δ1, leu2Δ0, ura3Δ0, LYS+, can1Δ::STE2pr-LEU2, lyp1Δ::, tor1-1, fpr1Δ::klURA, Tub4-(6)-RFP-(24)-FKBP::natNT2, COG6-FRB::hphNT1, YBR164C-GFP::HIS3* | This study | 5.59 | 2.82 |
| OGYSGA1664 | *MATα, his3Δ1, leu2Δ0, ura3Δ0, LYS+, can1Δ::STE2pr-LEU2, lyp1Δ::, tor1-1, fpr1Δ::klURA, Tub4-(6)-RFP-(24)-FKBP::natNT2, COG6-FRB::hphNT1, YOR070C-GFP::HIS3* | This study | 0.82 | 2.09 |

| | | | | |
|---|---|---|---|---|
| OGYSGA1665 | *MATa, his3Δ1, leu2Δ0, ura3Δ0, LYS+, can1Δ::STE2pr-LEU2, lyp1Δ::, tor1-1, fpr1Δ::klURA, Tub4-(6)-RFP-(24)-FKBP::natNT2, COG6-FRB::hphNT1, YGR261C-GFP::HIS3* | This study | 1.36 | 2.55 |
| OGYSGA1666 | *MATa, his3Δ1, leu2Δ0, ura3Δ0, LYS+, can1Δ::STE2pr-LEU2, lyp1Δ::, tor1-1, fpr1Δ::klURA, Tub4-(6)-RFP-(24)-FKBP::natNT2, COG6-FRB::hphNT1, YDR137W-GFP::HIS3* | This study | 2.59 | 1.66 |
| OGYSGA1667 | *MATa, his3Δ1, leu2Δ0, ura3Δ0, LYS+, can1Δ::STE2pr-LEU2, lyp1Δ::, tor1-1, fpr1Δ::klURA, Tub4-(6)-RFP-(24)-FKBP::natNT2, COG6-FRB::hphNT1, YCR053W-GFP::HIS3* | This study | 0 | 0 |
| OGYSGA1668 | *MATa, his3Δ1, leu2Δ0, ura3Δ0, LYS+, can1Δ::STE2pr-LEU2, lyp1Δ::, tor1-1, fpr1Δ::klURA, Tub4-(6)-RFP-(24)-FKBP::natNT2, COG6-FRB::hphNT1, YAL019W-GFP::HIS3* | This study | 0.17 | 1.55 |
| OGYSGA1669 | *MATa, his3Δ1, leu2Δ0, ura3Δ0, LYS+, can1Δ::STE2pr-LEU2, lyp1Δ::, tor1-1, fpr1Δ::klURA, Tub4-(6)-RFP-(24)-FKBP::natNT2, COG6-FRB::hphNT1, YBL102W-GFP::HIS3* | This study | 0 | 32.68 |
| OGYSGA1670 | *MATa, his3Δ1, leu2Δ0, ura3Δ0, LYS+, can1Δ::STE2pr-LEU2, lyp1Δ::, tor1-1, fpr1Δ::klURA, Tub4-(6)-RFP-(24)-FKBP::natNT2, COG6-FRB::hphNT1, YDR153C-GFP::HIS3* | This study | 1.49 | 1.53 |
| OGYSGA1671 | *MATa, his3Δ1, leu2Δ0, ura3Δ0, LYS+, can1Δ::STE2pr-LEU2, lyp1Δ::, tor1-1, fpr1Δ::klURA, Tub4-(6)-RFP-(24)-FKBP::natNT2, COG6-FRB::hphNT1, YLR360W-GFP::HIS3* | This study | 0 | 0 |
| OGYSGA1672 | *MATa, his3Δ1, leu2Δ0, ura3Δ0, LYS+, can1Δ::STE2pr-LEU2, lyp1Δ::, tor1-1, fpr1Δ::klURA, Tub4-(6)-RFP-(24)-FKBP::natNT2, COG6-FRB::hphNT1, YPL195W-GFP::HIS3* | This study | 0.68 | 2.94 |
| OGYSGA1673 | *MATa, his3Δ1, leu2Δ0, ura3Δ0, LYS+, can1Δ::STE2pr-LEU2, lyp1Δ::, tor1-1, fpr1Δ::klURA, Tub4-(6)-RFP-(24)-FKBP::natNT2, COG6-FRB::hphNT1, YER074W-GFP::HIS3* | This study | 0.41 | 0 |
| OGYSGA1674 | *MATa, his3Δ1, leu2Δ0, ura3Δ0, LYS+, can1Δ::STE2pr-LEU2, lyp1Δ::, tor1-1, fpr1Δ::klURA, Tub4-(6)-RFP-(24)-FKBP::natNT2, COG6-FRB::hphNT1, YOR132W-GFP::HIS3* | This study | 0 | 1.28 |
| OGYSGA1675 | *MATa, his3Δ1, leu2Δ0, ura3Δ0, LYS+, can1Δ::STE2pr-LEU2, lyp1Δ::, tor1-1, fpr1Δ::klURA, Tub4-(6)-RFP-(24)-FKBP::natNT2, COG6-FRB::hphNT1, YNL297C-GFP::HIS3* | This study | 2.88 | 3.65 |
| OGYSGA1676 | *MATa, his3Δ1, leu2Δ0, ura3Δ0, LYS+, can1Δ::STE2pr-LEU2, lyp1Δ::, tor1-1, fpr1Δ::klURA, Tub4-(6)-RFP-(24)-FKBP::natNT2, COG6-FRB::hphNT1, YMR272C-GFP::HIS3* | This study | 1.68 | 1.01 |
| OGYSGA1677 | *MATa, his3Δ1, leu2Δ0, ura3Δ0, LYS+, can1Δ::STE2pr-LEU2, lyp1Δ::, tor1-1, fpr1Δ::klURA, Tub4-(6)-RFP-(24)-FKBP::natNT2, COG6-FRB::hphNT1, YKR094C-GFP::HIS3* | This study | 0.24 | 0 |
| OGYSGA1678 | *MATa, his3Δ1, leu2Δ0, ura3Δ0, LYS+, can1Δ::STE2pr-LEU2, lyp1Δ::, tor1-1, fpr1Δ::klURA, Tub4-(6)-RFP-(24)-FKBP::natNT2, COG6-FRB::hphNT1, YPL120W-GFP::HIS3* | This study | 1.25 | 0.53 |
| OGYSGA1679 | *MATa, his3Δ1, leu2Δ0, ura3Δ0, LYS+, can1Δ::STE2pr-LEU2, lyp1Δ::, tor1-1, fpr1Δ::klURA, Tub4-(6)-RFP-(24)-FKBP::natNT2, COG6-FRB::hphNT1, YLR039C-GFP::HIS3* | This study | 1.44 | 7.3 |
| OGYSGA1680 | *MATa, his3Δ1, leu2Δ0, ura3Δ0, LYS+, can1Δ::STE2pr-LEU2, lyp1Δ::, tor1-1, fpr1Δ::klURA, Tub4-(6)-RFP-(24)-FKBP::natNT2, COG6-FRB::hphNT1, YDL115C-GFP::HIS3* | This study | 0 | 0 |
| OGYSGA1681 | *MATa, his3Δ1, leu2Δ0, ura3Δ0, LYS+, can1Δ::STE2pr-LEU2, lyp1Δ::, tor1-1, fpr1Δ::klURA, Tub4-(6)-RFP-(24)-FKBP::natNT2, COG6-FRB::hphNT1, YHR012W-GFP::HIS3* | This study | 0 | 3.76 |
| OGYSGA1682 | *MATa, his3Δ1, leu2Δ0, ura3Δ0, LYS+, can1Δ::STE2pr-LEU2, lyp1Δ::, tor1-1, fpr1Δ::klURA, Tub4-(6)-RFP-(24)-FKBP::natNT2, COG6-FRB::hphNT1, YGL225W-GFP::HIS3* | This study | 1.85 | 0.41 |
| OGYSGA1683 | *MATa, his3Δ1, leu2Δ0, ura3Δ0, LYS+, can1Δ::STE2pr-LEU2, lyp1Δ::, tor1-1, fpr1Δ::klURA, Tub4-(6)-RFP-(24)-FKBP::natNT2, COG6-FRB::hphNT1, YOL009C-GFP::HIS3* | This study | 9.78 | 6.85 |
| OGYSGA1684 | *MATa, his3Δ1, leu2Δ0, ura3Δ0, LYS+, can1Δ::STE2pr-LEU2, lyp1Δ::, tor1-1, fpr1Δ::klURA, Tub4-(6)-RFP-(24)-FKBP::natNT2, COG6-FRB::hphNT1, YDR310C-GFP::HIS3* | This study | 1.22 | 1.33 |
| OGYSGA1685 | *MATa, his3Δ1, leu2Δ0, ura3Δ0, LYS+, can1Δ::STE2pr-LEU2, lyp1Δ::, tor1-1, fpr1Δ::klURA, Tub4-(6)-RFP-(24)-FKBP::natNT2, COG6-FRB::hphNT1, YIL160C-GFP::HIS3* | This study | 0 | 0 |
| OGYSGA1686 | *MATa, his3Δ1, leu2Δ0, ura3Δ0, LYS+, can1Δ::STE2pr-LEU2, lyp1Δ::, tor1-1, fpr1Δ::klURA, Tub4-(6)-RFP-(24)-FKBP::natNT2, COG6-FRB::hphNT1, YIL090W-GFP::HIS3* | This study | 1.22 | 0.48 |
| OGYSGA1687 | *MATa, his3Δ1, leu2Δ0, ura3Δ0, LYS+, can1Δ::STE2pr-LEU2, lyp1Δ::, tor1-1, fpr1Δ::klURA, Tub4-(6)-RFP-(24)-FKBP::natNT2, COG6-FRB::hphNT1, YDR202C-GFP::HIS3* | This study | 0 | 3.09 |
| OGYSGA1688 | *MATa, his3Δ1, leu2Δ0, ura3Δ0, LYS+, can1Δ::STE2pr-LEU2, lyp1Δ::, tor1-1, fpr1Δ::klURA, Tub4-(6)-RFP-(24)-FKBP::natNT2, COG6-FRB::hphNT1,, YAL011W-GFP::HIS3* | This study | 0.29 | 0.06 |
| OGYSGA1689 | *MATa, his3Δ1, leu2Δ0, ura3Δ0, LYS+, can1Δ::STE2pr-LEU2, lyp1Δ::, tor1-1, fpr1Δ::klURA, Tub4-(6)-RFP-(24)-FKBP::natNT2, COG6-FRB::hphNT1, YDR293C-GFP::HIS3* | This study | 0 | 0 |
| OGYSGA1690 | *MATa, his3Δ1, leu2Δ0, ura3Δ0, LYS+, can1Δ::STE2pr-LEU2, lyp1Δ::, tor1-1, fpr1Δ::klURA, Tub4-(6)-RFP-(24)-FKBP::natNT2, COG6-FRB::hphNT1, YOL111C-GFP::HIS3* | This study | 1.31 | 3.35 |
| OGYSGA1691 | *MATa, his3Δ1, leu2Δ0, ura3Δ0, LYS+, can1Δ::STE2pr-LEU2, lyp1Δ::, tor1-1, fpr1Δ::klURA, Tub4-(6)-RFP-(24)-FKBP::natNT2, COG6-FRB::hphNT1, YOR007C-GFP::HIS3* | This study | 0 | 1.02 |
| OGYSGA1692 | *MATa, his3Δ1, leu2Δ0, ura3Δ0, LYS+, can1Δ::STE2pr-LEU2, lyp1Δ::, tor1-1, fpr1Δ::klURA, Tub4-(6)-RFP-(24)-FKBP::natNT2, COG6-FRB::hphNT1, YOR327C-GFP::HIS3* | This study | 38.21 | 27.24 |
| OGYSGA1693 | *MATa, his3Δ1, leu2Δ0, ura3Δ0, LYS+, can1Δ::STE2pr-LEU2, lyp1Δ::, tor1-1, fpr1Δ::klURA, Tub4-(6)-RFP-(24)-FKBP::natNT2, COG6-FRB::hphNT1, YEL042W-GFP::HIS3* | This study | 0 | 0 |
| OGYSGA1694 | *MATa, his3Δ1, leu2Δ0, ura3Δ0, LYS+, can1Δ::STE2pr-LEU2, lyp1Δ::, tor1-1, fpr1Δ::klURA, Tub4-(6)-RFP-(24)-FKBP::natNT2, COG6-FRB::hphNT1, YDL074C-GFP::HIS3* | This study | 0.02 | 1.17 |
| OGYSGA1695 | *MATa, his3Δ1, leu2Δ0, ura3Δ0, LYS+, can1Δ::STE2pr-LEU2, lyp1Δ::, tor1-1, fpr1Δ::klURA, Tub4-(6)-RFP-(24)-FKBP::natNT2, COG6-FRB::hphNT1, YJL053W-GFP::HIS3* | This study | 0.7 | 1.83 |
| OGYSGA1696 | *MATa, his3Δ1, leu2Δ0, ura3Δ0, LYS+, can1Δ::STE2pr-LEU2, lyp1Δ::, tor1-1, fpr1Δ::klURA, Tub4-(6)-RFP-(24)-FKBP::natNT2, COG6-FRB::hphNT1, YDR334W-GFP::HIS3* | This study | 0.25 | 0.27 |
| OGYSGA1697 | *MATa, his3Δ1, leu2Δ0, ura3Δ0, LYS+, can1Δ::STE2pr-LEU2, lyp1Δ::, tor1-1, fpr1Δ::klURA, Tub4-(6)-RFP-(24)-FKBP::natNT2, COG6-FRB::hphNT1, YML041C-GFP::HIS3* | This study | 0 | 0.05 |
| OGYSGA1698 | *MATa, his3Δ1, leu2Δ0, ura3Δ0, LYS+, can1Δ::STE2pr-LEU2, lyp1Δ::, tor1-1, fpr1Δ::klURA, Tub4-(6)-RFP-(24)-FKBP::natNT2, COG6-FRB::hphNT1, YHR004C-GFP::HIS3* | This study | 0.14 | 0.25 |
| OGYSGA1699 | *MATa, his3Δ1, leu2Δ0, ura3Δ0, LYS+, can1Δ::STE2pr-LEU2, lyp1Δ::, tor1-1, fpr1Δ::klURA, Tub4-(6)-RFP-(24)-FKBP::natNT2, COG6-FRB::hphNT1, YBR288C-GFP::HIS3* | This study | 1.03 | 1.42 |
| OGYSGA1700 | *MATa, his3Δ1, leu2Δ0, ura3Δ0, LYS+, can1Δ::STE2pr-LEU2, lyp1Δ::, tor1-1, fpr1Δ::klURA, Tub4-(6)-RFP-(24)-FKBP::natNT2, COG6-FRB::hphNT1, YNR051C-GFP::HIS3* | This study | 1.18 | 0.33 |
| OGYSGA1701 | *MATa, his3Δ1, leu2Δ0, ura3Δ0, LYS+, can1Δ::STE2pr-LEU2, lyp1Δ::, tor1-1, fpr1Δ::klURA, Tub4-(6)-RFP-(24)-FKBP::natNT2, COG6-FRB::hphNT1, YDL145C-GFP::HIS3* | This study | 2.47 | 1.21 |
| OGYSGA1702 | *MATa, his3Δ1, leu2Δ0, ura3Δ0, LYS+, can1Δ::STE2pr-LEU2, lyp1Δ::, tor1-1, fpr1Δ::klURA, Tub4-(6)-RFP-(24)-FKBP::natNT2, COG6-FRB::hphNT1, YJL154C-GFP::HIS3* | This study | 1.62 | 1.34 |
| OGYSGA1703 | *MATa, his3Δ1, leu2Δ0, ura3Δ0, LYS+, can1Δ::STE2pr-LEU2, lyp1Δ::, tor1-1, fpr1Δ::klURA, Tub4-(6)-RFP-(24)-FKBP::natNT2, COG6-FRB::hphNT1, YOR069W-GFP::HIS3* | This study | 0.97 | 3.24 |
| OGYSGA1704 | *MATa, his3Δ1, leu2Δ0, ura3Δ0, LYS+, can1Δ::STE2pr-LEU2, lyp1Δ::, tor1-1, fpr1Δ::klURA, Tub4-(6)-RFP-(24)-FKBP::natNT2, COG6-FRB::hphNT1, YLL040C-GFP::HIS3* | This study | 1.01 | 2.79 |

| | | | | |
|---|---|---|---|---|
| OGYSGA1705 | *MATa, his3Δ1, leu2Δ0, ura3Δ0, LYS+, can1Δ::STE2pr-LEU2, lyp1Δ::, tor1-1, fpr1Δ::klURA, Tub4-(6)-RFP-(24)-FKBP::natNT2, COG6-FRB::hphNT1, YJR073C-GFP::HIS3* | This study | 1 | 0.46 |
| OGYSGA1706 | *MATa, his3Δ1, leu2Δ0, ura3Δ0, LYS+, can1Δ::STE2pr-LEU2, lyp1Δ::, tor1-1, fpr1Δ::klURA, Tub4-(6)-RFP-(24)-FKBP::natNT2, COG6-FRB::hphNT1, YDR320C-GFP::HIS3* | This study | 0.29 | 0.74 |
| OGYSGA1707 | *MATa, his3Δ1, leu2Δ0, ura3Δ0, LYS+, can1Δ::STE2pr-LEU2, lyp1Δ::, tor1-1, fpr1Δ::klURA, Tub4-(6)-RFP-(24)-FKBP::natNT2, COG6-FRB::hphNT1, YKL212W-GFP::HIS3* | This study | 0.58 | 0.25 |
| OGYSGA1708 | *MATa, his3Δ1, leu2Δ0, ura3Δ0, LYS+, can1Δ::STE2pr-LEU2, lyp1Δ::, tor1-1, fpr1Δ::klURA, Tub4-(6)-RFP-(24)-FKBP::natNT2, COG6-FRB::hphNT1, YOR109W-GFP::HIS3* | This study | 2.59 | 1.87 |
| OGYSGA1709 | *MATa, his3Δ1, leu2Δ0, ura3Δ0, LYS+, can1Δ::STE2pr-LEU2, lyp1Δ::, tor1-1, fpr1Δ::klURA, Tub4-(6)-RFP-(24)-FKBP::natNT2, COG6-FRB::hphNT1, YKL004W-GFP::HIS3* | This study | 1.87 | 3.46 |
| OGYSGA1710 | *MATa, his3Δ1, leu2Δ0, ura3Δ0, LYS+, can1Δ::STE2pr-LEU2, lyp1Δ::, tor1-1, fpr1Δ::klURA, Tub4-(6)-RFP-(24)-FKBP::natNT2, COG6-FRB::hphNT1, YIL033C-GFP::HIS3* | This study | 0 | 0.36 |
| OGYSGA1711 | *MATa, his3Δ1, leu2Δ0, ura3Δ0, LYS+, can1Δ::STE2pr-LEU2, lyp1Δ::, tor1-1, fpr1Δ::klURA, Tub4-(6)-RFP-(24)-FKBP::natNT2, COG6-FRB::hphNT1, YFL024C-GFP::HIS3* | This study | 1.62 | 0.71 |
| OGYSGA1712 | *MATa, his3Δ1, leu2Δ0, ura3Δ0, LYS+, can1Δ::STE2pr-LEU2, lyp1Δ::, tor1-1, fpr1Δ::klURA, Tub4-(6)-RFP-(24)-FKBP::natNT2, COG6-FRB::hphNT1, YOR123C-GFP::HIS3* | This study | 1.17 | 0.8 |
| OGYSGA1713 | *MATa, his3Δ1, leu2Δ0, ura3Δ0, LYS+, can1Δ::STE2pr-LEU2, lyp1Δ::, tor1-1, fpr1Δ::klURA, Tub4-(6)-RFP-(24)-FKBP::natNT2, COG6-FRB::hphNT1, YDR146C-GFP::HIS3* | This study | 0 | 0 |
| OGYSGA1714 | *MATa, his3Δ1, leu2Δ0, ura3Δ0, LYS+, can1Δ::STE2pr-LEU2, lyp1Δ::, tor1-1, fpr1Δ::klURA, Tub4-(6)-RFP-(24)-FKBP::natNT2, COG6-FRB::hphNT1, YER031C-GFP::HIS3* | This study | 0 | 0 |
| OGYSGA1715 | *MATa, his3Δ1, leu2Δ0, ura3Δ0, LYS+, can1Δ::STE2pr-LEU2, lyp1Δ::, tor1-1, fpr1Δ::klURA, Tub4-(6)-RFP-(24)-FKBP::natNT2, COG6-FRB::hphNT1, YGL115W-GFP::HIS3* | This study | 0 | 0 |
| OGYSGA1716 | *MATa, his3Δ1, leu2Δ0, ura3Δ0, LYS+, can1Δ::STE2pr-LEU2, lyp1Δ::, tor1-1, fpr1Δ::klURA, Tub4-(6)-RFP-(24)-FKBP::natNT2, COG6-FRB::hphNT1, YOR089C-GFP::HIS3* | This study | 0 | 0 |
| OGYSGA1718 | *MATa, his3Δ1, leu2Δ0, ura3Δ0, LYS+, can1Δ::STE2pr-LEU2, lyp1Δ::, tor1-1, fpr1Δ::klURA, Tub4-(6)-RFP-(24)-FKBP::natNT2, COG6-FRB::hphNT1, YKL119C-GFP::HIS3* | This study | 3.12 | 0.32 |
| OGYSGA1719 | *MATa, his3Δ1, leu2Δ0, ura3Δ0, LYS+, can1Δ::STE2pr-LEU2, lyp1Δ::, tor1-1, fpr1Δ::klURA, Tub4-(6)-RFP-(24)-FKBP::natNT2, COG6-FRB::hphNT1, YMR123W-GFP::HIS3* | This study | 0.31 | 0 |
| OGYSGA1720 | *MATa, his3Δ1, leu2Δ0, ura3Δ0, LYS+, can1Δ::STE2pr-LEU2, lyp1Δ::, tor1-1, fpr1Δ::klURA, Tub4-(6)-RFP-(24)-FKBP::natNT2, COG6-FRB::hphNT1, YEL002C-GFP::HIS3* | This study | 0.82 | 0.06 |
| OGYSGA1721 | *MATa, his3Δ1, leu2Δ0, ura3Δ0, LYS+, can1Δ::STE2pr-LEU2, lyp1Δ::, tor1-1, fpr1Δ::klURA, Tub4-(6)-RFP-(24)-FKBP::natNT2, COG6-FRB::hphNT1, YBR036C-GFP::HIS3* | This study | 0.39 | 1.69 |
| OGYSGA1722 | *MATa, his3Δ1, leu2Δ0, ura3Δ0, LYS+, can1Δ::STE2pr-LEU2, lyp1Δ::, tor1-1, fpr1Δ::klURA, Tub4-(6)-RFP-(24)-FKBP::natNT2, COG6-FRB::hphNT1, YEL031W-GFP::HIS3* | This study | 0.58 | 0 |
| OGYSGA1723 | *MATa, his3Δ1, leu2Δ0, ura3Δ0, LYS+, can1Δ::STE2pr-LEU2, lyp1Δ::, tor1-1, fpr1Δ::klURA, Tub4-(6)-RFP-(24)-FKBP::natNT2, COG6-FRB::hphNT1, YHR090C-GFP::HIS3* | This study | 0 | 0 |
| OGYSGA1725 | *MATa, his3Δ1, leu2Δ0, ura3Δ0, LYS+, can1Δ::STE2pr-LEU2, lyp1Δ::, tor1-1, fpr1Δ::klURA, Tub4-(6)-RFP-(24)-FKBP::natNT2, COG6-FRB::hphNT1, YDL185W-GFP::HIS3* | This study | 1.18 | 0 |
| OGYSGA1726 | *MATa, his3Δ1, leu2Δ0, ura3Δ0, LYS+, can1Δ::STE2pr-LEU2, lyp1Δ::, tor1-1, fpr1Δ::klURA, Tub4-(6)-RFP-(24)-FKBP::natNT2, COG6-FRB::hphNT1, YPL057C-GFP::HIS3* | This study | 4.02 | 3.58 |
| OGYSGA1727 | *MATa, his3Δ1, leu2Δ0, ura3Δ0, LYS+, can1Δ::STE2pr-LEU2, lyp1Δ::, tor1-1, fpr1Δ::klURA, Tub4-(6)-RFP-(24)-FKBP::natNT2, COG6-FRB::hphNT1, YJL183W-GFP::HIS3* | This study | 2.48 | 4.16 |
| OGYSGA1728 | *MATa, his3Δ1, leu2Δ0, ura3Δ0, LYS+, can1Δ::STE2pr-LEU2, lyp1Δ::, tor1-1, fpr1Δ::klURA, Tub4-(6)-RFP-(24)-FKBP::natNT2, COG6-FRB::hphNT1, YBR080C-GFP::HIS3* | This study | 2.12 | 1.55 |
| OGYSGA1729 | *MATa, his3Δ1, leu2Δ0, ura3Δ0, LYS+, can1Δ::STE2pr-LEU2, lyp1Δ::, tor1-1, fpr1Δ::klURA, Tub4-(6)-RFP-(24)-FKBP::natNT2, COG6-FRB::hphNT1, YBR168W-GFP::HIS3* | This study | 0.7 | 0.97 |
| OGYSGA1730 | *MATa, his3Δ1, leu2Δ0, ura3Δ0, LYS+, can1Δ::STE2pr-LEU2, lyp1Δ::, tor1-1, fpr1Δ::klURA, Tub4-(6)-RFP-(24)-FKBP::natNT2, COG6-FRB::hphNT1, YLR350W-GFP::HIS3* | This study | 0.41 | 0.97 |
| OGYSGA1731 | *MATa, his3Δ1, leu2Δ0, ura3Δ0, LYS+, can1Δ::STE2pr-LEU2, lyp1Δ::, tor1-1, fpr1Δ::klURA, Tub4-(6)-RFP-(24)-FKBP::natNT2, COG6-FRB::hphNT1, YNL169C-GFP::HIS3* | This study | 1.46 | 2.09 |
| OGYSGA1732 | *MATa, his3Δ1, leu2Δ0, ura3Δ0, LYS+, can1Δ::STE2pr-LEU2, lyp1Δ::, tor1-1, fpr1Δ::klURA, Tub4-(6)-RFP-(24)-FKBP::natNT2, COG6-FRB::hphNT1, YHR181W-GFP::HIS3* | This study | 0.93 | 1.61 |
| OGYSGA1733 | *MATa, his3Δ1, leu2Δ0, ura3Δ0, LYS+, can1Δ::STE2pr-LEU2, lyp1Δ::, tor1-1, fpr1Δ::klURA, Tub4-(6)-RFP-(24)-FKBP::natNT2, COG6-FRB::hphNT1, YLR085C-GFP::HIS3* | This study | 0 | 0 |
| OGYSGA1734 | *MATa, his3Δ1, leu2Δ0, ura3Δ0, LYS+, can1Δ::STE2pr-LEU2, lyp1Δ::, tor1-1, fpr1Δ::klURA, Tub4-(6)-RFP-(24)-FKBP::natNT2, COG6-FRB::hphNT1, YGL167C-GFP::HIS3* | This study | 0.87 | 0.46 |
| OGYSGA1735 | *MATa, his3Δ1, leu2Δ0, ura3Δ0, LYS+, can1Δ::STE2pr-LEU2, lyp1Δ::, tor1-1, fpr1Δ::klURA, Tub4-(6)-RFP-(24)-FKBP::natNT2, COG6-FRB::hphNT1, YLL038C-GFP::HIS3* | This study | 0 | 0 |
| OGYSGA1736 | *MATa, his3Δ1, leu2Δ0, ura3Δ0, LYS+, can1Δ::STE2pr-LEU2, lyp1Δ::, tor1-1, fpr1Δ::klURA, Tub4-(6)-RFP-(24)-FKBP::natNT2, COG6-FRB::hphNT1, YGR089W-GFP::HIS3* | This study | 1.91 | 1.36 |
| OGYSGA1737 | *MATa, his3Δ1, leu2Δ0, ura3Δ0, LYS+, can1Δ::STE2pr-LEU2, lyp1Δ::, tor1-1, fpr1Δ::klURA, Tub4-(6)-RFP-(24)-FKBP::natNT2, COG6-FRB::hphNT1, YMR004W-GFP::HIS3* | This study | 3.47 | 1.06 |
| OGYSGA1738 | *MATa, his3Δ1, leu2Δ0, ura3Δ0, LYS+, can1Δ::STE2pr-LEU2, lyp1Δ::, tor1-1, fpr1Δ::klURA, Tub4-(6)-RFP-(24)-FKBP::natNT2, COG6-FRB::hphNT1, YBR231C-GFP::HIS3* | This study | 0.6 | 1.26 |
| OGYSGA1739 | *MATa, his3Δ1, leu2Δ0, ura3Δ0, LYS+, can1Δ::STE2pr-LEU2, lyp1Δ::, tor1-1, fpr1Δ::klURA, Tub4-(6)-RFP-(24)-FKBP::natNT2, COG6-FRB::hphNT1, YMR307W-GFP::HIS3* | This study | 0.52 | 0 |
| OGYSGA1740 | *MATa, his3Δ1, leu2Δ0, ura3Δ0, LYS+, can1Δ::STE2pr-LEU2, lyp1Δ::, tor1-1, fpr1Δ::klURA, Tub4-(6)-RFP-(24)-FKBP::natNT2, COG6-FRB::hphNT1, YGR106C-GFP::HIS3* | This study | 0 | 3.12 |
| OGYSGA1741 | *MATa, his3Δ1, leu2Δ0, ura3Δ0, LYS+, can1Δ::STE2pr-LEU2, lyp1Δ::, tor1-1, fpr1Δ::klURA, Tub4-(6)-RFP-(24)-FKBP::natNT2, COG6-FRB::hphNT1, YJL024C-GFP::HIS3* | This study | 0.79 | 1.24 |
| OGYSGA1742 | *MATa, his3Δ1, leu2Δ0, ura3Δ0, LYS+, can1Δ::STE2pr-LEU2, lyp1Δ::, tor1-1, fpr1Δ::klURA, Tub4-(6)-RFP-(24)-FKBP::natNT2, COG6-FRB::hphNT1, YOR244W-GFP::HIS3* | This study | 0.2 | 0.83 |
| OGYSGA1743 | *MATa, his3Δ1, leu2Δ0, ura3Δ0, LYS+, can1Δ::STE2pr-LEU2, lyp1Δ::, tor1-1, fpr1Δ::klURA, Tub4-(6)-RFP-(24)-FKBP::natNT2, COG6-FRB::hphNT1, YOL012C-GFP::HIS3* | This study | 0.97 | 0.74 |
| OGYSGA1744 | *MATa, his3Δ1, leu2Δ0, ura3Δ0, LYS+, can1Δ::STE2pr-LEU2, lyp1Δ::, tor1-1, fpr1Δ::klURA, Tub4-(6)-RFP-(24)-FKBP::natNT2, COG6-FRB::hphNT1, YHR059W-GFP::HIS3* | This study | 1.78 | 0.8 |
| OGYSGA1745 | *MATa, his3Δ1, leu2Δ0, ura3Δ0, LYS+, can1Δ::STE2pr-LEU2, lyp1Δ::, tor1-1, fpr1Δ::klURA, Tub4-(6)-RFP-(24)-FKBP::natNT2, COG6-FRB::hphNT1, YGR105W-GFP::HIS3* | This study | 0.26 | 0.75 |
| OGYSGA1746 | *MATa, his3Δ1, leu2Δ0, ura3Δ0, LYS+, can1Δ::STE2pr-LEU2, lyp1Δ::, tor1-1, fpr1Δ::klURA, Tub4-(6)-RFP-(24)-FKBP::natNT2, COG6-FRB::hphNT1, YCL001W-GFP::HIS3* | This study | 0.88 | 1.02 |

| | | | | |
|---|---|---|---|---|
| OGYSGA1747 | MATa, his3Δ1, leu2Δ0, ura3Δ0, LYS+, can1Δ::STE2pr-LEU2, lyp1Δ::, tor1-1, fpr1Δ::klURA, Tub4-(6)-RFP-(24)-FKBP::natNT2, COG6-FRB::hphNT1, YNL322C-GFP::HIS3 | This study | 0.51 | 10.22 |
| OGYSGA1748 | MATa, his3Δ1, leu2Δ0, ura3Δ0, LYS+, can1Δ::STE2pr-LEU2, lyp1Δ::, tor1-1, fpr1Δ::klURA, Tub4-(6)-RFP-(24)-FKBP::natNT2, COG6-FRB::hphNT1, YPR057W-GFP::HIS3 | This study | 0 | 0 |
| OGYSGA1749 | MATa, his3Δ1, leu2Δ0, ura3Δ0, LYS+, can1Δ::STE2pr-LEU2, lyp1Δ::, tor1-1, fpr1Δ::klURA, Tub4-(6)-RFP-(24)-FKBP::natNT2, COG6-FRB::hphNT1, YIR005W-GFP::HIS3 | This study | 0 | 0 |
| OGYSGA1750 | MATa, his3Δ1, leu2Δ0, ura3Δ0, LYS+, can1Δ::STE2pr-LEU2, lyp1Δ::, tor1-1, fpr1Δ::klURA, Tub4-(6)-RFP-(24)-FKBP::natNT2, COG6-FRB::hphNT1, YGL212W-GFP::HIS3 | This study | 0 | 0 |
| OGYSGA1751 | MATa, his3Δ1, leu2Δ0, ura3Δ0, LYS+, can1Δ::STE2pr-LEU2, lyp1Δ::, tor1-1, fpr1Δ::klURA, Tub4-(6)-RFP-(24)-FKBP::natNT2, COG6-FRB::hphNT1, YCR063W-GFP::HIS3 | This study | 0 | 0.61 |
| OGYSGA1752 | MATa, his3Δ1, leu2Δ0, ura3Δ0, LYS+, can1Δ::STE2pr-LEU2, lyp1Δ::, tor1-1, fpr1Δ::klURA, Tub4-(6)-RFP-(24)-FKBP::natNT2, COG6-FRB::hphNT1, YPL234C-GFP::HIS3 | This study | 3.17 | 0.6 |
| OGYSGA1753 | MATa, his3Δ1, leu2Δ0, ura3Δ0, LYS+, can1Δ::STE2pr-LEU2, lyp1Δ::, tor1-1, fpr1Δ::klURA, Tub4-(6)-RFP-(24)-FKBP::natNT2, COG6-FRB::hphNT1, YDR331W-GFP::HIS3 | This study | 0.53 | 0.68 |
| OGYSGA1754 | MATa, his3Δ1, leu2Δ0, ura3Δ0, LYS+, can1Δ::STE2pr-LEU2, lyp1Δ::, tor1-1, fpr1Δ::klURA, Tub4-(6)-RFP-(24)-FKBP::natNT2, COG6-FRB::hphNT1, YJL062W-GFP::HIS3 | This study | 0.72 | 0.64 |
| OGYSGA1755 | MATa, his3Δ1, leu2Δ0, ura3Δ0, LYS+, can1Δ::STE2pr-LEU2, lyp1Δ::, tor1-1, fpr1Δ::klURA, Tub4-(6)-RFP-(24)-FKBP::natNT2, COG6-FRB::hphNT1, YJR075W-GFP::HIS3 | This study | 4.44 | 2.09 |
| OGYSGA1756 | MATa, his3Δ1, leu2Δ0, ura3Δ0, LYS+, can1Δ::STE2pr-LEU2, lyp1Δ::, tor1-1, fpr1Δ::klURA, Tub4-(6)-RFP-(24)-FKBP::natNT2, COG6-FRB::hphNT1, YKL139W-GFP::HIS3 | This study | 0.25 | 1.76 |
| OGYSGA1757 | MATa, his3Δ1, leu2Δ0, ura3Δ0, LYS+, can1Δ::STE2pr-LEU2, lyp1Δ::, tor1-1, fpr1Δ::klURA, Tub4-(6)-RFP-(24)-FKBP::natNT2, COG6-FRB::hphNT1, YML097C-GFP::HIS3 | This study | 0 | 0 |
| OGYSGA1758 | MATa, his3Δ1, leu2Δ0, ura3Δ0, LYS+, can1Δ::STE2pr-LEU2, lyp1Δ::, tor1-1, fpr1Δ::klURA, Tub4-(6)-RFP-(24)-FKBP::natNT2, COG6-FRB::hphNT1, YKL048C-GFP::HIS3 | This study | 0 | 0 |
| OGYSGA1759 | MATa, his3Δ1, leu2Δ0, ura3Δ0, LYS+, can1Δ::STE2pr-LEU2, lyp1Δ::, tor1-1, fpr1Δ::klURA, Tub4-(6)-RFP-(24)-FKBP::natNT2, COG6-FRB::hphNT1, YLR200W-GFP::HIS3 | This study | 0 | 0 |
| OGYSGA1760 | MATa, his3Δ1, leu2Δ0, ura3Δ0, LYS+, can1Δ::STE2pr-LEU2, lyp1Δ::, tor1-1, fpr1Δ::klURA, Tub4-(6)-RFP-(24)-FKBP::natNT2, COG6-FRB::hphNT1, YDR225W-GFP::HIS3 | This study | 0 | 0 |
| OGYSGA1761 | MATa, his3Δ1, leu2Δ0, ura3Δ0, LYS+, can1Δ::STE2pr-LEU2, lyp1Δ::, tor1-1, fpr1Δ::klURA, Tub4-(6)-RFP-(24)-FKBP::natNT2, COG6-FRB::hphNT1, YPL069C-GFP::HIS3 | This study | 0 | 0 |
| OGYSGA1762 | MATa, his3Δ1, leu2Δ0, ura3Δ0, LYS+, can1Δ::STE2pr-LEU2, lyp1Δ::, tor1-1, fpr1Δ::klURA, Tub4-(6)-RFP-(24)-FKBP::natNT2, COG6-FRB::hphNT1, YKR001C-GFP::HIS3 | This study | 1.23 | 1.13 |
| OGYSGA1763 | MATa, his3Δ1, leu2Δ0, ura3Δ0, LYS+, can1Δ::STE2pr-LEU2, lyp1Δ::, tor1-1, fpr1Δ::klURA, Tub4-(6)-RFP-(24)-FKBP::natNT2, COG6-FRB::hphNT1, YGR157W-GFP::HIS3 | This study | 1.7 | 2.65 |
| OGYSGA1764 | MATa, his3Δ1, leu2Δ0, ura3Δ0, LYS+, can1Δ::STE2pr-LEU2, lyp1Δ::, tor1-1, fpr1Δ::klURA, Tub4-(6)-RFP-(24)-FKBP::natNT2, COG6-FRB::hphNT1, YPL158C-GFP::HIS3 | This study | 0 | 0.05 |
| OGYSGA1765 | MATa, his3Δ1, leu2Δ0, ura3Δ0, LYS+, can1Δ::STE2pr-LEU2, lyp1Δ::, tor1-1, fpr1Δ::klURA, Tub4-(6)-RFP-(24)-FKBP::natNT2, COG6-FRB::hphNT1, YIL076W-GFP::HIS3 | This study | 1.45 | 1.46 |
| OGYSGA1766 | MATa, his3Δ1, leu2Δ0, ura3Δ0, LYS+, can1Δ::STE2pr-LEU2, lyp1Δ::, tor1-1, fpr1Δ::klURA, Tub4-(6)-RFP-(24)-FKBP::natNT2, COG6-FRB::hphNT1, YDR434W-GFP::HIS3 | This study | 0.39 | 0.71 |
| OGYSGA1767 | MATa, his3Δ1, leu2Δ0, ura3Δ0, LYS+, can1Δ::STE2pr-LEU2, lyp1Δ::, tor1-1, fpr1Δ::klURA, Tub4-(6)-RFP-(24)-FKBP::natNT2, COG6-FRB::hphNT1, YPR017C-GFP::HIS3 | This study | 0 | 0 |
| OGYSGA1768 | MATa, his3Δ1, leu2Δ0, ura3Δ0, LYS+, can1Δ::STE2pr-LEU2, lyp1Δ::, tor1-1, fpr1Δ::klURA, Tub4-(6)-RFP-(24)-FKBP::natNT2, COG6-FRB::hphNT1, YLL006W-GFP::HIS3 | This study | 0 | 0.26 |
| OGYSGA1769 | MATa, his3Δ1, leu2Δ0, ura3Δ0, LYS+, can1Δ::STE2pr-LEU2, lyp1Δ::, tor1-1, fpr1Δ::klURA, Tub4-(6)-RFP-(24)-FKBP::natNT2, COG6-FRB::hphNT1, YDR485C-GFP::HIS3 | This study | 1.11 | 1.53 |
| OGYSGA1770 | MATa, his3Δ1, leu2Δ0, ura3Δ0, LYS+, can1Δ::STE2pr-LEU2, lyp1Δ::, tor1-1, fpr1Δ::klURA, Tub4-(6)-RFP-(24)-FKBP::natNT2, COG6-FRB::hphNT1, YJL034W-GFP::HIS3 | This study | 0 | 0 |
| OGYSGA1771 | MATa, his3Δ1, leu2Δ0, ura3Δ0, LYS+, can1Δ::STE2pr-LEU2, lyp1Δ::, tor1-1, fpr1Δ::klURA, Tub4-(6)-RFP-(24)-FKBP::natNT2, COG6-FRB::hphNT1, YJL210W-GFP::HIS3 | This study | 0.75 | 0.97 |
| OGYSGA1772 | MATa, his3Δ1, leu2Δ0, ura3Δ0, LYS+, can1Δ::STE2pr-LEU2, lyp1Δ::, tor1-1, fpr1Δ::klURA, Tub4-(6)-RFP-(24)-FKBP::natNT2, COG6-FRB::hphNT1, YER111C-GFP::HIS3 | This study | 1.5 | 0.43 |
| OGYSGA1773 | MATa, his3Δ1, leu2Δ0, ura3Δ0, LYS+, can1Δ::STE2pr-LEU2, lyp1Δ::, tor1-1, fpr1Δ::klURA, Tub4-(6)-RFP-(24)-FKBP::natNT2, COG6-FRB::hphNT1, YDR186C-GFP::HIS3 | This study | 0 | 0 |
| OGYSGA1774 | MATa, his3Δ1, leu2Δ0, ura3Δ0, LYS+, can1Δ::STE2pr-LEU2, lyp1Δ::, tor1-1, fpr1Δ::klURA, Tub4-(6)-RFP-(24)-FKBP::natNT2, COG6-FRB::hphNT1, YNL008C-GFP::HIS3 | This study | 1.03 | 1.67 |
| OGYSGA1775 | MATa, his3Δ1, leu2Δ0, ura3Δ0, LYS+, can1Δ::STE2pr-LEU2, lyp1Δ::, tor1-1, fpr1Δ::klURA, Tub4-(6)-RFP-(24)-FKBP::natNT2, COG6-FRB::hphNT1, YOR112W-GFP::HIS3 | This study | 1.13 | 0 |
| OGYSGA1776 | MATa, his3Δ1, leu2Δ0, ura3Δ0, LYS+, can1Δ::STE2pr-LEU2, lyp1Δ::, tor1-1, fpr1Δ::klURA, Tub4-(6)-RFP-(24)-FKBP::natNT2, COG6-FRB::hphNT1, YLR208W-GFP::HIS3 | This study | 1.41 | 1 |
| OGYSGA1777 | MATa, his3Δ1, leu2Δ0, ura3Δ0, LYS+, can1Δ::STE2pr-LEU2, lyp1Δ::, tor1-1, fpr1Δ::klURA, Tub4-(6)-RFP-(24)-FKBP::natNT2, COG6-FRB::hphNT1, YER151C-GFP::HIS3 | This study | 0 | 0.14 |
| OGYSGA1778 | MATa, his3Δ1, leu2Δ0, ura3Δ0, LYS+, can1Δ::STE2pr-LEU2, lyp1Δ::, tor1-1, fpr1Δ::klURA, Tub4-(6)-RFP-(24)-FKBP::natNT2, COG6-FRB::hphNT1, YJL002C-GFP::HIS3 | This study | 0.73 | 1.47 |
| OGYSGA1779 | MATa, his3Δ1, leu2Δ0, ura3Δ0, LYS+, can1Δ::STE2pr-LEU2, lyp1Δ::, tor1-1, fpr1Δ::klURA, Tub4-(6)-RFP-(24)-FKBP::natNT2, COG6-FRB::hphNT1, YAL009W-GFP::HIS3 | This study | 0.68 | 1.85 |
| OGYSGA1780 | MATa, his3Δ1, leu2Δ0, ura3Δ0, LYS+, can1Δ::STE2pr-LEU2, lyp1Δ::, tor1-1, fpr1Δ::klURA, Tub4-(6)-RFP-(24)-FKBP::natNT2, COG6-FRB::hphNT1, YHL033C-GFP::HIS3 | This study | 0 | 0 |
| OGYSGA1781 | MATa, his3Δ1, leu2Δ0, ura3Δ0, LYS+, can1Δ::STE2pr-LEU2, lyp1Δ::, tor1-1, fpr1Δ::klURA, Tub4-(6)-RFP-(24)-FKBP::natNT2, COG6-FRB::hphNT1, YEL043W-GFP::HIS3 | This study | 0 | 0 |
| OGYSGA1782 | MATa, his3Δ1, leu2Δ0, ura3Δ0, LYS+, can1Δ::STE2pr-LEU2, lyp1Δ::, tor1-1, fpr1Δ::klURA, Tub4-(6)-RFP-(24)-FKBP::natNT2, COG6-FRB::hphNT1, YJL082W-GFP::HIS3 | This study | 0.43 | 0 |
| OGYSGA1783 | MATa, his3Δ1, leu2Δ0, ura3Δ0, LYS+, can1Δ::STE2pr-LEU2, lyp1Δ::, tor1-1, fpr1Δ::klURA, Tub4-(6)-RFP-(24)-FKBP::natNT2, COG6-FRB::hphNT1, YPL105C-GFP::HIS3 | This study | 0 | 0 |
| OGYSGA1784 | MATa, his3Δ1, leu2Δ0, ura3Δ0, LYS+, can1Δ::STE2pr-LEU2, lyp1Δ::, tor1-1, fpr1Δ::klURA, Tub4-(6)-RFP-(24)-FKBP::natNT2, COG6-FRB::hphNT1, YBL022C-GFP::HIS3 | This study | 0 | 1.98 |
| OGYSGA1785 | MATa, his3Δ1, leu2Δ0, ura3Δ0, LYS+, can1Δ::STE2pr-LEU2, lyp1Δ::, tor1-1, fpr1Δ::klURA, Tub4-(6)-RFP-(24)-FKBP::natNT2, COG6-FRB::hphNT1, YDR372C-GFP::HIS3 | This study | 0.89 | 13.71 |
| OGYSGA1786 | MATa, his3Δ1, leu2Δ0, ura3Δ0, LYS+, can1Δ::STE2pr-LEU2, lyp1Δ::, tor1-1, fpr1Δ::klURA, Tub4-(6)-RFP-(24)-FKBP::natNT2, COG6-FRB::hphNT1, YJR033C-GFP::HIS3 | This study | 0.51 | 0.37 |

| | | | | |
|---|---|---|---|---|
| OGYSGA1787 | *MATa, his3Δ1, leu2Δ0, ura3Δ0, LYS+, can1Δ::STE2pr-LEU2, lyp1Δ::, tor1-1, fpr1Δ::klURA, Tub4-(6)-RFP-(24)-FKBP::natNT2, COG6-FRB::hphNT1, YDR141C-GFP::HIS3* | This study | 1.65 | 3.46 |
| OGYSGA1788 | *MATa, his3Δ1, leu2Δ0, ura3Δ0, LYS+, can1Δ::STE2pr-LEU2, lyp1Δ::, tor1-1, fpr1Δ::klURA, Tub4-(6)-RFP-(24)-FKBP::natNT2, COG6-FRB::hphNT1, YGL219C-GFP::HIS3* | This study | 0 | 0.32 |
| OGYSGA1789 | *MATa, his3Δ1, leu2Δ0, ura3Δ0, LYS+, can1Δ::STE2pr-LEU2, lyp1Δ::, tor1-1, fpr1Δ::klURA, Tub4-(6)-RFP-(24)-FKBP::natNT2, COG6-FRB::hphNT1, YBR127C-GFP::HIS3* | This study | 0 | 0 |
| OGYSGA1790 | *MATa, his3Δ1, leu2Δ0, ura3Δ0, LYS+, can1Δ::STE2pr-LEU2, lyp1Δ::, tor1-1, fpr1Δ::klURA, Tub4-(6)-RFP-(24)-FKBP::natNT2, COG6-FRB::hphNT1, YOR078W-GFP::HIS3* | This study | 0 | 0.7 |
| OGYSGA1791 | *MATa, his3Δ1, leu2Δ0, ura3Δ0, LYS+, can1Δ::STE2pr-LEU2, lyp1Δ::, tor1-1, fpr1Δ::klURA, Tub4-(6)-RFP-(24)-FKBP::natNT2, COG6-FRB::hphNT1, YDR432W-GFP::HIS3* | This study | 0.19 | 0 |
| OGYSGA1792 | *MATa, his3Δ1, leu2Δ0, ura3Δ0, LYS+, can1Δ::STE2pr-LEU2, lyp1Δ::, tor1-1, fpr1Δ::klURA, Tub4-(6)-RFP-(24)-FKBP::natNT2, COG6-FRB::hphNT1, YDR069C-GFP::HIS3* | This study | 0 | 0 |
| OGYSGA1793 | *MATa, his3Δ1, leu2Δ0, ura3Δ0, LYS+, can1Δ::STE2pr-LEU2, lyp1Δ::, tor1-1, fpr1Δ::klURA, Tub4-(6)-RFP-(24)-FKBP::natNT2, COG6-FRB::hphNT1, YKL110C-GFP::HIS3* | This study | 0 | 0 |
| OGYSGA1794 | *MATa, his3Δ1, leu2Δ0, ura3Δ0, LYS+, can1Δ::STE2pr-LEU2, lyp1Δ::, tor1-1, fpr1Δ::klURA, Tub4-(6)-RFP-(24)-FKBP::natNT2, COG6-FRB::hphNT1, YLR260W-GFP::HIS3* | This study | 0 | 0 |
| OGYSGA1795 | *MATa, his3Δ1, leu2Δ0, ura3Δ0, LYS+, can1Δ::STE2pr-LEU2, lyp1Δ::, tor1-1, fpr1Δ::klURA, Tub4-(6)-RFP-(24)-FKBP::natNT2, COG6-FRB::hphNT1, YMR060C-GFP::HIS3* | This study | 0 | 0 |
| OGYSGA1796 | *MATa, his3Δ1, leu2Δ0, ura3Δ0, LYS+, can1Δ::STE2pr-LEU2, lyp1Δ::, tor1-1, fpr1Δ::klURA, Tub4-(6)-RFP-(24)-FKBP::natNT2, COG6-FRB::hphNT1, YGR072W-GFP::HIS3* | This study | 0 | 0 |
| OGYSGA1797 | *MATa, his3Δ1, leu2Δ0, ura3Δ0, LYS+, can1Δ::STE2pr-LEU2, lyp1Δ::, tor1-1, fpr1Δ::klURA, Tub4-(6)-RFP-(24)-FKBP::natNT2, COG6-FRB::hphNT1, YCR094W-GFP::HIS3* | This study | 2.96 | 3.26 |
| OGYSGA1798 | *MATa, his3Δ1, leu2Δ0, ura3Δ0, LYS+, can1Δ::STE2pr-LEU2, lyp1Δ::, tor1-1, fpr1Δ::klURA, Tub4-(6)-RFP-(24)-FKBP::natNT2, COG6-FRB::hphNT1, YDR297W-GFP::HIS3* | This study | 0.56 | 0.8 |
| OGYSGA1799 | *MATa, his3Δ1, leu2Δ0, ura3Δ0, LYS+, can1Δ::STE2pr-LEU2, lyp1Δ::, tor1-1, fpr1Δ::klURA, Tub4-(6)-RFP-(24)-FKBP::natNT2, COG6-FRB::hphNT1, YER089C-GFP::HIS3* | This study | 0 | 0 |
| OGYSGA1800 | *MATa, his3Δ1, leu2Δ0, ura3Δ0, LYS+, can1Δ::STE2pr-LEU2, lyp1Δ::, tor1-1, fpr1Δ::klURA, Tub4-(6)-RFP-(24)-FKBP::natNT2, COG6-FRB::hphNT1, YLR110C-GFP::HIS3* | This study | 0 | 0 |
| OGYSGA1801 | *MATa, his3Δ1, leu2Δ0, ura3Δ0, LYS+, can1Δ::STE2pr-LEU2, lyp1Δ::, tor1-1, fpr1Δ::klURA, Tub4-(6)-RFP-(24)-FKBP::natNT2, COG6-FRB::hphNT1, YNL141W-GFP::HIS3* | This study | 0 | 0 |
| OGYSGA1802 | *MATa, his3Δ1, leu2Δ0, ura3Δ0, LYS+, can1Δ::STE2pr-LEU2, lyp1Δ::, tor1-1, fpr1Δ::klURA, Tub4-(6)-RFP-(24)-FKBP::natNT2, COG6-FRB::hphNT1, YNL238W-GFP::HIS3* | This study | 2.32 | 1.83 |
| OGYSGA1803 | *MATa, his3Δ1, leu2Δ0, ura3Δ0, LYS+, can1Δ::STE2pr-LEU2, lyp1Δ::, tor1-1, fpr1Δ::klURA, Tub4-(6)-RFP-(24)-FKBP::natNT2, COG6-FRB::hphNT1, YKL006W-GFP::HIS3* | This study | 0.38 | 0 |
| OGYSGA1804 | *MATa, his3Δ1, leu2Δ0, ura3Δ0, LYS+, can1Δ::STE2pr-LEU2, lyp1Δ::, tor1-1, fpr1Δ::klURA, Tub4-(6)-RFP-(24)-FKBP::natNT2, COG6-FRB::hphNT1, YDL226C-GFP::HIS3* | This study | 0.73 | 0.76 |
| OGYSGA1805 | *MATa, his3Δ1, leu2Δ0, ura3Δ0, LYS+, can1Δ::STE2pr-LEU2, lyp1Δ::, tor1-1, fpr1Δ::klURA, Tub4-(6)-RFP-(24)-FKBP::natNT2, COG6-FRB::hphNT1, YNL251C-GFP::HIS3* | This study | 0.5 | 0.79 |
| OGYSGA1806 | *MATa, his3Δ1, leu2Δ0, ura3Δ0, LYS+, can1Δ::STE2pr-LEU2, lyp1Δ::, tor1-1, fpr1Δ::klURA, Tub4-(6)-RFP-(24)-FKBP::natNT2, VPS3-FRB::hphNT1, YNL243W-GFP::HIS3* | This study | 0 | 0 |
| OGYSGA1807 | *MATa, his3Δ1, leu2Δ0, ura3Δ0, LYS+, can1Δ::STE2pr-LEU2, lyp1Δ::, tor1-1, fpr1Δ::klURA, Tub4-(6)-RFP-(24)-FKBP::natNT2, VPS3-FRB::hphNT1, YJL115W-GFP::HIS3* | This study | 0.13 | 0.79 |
| OGYSGA1808 | *MATa, his3Δ1, leu2Δ0, ura3Δ0, LYS+, can1Δ::STE2pr-LEU2, lyp1Δ::, tor1-1, fpr1Δ::klURA, Tub4-(6)-RFP-(24)-FKBP::natNT2, VPS3-FRB::hphNT1, YOR244W-GFP::HIS3* | This study | 0.56 | 0.32 |
| OGYSGA1809 | *MATa, his3Δ1, leu2Δ0, ura3Δ0, LYS+, can1Δ::STE2pr-LEU2, lyp1Δ::, tor1-1, fpr1Δ::klURA, Tub4-(6)-RFP-(24)-FKBP::natNT2, VPS3-FRB::hphNT1, YGL020C-GFP::HIS3* | This study | 0.64 | 0.94 |
| OGYSGA1810 | *MATa, his3Δ1, leu2Δ0, ura3Δ0, LYS+, can1Δ::STE2pr-LEU2, lyp1Δ::, tor1-1, fpr1Δ::klURA, Tub4-(6)-RFP-(24)-FKBP::natNT2, VPS3-FRB::hphNT1, YDL185W-GFP::HIS3* | This study | 10.56 | 0 |
| OGYSGA1811 | *MATa, his3Δ1, leu2Δ0, ura3Δ0, LYS+, can1Δ::STE2pr-LEU2, lyp1Δ::, tor1-1, fpr1Δ::klURA, Tub4-(6)-RFP-(24)-FKBP::natNT2, VPS3-FRB::hphNT1, YKL113C-GFP::HIS3* | This study | 84.06 | 2.51 |
| OGYSGA1812 | *MATa, his3Δ1, leu2Δ0, ura3Δ0, LYS+, can1Δ::STE2pr-LEU2, lyp1Δ::, tor1-1, fpr1Δ::klURA, Tub4-(6)-RFP-(24)-FKBP::natNT2, VPS3-FRB::hphNT1, YER083C-GFP::HIS3* | This study | 0.52 | 0.46 |
| OGYSGA1813 | *MATa, his3Δ1, leu2Δ0, ura3Δ0, LYS+, can1Δ::STE2pr-LEU2, lyp1Δ::, tor1-1, fpr1Δ::klURA, Tub4-(6)-RFP-(24)-FKBP::natNT2, VPS3-FRB::hphNT1, YDL040C-GFP::HIS3* | This study | 0 | 0.05 |
| OGYSGA1814 | *MATa, his3Δ1, leu2Δ0, ura3Δ0, LYS+, can1Δ::STE2pr-LEU2, lyp1Δ::, tor1-1, fpr1Δ::klURA, Tub4-(6)-RFP-(24)-FKBP::natNT2, VPS3-FRB::hphNT1, YPL153C-GFP::HIS3* | This study | 3.67 | 15.69 |
| OGYSGA1815 | *MATa, his3Δ1, leu2Δ0, ura3Δ0, LYS+, can1Δ::STE2pr-LEU2, lyp1Δ::, tor1-1, fpr1Δ::klURA, Tub4-(6)-RFP-(24)-FKBP::natNT2, VPS3-FRB::hphNT1, YCL016C-GFP::HIS3* | This study | 0.94 | 2.08 |
| OGYSGA1816 | *MATa, his3Δ1, leu2Δ0, ura3Δ0, LYS+, can1Δ::STE2pr-LEU2, lyp1Δ::, tor1-1, fpr1Δ::klURA, Tub4-(6)-RFP-(24)-FKBP::natNT2, VPS3-FRB::hphNT1, YHR013C-GFP::HIS3* | This study | 0 | 0 |
| OGYSGA1817 | *MATa, his3Δ1, leu2Δ0, ura3Δ0, LYS+, can1Δ::STE2pr-LEU2, lyp1Δ::, tor1-1, fpr1Δ::klURA, Tub4-(6)-RFP-(24)-FKBP::natNT2, VPS3-FRB::hphNT1, YHR200W-GFP::HIS3* | This study | 0.52 | 0.74 |
| OGYSGA1818 | *MATa, his3Δ1, leu2Δ0, ura3Δ0, LYS+, can1Δ::STE2pr-LEU2, lyp1Δ::, tor1-1, fpr1Δ::klURA, Tub4-(6)-RFP-(24)-FKBP::natNT2, VPS3-FRB::hphNT1, YMR078C-GFP::HIS3* | This study | 1.06 | 1.5 |
| OGYSGA1819 | *MATa, his3Δ1, leu2Δ0, ura3Δ0, LYS+, can1Δ::STE2pr-LEU2, lyp1Δ::, tor1-1, fpr1Δ::klURA, Tub4-(6)-RFP-(24)-FKBP::natNT2, VPS3-FRB::hphNT1, YGR002C-GFP::HIS3* | This study | 1.07 | 1.06 |
| OGYSGA1820 | *MATa, his3Δ1, leu2Δ0, ura3Δ0, LYS+, can1Δ::STE2pr-LEU2, lyp1Δ::, tor1-1, fpr1Δ::klURA, Tub4-(6)-RFP-(24)-FKBP::natNT2, VPS3-FRB::hphNT1, YGL058W-GFP::HIS3* | This study | 0 | 0.24 |
| OGYSGA1821 | *MATa, his3Δ1, leu2Δ0, ura3Δ0, LYS+, can1Δ::STE2pr-LEU2, lyp1Δ::, tor1-1, fpr1Δ::klURA, Tub4-(6)-RFP-(24)-FKBP::natNT2, VPS3-FRB::hphNT1, YJR043C-GFP::HIS3* | This study | 0 | 0 |
| OGYSGA1822 | *MATa, his3Δ1, leu2Δ0, ura3Δ0, LYS+, can1Δ::STE2pr-LEU2, lyp1Δ::, tor1-1, fpr1Δ::klURA, Tub4-(6)-RFP-(24)-FKBP::natNT2, VPS3-FRB::hphNT1, YGR198W-GFP::HIS3* | This study | 0.06 | 0.11 |
| OGYSGA1823 | *MATa, his3Δ1, leu2Δ0, ura3Δ0, LYS+, can1Δ::STE2pr-LEU2, lyp1Δ::, tor1-1, fpr1Δ::klURA, Tub4-(6)-RFP-(24)-FKBP::natNT2, VPS3-FRB::hphNT1, YGL095C-GFP::HIS3* | This study | 23.33 | 82.04 |
| OGYSGA1826 | *MATa, his3Δ1, leu2Δ0, ura3Δ0, LYS+, can1Δ::STE2pr-LEU2, lyp1Δ::, tor1-1, fpr1Δ::klURA, Tub4-(6)-RFP-(24)-FKBP::natNT2, VPS3-FRB::hphNT1, YFL024C-GFP::HIS3* | This study | 0.82 | 0.69 |
| OGYSGA1827 | *MATa, his3Δ1, leu2Δ0, ura3Δ0, LYS+, can1Δ::STE2pr-LEU2, lyp1Δ::, tor1-1, fpr1Δ::klURA, Tub4-(6)-RFP-(24)-FKBP::natNT2, VPS3-FRB::hphNT1, YGL116W-GFP::HIS3* | This study | 0 | 0 |
| OGYSGA1828 | *MATa, his3Δ1, leu2Δ0, ura3Δ0, LYS+, can1Δ::STE2pr-LEU2, lyp1Δ::, tor1-1, fpr1Δ::klURA, Tub4-(6)-RFP-(24)-FKBP::natNT2, VPS3-FRB::hphNT1, YHR090C-GFP::HIS3* | This study | 0.12 | 0.02 |

| | | | | |
|---|---|---|---|---|
| OGYSGA1829 | *MATa, his3Δ1, leu2Δ0, ura3Δ0, LYS+, can1Δ::STE2pr-LEU2, lyp1Δ::, tor1-1, fpr1Δ::klURA, Tub4-(6)-RFP-(24)-FKBP::natNT2, VPS3-FRB::hphNT1, YJL081C-GFP::HIS3* | This study | 0 | 0 |
| OGYSGA1830 | *MATa, his3Δ1, leu2Δ0, ura3Δ0, LYS+, can1Δ::STE2pr-LEU2, lyp1Δ::, tor1-1, fpr1Δ::klURA, Tub4-(6)-RFP-(24)-FKBP::natNT2, VPS3-FRB::hphNT1, YDL020C-GFP::HIS3* | This study | 0.44 | 0 |
| OGYSGA1831 | *MATa, his3Δ1, leu2Δ0, ura3Δ0, LYS+, can1Δ::STE2pr-LEU2, lyp1Δ::, tor1-1, fpr1Δ::klURA, Tub4-(6)-RFP-(24)-FKBP::natNT2, VPS3-FRB::hphNT1, YPL086C-GFP::HIS3* | This study | 0 | 0 |
| OGYSGA1832 | *MATa, his3Δ1, leu2Δ0, ura3Δ0, LYS+, can1Δ::STE2pr-LEU2, lyp1Δ::, tor1-1, fpr1Δ::klURA, Tub4-(6)-RFP-(24)-FKBP::natNT2, VPS3-FRB::hphNT1, YOL115W-GFP::HIS3* | This study | 0 | 5.7 |
| OGYSGA1833 | *MATa, his3Δ1, leu2Δ0, ura3Δ0, LYS+, can1Δ::STE2pr-LEU2, lyp1Δ::, tor1-1, fpr1Δ::klURA, Tub4-(6)-RFP-(24)-FKBP::natNT2, VPS3-FRB::hphNT1, YNL021W-GFP::HIS3* | This study | 0.83 | 0.52 |
| OGYSGA1834 | *MATa, his3Δ1, leu2Δ0, ura3Δ0, LYS+, can1Δ::STE2pr-LEU2, lyp1Δ::, tor1-1, fpr1Δ::klURA, Tub4-(6)-RFP-(24)-FKBP::natNT2, VPS3-FRB::hphNT1, YHL002W-GFP::HIS3* | This study | 0 | 1072.33 |
| OGYSGA1835 | *MATa, his3Δ1, leu2Δ0, ura3Δ0, LYS+, can1Δ::STE2pr-LEU2, lyp1Δ::, tor1-1, fpr1Δ::klURA, Tub4-(6)-RFP-(24)-FKBP::natNT2, VPS3-FRB::hphNT1, YDR150W-GFP::HIS3* | This study | 0 | 0 |
| OGYSGA1837 | *MATa, his3Δ1, leu2Δ0, ura3Δ0, LYS+, can1Δ::STE2pr-LEU2, lyp1Δ::, tor1-1, fpr1Δ::klURA, Tub4-(6)-RFP-(24)-FKBP::natNT2, VPS3-FRB::hphNT1, YPL084W-GFP::HIS3* | This study | 273.4 | 122.09 |
| OGYSGA1838 | *MATa, his3Δ1, leu2Δ0, ura3Δ0, LYS+, can1Δ::STE2pr-LEU2, lyp1Δ::, tor1-1, fpr1Δ::klURA, Tub4-(6)-RFP-(24)-FKBP::natNT2, VPS3-FRB::hphNT1, YNL323W-GFP::HIS3* | This study | 0 | 0.04 |
| OGYSGA1839 | *MATa, his3Δ1, leu2Δ0, ura3Δ0, LYS+, can1Δ::STE2pr-LEU2, lyp1Δ::, tor1-1, fpr1Δ::klURA, Tub4-(6)-RFP-(24)-FKBP::natNT2, VPS3-FRB::hphNT1, YMR236W-GFP::HIS3* | This study | 0.87 | 0.98 |
| OGYSGA1840 | *MATa, his3Δ1, leu2Δ0, ura3Δ0, LYS+, can1Δ::STE2pr-LEU2, lyp1Δ::, tor1-1, fpr1Δ::klURA, Tub4-(6)-RFP-(24)-FKBP::natNT2, VPS3-FRB::hphNT1, YMR161W-GFP::HIS3* | This study | 0.72 | 0.24 |
| OGYSGA1841 | *MATa, his3Δ1, leu2Δ0, ura3Δ0, LYS+, can1Δ::STE2pr-LEU2, lyp1Δ::, tor1-1, fpr1Δ::klURA, Tub4-(6)-RFP-(24)-FKBP::natNT2, VPS3-FRB::hphNT1, YGR122W-GFP::HIS3* | This study | 0.39 | 4.81 |
| OGYSGA1842 | *MATa, his3Δ1, leu2Δ0, ura3Δ0, LYS+, can1Δ::STE2pr-LEU2, lyp1Δ::, tor1-1, fpr1Δ::klURA, Tub4-(6)-RFP-(24)-FKBP::natNT2, VPS3-FRB::hphNT1, YCR044C-GFP::HIS3* | This study | 0.5 | 0.59 |
| OGYSGA1843 | *MATa, his3Δ1, leu2Δ0, ura3Δ0, LYS+, can1Δ::STE2pr-LEU2, lyp1Δ::, tor1-1, fpr1Δ::klURA, Tub4-(6)-RFP-(24)-FKBP::natNT2, VPS3-FRB::hphNT1, YMR022W-GFP::HIS3* | This study | 1.1 | 1.41 |
| OGYSGA1844 | *MATa, his3Δ1, leu2Δ0, ura3Δ0, LYS+, can1Δ::STE2pr-LEU2, lyp1Δ::, tor1-1, fpr1Δ::klURA, Tub4-(6)-RFP-(24)-FKBP::natNT2, VPS3-FRB::hphNT1, YJL062W-GFP::HIS3* | This study | 0.56 | 0.46 |
| OGYSGA1846 | *MATa, his3Δ1, leu2Δ0, ura3Δ0, LYS+, can1Δ::STE2pr-LEU2, lyp1Δ::, tor1-1, fpr1Δ::klURA, Tub4-(6)-RFP-(24)-FKBP::natNT2, VPS3-FRB::hphNT1, YML097C-GFP::HIS3* | This study | 10.73 | 0 |
| OGYSGA1847 | *MATa, his3Δ1, leu2Δ0, ura3Δ0, LYS+, can1Δ::STE2pr-LEU2, lyp1Δ::, tor1-1, fpr1Δ::klURA, Tub4-(6)-RFP-(24)-FKBP::natNT2, VAM6-FRB::hphNT1, YNL243W-GFP::HIS3* | This study | 0.89 | 0.94 |
| OGYSGA1848 | *MATa, his3Δ1, leu2Δ0, ura3Δ0, LYS+, can1Δ::STE2pr-LEU2, lyp1Δ::, tor1-1, fpr1Δ::klURA, Tub4-(6)-RFP-(24)-FKBP::natNT2, VAM6-FRB::hphNT1, YDL185W-GFP::HIS3* | This study | 1.75 | 0 |
| OGYSGA1849 | *MATa, his3Δ1, leu2Δ0, ura3Δ0, LYS+, can1Δ::STE2pr-LEU2, lyp1Δ::, tor1-1, fpr1Δ::klURA, Tub4-(6)-RFP-(24)-FKBP::natNT2, VAM6-FRB::hphNT1, YOR244W-GFP::HIS3* | This study | 0.29 | 0.91 |
| OGYSGA1850 | *MATa, his3Δ1, leu2Δ0, ura3Δ0, LYS+, can1Δ::STE2pr-LEU2, lyp1Δ::, tor1-1, fpr1Δ::klURA, Tub4-(6)-RFP-(24)-FKBP::natNT2, VAM6-FRB::hphNT1, YGL020C-GFP::HIS3* | This study | 0.51 | 0.43 |
| OGYSGA1851 | *MATa, his3Δ1, leu2Δ0, ura3Δ0, LYS+, can1Δ::STE2pr-LEU2, lyp1Δ::, tor1-1, fpr1Δ::klURA, Tub4-(6)-RFP-(24)-FKBP::natNT2, VAM6-FRB::hphNT1, YER083C-GFP::HIS3* | This study | 0.36 | 0.82 |
| OGYSGA1852 | *MATa, his3Δ1, leu2Δ0, ura3Δ0, LYS+, can1Δ::STE2pr-LEU2, lyp1Δ::, tor1-1, fpr1Δ::klURA, Tub4-(6)-RFP-(24)-FKBP::natNT2, VAM6-FRB::hphNT1, YJL115W-GFP::HIS3* | This study | 0 | 0.78 |
| OGYSGA1853 | *MATa, his3Δ1, leu2Δ0, ura3Δ0, LYS+, can1Δ::STE2pr-LEU2, lyp1Δ::, tor1-1, fpr1Δ::klURA, Tub4-(6)-RFP-(24)-FKBP::natNT2, VAM6-FRB::hphNT1, YKL113C-GFP::HIS3* | This study | 0 | 2.12 |
| OGYSGA1854 | *MATa, his3Δ1, leu2Δ0, ura3Δ0, LYS+, can1Δ::STE2pr-LEU2, lyp1Δ::, tor1-1, fpr1Δ::klURA, Tub4-(6)-RFP-(24)-FKBP::natNT2, VAM6-FRB::hphNT1, YGL095C-GFP::HIS3* | This study | 0.23 | 1.01 |
| OGYSGA1855 | *MATa, his3Δ1, leu2Δ0, ura3Δ0, LYS+, can1Δ::STE2pr-LEU2, lyp1Δ::, tor1-1, fpr1Δ::klURA, Tub4-(6)-RFP-(24)-FKBP::natNT2, VAM6-FRB::hphNT1, YFL024C-GFP::HIS3* | This study | 4.96 | 0.93 |
| OGYSGA1856 | *MATa, his3Δ1, leu2Δ0, ura3Δ0, LYS+, can1Δ::STE2pr-LEU2, lyp1Δ::, tor1-1, fpr1Δ::klURA, Tub4-(6)-RFP-(24)-FKBP::natNT2, VAM6-FRB::hphNT1, YDL040C-GFP::HIS3* | This study | 0 | 5.71 |
| OGYSGA1857 | *MATa, his3Δ1, leu2Δ0, ura3Δ0, LYS+, can1Δ::STE2pr-LEU2, lyp1Δ::, tor1-1, fpr1Δ::klURA, Tub4-(6)-RFP-(24)-FKBP::natNT2, VAM6-FRB::hphNT1, YGL116W-GFP::HIS3* | This study | 0 | 0.59 |
| OGYSGA1858 | *MATa, his3Δ1, leu2Δ0, ura3Δ0, LYS+, can1Δ::STE2pr-LEU2, lyp1Δ::, tor1-1, fpr1Δ::klURA, Tub4-(6)-RFP-(24)-FKBP::natNT2, VAM6-FRB::hphNT1, YPL153C-GFP::HIS3* | This study | 5.71 | 11.33 |
| OGYSGA1859 | *MATa, his3Δ1, leu2Δ0, ura3Δ0, LYS+, can1Δ::STE2pr-LEU2, lyp1Δ::, tor1-1, fpr1Δ::klURA, Tub4-(6)-RFP-(24)-FKBP::natNT2, VAM6-FRB::hphNT1, YHR090C-GFP::HIS3* | This study | 0.14 | 0.26 |
| OGYSGA1860 | *MATa, his3Δ1, leu2Δ0, ura3Δ0, LYS+, can1Δ::STE2pr-LEU2, lyp1Δ::, tor1-1, fpr1Δ::klURA, Tub4-(6)-RFP-(24)-FKBP::natNT2, VAM6-FRB::hphNT1, YCL016C-GFP::HIS3* | This study | 1.95 | 2.41 |
| OGYSGA1861 | *MATa, his3Δ1, leu2Δ0, ura3Δ0, LYS+, can1Δ::STE2pr-LEU2, lyp1Δ::, tor1-1, fpr1Δ::klURA, Tub4-(6)-RFP-(24)-FKBP::natNT2, VAM6-FRB::hphNT1, YGL058W-GFP::HIS3* | This study | 0 | 0.78 |
| OGYSGA1862 | *MATa, his3Δ1, leu2Δ0, ura3Δ0, LYS+, can1Δ::STE2pr-LEU2, lyp1Δ::, tor1-1, fpr1Δ::klURA, Tub4-(6)-RFP-(24)-FKBP::natNT2, VAM6-FRB::hphNT1, YHR013C-GFP::HIS3* | This study | 0 | 0 |
| OGYSGA1863 | *MATa, his3Δ1, leu2Δ0, ura3Δ0, LYS+, can1Δ::STE2pr-LEU2, lyp1Δ::, tor1-1, fpr1Δ::klURA, Tub4-(6)-RFP-(24)-FKBP::natNT2, VAM6-FRB::hphNT1, YMR078C-GFP::HIS3* | This study | 1.36 | 1.22 |
| OGYSGA1864 | *MATa, his3Δ1, leu2Δ0, ura3Δ0, LYS+, can1Δ::STE2pr-LEU2, lyp1Δ::, tor1-1, fpr1Δ::klURA, Tub4-(6)-RFP-(24)-FKBP::natNT2, VAM6-FRB::hphNT1, YHR200W-GFP::HIS3* | This study | 1.17 | 0.94 |
| OGYSGA1865 | *MATa, his3Δ1, leu2Δ0, ura3Δ0, LYS+, can1Δ::STE2pr-LEU2, lyp1Δ::, tor1-1, fpr1Δ::klURA, Tub4-(6)-RFP-(24)-FKBP::natNT2, VAM6-FRB::hphNT1, YDR023W-GFP::HIS3* | This study | 0 | 0 |
| OGYSGA1866 | *MATa, his3Δ1, leu2Δ0, ura3Δ0, LYS+, can1Δ::STE2pr-LEU2, lyp1Δ::, tor1-1, fpr1Δ::klURA, Tub4-(6)-RFP-(24)-FKBP::natNT2, VAM6-FRB::hphNT1, YDL074C-GFP::HIS3* | This study | 0 | 0 |
| OGYSGA1867 | *MATa, his3Δ1, leu2Δ0, ura3Δ0, LYS+, can1Δ::STE2pr-LEU2, lyp1Δ::, tor1-1, fpr1Δ::klURA, Tub4-(6)-RFP-(24)-FKBP::natNT2, VAM6-FRB::hphNT1, YPL055C-GFP::HIS3* | This study | 0 | 1.72 |
| OGYSGA1868 | *MATa, his3Δ1, leu2Δ0, ura3Δ0, LYS+, can1Δ::STE2pr-LEU2, lyp1Δ::, tor1-1, fpr1Δ::klURA, Tub4-(6)-RFP-(24)-FKBP::natNT2, VAM6-FRB::hphNT1, YBR164C-GFP::HIS3* | This study | 0.32 | 0.34 |
| OGYSGA1869 | *MATa, his3Δ1, leu2Δ0, ura3Δ0, LYS+, can1Δ::STE2pr-LEU2, lyp1Δ::, tor1-1, fpr1Δ::klURA, Tub4-(6)-RFP-(24)-FKBP::natNT2, VAM6-FRB::hphNT1, YJL092W-GFP::HIS3* | This study | 0 | 0 |
| OGYSGA1870 | *MATa, his3Δ1, leu2Δ0, ura3Δ0, LYS+, can1Δ::STE2pr-LEU2, lyp1Δ::, tor1-1, fpr1Δ::klURA, Tub4-(6)-RFP-(24)-FKBP::natNT2, VAM6-FRB::hphNT1, YOR141C-GFP::HIS3* | This study | 0 | 0 |

| | | | | |
|---|---|---|---|---|
| OGYSGA1871 | *MATa, his3Δ1, leu2Δ0, ura3Δ0, LYS+, can1Δ::STE2pr-LEU2, lyp1Δ::, tor1-1, fpr1Δ::klURA, Tub4-(6)-RFP-(24)-FKBP::natNT2, VAM6-FRB::hphNT1, YGR106C-GFP::HIS3* | This study | 3.36 | 1.28 |
| OGYSGA1872 | *MATa, his3Δ1, leu2Δ0, ura3Δ0, LYS+, can1Δ::STE2pr-LEU2, lyp1Δ::, tor1-1, fpr1Δ::klURA, Tub4-(6)-RFP-(24)-FKBP::natNT2, VAM6-FRB::hphNT1, YGR198W-GFP::HIS3* | This study | 1.05 | 0.4 |
| OGYSGA1873 | *MATa, his3Δ1, leu2Δ0, ura3Δ0, LYS+, can1Δ::STE2pr-LEU2, lyp1Δ::, tor1-1, fpr1Δ::klURA, Tub4-(6)-RFP-(24)-FKBP::natNT2, VAM6-FRB::hphNT1, YGR105W-GFP::HIS3* | This study | 2.91 | 0 |
| OGYSGA1874 | *MATa, his3Δ1, leu2Δ0, ura3Δ0, LYS+, can1Δ::STE2pr-LEU2, lyp1Δ::, tor1-1, fpr1Δ::klURA, Tub4-(6)-RFP-(24)-FKBP::natNT2, VAM6-FRB::hphNT1, YDR150W-GFP::HIS3* | This study | 0 | 1.58 |
| OGYSGA1875 | *MATa, his3Δ1, leu2Δ0, ura3Δ0, LYS+, can1Δ::STE2pr-LEU2, lyp1Δ::, tor1-1, fpr1Δ::klURA, Tub4-(6)-RFP-(24)-FKBP::natNT2, VAM6-FRB::hphNT1, YGR061C-GFP::HIS3* | This study | 0 | 0 |
| OGYSGA1876 | *MATa, his3Δ1, leu2Δ0, ura3Δ0, LYS+, can1Δ::STE2pr-LEU2, lyp1Δ::, tor1-1, fpr1Δ::klURA, Tub4-(6)-RFP-(24)-FKBP::natNT2, VAM6-FRB::hphNT1, YBR152W-GFP::HIS3* | This study | 0 | 0 |
| OGYSGA1877 | *MATa, his3Δ1, leu2Δ0, ura3Δ0, LYS+, can1Δ::STE2pr-LEU2, lyp1Δ::, tor1-1, fpr1Δ::klURA, Tub4-(6)-RFP-(24)-FKBP::natNT2, VAM6-FRB::hphNT1, YLL029W-GFP::HIS3* | This study | 0 | 0 |
| OGYSGA1878 | *MATa, his3Δ1, leu2Δ0, ura3Δ0, LYS+, can1Δ::STE2pr-LEU2, lyp1Δ::, tor1-1, fpr1Δ::klURA, Tub4-(6)-RFP-(24)-FKBP::natNT2, VAM6-FRB::hphNT1, YGR002C-GFP::HIS3* | This study | 1.18 | 0.89 |
| OGYSGA1879 | *MATa, his3Δ1 leu2Δ0 ura3Δ0 LYS+, can1Δ::STE2pr-LEU2, lyp1Δ::, tor1-1, fpr1Δ::klURA, Tub4-(6)-RFP-(24)-FKBP::natNT2, VAM6-FRB::hphNT1, YJL081C-GFP::HIS3* | This study | 0 | 0 |
| OGYSGA1880 | *MATa, his3Δ1 leu2Δ0 ura3Δ0 LYS+, can1Δ::STE2pr-LEU2, lyp1Δ::, tor1-1, fpr1Δ::klURA, Tub4-(6)-RFP-(24)-FKBP::natNT2, VAM6-FRB::hphNT1, YLR114C-GFP::HIS3* | This study | 0 | 0 |
| OGYSGA1881 | *MATa, his3Δ1 leu2Δ0 ura3Δ0 LYS+, can1Δ::STE2pr-LEU2, lyp1Δ::, tor1-1, fpr1Δ::klURA, Tub4-(6)-RFP-(24)-FKBP::natNT2, VAM6-FRB::hphNT1, YJR043C-GFP::HIS3* | This study | 0 | 0 |
| OGYSGA1882 | *MATa, his3Δ1 leu2Δ0 ura3Δ0 LYS+, can1Δ::STE2pr-LEU2, lyp1Δ::, tor1-1, fpr1Δ::klURA, Tub4-(6)-RFP-(24)-FKBP::natNT2, VAM6-FRB::hphNT1, YLR153C-GFP::HIS3* | This study | 1.05 | 0 |
| OGYSGA1883 | *MATa, his3Δ1 leu2Δ0 ura3Δ0 LYS+, can1Δ::STE2pr-LEU2, lyp1Δ::, tor1-1, fpr1Δ::klURA, Tub4-(6)-RFP-(24)-FKBP::natNT2, VAM6-FRB::hphNT1, YNL056W-GFP::HIS3* | This study | 0 | 0 |
| OGYSGA1884 | *MATa, his3Δ1 leu2Δ0 ura3Δ0 LYS+, can1Δ::STE2pr-LEU2, lyp1Δ::, tor1-1, fpr1Δ::klURA, Tub4-(6)-RFP-(24)-FKBP::natNT2, VAM6-FRB::hphNT1, YGR155W-GFP::HIS3* | This study | 0.24 | 0 |
| OGYSGA1885 | *MATa, his3Δ1 leu2Δ0 ura3Δ0 LYS+, can1Δ::STE2pr-LEU2, lyp1Δ::, tor1-1, fpr1Δ::klURA, Tub4-(6)-RFP-(24)-FKBP::natNT2, VAM6-FRB::hphNT1, YNL032W-GFP::HIS3* | This study | 0 | 0 |
| OGYSGA1886 | *MATa, his3Δ1 leu2Δ0 ura3Δ0 LYS+, can1Δ::STE2pr-LEU2, lyp1Δ::, tor1-1, fpr1Δ::klURA, Tub4-(6)-RFP-(24)-FKBP::natNT2, VAM6-FRB::hphNT1, YOL115W-GFP::HIS3* | This study | 0 | 0 |
| OGYSGA1887 | *MATa, his3Δ1 leu2Δ0 ura3Δ0 LYS+, can1Δ::STE2pr-LEU2, lyp1Δ::, tor1-1, fpr1Δ::klURA, Tub4-(6)-RFP-(24)-FKBP::natNT2, VAM6-FRB::hphNT1, YJL154C-GFP::HIS3* | This study | 5.03 | 5.3 |
| OGYSGA1888 | *MATa, his3Δ1 leu2Δ0 ura3Δ0 LYS+, can1Δ::STE2pr-LEU2, lyp1Δ::, tor1-1, fpr1Δ::klURA, Tub4-(6)-RFP-(24)-FKBP::natNT2, VAM6-FRB::hphNT1, YOR069W-GFP::HIS3* | This study | 5.66 | 9.07 |
| OGYSGA1889 | *MATa, his3Δ1 leu2Δ0 ura3Δ0 LYS+, can1Δ::STE2pr-LEU2, lyp1Δ::, tor1-1, fpr1Δ::klURA, Tub4-(6)-RFP-(24)-FKBP::natNT2, VAM6-FRB::hphNT1, YKL009W-GFP::HIS3* | This study | 0.08 | 1.03 |
| OGYSGA1890 | *MATa, his3Δ1 leu2Δ0 ura3Δ0 LYS+, can1Δ::STE2pr-LEU2, lyp1Δ::, tor1-1, fpr1Δ::klURA, Tub4-(6)-RFP-(24)-FKBP::natNT2, VAM6-FRB::hphNT1, YNL021W-GFP::HIS3* | This study | 0.73 | 2.09 |
| OGYSGA1891 | *MATa, his3Δ1 leu2Δ0 ura3Δ0 LYS+, can1Δ::STE2pr-LEU2, lyp1Δ::, tor1-1, fpr1Δ::klURA, Tub4-(6)-RFP-(24)-FKBP::natNT2, VAM6-FRB::hphNT1, YHR012W-GFP::HIS3* | This study | 0 | 0 |
| OGYSGA1892 | *MATa, his3Δ1 leu2Δ0 ura3Δ0 LYS+, can1Δ::STE2pr-LEU2, lyp1Δ::, tor1-1, fpr1Δ::klURA, Tub4-(6)-RFP-(24)-FKBP::natNT2, VAM6-FRB::hphNT1, YGL061C-GFP::HIS3* | This study | 0 | 0 |
| OGYSGA1893 | *MATa, his3Δ1 leu2Δ0 ura3Δ0 LYS+, can1Δ::STE2pr-LEU2, lyp1Δ::, tor1-1, fpr1Δ::klURA, Tub4-(6)-RFP-(24)-FKBP::natNT2, VAM6-FRB::hphNT1, YPL145C-GFP::HIS3* | This study | 7.47 | 0 |
| OGYSGA1894 | *MATa, his3Δ1 leu2Δ0 ura3Δ0 LYS+, can1Δ::STE2pr-LEU2, lyp1Δ::, tor1-1, fpr1Δ::klURA, Tub4-(6)-RFP-(24)-FKBP::natNT2, VAM6-FRB::hphNT1, YDL020C-GFP::HIS3* | This study | 1.95 | 0 |
| OGYSGA1895 | *MATa, his3Δ1 leu2Δ0 ura3Δ0 LYS+, can1Δ::STE2pr-LEU2, lyp1Δ::, tor1-1, fpr1Δ::klURA, Tub4-(6)-RFP-(24)-FKBP::natNT2, VAM6-FRB::hphNT1, YLR190W-GFP::HIS3* | This study | 0 | 0.32 |
| OGYSGA1896 | *MATa, his3Δ1 leu2Δ0 ura3Δ0 LYS+, can1Δ::STE2pr-LEU2, lyp1Δ::, tor1-1, fpr1Δ::klURA, Tub4-(6)-RFP-(24)-FKBP::natNT2, VAM6-FRB::hphNT1, YJL004C-GFP::HIS3* | This study | 0.19 | 0.26 |
| OGYSGA1897 | *MATa, his3Δ1 leu2Δ0 ura3Δ0 LYS+, can1Δ::STE2pr-LEU2, lyp1Δ::, tor1-1, fpr1Δ::klURA, Tub4-(6)-RFP-(24)-FKBP::natNT2, VAM6-FRB::hphNT1, YAL010C-GFP::HIS3* | This study | 0.11 | 0.07 |
| OGYSGA1898 | *MATa, his3Δ1 leu2Δ0 ura3Δ0 LYS+, can1Δ::STE2pr-LEU2, lyp1Δ::, tor1-1, fpr1Δ::klURA, Tub4-(6)-RFP-(24)-FKBP::natNT2, VAM6-FRB::hphNT1, YPL051W-GFP::HIS3* | This study | 0.03 | 4.1 |
| OGYSGA1899 | *MATa, his3Δ1 leu2Δ0 ura3Δ0 LYS+, can1Δ::STE2pr-LEU2, lyp1Δ::, tor1-1, fpr1Δ::klURA, Tub4-(6)-RFP-(24)-FKBP::natNT2, VAM6-FRB::hphNT1, YJL166W-GFP::HIS3* | This study | 0.36 | 0.53 |
| OGYSGA1935 | *MATa, his3Δ1 leu2Δ0 ura3Δ0 LYS+, can1Δ::STE2pr-LEU2, lyp1Δ::, tor1-1, fpr1Δ::klURA, Tub4-(6)-RFP-(24)-FKBP::natNT2, TRS130-FRB::hphNT1, YMR307W-GFP::HIS3* | This study | 0.12 | 0 |
| OGYSGA1936 | *MATa, his3Δ1 leu2Δ0 ura3Δ0 LYS+, can1Δ::STE2pr-LEU2, lyp1Δ::, tor1-1, fpr1Δ::klURA, Tub4-(6)-RFP-(24)-FKBP::natNT2, TRS130-FRB::hphNT1, YNL267W-GFP::HIS3* | This study | 0.12 | 0 |
| OGYSGA1937 | *MATa, his3Δ1 leu2Δ0 ura3Δ0 LYS+, can1Δ::STE2pr-LEU2, lyp1Δ::, tor1-1, fpr1Δ::klURA, Tub4-(6)-RFP-(24)-FKBP::natNT2, TRS130-FRB::hphNT1, YGL210W-GFP::HIS3* | This study | 0 | 0 |
| OGYSGA1938 | *MATa, his3Δ1 leu2Δ0 ura3Δ0 LYS+, can1Δ::STE2pr-LEU2, lyp1Δ::, tor1-1, fpr1Δ::klURA, Tub4-(6)-RFP-(24)-FKBP::natNT2, TRS130-FRB::hphNT1, YOR216C-GFP::HIS3* | This study | 0.4 | 0.45 |
| OGYSGA1939 | *MATa, his3Δ1 leu2Δ0 ura3Δ0 LYS+, can1Δ::STE2pr-LEU2, lyp1Δ::, tor1-1, fpr1Δ::klURA, Tub4-(6)-RFP-(24)-FKBP::natNT2, TRS130-FRB::hphNT1, YMR123W-GFP::HIS3* | This study | 0.91 | 0 |
| OGYSGA1940 | *MATa, his3Δ1 leu2Δ0 ura3Δ0 LYS+, can1Δ::STE2pr-LEU2, lyp1Δ::, tor1-1, fpr1Δ::klURA, Tub4-(6)-RFP-(24)-FKBP::natNT2, TRS130-FRB::hphNT1, YBR164C-GFP::HIS3* | This study | 0 | 0 |
| OGYSGA1941 | *MATa, his3Δ1 leu2Δ0 ura3Δ0 LYS+, can1Δ::STE2pr-LEU2, lyp1Δ::, tor1-1, fpr1Δ::klURA, Tub4-(6)-RFP-(24)-FKBP::natNT2, TRS130-FRB::hphNT1, YLR268W-GFP::HIS3* | This study | 1.74 | 0.88 |
| OGYSGA1942 | *MATa, his3Δ1 leu2Δ0 ura3Δ0 LYS+, can1Δ::STE2pr-LEU2, lyp1Δ::, tor1-1, fpr1Δ::klURA, Tub4-(6)-RFP-(24)-FKBP::natNT2, TRS130-FRB::hphNT1, YJR118C-GFP::HIS3* | This study | 1.9 | 0.28 |
| OGYSGA1943 | *MATa, his3Δ1 leu2Δ0 ura3Δ0 LYS+, can1Δ::STE2pr-LEU2, lyp1Δ::, tor1-1, fpr1Δ::klURA, Tub4-(6)-RFP-(24)-FKBP::natNT2, TRS130-FRB::hphNT1, YMR272C-GFP::HIS3* | This study | 1.41 | 2.92 |
| OGYSGA1944 | *MATa, his3Δ1 leu2Δ0 ura3Δ0 LYS+, can1Δ::STE2pr-LEU2, lyp1Δ::, tor1-1, fpr1Δ::klURA, Tub4-(6)-RFP-(24)-FKBP::natNT2, TRS130-FRB::hphNT1, YLR262C-GFP::HIS3* | This study | 0 | 0 |
| OGYSGA1945 | *MATa, his3Δ1 leu2Δ0 ura3Δ0 LYS+, can1Δ::STE2pr-LEU2, lyp1Δ::, tor1-1, fpr1Δ::klURA, Tub4-(6)-RFP-(24)-FKBP::natNT2, TRS130-FRB::hphNT1, YMR079W-GFP::HIS3* | This study | 0 | 0 |

| | | | | |
|---|---|---|---|---|
| OGYSGA1946 | *MATa, his3Δ1 leu2Δ0 ura3Δ0 LYS+, can1Δ::STE2pr-LEU2, lyp1Δ::, tor1-1, fpr1Δ::klURA, Tub4-(6)-RFP-(24)-FKBP::natNT2, TRS130-FRB::hphNT1, YPL051W-GFP::HIS3* | This study | 0 | 5.94 |
| OGYSGA1947 | *MATa, his3Δ1 leu2Δ0 ura3Δ0 LYS+, can1Δ::STE2pr-LEU2, lyp1Δ::, tor1-1, fpr1Δ::klURA, Tub4-(6)-RFP-(24)-FKBP::natNT2, TRS130-FRB::hphNT1, YGL054C-GFP::HIS3* | This study | 3.69 | 0 |
| OGYSGA1948 | *MATa, his3Δ1 leu2Δ0 ura3Δ0 LYS+, can1Δ::STE2pr-LEU2, lyp1Δ::, tor1-1, fpr1Δ::klURA, Tub4-(6)-RFP-(24)-FKBP::natNT2, TRS130-FRB::hphNT1, YPL234C-GFP::HIS3* | This study | 4.57 | 1.57 |
| OGYSGA1949 | *MATa, his3Δ1 leu2Δ0 ura3Δ0 LYS+, can1Δ::STE2pr-LEU2, lyp1Δ::, tor1-1, fpr1Δ::klURA, Tub4-(6)-RFP-(24)-FKBP::natNT2, TRS130-FRB::hphNT1, YKL119C-GFP::HIS3* | This study | 0.26 | 2.74 |
| OGYSGA1950 | *MATa, his3Δ1 leu2Δ0 ura3Δ0 LYS+, can1Δ::STE2pr-LEU2, lyp1Δ::, tor1-1, fpr1Δ::klURA, Tub4-(6)-RFP-(24)-FKBP::natNT2, TRS130-FRB::hphNT1, YAL026C-GFP::HIS3* | This study | 7.23 | 5.68 |
| OGYSGA1951 | *MATa, his3Δ1 leu2Δ0 ura3Δ0 LYS+, can1Δ::STE2pr-LEU2, lyp1Δ::, tor1-1, fpr1Δ::klURA, Tub4-(6)-RFP-(24)-FKBP::natNT2, TRS130-FRB::hphNT1, YOR070C-GFP::HIS3* | This study | 0 | 0 |
| OGYSGA1952 | *MATa, his3Δ1 leu2Δ0 ura3Δ0 LYS+, can1Δ::STE2pr-LEU2, lyp1Δ::, tor1-1, fpr1Δ::klURA, Tub4-(6)-RFP-(24)-FKBP::natNT2, TRS130-FRB::hphNT1, YKR001C-GFP::HIS3* | This study | 11.65 | 3.27 |
| OGYSGA1953 | *MATa, his3Δ1 leu2Δ0 ura3Δ0 LYS+, can1Δ::STE2pr-LEU2, lyp1Δ::, tor1-1, fpr1Δ::klURA, Tub4-(6)-RFP-(24)-FKBP::natNT2, TRS130-FRB::hphNT1, YPL057C-GFP::HIS3* | This study | 7.14 | 3.4 |
| OGYSGA1954 | *MATa, his3Δ1 leu2Δ0 ura3Δ0 LYS+, can1Δ::STE2pr-LEU2, lyp1Δ::, tor1-1, fpr1Δ::klURA, Tub4-(6)-RFP-(24)-FKBP::natNT2, TRS130-FRB::hphNT1, YLR350W-GFP::HIS3* | This study | 2.52 | 0 |
| OGYSGA1955 | *MATa, his3Δ1 leu2Δ0 ura3Δ0 LYS+, can1Δ::STE2pr-LEU2, lyp1Δ::, tor1-1, fpr1Δ::klURA, Tub4-(6)-RFP-(24)-FKBP::natNT2, TRS130-FRB::hphNT1, YMR190C-GFP::HIS3* | This study | 11.92 | 0 |
| OGYSGA1956 | *MATa, his3Δ1 leu2Δ0 ura3Δ0 LYS+, can1Δ::STE2pr-LEU2, lyp1Δ::, tor1-1, fpr1Δ::klURA, Tub4-(6)-RFP-(24)-FKBP::natNT2, TRS130-FRB::hphNT1, YLR039C-GFP::HIS3* | This study | 0.92 | 2.5 |
| OGYSGA1957 | *MATa, his3Δ1 leu2Δ0 ura3Δ0 LYS+, can1Δ::STE2pr-LEU2, lyp1Δ::, tor1-1, fpr1Δ::klURA, Tub4-(6)-RFP-(24)-FKBP::natNT2, TRS130-FRB::hphNT1, YIL004C-GFP::HIS3* | This study | 0 | 0 |
| OGYSGA1958 | *MATa, his3Δ1 leu2Δ0 ura3Δ0 LYS+, can1Δ::STE2pr-LEU2, lyp1Δ::, tor1-1, fpr1Δ::klURA, Tub4-(6)-RFP-(24)-FKBP::natNT2, TRS130-FRB::hphNT1, YBR080C-GFP::HIS3* | This study | 1 | 45.08 |
| OGYSGA1959 | *MATa, his3Δ1 leu2Δ0 ura3Δ0 LYS+, can1Δ::STE2pr-LEU2, lyp1Δ::, tor1-1, fpr1Δ::klURA, Tub4-(6)-RFP-(24)-FKBP::natNT2, TRS130-FRB::hphNT1, YDL226C-GFP::HIS3* | This study | 1.03 | 0 |
| OGYSGA1960 | *MATa, his3Δ1 leu2Δ0 ura3Δ0 LYS+, can1Δ::STE2pr-LEU2, lyp1Δ::, tor1-1, fpr1Δ::klURA, Tub4-(6)-RFP-(24)-FKBP::natNT2, TRS130-FRB::hphNT1, YJL123C-GFP::HIS3* | This study | 0.49 | 0.53 |
| OGYSGA1961 | *MATa, his3Δ1 leu2Δ0 ura3Δ0 LYS+, can1Δ::STE2pr-LEU2, lyp1Δ::, tor1-1, fpr1Δ::klURA, Tub4-(6)-RFP-(24)-FKBP::natNT2, TRS130-FRB::hphNT1, YOR144C-GFP::HIS3* | This study | 0 | 0 |
| OGYSGA1962 | *MATa, his3Δ1 leu2Δ0 ura3Δ0 LYS+, can1Δ::STE2pr-LEU2, lyp1Δ::, tor1-1, fpr1Δ::klURA, Tub4-(6)-RFP-(24)-FKBP::natNT2, TRS130-FRB::hphNT1, YDR372C-GFP::HIS3* | This study | 0 | 0 |
| OGYSGA1963 | *MATa, his3Δ1 leu2Δ0 ura3Δ0 LYS+, can1Δ::STE2pr-LEU2, lyp1Δ::, tor1-1, fpr1Δ::klURA, Tub4-(6)-RFP-(24)-FKBP::natNT2, TRS130-FRB::hphNT1, YGL180W-GFP::HIS3* | This study | 0 | 0 |
| OGYSGA1964 | *MATa, his3Δ1 leu2Δ0 ura3Δ0 LYS+, can1Δ::STE2pr-LEU2, lyp1Δ::, tor1-1, fpr1Δ::klURA, Tub4-(6)-RFP-(24)-FKBP::natNT2, TRS130-FRB::hphNT1, YLL040C-GFP::HIS3* | This study | 6.2 | 5.73 |
| OGYSGA1965 | *MATa, his3Δ1 leu2Δ0 ura3Δ0 LYS+, can1Δ::STE2pr-LEU2, lyp1Δ::, tor1-1, fpr1Δ::klURA, Tub4-(6)-RFP-(24)-FKBP::natNT2, TRS130-FRB::hphNT1, YPL120W-GFP::HIS3* | This study | 5.48 | 38.75 |
| OGYSGA1966 | *MATa, his3Δ1 leu2Δ0 ura3Δ0 LYS+, can1Δ::STE2pr-LEU2, lyp1Δ::, tor1-1, fpr1Δ::klURA, Tub4-(6)-RFP-(24)-FKBP::natNT2, TRS130-FRB::hphNT1, YKL004W-GFP::HIS3* | This study | 1.84 | 0 |
| OGYSGA1967 | *MATa, his3Δ1 leu2Δ0 ura3Δ0 LYS+, can1Δ::STE2pr-LEU2, lyp1Δ::, tor1-1, fpr1Δ::klURA, Tub4-(6)-RFP-(24)-FKBP::natNT2, TRS130-FRB::hphNT1, YNL238W-GFP::HIS3* | This study | 3.18 | 3.51 |
| OGYSGA1968 | *MATa, his3Δ1 leu2Δ0 ura3Δ0 LYS+, can1Δ::STE2pr-LEU2, lyp1Δ::, tor1-1, fpr1Δ::klURA, Tub4-(6)-RFP-(24)-FKBP::natNT2, TRS130-FRB::hphNT1, YGL084C-GFP::HIS3* | This study | 1.24 | 6.13 |
| OGYSGA1969 | *MATa, his3Δ1 leu2Δ0 ura3Δ0 LYS+, can1Δ::STE2pr-LEU2, lyp1Δ::, tor1-1, fpr1Δ::klURA, Tub4-(6)-RFP-(24)-FKBP::natNT2, TRS130-FRB::hphNT1, YCR044C-GFP::HIS3* | This study | 1.36 | 0 |
| OGYSGA1970 | *MATa, his3Δ1 leu2Δ0 ura3Δ0 LYS+, can1Δ::STE2pr-LEU2, lyp1Δ::, tor1-1, fpr1Δ::klURA, Tub4-(6)-RFP-(24)-FKBP::natNT2, TRS130-FRB::hphNT1, YGL086W-GFP::HIS3* | This study | 0 | 0 |
| OGYSGA1971 | *MATa, his3Δ1 leu2Δ0 ura3Δ0 LYS+, can1Δ::STE2pr-LEU2, lyp1Δ::, tor1-1, fpr1Δ::klURA, Tub4-(6)-RFP-(24)-FKBP::natNT2, TRS130-FRB::hphNT1, YDR137W-GFP::HIS3* | This study | 1.74 | 5 |
| OGYSGA1972 | *MATa, his3Δ1 leu2Δ0 ura3Δ0 LYS+, can1Δ::STE2pr-LEU2, lyp1Δ::, tor1-1, fpr1Δ::klURA, Tub4-(6)-RFP-(24)-FKBP::natNT2, TRS130-FRB::hphNT1, YML008C-GFP::HIS3* | This study | 3.15 | 0 |
| OGYSGA1973 | *MATa, his3Δ1 leu2Δ0 ura3Δ0 LYS+, can1Δ::STE2pr-LEU2, lyp1Δ::, tor1-1, fpr1Δ::klURA, Tub4-(6)-RFP-(24)-FKBP::natNT2, TRS130-FRB::hphNT1, YPL158C-GFP::HIS3* | This study | 0 | 1.58 |
| OGYSGA1974 | *MATa, his3Δ1 leu2Δ0 ura3Δ0 LYS+, can1Δ::STE2pr-LEU2, lyp1Δ::, tor1-1, fpr1Δ::klURA, Tub4-(6)-RFP-(24)-FKBP::natNT2, TRS130-FRB::hphNT1, YNL169C-GFP::HIS3* | This study | 0 | 0.84 |
| OGYSGA1975 | *MATa, his3Δ1 leu2Δ0 ura3Δ0 LYS+, can1Δ::STE2pr-LEU2, lyp1Δ::, tor1-1, fpr1Δ::klURA, Tub4-(6)-RFP-(24)-FKBP::natNT2, TRS130-FRB::hphNT1, YLL006W-GFP::HIS3* | This study | 0 | 5.12 |
| OGYSGA1976 | *MATa, his3Δ1 leu2Δ0 ura3Δ0 LYS+, can1Δ::STE2pr-LEU2, lyp1Δ::, tor1-1, fpr1Δ::klURA, Tub4-(6)-RFP-(24)-FKBP::natNT2, TRS130-FRB::hphNT1, YML041C-GFP::HIS3* | This study | 1.5 | 0.18 |
| OGYSGA1977 | *MATa, his3Δ1 leu2Δ0 ura3Δ0 LYS+, can1Δ::STE2pr-LEU2, lyp1Δ::, tor1-1, fpr1Δ::klURA, Tub4-(6)-RFP-(24)-FKBP::natNT2, TRS130-FRB::hphNT1, YGL020C-GFP::HIS3* | This study | 0 | 0 |
| OGYSGA1978 | *MATa, his3Δ1 leu2Δ0 ura3Δ0 LYS+, can1Δ::STE2pr-LEU2, lyp1Δ::, tor1-1, fpr1Δ::klURA, Tub4-(6)-RFP-(24)-FKBP::natNT2, TRS130-FRB::hphNT1, YOL012C-GFP::HIS3* | This study | 1.22 | 3.61 |
| OGYSGA1979 | *MATa, his3Δ1 leu2Δ0 ura3Δ0 LYS+, can1Δ::STE2pr-LEU2, lyp1Δ::, tor1-1, fpr1Δ::klURA, Tub4-(6)-RFP-(24)-FKBP::natNT2, TRS85-FRB::hphNT1, YOR216C-GFP::HIS3* | This study | 2.01 | 3.58 |
| OGYSGA1980 | *MATa, his3Δ1 leu2Δ0 ura3Δ0 LYS+, can1Δ::STE2pr-LEU2, lyp1Δ::, tor1-1, fpr1Δ::klURA, Tub4-(6)-RFP-(24)-FKBP::natNT2, TRS85-FRB::hphNT1, YMR307W-GFP::HIS3* | This study | 0.36 | 0 |
| OGYSGA1981 | *MATa, his3Δ1 leu2Δ0 ura3Δ0 LYS+, can1Δ::STE2pr-LEU2, lyp1Δ::, tor1-1, fpr1Δ::klURA, Tub4-(6)-RFP-(24)-FKBP::natNT2, TRS85-FRB::hphNT1, YMR272C-GFP::HIS3* | This study | 0.98 | 1.21 |
| OGYSGA1982 | *MATa, his3Δ1 leu2Δ0 ura3Δ0 LYS+, can1Δ::STE2pr-LEU2, lyp1Δ::, tor1-1, fpr1Δ::klURA, Tub4-(6)-RFP-(24)-FKBP::natNT2, TRS85-FRB::hphNT1, YLR262C-GFP::HIS3* | This study | 0 | 0 |
| OGYSGA1983 | *MATa, his3Δ1 leu2Δ0 ura3Δ0 LYS+, can1Δ::STE2pr-LEU2, lyp1Δ::, tor1-1, fpr1Δ::klURA, Tub4-(6)-RFP-(24)-FKBP::natNT2, TRS85-FRB::hphNT1, YGL054C-GFP::HIS3* | This study | 1.59 | 0 |
| OGYSGA1984 | *MATa, his3Δ1 leu2Δ0 ura3Δ0 LYS+, can1Δ::STE2pr-LEU2, lyp1Δ::, tor1-1, fpr1Δ::klURA, Tub4-(6)-RFP-(24)-FKBP::natNT2, TRS85-FRB::hphNT1, YPL051W-GFP::HIS3* | This study | 7.16 | 3.23 |
| OGYSGA1985 | *MATa, his3Δ1 leu2Δ0 ura3Δ0 LYS+, can1Δ::STE2pr-LEU2, lyp1Δ::, tor1-1, fpr1Δ::klURA, Tub4-(6)-RFP-(24)-FKBP::natNT2, TRS85-FRB::hphNT1, YMR123W-GFP::HIS3* | This study | 0.72 | 0 |

| | | | | |
|---|---|---|---|---|
| OGYSGA1986 | *MATa, his3Δ1 leu2Δ0 ura3Δ0 LYS+, can1Δ::STE2pr-LEU2, lyp1Δ::, tor1-1, fpr1Δ::klURA, Tub4-(6)-RFP-(24)-FKBP::natNT2, TRS85-FRB::hphNT1, YBR164C-GFP::HIS3* | This study | 2.85 | 3.07 |
| OGYSGA1987 | *MATa, his3Δ1 leu2Δ0 ura3Δ0 LYS+, can1Δ::STE2pr-LEU2, lyp1Δ::, tor1-1, fpr1Δ::klURA, Tub4-(6)-RFP-(24)-FKBP::natNT2, TRS85-FRB::hphNT1, YOR070C-GFP::HIS3* | This study | 0.84 | 3.61 |
| OGYSGA1988 | *MATa, his3Δ1 leu2Δ0 ura3Δ0 LYS+, can1Δ::STE2pr-LEU2, lyp1Δ::, tor1-1, fpr1Δ::klURA, Tub4-(6)-RFP-(24)-FKBP::natNT2, TRS85-FRB::hphNT1, YKR001C-GFP::HIS3* | This study | 1.31 | 0.31 |
| OGYSGA1989 | *MATa, his3Δ1 leu2Δ0 ura3Δ0 LYS+, can1Δ::STE2pr-LEU2, lyp1Δ::, tor1-1, fpr1Δ::klURA, Tub4-(6)-RFP-(24)-FKBP::natNT2, TRS85-FRB::hphNT1, YPL057C-GFP::HIS3* | This study | 0.54 | 0.42 |
| OGYSGA1990 | *MATa, his3Δ1 leu2Δ0 ura3Δ0 LYS+, can1Δ::STE2pr-LEU2, lyp1Δ::, tor1-1, fpr1Δ::klURA, Tub4-(6)-RFP-(24)-FKBP::natNT2, TRS85-FRB::hphNT1, YLR350W-GFP::HIS3* | This study | 1.62 | 1.39 |
| OGYSGA1991 | *MATa, his3Δ1 leu2Δ0 ura3Δ0 LYS+, can1Δ::STE2pr-LEU2, lyp1Δ::, tor1-1, fpr1Δ::klURA, Tub4-(6)-RFP-(24)-FKBP::natNT2, TRS85-FRB::hphNT1, YLR268W-GFP::HIS3* | This study | 2.16 | 1.46 |
| OGYSGA1992 | *MATa, his3Δ1 leu2Δ0 ura3Δ0 LYS+, can1Δ::STE2pr-LEU2, lyp1Δ::, tor1-1, fpr1Δ::klURA, Tub4-(6)-RFP-(24)-FKBP::natNT2, TRS85-FRB::hphNT1, YJR118C-GFP::HIS3* | This study | 0 | 0 |
| OGYSGA1993 | *MATa, his3Δ1 leu2Δ0 ura3Δ0 LYS+, can1Δ::STE2pr-LEU2, lyp1Δ::, tor1-1, fpr1Δ::klURA, Tub4-(6)-RFP-(24)-FKBP::natNT2, TRS85-FRB::hphNT1, YLR039C-GFP::HIS3* | This study | 1.58 | 4.67 |
| OGYSGA1994 | *MATa, his3Δ1 leu2Δ0 ura3Δ0 LYS+, can1Δ::STE2pr-LEU2, lyp1Δ::, tor1-1, fpr1Δ::klURA, Tub4-(6)-RFP-(24)-FKBP::natNT2, TRS85-FRB::hphNT1, YBR080C-GFP::HIS3* | This study | 3.72 | 1.19 |
| OGYSGA1995 | *MATa, his3Δ1 leu2Δ0 ura3Δ0 LYS+, can1Δ::STE2pr-LEU2, lyp1Δ::, tor1-1, fpr1Δ::klURA, Tub4-(6)-RFP-(24)-FKBP::natNT2, TRS85-FRB::hphNT1, YJL123C-GFP::HIS3* | This study | 4.96 | 2.2 |
| OGYSGA1996 | *MATa, his3Δ1 leu2Δ0 ura3Δ0 LYS+, can1Δ::STE2pr-LEU2, lyp1Δ::, tor1-1, fpr1Δ::klURA, Tub4-(6)-RFP-(24)-FKBP::natNT2, TRS85-FRB::hphNT1, YPL234C-GFP::HIS3* | This study | 0 | 0 |
| OGYSGA1997 | *MATa, his3Δ1 leu2Δ0 ura3Δ0 LYS+, can1Δ::STE2pr-LEU2, lyp1Δ::, tor1-1, fpr1Δ::klURA, Tub4-(6)-RFP-(24)-FKBP::natNT2, TRS85-FRB::hphNT1, YKL119C-GFP::HIS3* | This study | 0 | 3.21 |
| OGYSGA1998 | *MATa, his3Δ1 leu2Δ0 ura3Δ0 LYS+, can1Δ::STE2pr-LEU2, lyp1Δ::, tor1-1, fpr1Δ::klURA, Tub4-(6)-RFP-(24)-FKBP::natNT2, TRS85-FRB::hphNT1, YLL040C-GFP::HIS3* | This study | 0 | 0 |
| OGYSGA1999 | *MATa, his3Δ1 leu2Δ0 ura3Δ0 LYS+, can1Δ::STE2pr-LEU2, lyp1Δ::, tor1-1, fpr1Δ::klURA, Tub4-(6)-RFP-(24)-FKBP::natNT2, TRS85-FRB::hphNT1, YPL120W-GFP::HIS3* | This study | 0 | 0 |
| OGYSGA2000 | *MATa, his3Δ1 leu2Δ0 ura3Δ0 LYS+, can1Δ::STE2pr-LEU2, lyp1Δ::, tor1-1, fpr1Δ::klURA, Tub4-(6)-RFP-(24)-FKBP::natNT2, TRS85-FRB::hphNT1, YAL026C-GFP::HIS3* | This study | 258.36 | 39.51 |
| OGYSGA2001 | *MATa, his3Δ1 leu2Δ0 ura3Δ0 LYS+, can1Δ::STE2pr-LEU2, lyp1Δ::, tor1-1, fpr1Δ::klURA, Tub4-(6)-RFP-(24)-FKBP::natNT2, TRS85-FRB::hphNT1, YKL004W-GFP::HIS3* | This study | 3.84 | 1.26 |
| OGYSGA2002 | *MATa, his3Δ1 leu2Δ0 ura3Δ0 LYS+, can1Δ::STE2pr-LEU2, lyp1Δ::, tor1-1, fpr1Δ::klURA, Tub4-(6)-RFP-(24)-FKBP::natNT2, TRS85-FRB::hphNT1, YMR190C-GFP::HIS3* | This study | 0 | 0 |
| OGYSGA2003 | *MATa, his3Δ1 leu2Δ0 ura3Δ0 LYS+, can1Δ::STE2pr-LEU2, lyp1Δ::, tor1-1, fpr1Δ::klURA, Tub4-(6)-RFP-(24)-FKBP::natNT2, TRS85-FRB::hphNT1, YDR137W-GFP::HIS3* | This study | 2.49 | 1.44 |
| OGYSGA2004 | *MATa, his3Δ1 leu2Δ0 ura3Δ0 LYS+, can1Δ::STE2pr-LEU2, lyp1Δ::, tor1-1, fpr1Δ::klURA, Tub4-(6)-RFP-(24)-FKBP::natNT2, TRS85-FRB::hphNT1, YIL004C-GFP::HIS3* | This study | 0 | 3.24 |
| OGYSGA2005 | *MATa, his3Δ1 leu2Δ0 ura3Δ0 LYS+, can1Δ::STE2pr-LEU2, lyp1Δ::, tor1-1, fpr1Δ::klURA, Tub4-(6)-RFP-(24)-FKBP::natNT2, TRS85-FRB::hphNT1, YPL158C-GFP::HIS3* | This study | 0 | 0 |
| OGYSGA2006 | *MATa, his3Δ1 leu2Δ0 ura3Δ0 LYS+, can1Δ::STE2pr-LEU2, lyp1Δ::, tor1-1, fpr1Δ::klURA, Tub4-(6)-RFP-(24)-FKBP::natNT2, TRS85-FRB::hphNT1, YNL169C-GFP::HIS3* | This study | 0.28 | 1.8 |
| OGYSGA2007 | *MATa, his3Δ1 leu2Δ0 ura3Δ0 LYS+, can1Δ::STE2pr-LEU2, lyp1Δ::, tor1-1, fpr1Δ::klURA, Tub4-(6)-RFP-(24)-FKBP::natNT2, TRS85-FRB::hphNT1, YML041C-GFP::HIS3* | This study | 4.61 | 1.14 |
| OGYSGA2008 | *MATa, his3Δ1 leu2Δ0 ura3Δ0 LYS+, can1Δ::STE2pr-LEU2, lyp1Δ::, tor1-1, fpr1Δ::klURA, Tub4-(6)-RFP-(24)-FKBP::natNT2, TRS85-FRB::hphNT1, YGL020C-GFP::HIS3* | This study | 0 | 0 |
| OGYSGA2009 | *MATa, his3Δ1 leu2Δ0 ura3Δ0 LYS+, can1Δ::STE2pr-LEU2, lyp1Δ::, tor1-1, fpr1Δ::klURA, Tub4-(6)-RFP-(24)-FKBP::natNT2, TRS85-FRB::hphNT1, YOL012C-GFP::HIS3* | This study | 0.24 | 0.98 |
| OGYSGA2010 | *MATa, his3Δ1 leu2Δ0 ura3Δ0 LYS+, can1Δ::STE2pr-LEU2, lyp1Δ::, tor1-1, fpr1Δ::klURA, Tub4-(6)-RFP-(24)-FKBP::natNT2, TRS85-FRB::hphNT1, YNR051C-GFP::HIS3* | This study | 0 | 0 |
| OGYSGA2011 | *MATa, his3Δ1 leu2Δ0 ura3Δ0 LYS+, can1Δ::STE2pr-LEU2, lyp1Δ::, tor1-1, fpr1Δ::klURA, Tub4-(6)-RFP-(24)-FKBP::natNT2, TRS85-FRB::hphNT1, YKL212W-GFP::HIS3* | This study | 0.35 | 0.18 |
| OGYSGA2012 | *MATa, his3Δ1 leu2Δ0 ura3Δ0 LYS+, can1Δ::STE2pr-LEU2, lyp1Δ::, tor1-1, fpr1Δ::klURA, Tub4-(6)-RFP-(24)-FKBP::natNT2, TRS85-FRB::hphNT1, YLR418C-GFP::HIS3* | This study | 0 | 0 |
| OGYSGA2013 | *MATa, his3Δ1 leu2Δ0 ura3Δ0 LYS+, can1Δ::STE2pr-LEU2, lyp1Δ::, tor1-1, fpr1Δ::klURA, Tub4-(6)-RFP-(24)-FKBP::natNT2, TRS85-FRB::hphNT1, YNL044W-GFP::HIS3* | This study | 11.06 | 0 |
| OGYSGA2014 | *MATa, his3Δ1 leu2Δ0 ura3Δ0 LYS+, can1Δ::STE2pr-LEU2, lyp1Δ::, tor1-1, fpr1Δ::klURA, Tub4-(6)-RFP-(24)-FKBP::natNT2, TRS85-FRB::hphNT1, YMR010W-GFP::HIS3* | This study | 0.51 | 1.83 |
| OGYSGA2015 | *MATa, his3Δ1 leu2Δ0 ura3Δ0 LYS+, can1Δ::STE2pr-LEU2, lyp1Δ::, tor1-1, fpr1Δ::klURA, Tub4-(6)-RFP-(24)-FKBP::natNT2, TRS85-FRB::hphNT1, YEL043W-GFP::HIS3* | This study | 0 | 0 |
| OGYSGA2016 | *MATa, his3Δ1 leu2Δ0 ura3Δ0 LYS+, can1Δ::STE2pr-LEU2, lyp1Δ::, tor1-1, fpr1Δ::klURA, Tub4-(6)-RFP-(24)-FKBP::natNT2, TRS85-FRB::hphNT1, YDL058W-GFP::HIS3* | This study | 4.53 | 0.92 |
| OGYSGA2017 | *MATa, his3Δ1 leu2Δ0 ura3Δ0 LYS+, can1Δ::STE2pr-LEU2, lyp1Δ::, tor1-1, fpr1Δ::klURA, Tub4-(6)-RFP-(24)-FKBP::natNT2, TRS85-FRB::hphNT1, YNL267W-GFP::HIS3* | This study | 0 | 0 |
| OGYSGA2018 | *MATa, his3Δ1 leu2Δ0 ura3Δ0 LYS+, can1Δ::STE2pr-LEU2, lyp1Δ::, tor1-1, fpr1Δ::klURA, Tub4-(6)-RFP-(24)-FKBP::natNT2, TRS85-FRB::hphNT1, YDL099W-GFP::HIS3* | This study | 0 | 0 |
| OGYSGA2019 | *MATa, his3Δ1 leu2Δ0 ura3Δ0 LYS+, can1Δ::STE2pr-LEU2, lyp1Δ::, tor1-1, fpr1Δ::klURA, Tub4-(6)-RFP-(24)-FKBP::natNT2, TRS85-FRB::hphNT1, YIL039W-GFP::HIS3* | This study | 0 | 0 |
| OGYSGA2020 | *MATa, his3Δ1 leu2Δ0 ura3Δ0 LYS+, can1Δ::STE2pr-LEU2, lyp1Δ::, tor1-1, fpr1Δ::klURA, Tub4-(6)-RFP-(24)-FKBP::natNT2, TRS85-FRB::hphNT1, YML012W-GFP::HIS3* | This study | 0.99 | 0 |
| OGYSGA2021 | *MATa, his3Δ1 leu2Δ0 ura3Δ0 LYS+, can1Δ::STE2pr-LEU2, lyp1Δ::, tor1-1, fpr1Δ::klURA, Tub4-(6)-RFP-(24)-FKBP::natNT2, TRS85-FRB::hphNT1, YDR372C-GFP::HIS3* | This study | 0 | 0 |
| OGYSGA2022 | *MATa, his3Δ1 leu2Δ0 ura3Δ0 LYS+, can1Δ::STE2pr-LEU2, lyp1Δ::, tor1-1, fpr1Δ::klURA, Tub4-(6)-RFP-(24)-FKBP::natNT2, TRS85-FRB::hphNT1, YGL200C-GFP::HIS3* | This study | 0.61 | 4.64 |
| OGYSGA2023 | *MATa, his3Δ1 leu2Δ0 ura3Δ0 LYS+, can1Δ::STE2pr-LEU2, lyp1Δ::, tor1-1, fpr1Δ::klURA, Tub4-(6)-RFP-(24)-FKBP::natNT2, TRS85-FRB::hphNT1, YJL117W-GFP::HIS3* | This study | 0 | 0 |
| OGYSGA2024 | *MATa, his3Δ1 leu2Δ0 ura3Δ0 LYS+, can1Δ::STE2pr-LEU2, lyp1Δ::, tor1-1, fpr1Δ::klURA, Tub4-(6)-RFP-(24)-FKBP::natNT2, TRS85-FRB::hphNT1, YEL042W-GFP::HIS3* | This study | 8.47 | 12.15 |
| OGYSGA2025 | *MATa, his3Δ1 leu2Δ0 ura3Δ0 LYS+, can1Δ::STE2pr-LEU2, lyp1Δ::, tor1-1, fpr1Δ::klURA, Tub4-(6)-RFP-(24)-FKBP::natNT2, TRS85-FRB::hphNT1, YKL069W-GFP::HIS3* | This study | 0 | 0 |

| | | | | |
|---|---|---|---|---|
| OGYSGA2026 | *MATa, his3Δ1 leu2Δ0 ura3Δ0 LYS+, can1Δ::STE2pr-LEU2, lyp1Δ::, tor1-1, fpr1Δ::klURA, Tub4-(6)-RFP-(24)-FKBP::natNT2, TRS85-FRB::hphNT1, YJR032W-GFP::HIS3* | This study | 0 | 0 |
| OGYSGA2027 | *MATa, his3Δ1 leu2Δ0 ura3Δ0 LYS+, can1Δ::STE2pr-LEU2, lyp1Δ::, tor1-1, fpr1Δ::klURA, Tub4-(6)-RFP-(24)-FKBP::natNT2, TRS85-FRB::hphNT1, YLR260W-GFP::HIS3* | This study | 0 | 0 |
| OGYSGA2028 | *MATa, his3Δ1 leu2Δ0 ura3Δ0 LYS+, can1Δ::STE2pr-LEU2, lyp1Δ::, tor1-1, fpr1Δ::klURA, Tub4-(6)-RFP-(24)-FKBP::natNT2, TRS85-FRB::hphNT1, YEL051W-GFP::HIS3* | This study | 0 | 2.51 |
| OGYSGA2029 | *MATa, his3Δ1 leu2Δ0 ura3Δ0 LYS+, can1Δ::STE2pr-LEU2, lyp1Δ::, tor1-1, fpr1Δ::klURA, Tub4-(6)-RFP-(24)-FKBP::natNT2, TRS85-FRB::hphNT1, YNL238W-GFP::HIS3* | This study | 5.12 | 7.41 |
| OGYSGA2030 | *MATa, his3Δ1 leu2Δ0 ura3Δ0 LYS+, can1Δ::STE2pr-LEU2, lyp1Δ::, tor1-1, fpr1Δ::klURA, Tub4-(6)-RFP-(24)-FKBP::natNT2, TRS85-FRB::hphNT1, YMR079W-GFP::HIS3* | This study | 0 | 0 |
| OGYSGA2031 | *MATa, his3Δ1 leu2Δ0 ura3Δ0 LYS+, can1Δ::STE2pr-LEU2, lyp1Δ::, tor1-1, fpr1Δ::klURA, Tub4-(6)-RFP-(24)-FKBP::natNT2, TRS85-FRB::hphNT1, YGL086W-GFP::HIS3* | This study | 0 | 0 |
| OGYSGA2032 | *MATa, his3Δ1 leu2Δ0 ura3Δ0 LYS+, can1Δ::STE2pr-LEU2, lyp1Δ::, tor1-1, fpr1Δ::klURA, Tub4-(6)-RFP-(24)-FKBP::natNT2, TRS85-FRB::hphNT1, YML008C-GFP::HIS3* | This study | 0.94 | 0 |
| OGYSGA2033 | *MATa, his3Δ1 leu2Δ0 ura3Δ0 LYS+, can1Δ::STE2pr-LEU2, lyp1Δ::, tor1-1, fpr1Δ::klURA, Tub4-(6)-RFP-(24)-FKBP::natNT2, TRS85-FRB::hphNT1, YJL004C-GFP::HIS3* | This study | 0 | 0 |
| OGYSGA2034 | *MATa, his3Δ1 leu2Δ0 ura3Δ0 LYS+, can1Δ::STE2pr-LEU2, lyp1Δ::, tor1-1, fpr1Δ::klURA, Tub4-(6)-RFP-(24)-FKBP::natNT2, TRS85-FRB::hphNT1, YDR320C-GFP::HIS3* | This study | 9 | 107.51 |
| OGYSGA2035 | *MATa, his3Δ1 leu2Δ0 ura3Δ0 LYS+, can1Δ::STE2pr-LEU2, lyp1Δ::, tor1-1, fpr1Δ::klURA, Tub4-(6)-RFP-(24)-FKBP::natNT2, TRS85-FRB::hphNT1, YMR216C-GFP::HIS3* | This study | 0 | 0 |
| OGYSGA2036 | *MATa, his3Δ1 leu2Δ0 ura3Δ0 LYS+, can1Δ::STE2pr-LEU2, lyp1Δ::, tor1-1, fpr1Δ::klURA, Tub4-(6)-RFP-(24)-FKBP::natNT2, TRS85-FRB::hphNT1, YGL084C-GFP::HIS3* | This study | 0 | 5.48 |
| OGYSGA2037 | *MATa, his3Δ1 leu2Δ0 ura3Δ0 LYS+, can1Δ::STE2pr-LEU2, lyp1Δ::, tor1-1, fpr1Δ::klURA, Tub4-(6)-RFP-(24)-FKBP::natNT2, TRS85-FRB::hphNT1, YOR254C-GFP::HIS3* | This study | 0 | 0 |
| OGYSGA2038 | *MATa, his3Δ1 leu2Δ0 ura3Δ0 LYS+, can1Δ::STE2pr-LEU2, lyp1Δ::, tor1-1, fpr1Δ::klURA, Tub4-(6)-RFP-(24)-FKBP::natNT2, TRS85-FRB::hphNT1, YGR092W-GFP::HIS3* | This study | 3.56 | 2.22 |
| OGYSGA2039 | *MATa, his3Δ1 leu2Δ0 ura3Δ0 LYS+, can1Δ::STE2pr-LEU2, lyp1Δ::, tor1-1, fpr1Δ::klURA, Tub4-(6)-RFP-(24)-FKBP::natNT2, TRS85-FRB::hphNT1, YJR060W-GFP::HIS3* | This study | 1.4 | 1.03 |
| OGYSGA2040 | *MATa, his3Δ1 leu2Δ0 ura3Δ0 LYS+, can1Δ::STE2pr-LEU2, lyp1Δ::, tor1-1, fpr1Δ::klURA, Tub4-(6)-RFP-(24)-FKBP::natNT2, TRS85-FRB::hphNT1, YLL006W-GFP::HIS3* | This study | 2.17 | 2.63 |
| OGYSGA2041 | *MATa, his3Δ1 leu2Δ0 ura3Δ0 LYS+, can1Δ::STE2pr-LEU2, lyp1Δ::, tor1-1, fpr1Δ::klURA, Tub4-(6)-RFP-(24)-FKBP::natNT2, TRS85-FRB::hphNT1, YCR044C-GFP::HIS3* | This study | 0.47 | 0 |
| OGYSGA2042 | *MATa, his3Δ1 leu2Δ0 ura3Δ0 LYS+, can1Δ::STE2pr-LEU2, lyp1Δ::, tor1-1, fpr1Δ::klURA, Tub4-(6)-RFP-(24)-FKBP::natNT2, TRS85-FRB::hphNT1, YNL297C-GFP::HIS3* | This study | 2.42 | 4.79 |
| OGYSGA2043 | *MATa, his3Δ1 leu2Δ0 ura3Δ0 LYS+, can1Δ::STE2pr-LEU2, lyp1Δ::, tor1-1, fpr1Δ::klURA, Tub4-(6)-RFP-(24)-FKBP::natNT2, TRS85-FRB::hphNT1, YCR034W-GFP::HIS3* | This study | 1.42 | 0 |
| OGYSGA2044 | *MATa, his3Δ1 leu2Δ0 ura3Δ0 LYS+, can1Δ::STE2pr-LEU2, lyp1Δ::, tor1-1, fpr1Δ::klURA, Tub4-(6)-RFP-(24)-FKBP::natNT2, VPS53-FRB::hphNT1, YCR017C-GFP::HIS3* | This study | 0 | 0 |
| OGYSGA2045 | *MATa, his3Δ1 leu2Δ0 ura3Δ0 LYS+, can1Δ::STE2pr-LEU2, lyp1Δ::, tor1-1, fpr1Δ::klURA, Tub4-(6)-RFP-(24)-FKBP::natNT2, VPS53-FRB::hphNT1, YNL243W-GFP::HIS3* | This study | 0 | 0 |
| OGYSGA2046 | *MATa, his3Δ1 leu2Δ0 ura3Δ0 LYS+, can1Δ::STE2pr-LEU2, lyp1Δ::, tor1-1, fpr1Δ::klURA, Tub4-(6)-RFP-(24)-FKBP::natNT2, VPS53-FRB::hphNT1, YOR164C-GFP::HIS3* | This study | 0 | 0 |
| OGYSGA2047 | *MATa, his3Δ1 leu2Δ0 ura3Δ0 LYS+, can1Δ::STE2pr-LEU2, lyp1Δ::, tor1-1, fpr1Δ::klURA, Tub4-(6)-RFP-(24)-FKBP::natNT2, VPS53-FRB::hphNT1, YCL001W-GFP::HIS3* | This study | 1.86 | 1.07 |
| OGYSGA2048 | *MATa, his3Δ1 leu2Δ0 ura3Δ0 LYS+, can1Δ::STE2pr-LEU2, lyp1Δ::, tor1-1, fpr1Δ::klURA, Tub4-(6)-RFP-(24)-FKBP::natNT2, VPS53-FRB::hphNT1, YBR245C-GFP::HIS3* | This study | 3.09 | 1.14 |
| OGYSGA2049 | *MATa, his3Δ1 leu2Δ0 ura3Δ0 LYS+, can1Δ::STE2pr-LEU2, lyp1Δ::, tor1-1, fpr1Δ::klURA, Tub4-(6)-RFP-(24)-FKBP::natNT2, VPS53-FRB::hphNT1, YMR079W-GFP::HIS3* | This study | 0 | 0 |
| OGYSGA2050 | *MATa, his3Δ1 leu2Δ0 ura3Δ0 LYS+, can1Δ::STE2pr-LEU2, lyp1Δ::, tor1-1, fpr1Δ::klURA, Tub4-(6)-RFP-(24)-FKBP::natNT2, VPS53-FRB::hphNT1, YMR216C-GFP::HIS3* | This study | 0 | 0 |
| OGYSGA2051 | *MATa, his3Δ1 leu2Δ0 ura3Δ0 LYS+, can1Δ::STE2pr-LEU2, lyp1Δ::, tor1-1, fpr1Δ::klURA, Tub4-(6)-RFP-(24)-FKBP::natNT2, VPS53-FRB::hphNT1, YKR001C-GFP::HIS3* | This study | 1.14 | 1.43 |
| OGYSGA2052 | *MATa, his3Δ1 leu2Δ0 ura3Δ0 LYS+, can1Δ::STE2pr-LEU2, lyp1Δ::, tor1-1, fpr1Δ::klURA, Tub4-(6)-RFP-(24)-FKBP::natNT2, VPS53-FRB::hphNT1, YLR372W-GFP::HIS3* | This study | 0.6 | 0 |
| OGYSGA2053 | *MATa, his3Δ1 leu2Δ0 ura3Δ0 LYS+, can1Δ::STE2pr-LEU2, lyp1Δ::, tor1-1, fpr1Δ::klURA, Tub4-(6)-RFP-(24)-FKBP::natNT2, VPS53-FRB::hphNT1, YLR350W-GFP::HIS3* | This study | 2.63 | 1.52 |
| OGYSGA2054 | *MATa, his3Δ1 leu2Δ0 ura3Δ0 LYS+, can1Δ::STE2pr-LEU2, lyp1Δ::, tor1-1, fpr1Δ::klURA, Tub4-(6)-RFP-(24)-FKBP::natNT2, VPS53-FRB::hphNT1, YGL020C-GFP::HIS3* | This study | 3.68 | 0.94 |
| OGYSGA2055 | *MATa, his3Δ1 leu2Δ0 ura3Δ0 LYS+, can1Δ::STE2pr-LEU2, lyp1Δ::, tor1-1, fpr1Δ::klURA, Tub4-(6)-RFP-(24)-FKBP::natNT2, VPS53-FRB::hphNT1, YER083C-GFP::HIS3* | This study | 9.22 | 1.13 |
| OGYSGA2056 | *MATa, his3Δ1 leu2Δ0 ura3Δ0 LYS+, can1Δ::STE2pr-LEU2, lyp1Δ::, tor1-1, fpr1Δ::klURA, Tub4-(6)-RFP-(24)-FKBP::natNT2, VPS53-FRB::hphNT1, YMR161W-GFP::HIS3* | This study | 5.29 | 0.39 |
| OGYSGA2057 | *MATa, his3Δ1 leu2Δ0 ura3Δ0 LYS+, can1Δ::STE2pr-LEU2, lyp1Δ::, tor1-1, fpr1Δ::klURA, Tub4-(6)-RFP-(24)-FKBP::natNT2, VPS53-FRB::hphNT1, YER122C-GFP::HIS3* | This study | 3.61 | 2.12 |
| OGYSGA2058 | *MATa, his3Δ1 leu2Δ0 ura3Δ0 LYS+, can1Δ::STE2pr-LEU2, lyp1Δ::, tor1-1, fpr1Δ::klURA, Tub4-(6)-RFP-(24)-FKBP::natNT2, VPS53-FRB::hphNT1, YJR117W-GFP::HIS3* | This study | 2.03 | 3.08 |
| OGYSGA2059 | *MATa, his3Δ1 leu2Δ0 ura3Δ0 LYS+, can1Δ::STE2pr-LEU2, lyp1Δ::, tor1-1, fpr1Δ::klURA, Tub4-(6)-RFP-(24)-FKBP::natNT2, VPS53-FRB::hphNT1, YKR029C-GFP::HIS3* | This study | 2.83 | 0.34 |
| OGYSGA2060 | *MATa, his3Δ1 leu2Δ0 ura3Δ0 LYS+, can1Δ::STE2pr-LEU2, lyp1Δ::, tor1-1, fpr1Δ::klURA, Tub4-(6)-RFP-(24)-FKBP::natNT2, VPS53-FRB::hphNT1, YGR063C-GFP::HIS3* | This study | 6.34 | 1.3 |
| OGYSGA2061 | *MATa, his3Δ1 leu2Δ0 ura3Δ0 LYS+, can1Δ::STE2pr-LEU2, lyp1Δ::, tor1-1, fpr1Δ::klURA, Tub4-(6)-RFP-(24)-FKBP::natNT2, VPS53-FRB::hphNT1, YML097C-GFP::HIS3* | This study | 0.91 | 0.33 |
| OGYSGA2062 | *MATa, his3Δ1 leu2Δ0 ura3Δ0 LYS+, can1Δ::STE2pr-LEU2, lyp1Δ::, tor1-1, fpr1Δ::klURA, Tub4-(6)-RFP-(24)-FKBP::natNT2, VPS53-FRB::hphNT1, YOL001W-GFP::HIS3* | This study | 0 | 0 |
| OGYSGA2063 | *MATa, his3Δ1 leu2Δ0 ura3Δ0 LYS+, can1Δ::STE2pr-LEU2, lyp1Δ::, tor1-1, fpr1Δ::klURA, Tub4-(6)-RFP-(24)-FKBP::natNT2, VPS53-FRB::hphNT1, YOR070C-GFP::HIS3* | This study | 8.88 | 10.61 |
| OGYSGA2064 | *MATa, his3Δ1 leu2Δ0 ura3Δ0 LYS+, can1Δ::STE2pr-LEU2, lyp1Δ::, tor1-1, fpr1Δ::klURA, Tub4-(6)-RFP-(24)-FKBP::natNT2, VPS53-FRB::hphNT1, YLR056W-GFP::HIS3* | This study | 2.84 | 0.41 |
| OGYSGA2065 | *MATa, his3Δ1 leu2Δ0 ura3Δ0 LYS+, can1Δ::STE2pr-LEU2, lyp1Δ::, tor1-1, fpr1Δ::klURA, Tub4-(6)-RFP-(24)-FKBP::natNT2, VPS53-FRB::hphNT1, YDL185W-GFP::HIS3* | This study | 2.73 | 0 |

| | | | | |
|---|---|---|---|---|
| OGYSGA2066 | *MATa, his3Δ1 leu2Δ0 ura3Δ0 LYS+, can1Δ::STE2pr-LEU2, lyp1Δ::, tor1-1, fpr1Δ::klURA, Tub4-(6)-RFP-(24)-FKBP::natNT2, VPS53-FRB::hphNT1, YGL058W-GFP::HIS3* | This study | 2.61 | 1.21 |
| OGYSGA2067 | *MATa, his3Δ1 leu2Δ0 ura3Δ0 LYS+, can1Δ::STE2pr-LEU2, lyp1Δ::, tor1-1, fpr1Δ::klURA, Tub4-(6)-RFP-(24)-FKBP::natNT2, VPS53-FRB::hphNT1, YJR073C-GFP::HIS3* | This study | 5.99 | 0 |
| OGYSGA2068 | *MATa, his3Δ1 leu2Δ0 ura3Δ0 LYS+, can1Δ::STE2pr-LEU2, lyp1Δ::, tor1-1, fpr1Δ::klURA, Tub4-(6)-RFP-(24)-FKBP::natNT2, VPS53-FRB::hphNT1, YDL074C-GFP::HIS3* | This study | 23.78 | 0.87 |
| OGYSGA2069 | *MATa, his3Δ1 leu2Δ0 ura3Δ0 LYS+, can1Δ::STE2pr-LEU2, lyp1Δ::, tor1-1, fpr1Δ::klURA, Tub4-(6)-RFP-(24)-FKBP::natNT2, VPS53-FRB::hphNT1, YPL055C-GFP::HIS3* | This study | 0 | 0 |
| OGYSGA2070 | *MATa, his3Δ1 leu2Δ0 ura3Δ0 LYS+, can1Δ::STE2pr-LEU2, lyp1Δ::, tor1-1, fpr1Δ::klURA, Tub4-(6)-RFP-(24)-FKBP::natNT2, VPS53-FRB::hphNT1, YDR056C-GFP::HIS3* | This study | 2.98 | 0.85 |
| OGYSGA2071 | *MATa, his3Δ1 leu2Δ0 ura3Δ0 LYS+, can1Δ::STE2pr-LEU2, lyp1Δ::, tor1-1, fpr1Δ::klURA, Tub4-(6)-RFP-(24)-FKBP::natNT2, VPS53-FRB::hphNT1, YOR244W-GFP::HIS3* | This study | 0.87 | 0.38 |
| OGYSGA2072 | *MATa, his3Δ1 leu2Δ0 ura3Δ0 LYS+, can1Δ::STE2pr-LEU2, lyp1Δ::, tor1-1, fpr1Δ::klURA, Tub4-(6)-RFP-(24)-FKBP::natNT2, VPS53-FRB::hphNT1, YGR105W-GFP::HIS3* | This study | 0 | 1.44 |
| OGYSGA2073 | *MATa, his3Δ1 leu2Δ0 ura3Δ0 LYS+, can1Δ::STE2pr-LEU2, lyp1Δ::, tor1-1, fpr1Δ::klURA, Tub4-(6)-RFP-(24)-FKBP::natNT2, VPS53-FRB::hphNT1, YAR002C-A -GFP::HIS3* | This study | 16.25 | 1.24 |
| OGYSGA2074 | *MATa, his3Δ1 leu2Δ0 ura3Δ0 LYS+, can1Δ::STE2pr-LEU2, lyp1Δ::, tor1-1, fpr1Δ::klURA, Tub4-(6)-RFP-(24)-FKBP::natNT2, VPS53-FRB::hphNT1, YBR041W-GFP::HIS3* | This study | 2.17 | 1.28 |
| OGYSGA2075 | *MATa, his3Δ1 leu2Δ0 ura3Δ0 LYS+, can1Δ::STE2pr-LEU2, lyp1Δ::, tor1-1, fpr1Δ::klURA, Tub4-(6)-RFP-(24)-FKBP::natNT2, VPS53-FRB::hphNT1, YDL099W-GFP::HIS3* | This study | 1.86 | 1 |
| OGYSGA2076 | *MATa, his3Δ1 leu2Δ0 ura3Δ0 LYS+, can1Δ::STE2pr-LEU2, lyp1Δ::, tor1-1, fpr1Δ::klURA, Tub4-(6)-RFP-(24)-FKBP::natNT2, VPS53-FRB::hphNT1, YML075C-GFP::HIS3* | This study | 1.7 | 0.43 |
| OGYSGA2077 | *MATa, his3Δ1 leu2Δ0 ura3Δ0 LYS+, can1Δ::STE2pr-LEU2, lyp1Δ::, tor1-1, fpr1Δ::klURA, Tub4-(6)-RFP-(24)-FKBP::natNT2, VPS53-FRB::hphNT1, YNL323W-GFP::HIS3* | This study | 28.38 | 25.42 |
| OGYSGA2078 | *MATa, his3Δ1 leu2Δ0 ura3Δ0 LYS+, can1Δ::STE2pr-LEU2, lyp1Δ::, tor1-1, fpr1Δ::klURA, Tub4-(6)-RFP-(24)-FKBP::natNT2, VPS53-FRB::hphNT1, YAL058W-GFP::HIS3* | This study | 4.77 | 0 |
| OGYSGA2079 | *MATa, his3Δ1 leu2Δ0 ura3Δ0 LYS+, can1Δ::STE2pr-LEU2, lyp1Δ::, tor1-1, fpr1Δ::klURA, Tub4-(6)-RFP-(24)-FKBP::natNT2, VPS53-FRB::hphNT1, YOL111C-GFP::HIS3* | This study | 0 | 2.71 |
| OGYSGA2080 | *MATa, his3Δ1 leu2Δ0 ura3Δ0 LYS+, can1Δ::STE2pr-LEU2, lyp1Δ::, tor1-1, fpr1Δ::klURA, Tub4-(6)-RFP-(24)-FKBP::natNT2, VPS53-FRB::hphNT1, YIL030C-GFP::HIS3* | This study | 3.36 | 1 |
| OGYSGA2081 | *MATa, his3Δ1 leu2Δ0 ura3Δ0 LYS+, can1Δ::STE2pr-LEU2, lyp1Δ::, tor1-1, fpr1Δ::klURA, Tub4-(6)-RFP-(24)-FKBP::natNT2, VPS53-FRB::hphNT1, YOL013C-GFP::HIS3* | This study | 14.06 | 1.61 |
| OGYSGA2082 | *MATa, his3Δ1 leu2Δ0 ura3Δ0 LYS+, can1Δ::STE2pr-LEU2, lyp1Δ::, tor1-1, fpr1Δ::klURA, Tub4-(6)-RFP-(24)-FKBP::natNT2, VPS53-FRB::hphNT1, YMR015C-GFP::HIS3* | This study | 2.67 | 0 |
| OGYSGA2083 | *MATa, his3Δ1 leu2Δ0 ura3Δ0 LYS+, can1Δ::STE2pr-LEU2, lyp1Δ::, tor1-1, fpr1Δ::klURA, Tub4-(6)-RFP-(24)-FKBP::natNT2, VPS53-FRB::hphNT1, YBR283C-GFP::HIS3* | This study | 2.84 | 0 |
| OGYSGA2084 | *MATa, his3Δ1 leu2Δ0 ura3Δ0 LYS+, can1Δ::STE2pr-LEU2, lyp1Δ::, tor1-1, fpr1Δ::klURA, Tub4-(6)-RFP-(24)-FKBP::natNT2, VPS53-FRB::hphNT1, YOR085W-GFP::HIS3* | This study | 21.58 | 0.48 |
| OGYSGA2085 | *MATa, his3Δ1 leu2Δ0 ura3Δ0 LYS+, can1Δ::STE2pr-LEU2, lyp1Δ::, tor1-1, fpr1Δ::klURA, Tub4-(6)-RFP-(24)-FKBP::natNT2, VPS53-FRB::hphNT1, YNL267W-GFP::HIS3* | This study | 69.47 | 0 |
| OGYSGA2086 | *MATa, his3Δ1 leu2Δ0 ura3Δ0 LYS+, can1Δ::STE2pr-LEU2, lyp1Δ::, tor1-1, fpr1Δ::klURA, Tub4-(6)-RFP-(24)-FKBP::natNT2, VPS53-FRB::hphNT1, YNR052C-GFP::HIS3* | This study | 0 | 0 |
| OGYSGA2087 | *MATa, his3Δ1 leu2Δ0 ura3Δ0 LYS+, can1Δ::STE2pr-LEU2, lyp1Δ::, tor1-1, fpr1Δ::klURA, Tub4-(6)-RFP-(24)-FKBP::natNT2, VPS53-FRB::hphNT1, YAL023C-GFP::HIS3* | This study | 0 | 0 |
| OGYSGA2088 | *MATa, his3Δ1 leu2Δ0 ura3Δ0 LYS+, can1Δ::STE2pr-LEU2, lyp1Δ::, tor1-1, fpr1Δ::klURA, Tub4-(6)-RFP-(24)-FKBP::natNT2, VPS53-FRB::hphNT1, YGR060W-GFP::HIS3* | This study | 1.58 | 0 |
| OGYSGA2089 | *MATa, his3Δ1 leu2Δ0 ura3Δ0 LYS+, can1Δ::STE2pr-LEU2, lyp1Δ::, tor1-1, fpr1Δ::klURA, Tub4-(6)-RFP-(24)-FKBP::natNT2, VPS53-FRB::hphNT1, YLL040C-GFP::HIS3* | This study | 0 | 0 |
| OGYSGA2090 | *MATa, his3Δ1 leu2Δ0 ura3Δ0 LYS+, can1Δ::STE2pr-LEU2, lyp1Δ::, tor1-1, fpr1Δ::klURA, Tub4-(6)-RFP-(24)-FKBP::natNT2, VPS53-FRB::hphNT1, YOR216C-GFP::HIS3* | This study | 1.92 | 0.57 |
| OGYSGA2091 | *MATa, his3Δ1 leu2Δ0 ura3Δ0 LYS+, can1Δ::STE2pr-LEU2, lyp1Δ::, tor1-1, fpr1Δ::klURA, Tub4-(6)-RFP-(24)-FKBP::natNT2, VPS53-FRB::hphNT1, YFL024C-GFP::HIS3* | This study | 7.63 | 2.99 |
| OGYSGA2092 | *MATa, his3Δ1 leu2Δ0 ura3Δ0 LYS+, can1Δ::STE2pr-LEU2, lyp1Δ::, tor1-1, fpr1Δ::klURA, Tub4-(6)-RFP-(24)-FKBP::natNT2, VPS53-FRB::hphNT1, YAL021C-GFP::HIS3* | This study | 0 | 0 |
| OGYSGA2093 | *MATa, his3Δ1 leu2Δ0 ura3Δ0 LYS+, can1Δ::STE2pr-LEU2, lyp1Δ::, tor1-1, fpr1Δ::klURA, Tub4-(6)-RFP-(24)-FKBP::natNT2, VPS53-FRB::hphNT1, YPL120W-GFP::HIS3* | This study | 0 | 0 |
| OGYSGA2094 | *MATa, his3Δ1 leu2Δ0 ura3Δ0 LYS+, can1Δ::STE2pr-LEU2, lyp1Δ::, tor1-1, fpr1Δ::klURA, Tub4-(6)-RFP-(24)-FKBP::natNT2, VPS53-FRB::hphNT1, YPR032W-GFP::HIS3* | This study | 0 | 0 |
| OGYSGA2095 | *MATa, his3Δ1 leu2Δ0 ura3Δ0 LYS+, can1Δ::STE2pr-LEU2, lyp1Δ::, tor1-1, fpr1Δ::klURA, Tub4-(6)-RFP-(24)-FKBP::natNT2, VPS53-FRB::hphNT1, YGL054C-GFP::HIS3* | This study | 4.48 | 1.4 |
| OGYSGA2096 | *MATa, his3Δ1 leu2Δ0 ura3Δ0 LYS+, can1Δ::STE2pr-LEU2, lyp1Δ::, tor1-1, fpr1Δ::klURA, Tub4-(6)-RFP-(24)-FKBP::natNT2, VPS53-FRB::hphNT1, YDL100C-GFP::HIS3* | This study | 8.09 | 0.41 |
| OGYSGA2097 | *MATa, his3Δ1 leu2Δ0 ura3Δ0 LYS+, can1Δ::STE2pr-LEU2, lyp1Δ::, tor1-1, fpr1Δ::klURA, Tub4-(6)-RFP-(24)-FKBP::natNT2, VPS53-FRB::hphNT1, YJL004C-GFP::HIS3* | This study | 4.7 | 4.02 |
| OGYSGA2098 | *MATa, his3Δ1 leu2Δ0 ura3Δ0 LYS+, can1Δ::STE2pr-LEU2, lyp1Δ::, tor1-1, fpr1Δ::klURA, Tub4-(6)-RFP-(24)-FKBP::natNT2, VPS53-FRB::hphNT1, YCR044C-GFP::HIS3* | This study | 10.97 | 0.63 |
| OGYSGA2099 | *MATa, his3Δ1 leu2Δ0 ura3Δ0 LYS+, can1Δ::STE2pr-LEU2, lyp1Δ::, tor1-1, fpr1Δ::klURA, Tub4-(6)-RFP-(24)-FKBP::natNT2, VPS53-FRB::hphNT1, YGL084C-GFP::HIS3* | This study | 6.51 | 1.93 |
| OGYSGA2100 | *MATa, his3Δ1 leu2Δ0 ura3Δ0 LYS+, can1Δ::STE2pr-LEU2, lyp1Δ::, tor1-1, fpr1Δ::klURA, Tub4-(6)-RFP-(24)-FKBP::natNT2, VPS53-FRB::hphNT1, YMR123W-GFP::HIS3* | This study | 1.09 | 0 |
| OGYSGA2101 | *MATa, his3Δ1 leu2Δ0 ura3Δ0 LYS+, can1Δ::STE2pr-LEU2, lyp1Δ::, tor1-1, fpr1Δ::klURA, Tub4-(6)-RFP-(24)-FKBP::natNT2, VPS53-FRB::hphNT1, YNL271C-GFP::HIS3* | This study | 0 | 0 |
| OGYSGA2102 | *MATa, his3Δ1 leu2Δ0 ura3Δ0 LYS+, can1Δ::STE2pr-LEU2, lyp1Δ::, tor1-1, fpr1Δ::klURA, Tub4-(6)-RFP-(24)-FKBP::natNT2, VPS53-FRB::hphNT1, YJR118C-GFP::HIS3* | This study | 1.33 | 2.24 |
| OGYSGA4423 | *MATa, his3Δ1 leu2Δ0 ura3Δ0 LYS+, can1Δ::STE2pr-LEU2, lyp1Δ::, tor1-1, fpr1Δ::klURA, Tub4-(6)-RFP-(24)-FKBP::natNT2, SEC3-FRB::hphNT1, YMR183C-GFP::HIS3* | This study | 0 | 81.77 |
| OGYSGA4424 | *MATa, his3Δ1 leu2Δ0 ura3Δ0 LYS+, can1Δ::STE2pr-LEU2, lyp1Δ::, tor1-1, fpr1Δ::klURA, Tub4-(6)-RFP-(24)-FKBP::natNT2, SEC3-FRB::hphNT1, YDR086C-GFP::HIS3* | This study | 0 | 0 |
| OGYSGA4425 | *MATa, his3Δ1 leu2Δ0 ura3Δ0 LYS+, can1Δ::STE2pr-LEU2, lyp1Δ::, tor1-1, fpr1Δ::klURA, Tub4-(6)-RFP-(24)-FKBP::natNT2, SEC3-FRB::hphNT1, YOR327C-GFP::HIS3* | This study | 2.44 | 0 |

| | | | | |
|---|---|---|---|---|
| OGYSGA4426 | *MATa, his3Δ1 leu2Δ0 ura3Δ0 LYS+, can1Δ::STE2pr-LEU2, lyp1Δ::, tor1-1, fpr1Δ::klURA, Tub4-(6)-RFP-(24)-FKBP::natNT2, SEC3-FRB::hphNT1, YNL267W-GFP::HIS3* | This study | 0.33 | 0 |
| OGYSGA4427 | *MATa, his3Δ1 leu2Δ0 ura3Δ0 LYS+, can1Δ::STE2pr-LEU2, lyp1Δ::, tor1-1, fpr1Δ::klURA, Tub4-(6)-RFP-(24)-FKBP::natNT2, SEC3-FRB::hphNT1, YJL145W-GFP::HIS3* | This study | 0 | 0 |
| OGYSGA4428 | *MATa, his3Δ1 leu2Δ0 ura3Δ0 LYS+, can1Δ::STE2pr-LEU2, lyp1Δ::, tor1-1, fpr1Δ::klURA, Tub4-(6)-RFP-(24)-FKBP::natNT2, SEC3-FRB::hphNT1, YPL232W-GFP::HIS3* | This study | 20.25 | 0 |
| OGYSGA4429 | *MATa, his3Δ1 leu2Δ0 ura3Δ0 LYS+, can1Δ::STE2pr-LEU2, lyp1Δ::, tor1-1, fpr1Δ::klURA, Tub4-(6)-RFP-(24)-FKBP::natNT2, SEC3-FRB::hphNT1, YPR017C-GFP::HIS3* | This study | 0.13 | 0 |
| OGYSGA4430 | *MATa, his3Δ1 leu2Δ0 ura3Δ0 LYS+, can1Δ::STE2pr-LEU2, lyp1Δ::, tor1-1, fpr1Δ::klURA, Tub4-(6)-RFP-(24)-FKBP::natNT2, SEC3-FRB::hphNT1, YDR363W-A-GFP::HIS3* | This study | 0 | 0 |
| OGYSGA4431 | *MATa, his3Δ1 leu2Δ0 ura3Δ0 LYS+, can1Δ::STE2pr-LEU2, lyp1Δ::, tor1-1, fpr1Δ::klURA, Tub4-(6)-RFP-(24)-FKBP::natNT2, SEC3-FRB::hphNT1, YPL069C-GFP::HIS3* | This study | 1.2 | 0.91 |
| OGYSGA4432 | *MATa, his3Δ1 leu2Δ0 ura3Δ0 LYS+, can1Δ::STE2pr-LEU2, lyp1Δ::, tor1-1, fpr1Δ::klURA, Tub4-(6)-RFP-(24)-FKBP::natNT2, SEC3-FRB::hphNT1, YOR122C-GFP::HIS3* | This study | 0 | 0 |
| OGYSGA4433 | *MATa, his3Δ1 leu2Δ0 ura3Δ0 LYS+, can1Δ::STE2pr-LEU2, lyp1Δ::, tor1-1, fpr1Δ::klURA, Tub4-(6)-RFP-(24)-FKBP::natNT2, SEC3-FRB::hphNT1, YLR096W-GFP::HIS3* | This study | 1.52 | 0 |
| OGYSGA4434 | *MATa, his3Δ1 leu2Δ0 ura3Δ0 LYS+, can1Δ::STE2pr-LEU2, lyp1Δ::, tor1-1, fpr1Δ::klURA, Tub4-(6)-RFP-(24)-FKBP::natNT2, SEC3-FRB::hphNT1, YLR378C-GFP::HIS3* | This study | 0 | 0 |
| OGYSGA4435 | *MATa, his3Δ1 leu2Δ0 ura3Δ0 LYS+, can1Δ::STE2pr-LEU2, lyp1Δ::, tor1-1, fpr1Δ::klURA, Tub4-(6)-RFP-(24)-FKBP::natNT2, SEC3-FRB::hphNT1, YDR122W-GFP::HIS3* | This study | 0 | 0 |
| OGYSGA4436 | *MATa, his3Δ1 leu2Δ0 ura3Δ0 LYS+, can1Δ::STE2pr-LEU2, lyp1Δ::, tor1-1, fpr1Δ::klURA, Tub4-(6)-RFP-(24)-FKBP::natNT2, SEC3-FRB::hphNT1, YER155C-GFP::HIS3* | This study | 0 | 0 |
| OGYSGA4437 | *MATa, his3Δ1 leu2Δ0 ura3Δ0 LYS+, can1Δ::STE2pr-LEU2, lyp1Δ::, tor1-1, fpr1Δ::klURA, Tub4-(6)-RFP-(24)-FKBP::natNT2, SEC3-FRB::hphNT1, YNL271C-GFP::HIS3* | This study | 0 | 0 |
| OGYSGA4438 | *MATa, his3Δ1 leu2Δ0 ura3Δ0 LYS+, can1Δ::STE2pr-LEU2, lyp1Δ::, tor1-1, fpr1Δ::klURA, Tub4-(6)-RFP-(24)-FKBP::natNT2, SEC3-FRB::hphNT1, YGL167C-GFP::HIS3* | This study | 0 | 0.53 |
| OGYSGA4439 | *MATa, his3Δ1 leu2Δ0 ura3Δ0 LYS+, can1Δ::STE2pr-LEU2, lyp1Δ::, tor1-1, fpr1Δ::klURA, Tub4-(6)-RFP-(24)-FKBP::natNT2, SEC3-FRB::hphNT1, YDR293C-GFP::HIS3* | This study | 0 | 0 |
| OGYSGA4440 | *MATa, his3Δ1 leu2Δ0 ura3Δ0 LYS+, can1Δ::STE2pr-LEU2, lyp1Δ::, tor1-1, fpr1Δ::klURA, Tub4-(6)-RFP-(24)-FKBP::natNT2, SEC3-FRB::hphNT1, YKL212W-GFP::HIS3* | This study | 0.65 | 0 |
| OGYSGA4441 | *MATa, his3Δ1 leu2Δ0 ura3Δ0 LYS+, can1Δ::STE2pr-LEU2, lyp1Δ::, tor1-1, fpr1Δ::klURA, Tub4-(6)-RFP-(24)-FKBP::natNT2, SEC3-FRB::hphNT1, YHR030C-GFP::HIS3* | This study | 0 | 0 |
| OGYSGA4442 | *MATa, his3Δ1 leu2Δ0 ura3Δ0 LYS+, can1Δ::STE2pr-LEU2, lyp1Δ::, tor1-1, fpr1Δ::klURA, Tub4-(6)-RFP-(24)-FKBP::natNT2, SEC3-FRB::hphNT1, YMR307W-GFP::HIS3* | This study | 0 | 0 |
| OGYSGA4443 | *MATa, his3Δ1 leu2Δ0 ura3Δ0 LYS+, can1Δ::STE2pr-LEU2, lyp1Δ::, tor1-1, fpr1Δ::klURA, Tub4-(6)-RFP-(24)-FKBP::natNT2, SEC3-FRB::hphNT1, YDL019C-GFP::HIS3* | This study | 0.46 | 0 |
| OGYSGA4444 | *MATa, his3Δ1 leu2Δ0 ura3Δ0 LYS+, can1Δ::STE2pr-LEU2, lyp1Δ::, tor1-1, fpr1Δ::klURA, Tub4-(6)-RFP-(24)-FKBP::natNT2, SEC3-FRB::hphNT1, YER098W-GFP::HIS3* | This study | 0 | 0 |
| OGYSGA4445 | *MATa, his3Δ1 leu2Δ0 ura3Δ0 LYS+, can1Δ::STE2pr-LEU2, lyp1Δ::, tor1-1, fpr1Δ::klURA, Tub4-(6)-RFP-(24)-FKBP::natNT2, SEC3-FRB::hphNT1, YER019C-A-GFP::HIS3* | This study | 0 | 0 |
| OGYSGA4446 | *MATa, his3Δ1 leu2Δ0 ura3Δ0 LYS+, can1Δ::STE2pr-LEU2, lyp1Δ::, tor1-1, fpr1Δ::klURA, Tub4-(6)-RFP-(24)-FKBP::natNT2, SEC3-FRB::hphNT1, YDR116C-GFP::HIS3* | This study | 0 | 0.67 |
| OGYSGA4447 | *MATa, his3Δ1 leu2Δ0 ura3Δ0 LYS+, can1Δ::STE2pr-LEU2, lyp1Δ::, tor1-1, fpr1Δ::klURA, Tub4-(6)-RFP-(24)-FKBP::natNT2, SEC3-FRB::hphNT1, YER114C-GFP::HIS3* | This study | 2.19 | 0 |
| OGYSGA4448 | *MATa, his3Δ1 leu2Δ0 ura3Δ0 LYS+, can1Δ::STE2pr-LEU2, lyp1Δ::, tor1-1, fpr1Δ::klURA, Tub4-(6)-RFP-(24)-FKBP::natNT2, SEC3-FRB::hphNT1, YIL046W-GFP::HIS3* | This study | 0.6 | 0 |
| OGYSGA4449 | *MATa, his3Δ1 leu2Δ0 ura3Δ0 LYS+, can1Δ::STE2pr-LEU2, lyp1Δ::, tor1-1, fpr1Δ::klURA, Tub4-(6)-RFP-(24)-FKBP::natNT2, SEC3-FRB::hphNT1, YNL293W-GFP::HIS3* | This study | 0 | 0 |
| OGYSGA4450 | *MATa, his3Δ1 leu2Δ0 ura3Δ0 LYS+, can1Δ::STE2pr-LEU2, lyp1Δ::, tor1-1, fpr1Δ::klURA, Tub4-(6)-RFP-(24)-FKBP::natNT2, SEC3-FRB::hphNT1, YBR135W-GFP::HIS3* | This study | 0 | 0 |
| OGYSGA4451 | *MATa, his3Δ1 leu2Δ0 ura3Δ0 LYS+, can1Δ::STE2pr-LEU2, lyp1Δ::, tor1-1, fpr1Δ::klURA, Tub4-(6)-RFP-(24)-FKBP::natNT2, SEC3-FRB::hphNT1, YDR099W-GFP::HIS3* | This study | 0 | 0 |
| OGYSGA4452 | *MATa, his3Δ1 leu2Δ0 ura3Δ0 LYS+, can1Δ::STE2pr-LEU2, lyp1Δ::, tor1-1, fpr1Δ::klURA, Tub4-(6)-RFP-(24)-FKBP::natNT2, SEC3-FRB::hphNT1, YMR149W-GFP::HIS3* | This study | 0.17 | 0.3 |
| OGYSGA4453 | *MATa, his3Δ1 leu2Δ0 ura3Δ0 LYS+, can1Δ::STE2pr-LEU2, lyp1Δ::, tor1-1, fpr1Δ::klURA, Tub4-(6)-RFP-(24)-FKBP::natNT2, SEC3-FRB::hphNT1, YBR088C-GFP::HIS3* | This study | 0 | 0 |
| OGYSGA4454 | *MATa, his3Δ1 leu2Δ0 ura3Δ0 LYS+, can1Δ::STE2pr-LEU2, lyp1Δ::, tor1-1, fpr1Δ::klURA, Tub4-(6)-RFP-(24)-FKBP::natNT2, SEC3-FRB::hphNT1, YMR079W-GFP::HIS3* | This study | 0.28 | 0 |
| OGYSGA4455 | *MATa, his3Δ1 leu2Δ0 ura3Δ0 LYS+, can1Δ::STE2pr-LEU2, lyp1Δ::, tor1-1, fpr1Δ::klURA, Tub4-(6)-RFP-(24)-FKBP::natNT2, SEC3-FRB::hphNT1, YBR126C-GFP::HIS3* | This study | 0 | 0 |
| OGYSGA4456 | *MATa, his3Δ1 leu2Δ0 ura3Δ0 LYS+, can1Δ::STE2pr-LEU2, lyp1Δ::, tor1-1, fpr1Δ::klURA, Tub4-(6)-RFP-(24)-FKBP::natNT2, SEC3-FRB::hphNT1, YPR024W-GFP::HIS3* | This study | 0 | 1.34 |
| OGYSGA4457 | *MATa, his3Δ1 leu2Δ0 ura3Δ0 LYS+, can1Δ::STE2pr-LEU2, lyp1Δ::, tor1-1, fpr1Δ::klURA, Tub4-(6)-RFP-(24)-FKBP::natNT2, SEC3-FRB::hphNT1, YER177W-GFP::HIS3* | This study | 0 | 0 |
| OGYSGA4458 | *MATa, his3Δ1 leu2Δ0 ura3Δ0 LYS+, can1Δ::STE2pr-LEU2, lyp1Δ::, tor1-1, fpr1Δ::klURA, Tub4-(6)-RFP-(24)-FKBP::natNT2, SEC3-FRB::hphNT1, YDR217C-GFP::HIS3* | This study | 0.94 | 0 |
| OGYSGA4459 | *MATa, his3Δ1 leu2Δ0 ura3Δ0 LYS+, can1Δ::STE2pr-LEU2, lyp1Δ::, tor1-1, fpr1Δ::klURA, Tub4-(6)-RFP-(24)-FKBP::natNT2, SEC3-FRB::hphNT1, YDR477W-GFP::HIS3* | This study | 0 | 0 |
| OGYSGA4460 | *MATa, his3Δ1 leu2Δ0 ura3Δ0 LYS+, can1Δ::STE2pr-LEU2, lyp1Δ::, tor1-1, fpr1Δ::klURA, Tub4-(6)-RFP-(24)-FKBP::natNT2, SEC3-FRB::hphNT1, YBR095C-GFP::HIS3* | This study | 0 | 1.67 |
| OGYSGA4461 | *MATa, his3Δ1 leu2Δ0 ura3Δ0 LYS+, can1Δ::STE2pr-LEU2, lyp1Δ::, tor1-1, fpr1Δ::klURA, Tub4-(6)-RFP-(24)-FKBP::natNT2, SEC3-FRB::hphNT1, YGL173C-GFP::HIS3* | This study | 4.21 | 0 |
| OGYSGA4462 | *MATa, his3Δ1 leu2Δ0 ura3Δ0 LYS+, can1Δ::STE2pr-LEU2, lyp1Δ::, tor1-1, fpr1Δ::klURA, Tub4-(6)-RFP-(24)-FKBP::natNT2, SEC3-FRB::hphNT1, YKL033W-GFP::HIS3* | This study | 1.44 | 0 |
| OGYSGA4463 | *MATa, his3Δ1 leu2Δ0 ura3Δ0 LYS+, can1Δ::STE2pr-LEU2, lyp1Δ::, tor1-1, fpr1Δ::klURA, Tub4-(6)-RFP-(24)-FKBP::natNT2, SEC3-FRB::hphNT1, YIL084C-GFP::HIS3* | This study | 0 | 0 |

Values of 0 indicate that no prey was detected in the anchoring platform.
