## Supplementary material for "Drs2 regulates TRAPPIII in Atg9 transport: exposing the interplay of P4-ATPases and Multisubunit Tethering Complexes": Supplementary Table 2.pdf

| Yeast strains and results of the PICT assay to determine the most efficient bait to anchor each MTC. | | | |
|---|---|---|---|
| **Strain** | **Genotype** | **Source** | **Interaction score** |
| OGYSGA6043 | MATa, his3Δ1, leu2Δ0, ura3Δ0, LYS+, can1Δ::STE2pr-LEU2, lyp1Δ::, tor1-1, fpr1Δ::klURA, Tub4-(6)-RFP-(24)-FKBP::natNT2 DSL1-FRB::hphNT1, TIP20-3xmyeGFP::kanMX4 | This study | 615.59 |
| OGYSGA6044 | MATa, his3Δ1, leu2Δ0, ura3Δ0, LYS+, can1Δ::STE2pr-LEU2, lyp1Δ::, tor1-1, fpr1Δ::klURA, Tub4-(6)-RFP-(24)-FKBP::natNT2 DSL1-FRB::hphNT1, SEC39-3xmyeGFP::kanMX4 | This study | -13.35 |
| OGYSGA6045 | MATa, his3Δ1, leu2Δ0, ura3Δ0, LYS+, can1Δ::STE2pr-LEU2, lyp1Δ::, tor1-1, fpr1Δ::klURA, Tub4-(6)-RFP-(24)-FKBP::natNT2 TIP20-FRB::hphNT1, DSL1-3xmyeGFP::kanMX4 | This study | -94.33 |
| OGYSGA6046 | MATa, his3Δ1, leu2Δ0, ura3Δ0, LYS+, can1Δ::STE2pr-LEU2, lyp1Δ::, tor1-1, fpr1Δ::klURA, Tub4-(6)-RFP-(24)-FKBP::natNT2 TIP20-FRB::hphNT1, SEC39-3xmyeGFP::kanMX4 | This study | -45.46 |
| OGYSGA6047 | MATa, his3Δ1, leu2Δ0, ura3Δ0, LYS+, can1Δ::STE2pr-LEU2, lyp1Δ::, tor1-1, fpr1Δ::klURA, Tub4-(6)-RFP-(24)-FKBP::natNT2 SEC39-FRB::hphNT1, DSL1-3xmyeGFP::kanMX4 | This study | -25.03 |
| OGYSGA6048 | MATa, his3Δ1, leu2Δ0, ura3Δ0, LYS+, can1Δ::STE2pr-LEU2, lyp1Δ::, tor1-1, fpr1Δ::klURA, Tub4-(6)-RFP-(24)-FKBP::natNT2 SEC39-FRB::hphNT1, TIP20-3xmyeGFP::kanMX4 | This study | -0.75 |
| OGYSGA6049 | MATa, his3Δ1, leu2Δ0, ura3Δ0, LYS+, can1Δ::STE2pr-LEU2, lyp1Δ::, tor1-1, fpr1Δ::klURA, Tub4-(6)-RFP-(24)-FKBP::natNT2 COG1-FRB::hphNT1, COG2-3xmyeGFP::kanMX4 | This study | -16.61 |
| OGYSGA6050 | MATa, his3Δ1, leu2Δ0, ura3Δ0, LYS+, can1Δ::STE2pr-LEU2, lyp1Δ::, tor1-1, fpr1Δ::klURA, Tub4-(6)-RFP-(24)-FKBP::natNT2 COG1-FRB::hphNT1, COG3-3xmyeGFP::kanMX4 | This study | NA |
| OGYSGA6051 | MATa, his3Δ1, leu2Δ0, ura3Δ0, LYS+, can1Δ::STE2pr-LEU2, lyp1Δ::, tor1-1, fpr1Δ::klURA, Tub4-(6)-RFP-(24)-FKBP::natNT2 COG1-FRB::hphNT1, COG4-3xmyeGFP::kanMX4 | This study | NA |
| OGYSGA6052 | MATa, his3Δ1, leu2Δ0, ura3Δ0, LYS+, can1Δ::STE2pr-LEU2, lyp1Δ::, tor1-1, fpr1Δ::klURA, Tub4-(6)-RFP-(24)-FKBP::natNT2 COG1-FRB::hphNT1, COG5-3xmyeGFP::kanMX4 | This study | NA |
| OGYSGA6053 | MATa, his3Δ1, leu2Δ0, ura3Δ0, LYS+, can1Δ::STE2pr-LEU2, lyp1Δ::, tor1-1, fpr1Δ::klURA, Tub4-(6)-RFP-(24)-FKBP::natNT2 COG1-FRB::hphNT1, COG6-3xmyeGFP::kanMX4 | This study | 542.47 |
| OGYSGA6054 | MATa, his3Δ1, leu2Δ0, ura3Δ0, LYS+, can1Δ::STE2pr-LEU2, lyp1Δ::, tor1-1, fpr1Δ::klURA, Tub4-(6)-RFP-(24)-FKBP::natNT2 COG1-FRB::hphNT1, COG7-3xmyeGFP::kanMX4 | This study | 268.77 |
| OGYSGA6055 | MATa, his3Δ1, leu2Δ0, ura3Δ0, LYS+, can1Δ::STE2pr-LEU2, lyp1Δ::, tor1-1, fpr1Δ::klURA, Tub4-(6)-RFP-(24)-FKBP::natNT2 COG1-FRB::hphNT1, COG8-3xmyeGFP::kanMX4 | This study | 560.03 |
| OGYSGA6056 | MATa, his3Δ1, leu2Δ0, ura3Δ0, LYS+, can1Δ::STE2pr-LEU2, lyp1Δ::, tor1-1, fpr1Δ::klURA, Tub4-(6)-RFP-(24)-FKBP::natNT2 COG2-FRB::hphNT1, COG1-3xmyeGFP::kanMX4 | This study | 591.97 |
| OGYSGA6057 | MATa, his3Δ1, leu2Δ0, ura3Δ0, LYS+, can1Δ::STE2pr-LEU2, lyp1Δ::, tor1-1, fpr1Δ::klURA, Tub4-(6)-RFP-(24)-FKBP::natNT2 COG2-FRB::hphNT1, COG3-3xmyeGFP::kanMX4 | This study | 525.09 |
| OGYSGA6058 | MATa, his3Δ1, leu2Δ0, ura3Δ0, LYS+, can1Δ::STE2pr-LEU2, lyp1Δ::, tor1-1, fpr1Δ::klURA, Tub4-(6)-RFP-(24)-FKBP::natNT2 COG2-FRB::hphNT1, COG4-3xmyeGFP::kanMX4 | This study | 515.09 |
| OGYSGA6059 | MATa, his3Δ1, leu2Δ0, ura3Δ0, LYS+, can1Δ::STE2pr-LEU2, lyp1Δ::, tor1-1, fpr1Δ::klURA, Tub4-(6)-RFP-(24)-FKBP::natNT2 COG2-FRB::hphNT1, COG5-3xmyeGFP::kanMX4 | This study | 612.78 |
| OGYSGA6060 | MATa, his3Δ1, leu2Δ0, ura3Δ0, LYS+, can1Δ::STE2pr-LEU2, lyp1Δ::, tor1-1, fpr1Δ::klURA, Tub4-(6)-RFP-(24)-FKBP::natNT2 COG2-FRB::hphNT1, COG6-3xmyeGFP::kanMX4 | This study | 522.08 |
| OGYSGA6061 | MATa, his3Δ1, leu2Δ0, ura3Δ0, LYS+, can1Δ::STE2pr-LEU2, lyp1Δ::, tor1-1, fpr1Δ::klURA, Tub4-(6)-RFP-(24)-FKBP::natNT2 COG2-FRB::hphNT1, COG7-3xmyeGFP::kanMX4 | This study | 278.83 |
| OGYSGA6062 | MATa, his3Δ1, leu2Δ0, ura3Δ0, LYS+, can1Δ::STE2pr-LEU2, lyp1Δ::, tor1-1, fpr1Δ::klURA, Tub4-(6)-RFP-(24)-FKBP::natNT2 COG2-FRB::hphNT1, COG8-3xmyeGFP::kanMX4 | This study | 561.6 |
| OGYSGA6063 | MATa, his3Δ1, leu2Δ0, ura3Δ0, LYS+, can1Δ::STE2pr-LEU2, lyp1Δ::, tor1-1, fpr1Δ::klURA, Tub4-(6)-RFP-(24)-FKBP::natNT2 COG3-FRB::hphNT1, COG1-3xmyeGFP::kanMX4 | This study | NA |
| OGYSGA6064 | MATa, his3Δ1, leu2Δ0, ura3Δ0, LYS+, can1Δ::STE2pr-LEU2, lyp1Δ::, tor1-1, fpr1Δ::klURA, Tub4-(6)-RFP-(24)-FKBP::natNT2 COG3-FRB::hphNT1, COG2-3xmyeGFP::kanMX4 | This study | 659.31 |
| OGYSGA6065 | MATa, his3Δ1, leu2Δ0, ura3Δ0, LYS+, can1Δ::STE2pr-LEU2, lyp1Δ::, tor1-1, fpr1Δ::klURA, Tub4-(6)-RFP-(24)-FKBP::natNT2 COG3-FRB::hphNT1, COG4-3xmyeGFP::kanMX4 | This study | 620.51 |
| OGYSGA6066 | MATa, his3Δ1, leu2Δ0, ura3Δ0, LYS+, can1Δ::STE2pr-LEU2, lyp1Δ::, tor1-1, fpr1Δ::klURA, Tub4-(6)-RFP-(24)-FKBP::natNT2 COG3-FRB::hphNT1, COG5-3xmyeGFP::kanMX4 | This study | 618.01 |
| OGYSGA6067 | MATa, his3Δ1, leu2Δ0, ura3Δ0, LYS+, can1Δ::STE2pr-LEU2, lyp1Δ::, tor1-1, fpr1Δ::klURA, Tub4-(6)-RFP-(24)-FKBP::natNT2 COG3-FRB::hphNT1, COG6-3xmyeGFP::kanMX4 | This study | 580.08 |
| OGYSGA6068 | MATa, his3Δ1, leu2Δ0, ura3Δ0, LYS+, can1Δ::STE2pr-LEU2, lyp1Δ::, tor1-1, fpr1Δ::klURA, Tub4-(6)-RFP-(24)-FKBP::natNT2 COG3-FRB::hphNT1, COG7-3xmyeGFP::kanMX4 | This study | 289.09 |
| OGYSGA6069 | MATa, his3Δ1, leu2Δ0, ura3Δ0, LYS+, can1Δ::STE2pr-LEU2, lyp1Δ::, tor1-1, fpr1Δ::klURA, Tub4-(6)-RFP-(24)-FKBP::natNT2 COG3-FRB::hphNT1, COG8-3xmyeGFP::kanMX4 | This study | 638.8 |
| OGYSGA6070 | MATa, his3Δ1, leu2Δ0, ura3Δ0, LYS+, can1Δ::STE2pr-LEU2, lyp1Δ::, tor1-1, fpr1Δ::klURA, Tub4-(6)-RFP-(24)-FKBP::natNT2 COG4-FRB::hphNT1, COG1-3xmyeGFP::kanMX4 | This study | NA |
| OGYSGA6071 | MATa, his3Δ1, leu2Δ0, ura3Δ0, LYS+, can1Δ::STE2pr-LEU2, lyp1Δ::, tor1-1, fpr1Δ::klURA, Tub4-(6)-RFP-(24)-FKBP::natNT2 COG4-FRB::hphNT1, COG2-3xmyeGFP::kanMX4 | This study | 627.81 |
| OGYSGA6072 | MATa, his3Δ1, leu2Δ0, ura3Δ0, LYS+, can1Δ::STE2pr-LEU2, lyp1Δ::, tor1-1, fpr1Δ::klURA, Tub4-(6)-RFP-(24)-FKBP::natNT2 COG4-FRB::hphNT1, COG3-3xmyeGFP::kanMX4 | This study | 381.97 |
| OGYSGA6073 | MATa, his3Δ1, leu2Δ0, ura3Δ0, LYS+, can1Δ::STE2pr-LEU2, lyp1Δ::, tor1-1, fpr1Δ::klURA, Tub4-(6)-RFP-(24)-FKBP::natNT2 COG4-FRB::hphNT1, COG5-3xmyeGFP::kanMX4 | This study | 243.72 |
| OGYSGA6074 | MATa, his3Δ1, leu2Δ0, ura3Δ0, LYS+, can1Δ::STE2pr-LEU2, lyp1Δ::, tor1-1, fpr1Δ::klURA, Tub4-(6)-RFP-(24)-FKBP::natNT2 COG4-FRB::hphNT1, COG6-3xmyeGFP::kanMX4 | This study | 324.62 |
| OGYSGA6075 | MATa, his3Δ1, leu2Δ0, ura3Δ0, LYS+, can1Δ::STE2pr-LEU2, lyp1Δ::, tor1-1, fpr1Δ::klURA, Tub4-(6)-RFP-(24)-FKBP::natNT2 COG4-FRB::hphNT1, COG7-3xmyeGFP::kanMX4 | This study | 68.82 |
| OGYSGA6076 | MATa, his3Δ1, leu2Δ0, ura3Δ0, LYS+, can1Δ::STE2pr-LEU2, lyp1Δ::, tor1-1, fpr1Δ::klURA, Tub4-(6)-RFP-(24)-FKBP::natNT2 COG4-FRB::hphNT1, COG8-3xmyeGFP::kanMX4 | This study | 287.81 |
| OGYSGA6077 | MATa, his3Δ1, leu2Δ0, ura3Δ0, LYS+, can1Δ::STE2pr-LEU2, lyp1Δ::, tor1-1, fpr1Δ::klURA, Tub4-(6)-RFP-(24)-FKBP::natNT2 COG5-FRB::hphNT1, COG1-3xmyeGFP::kanMX4 | This study | NA |
| OGYSGA6078 | MATa, his3Δ1, leu2Δ0, ura3Δ0, LYS+, can1Δ::STE2pr-LEU2, lyp1Δ::, tor1-1, fpr1Δ::klURA, Tub4-(6)-RFP-(24)-FKBP::natNT2 COG5-FRB::hphNT1, COG2-3xmyeGFP::kanMX4 | This study | 482.73 |
| OGYSGA6079 | MATa, his3Δ1, leu2Δ0, ura3Δ0, LYS+, can1Δ::STE2pr-LEU2, lyp1Δ::, tor1-1, fpr1Δ::klURA, Tub4-(6)-RFP-(24)-FKBP::natNT2 COG5-FRB::hphNT1, COG3-3xmyeGFP::kanMX4 | This study | 399.96 |
| OGYSGA6080 | MATa, his3Δ1, leu2Δ0, ura3Δ0, LYS+, can1Δ::STE2pr-LEU2, lyp1Δ::, tor1-1, fpr1Δ::klURA, Tub4-(6)-RFP-(24)-FKBP::natNT2 COG5-FRB::hphNT1, COG4-3xmyeGFP::kanMX4 | This study | 338 |

| | | | |
|---|---|---|---|
| OGYSGA6081 | *MATa, his3Δ1, leu2Δ0, ura3Δ0, LYS+, can1Δ::STE2pr-LEU2, lyp1Δ::, tor1-1, fpr1Δ::klURA, Tub4-(6)-RFP-(24)-FKBP::natNT2 COG5-FRB::hphNT1, COG6-3xmyeGFP::kanMX4* | This study | 488.86 |
| OGYSGA6082 | *MATa, his3Δ1, leu2Δ0, ura3Δ0, LYS+, can1Δ::STE2pr-LEU2, lyp1Δ::, tor1-1, fpr1Δ::klURA, Tub4-(6)-RFP-(24)-FKBP::natNT2 COG5-FRB::hphNT1, COG7-3xmyeGFP::kanMX4* | This study | 585.08 |
| OGYSGA6083 | *MATa, his3Δ1, leu2Δ0, ura3Δ0, LYS+, can1Δ::STE2pr-LEU2, lyp1Δ::, tor1-1, fpr1Δ::klURA, Tub4-(6)-RFP-(24)-FKBP::natNT2 COG5-FRB::hphNT1, COG8-3xmyeGFP::kanMX4* | This study | 520.38 |
| OGYSGA6084 | *MATa, his3Δ1, leu2Δ0, ura3Δ0, LYS+, can1Δ::STE2pr-LEU2, lyp1Δ::, tor1-1, fpr1Δ::klURA, Tub4-(6)-RFP-(24)-FKBP::natNT2 COG6-FRB::hphNT1, COG1-3xmyeGFP::kanMX4* | This study | 204.99 |
| OGYSGA6085 | *MATa, his3Δ1, leu2Δ0, ura3Δ0, LYS+, can1Δ::STE2pr-LEU2, lyp1Δ::, tor1-1, fpr1Δ::klURA, Tub4-(6)-RFP-(24)-FKBP::natNT2 COG6-FRB::hphNT1, COG2-3xmyeGFP::kanMX4* | This study | 559.35 |
| OGYSGA6086 | *MATa, his3Δ1, leu2Δ0, ura3Δ0, LYS+, can1Δ::STE2pr-LEU2, lyp1Δ::, tor1-1, fpr1Δ::klURA, Tub4-(6)-RFP-(24)-FKBP::natNT2 COG6-FRB::hphNT1, COG3-3xmyeGFP::kanMX4* | This study | 519 |
| OGYSGA6087 | *MATa, his3Δ1, leu2Δ0, ura3Δ0, LYS+, can1Δ::STE2pr-LEU2, lyp1Δ::, tor1-1, fpr1Δ::klURA, Tub4-(6)-RFP-(24)-FKBP::natNT2 COG6-FRB::hphNT1, COG4-3xmyeGFP::kanMX4* | This study | 445.83 |
| OGYSGA6088 | *MATa, his3Δ1, leu2Δ0, ura3Δ0, LYS+, can1Δ::STE2pr-LEU2, lyp1Δ::, tor1-1, fpr1Δ::klURA, Tub4-(6)-RFP-(24)-FKBP::natNT2 COG6-FRB::hphNT1, COG5-3xmyeGFP::kanMX4* | This study | 599.04 |
| OGYSGA6089 | *MATa, his3Δ1, leu2Δ0, ura3Δ0, LYS+, can1Δ::STE2pr-LEU2, lyp1Δ::, tor1-1, fpr1Δ::klURA, Tub4-(6)-RFP-(24)-FKBP::natNT2 COG6-FRB::hphNT1, COG7-3xmyeGFP::kanMX4* | This study | 275.15 |
| OGYSGA6090 | *MATa, his3Δ1, leu2Δ0, ura3Δ0, LYS+, can1Δ::STE2pr-LEU2, lyp1Δ::, tor1-1, fpr1Δ::klURA, Tub4-(6)-RFP-(24)-FKBP::natNT2 COG6-FRB::hphNT1, COG8-3xmyeGFP::kanMX4* | This study | 582.41 |
| OGYSGA6091 | *MATa, his3Δ1, leu2Δ0, ura3Δ0, LYS+, can1Δ::STE2pr-LEU2, lyp1Δ::, tor1-1, fpr1Δ::klURA, Tub4-(6)-RFP-(24)-FKBP::natNT2 COG7-FRB::hphNT1, COG1-3xmyeGFP::kanMX4* | This study | NA |
| OGYSGA6092 | *MATa, his3Δ1, leu2Δ0, ura3Δ0, LYS+, can1Δ::STE2pr-LEU2, lyp1Δ::, tor1-1, fpr1Δ::klURA, Tub4-(6)-RFP-(24)-FKBP::natNT2 COG7-FRB::hphNT1, COG2-3xmyeGFP::kanMX4* | This study | 191.27 |
| OGYSGA6093 | *MATa, his3Δ1, leu2Δ0, ura3Δ0, LYS+, can1Δ::STE2pr-LEU2, lyp1Δ::, tor1-1, fpr1Δ::klURA, Tub4-(6)-RFP-(24)-FKBP::natNT2 COG7-FRB::hphNT1, COG3-3xmyeGFP::kanMX4* | This study | 223.52 |
| OGYSGA6094 | *MATa, his3Δ1, leu2Δ0, ura3Δ0, LYS+, can1Δ::STE2pr-LEU2, lyp1Δ::, tor1-1, fpr1Δ::klURA, Tub4-(6)-RFP-(24)-FKBP::natNT2 COG7-FRB::hphNT1, COG4-3xmyeGFP::kanMX4* | This study | 119.95 |
| OGYSGA6095 | *MATa, his3Δ1, leu2Δ0, ura3Δ0, LYS+, can1Δ::STE2pr-LEU2, lyp1Δ::, tor1-1, fpr1Δ::klURA, Tub4-(6)-RFP-(24)-FKBP::natNT2 COG7-FRB::hphNT1, COG5-3xmyeGFP::kanMX4* | This study | 614.09 |
| OGYSGA6096 | *MATa, his3Δ1, leu2Δ0, ura3Δ0, LYS+, can1Δ::STE2pr-LEU2, lyp1Δ::, tor1-1, fpr1Δ::klURA, Tub4-(6)-RFP-(24)-FKBP::natNT2 COG7-FRB::hphNT1, COG6-3xmyeGFP::kanMX4* | This study | 336.62 |
| OGYSGA6097 | *MATa, his3Δ1, leu2Δ0, ura3Δ0, LYS+, can1Δ::STE2pr-LEU2, lyp1Δ::, tor1-1, fpr1Δ::klURA, Tub4-(6)-RFP-(24)-FKBP::natNT2 COG7-FRB::hphNT1, COG8-3xmyeGFP::kanMX4* | This study | 250.85 |
| OGYSGA6098 | *MATa, his3Δ1, leu2Δ0, ura3Δ0, LYS+, can1Δ::STE2pr-LEU2, lyp1Δ::, tor1-1, fpr1Δ::klURA, Tub4-(6)-RFP-(24)-FKBP::natNT2 COG8-FRB::hphNT1, COG1-3xmyeGFP::kanMX4* | This study | 4.99 |
| OGYSGA6099 | *MATa, his3Δ1, leu2Δ0, ura3Δ0, LYS+, can1Δ::STE2pr-LEU2, lyp1Δ::, tor1-1, fpr1Δ::klURA, Tub4-(6)-RFP-(24)-FKBP::natNT2 COG8-FRB::hphNT1, COG2-3xmyeGFP::kanMX4* | This study | 598.46 |
| OGYSGA6100 | *MATa, his3Δ1, leu2Δ0, ura3Δ0, LYS+, can1Δ::STE2pr-LEU2, lyp1Δ::, tor1-1, fpr1Δ::klURA, Tub4-(6)-RFP-(24)-FKBP::natNT2 COG8-FRB::hphNT1, COG3-3xmyeGFP::kanMX4* | This study | 491.67 |
| OGYSGA6101 | *MATa, his3Δ1, leu2Δ0, ura3Δ0, LYS+, can1Δ::STE2pr-LEU2, lyp1Δ::, tor1-1, fpr1Δ::klURA, Tub4-(6)-RFP-(24)-FKBP::natNT2 COG8-FRB::hphNT1, COG4-3xmyeGFP::kanMX4* | This study | 468.26 |
| OGYSGA6102 | *MATa, his3Δ1, leu2Δ0, ura3Δ0, LYS+, can1Δ::STE2pr-LEU2, lyp1Δ::, tor1-1, fpr1Δ::klURA, Tub4-(6)-RFP-(24)-FKBP::natNT2 COG8-FRB::hphNT1, COG5-3xmyeGFP::kanMX4* | This study | 631.7 |
| OGYSGA6103 | *MATa, his3Δ1, leu2Δ0, ura3Δ0, LYS+, can1Δ::STE2pr-LEU2, lyp1Δ::, tor1-1, fpr1Δ::klURA, Tub4-(6)-RFP-(24)-FKBP::natNT2 COG8-FRB::hphNT1, COG6-3xmyeGFP::kanMX4* | This study | 732.32 |
| OGYSGA6104 | *MATa, his3Δ1, leu2Δ0, ura3Δ0, LYS+, can1Δ::STE2pr-LEU2, lyp1Δ::, tor1-1, fpr1Δ::klURA, Tub4-(6)-RFP-(24)-FKBP::natNT2 COG8-FRB::hphNT1, COG7-3xmyeGFP::kanMX4* | This study | 288.24 |
| OGYSGA6105 | *MATa, his3Δ1, leu2Δ0, ura3Δ0, LYS+, can1Δ::STE2pr-LEU2, lyp1Δ::, tor1-1, fpr1Δ::klURA, Tub4-(6)-RFP-(24)-FKBP::natNT2 VPS8-FRB::hphNT1, VPS3-3xmyeGFP::kanMX4* | This study | 364.66 |
| OGYSGA6106 | *MATa, his3Δ1, leu2Δ0, ura3Δ0, LYS+, can1Δ::STE2pr-LEU2, lyp1Δ::, tor1-1, fpr1Δ::klURA, Tub4-(6)-RFP-(24)-FKBP::natNT2 VPS8-FRB::hphNT1, PEP5-3xmyeGFP::kanMX4* | This study | 379.43 |
| OGYSGA6107 | *MATa, his3Δ1, leu2Δ0, ura3Δ0, LYS+, can1Δ::STE2pr-LEU2, lyp1Δ::, tor1-1, fpr1Δ::klURA, Tub4-(6)-RFP-(24)-FKBP::natNT2 VPS8-FRB::hphNT1, VPS16-3xmyeGFP::kanMX4* | This study | NA |
| OGYSGA6108 | *MATa, his3Δ1, leu2Δ0, ura3Δ0, LYS+, can1Δ::STE2pr-LEU2, lyp1Δ::, tor1-1, fpr1Δ::klURA, Tub4-(6)-RFP-(24)-FKBP::natNT2 VPS8-FRB::hphNT1, PEP3-3xmyeGFP::kanMX4* | This study | 418.33 |
| OGYSGA6109 | *MATa, his3Δ1, leu2Δ0, ura3Δ0, LYS+, can1Δ::STE2pr-LEU2, lyp1Δ::, tor1-1, fpr1Δ::klURA, Tub4-(6)-RFP-(24)-FKBP::natNT2 VPS8-FRB::hphNT1, VPS33-3xmyeGFP::kanMX4* | This study | 366.2 |
| OGYSGA6110 | *MATa, his3Δ1, leu2Δ0, ura3Δ0, LYS+, can1Δ::STE2pr-LEU2, lyp1Δ::, tor1-1, fpr1Δ::klURA, Tub4-(6)-RFP-(24)-FKBP::natNT2 VPS3-FRB::hphNT1, VPS8-3xmyeGFP::kanMX4* | This study | 577.26 |
| OGYSGA6111 | *MATa, his3Δ1, leu2Δ0, ura3Δ0, LYS+, can1Δ::STE2pr-LEU2, lyp1Δ::, tor1-1, fpr1Δ::klURA, Tub4-(6)-RFP-(24)-FKBP::natNT2 VPS3-FRB::hphNT1, PEP5-3xmyeGFP::kanMX4* | This study | 587.65 |
| OGYSGA6112 | *MATa, his3Δ1, leu2Δ0, ura3Δ0, LYS+, can1Δ::STE2pr-LEU2, lyp1Δ::, tor1-1, fpr1Δ::klURA, Tub4-(6)-RFP-(24)-FKBP::natNT2 VPS3-FRB::hphNT1, VPS16-3xmyeGFP::kanMX4* | This study | 668.67 |
| OGYSGA6113 | *MATa, his3Δ1, leu2Δ0, ura3Δ0, LYS+, can1Δ::STE2pr-LEU2, lyp1Δ::, tor1-1, fpr1Δ::klURA, Tub4-(6)-RFP-(24)-FKBP::natNT2 VPS3-FRB::hphNT1, PEP3-3xmyeGFP::kanMX4* | This study | 764.89 |
| OGYSGA6114 | *MATa, his3Δ1, leu2Δ0, ura3Δ0, LYS+, can1Δ::STE2pr-LEU2, lyp1Δ::, tor1-1, fpr1Δ::klURA, Tub4-(6)-RFP-(24)-FKBP::natNT2 VPS3-FRB::hphNT1, VPS33-3xmyeGFP::kanMX4* | This study | 565.89 |
| OGYSGA6115 | *MATa, his3Δ1, leu2Δ0, ura3Δ0, LYS+, can1Δ::STE2pr-LEU2, lyp1Δ::, tor1-1, fpr1Δ::klURA, Tub4-(6)-RFP-(24)-FKBP::natNT2 PEP5-FRB::hphNT1, VPS8-3xmyeGFP::kanMX4* | This study | 525 |
| OGYSGA6116 | *MATa, his3Δ1, leu2Δ0, ura3Δ0, LYS+, can1Δ::STE2pr-LEU2, lyp1Δ::, tor1-1, fpr1Δ::klURA, Tub4-(6)-RFP-(24)-FKBP::natNT2 PEP5-FRB::hphNT1, VPS3-3xmyeGFP::kanMX4* | This study | 611.05 |
| OGYSGA6117 | *MATa, his3Δ1, leu2Δ0, ura3Δ0, LYS+, can1Δ::STE2pr-LEU2, lyp1Δ::, tor1-1, fpr1Δ::klURA, Tub4-(6)-RFP-(24)-FKBP::natNT2 PEP5-FRB::hphNT1, VPS16-3xmyeGFP::kanMX4* | This study | NA |
| OGYSGA6118 | *MATa, his3Δ1, leu2Δ0, ura3Δ0, LYS+, can1Δ::STE2pr-LEU2, lyp1Δ::, tor1-1, fpr1Δ::klURA, Tub4-(6)-RFP-(24)-FKBP::natNT2 PEP5-FRB::hphNT1, PEP3-3xmyeGFP::kanMX4* | This study | 970.45 |
| OGYSGA6119 | *MATa, his3Δ1, leu2Δ0, ura3Δ0, LYS+, can1Δ::STE2pr-LEU2, lyp1Δ::, tor1-1, fpr1Δ::klURA, Tub4-(6)-RFP-(24)-FKBP::natNT2 PEP5-FRB::hphNT1, VPS33-3xmyeGFP::kanMX4* | This study | 806.8 |
| OGYSGA6120 | *MATa, his3Δ1, leu2Δ0, ura3Δ0, LYS+, can1Δ::STE2pr-LEU2, lyp1Δ::, tor1-1, fpr1Δ::klURA, Tub4-(6)-RFP-(24)-FKBP::natNT2 VPS16-FRB::hphNT1, VPS8-3xmyeGFP::kanMX4* | This study | 367.36 |

| | | | |
|---|---|---|---|
| OGYSGA6121 | *MATa, his3Δ1, leu2Δ0, ura3Δ0, LYS+, can1Δ::STE2pr-LEU2, lyp1Δ::, tor1-1, fpr1Δ::klURA, Tub4-(6)-RFP-(24)-FKBP::natNT2 VPS16-FRB::hphNT1, VPS3-3xmyeGFP::kanMX4* | This study | 473.25 |
| OGYSGA6122 | *MATa, his3Δ1, leu2Δ0, ura3Δ0, LYS+, can1Δ::STE2pr-LEU2, lyp1Δ::, tor1-1, fpr1Δ::klURA, Tub4-(6)-RFP-(24)-FKBP::natNT2 VPS16-FRB::hphNT1, PEP5-3xmyeGFP::kanMX4* | This study | 619.71 |
| OGYSGA6123 | *MATa, his3Δ1, leu2Δ0, ura3Δ0, LYS+, can1Δ::STE2pr-LEU2, lyp1Δ::, tor1-1, fpr1Δ::klURA, Tub4-(6)-RFP-(24)-FKBP::natNT2 VPS16-FRB::hphNT1, PEP3-3xmyeGFP::kanMX4* | This study | 615.04 |
| OGYSGA6124 | *MATa, his3Δ1, leu2Δ0, ura3Δ0, LYS+, can1Δ::STE2pr-LEU2, lyp1Δ::, tor1-1, fpr1Δ::klURA, Tub4-(6)-RFP-(24)-FKBP::natNT2 VPS16-FRB::hphNT1, VPS33-3xmyeGFP::kanMX4* | This study | 727.21 |
| OGYSGA6125 | *MATa, his3Δ1, leu2Δ0, ura3Δ0, LYS+, can1Δ::STE2pr-LEU2, lyp1Δ::, tor1-1, fpr1Δ::klURA, Tub4-(6)-RFP-(24)-FKBP::natNT2 PEP3-FRB::hphNT1, VPS8-3xmyeGFP::kanMX4* | This study | 23.79 |
| OGYSGA6126 | *MATa, his3Δ1, leu2Δ0, ura3Δ0, LYS+, can1Δ::STE2pr-LEU2, lyp1Δ::, tor1-1, fpr1Δ::klURA, Tub4-(6)-RFP-(24)-FKBP::natNT2 PEP3-FRB::hphNT1, VPS3-3xmyeGFP::kanMX4* | This study | NA |
| OGYSGA6127 | *MATa, his3Δ1, leu2Δ0, ura3Δ0, LYS+, can1Δ::STE2pr-LEU2, lyp1Δ::, tor1-1, fpr1Δ::klURA, Tub4-(6)-RFP-(24)-FKBP::natNT2 PEP3-FRB::hphNT1, PEP5-3xmyeGFP::kanMX4* | This study | 676.39 |
| OGYSGA6128 | *MATa, his3Δ1, leu2Δ0, ura3Δ0, LYS+, can1Δ::STE2pr-LEU2, lyp1Δ::, tor1-1, fpr1Δ::klURA, Tub4-(6)-RFP-(24)-FKBP::natNT2 PEP3-FRB::hphNT1, VPS16-3xmyeGFP::kanMX4* | This study | NA |
| OGYSGA6129 | *MATa, his3Δ1, leu2Δ0, ura3Δ0, LYS+, can1Δ::STE2pr-LEU2, lyp1Δ::, tor1-1, fpr1Δ::klURA, Tub4-(6)-RFP-(24)-FKBP::natNT2 PEP3-FRB::hphNT1, VPS33-3xmyeGFP::kanMX4* | This study | NA |
| OGYSGA6130 | *MATa, his3Δ1, leu2Δ0, ura3Δ0, LYS+, can1Δ::STE2pr-LEU2, lyp1Δ::, tor1-1, fpr1Δ::klURA, Tub4-(6)-RFP-(24)-FKBP::natNT2 VPS33-FRB::hphNT1, VPS8-3xmyeGFP::kanMX4* | This study | 246.07 |
| OGYSGA6131 | *MATa, his3Δ1, leu2Δ0, ura3Δ0, LYS+, can1Δ::STE2pr-LEU2, lyp1Δ::, tor1-1, fpr1Δ::klURA, Tub4-(6)-RFP-(24)-FKBP::natNT2 VPS33-FRB::hphNT1, VPS3-3xmyeGFP::kanMX4* | This study | NA |
| OGYSGA6132 | *MATa, his3Δ1, leu2Δ0, ura3Δ0, LYS+, can1Δ::STE2pr-LEU2, lyp1Δ::, tor1-1, fpr1Δ::klURA, Tub4-(6)-RFP-(24)-FKBP::natNT2 VPS33-FRB::hphNT1, PEP5-3xmyeGFP::kanMX4* | This study | 377.1 |
| OGYSGA6133 | *MATa, his3Δ1, leu2Δ0, ura3Δ0, LYS+, can1Δ::STE2pr-LEU2, lyp1Δ::, tor1-1, fpr1Δ::klURA, Tub4-(6)-RFP-(24)-FKBP::natNT2 VPS33-FRB::hphNT1, VPS16-3xmyeGFP::kanMX4* | This study | 3.44 |
| OGYSGA6134 | *MATa, his3Δ1, leu2Δ0, ura3Δ0, LYS+, can1Δ::STE2pr-LEU2, lyp1Δ::, tor1-1, fpr1Δ::klURA, Tub4-(6)-RFP-(24)-FKBP::natNT2 VPS33-FRB::hphNT1, PEP3-3xmyeGFP::kanMX4* | This study | 304.544 |
| OGYSGA6135 | *MATa, his3Δ1, leu2Δ0, ura3Δ0, LYS+, can1Δ::STE2pr-LEU2, lyp1Δ::, tor1-1, fpr1Δ::klURA, Tub4-(6)-RFP-(24)-FKBP::natNT2 PEP5-FRB::hphNT1, VPS41-3xmyeGFP::kanMX4* | This study | 218.09 |
| OGYSGA6136 | *MATa, his3Δ1, leu2Δ0, ura3Δ0, LYS+, can1Δ::STE2pr-LEU2, lyp1Δ::, tor1-1, fpr1Δ::klURA, Tub4-(6)-RFP-(24)-FKBP::natNT2 PEP5-FRB::hphNT1, VAM6-3xmyeGFP::kanMX4* | This study | 114.75 |
| OGYSGA6137 | *MATa, his3Δ1, leu2Δ0, ura3Δ0, LYS+, can1Δ::STE2pr-LEU2, lyp1Δ::, tor1-1, fpr1Δ::klURA, Tub4-(6)-RFP-(24)-FKBP::natNT2 VPS16-FRB::hphNT1, VPS41-3xmyeGFP::kanMX4* | This study | 208.12 |
| OGYSGA6138 | *MATa, his3Δ1, leu2Δ0, ura3Δ0, LYS+, can1Δ::STE2pr-LEU2, lyp1Δ::, tor1-1, fpr1Δ::klURA, Tub4-(6)-RFP-(24)-FKBP::natNT2 VPS16-FRB::hphNT1, VAM6-3xmyeGFP::kanMX4* | This study | 61.88 |
| OGYSGA6139 | *MATa, his3Δ1, leu2Δ0, ura3Δ0, LYS+, can1Δ::STE2pr-LEU2, lyp1Δ::, tor1-1, fpr1Δ::klURA, Tub4-(6)-RFP-(24)-FKBP::natNT2 PEP3-FRB::hphNT1, VPS41-3xmyeGFP::kanMX4* | This study | NA |
| OGYSGA6140 | *MATa, his3Δ1, leu2Δ0, ura3Δ0, LYS+, can1Δ::STE2pr-LEU2, lyp1Δ::, tor1-1, fpr1Δ::klURA, Tub4-(6)-RFP-(24)-FKBP::natNT2 PEP3-FRB::hphNT1, VAM6-3xmyeGFP::kanMX4* | This study | NA |
| OGYSGA6141 | *MATa, his3Δ1, leu2Δ0, ura3Δ0, LYS+, can1Δ::STE2pr-LEU2, lyp1Δ::, tor1-1, fpr1Δ::klURA, Tub4-(6)-RFP-(24)-FKBP::natNT2 VPS33-FRB::hphNT1, VPS41-3xmyeGFP::kanMX4* | This study | NA |
| OGYSGA6142 | *MATa, his3Δ1, leu2Δ0, ura3Δ0, LYS+, can1Δ::STE2pr-LEU2, lyp1Δ::, tor1-1, fpr1Δ::klURA, Tub4-(6)-RFP-(24)-FKBP::natNT2 VPS33-FRB::hphNT1, VAM6-3xmyeGFP::kanMX4* | This study | NA |
| OGYSGA6143 | *MATa, his3Δ1, leu2Δ0, ura3Δ0, LYS+, can1Δ::STE2pr-LEU2, lyp1Δ::, tor1-1, fpr1Δ::klURA, Tub4-(6)-RFP-(24)-FKBP::natNT2 VPS41-FRB::hphNT1, PEP5-3xmyeGFP::kanMX4* | This study | 6.03 |
| OGYSGA6144 | *MATa, his3Δ1, leu2Δ0, ura3Δ0, LYS+, can1Δ::STE2pr-LEU2, lyp1Δ::, tor1-1, fpr1Δ::klURA, Tub4-(6)-RFP-(24)-FKBP::natNT2 VPS41-FRB::hphNT1, VPS16-3xmyeGFP::kanMX4* | This study | 2.16 |
| OGYSGA6145 | *MATa, his3Δ1, leu2Δ0, ura3Δ0, LYS+, can1Δ::STE2pr-LEU2, lyp1Δ::, tor1-1, fpr1Δ::klURA, Tub4-(6)-RFP-(24)-FKBP::natNT2 VPS41-FRB::hphNT1, PEP3-3xmyeGFP::kanMX4* | This study | 5.7 |
| OGYSGA6146 | *MATa, his3Δ1, leu2Δ0, ura3Δ0, LYS+, can1Δ::STE2pr-LEU2, lyp1Δ::, tor1-1, fpr1Δ::klURA, Tub4-(6)-RFP-(24)-FKBP::natNT2 VPS41-FRB::hphNT1, VPS33-3xmyeGFP::kanMX4* | This study | -0.24 |
| OGYSGA6147 | *MATa, his3Δ1, leu2Δ0, ura3Δ0, LYS+, can1Δ::STE2pr-LEU2, lyp1Δ::, tor1-1, fpr1Δ::klURA, Tub4-(6)-RFP-(24)-FKBP::natNT2 VPS41-FRB::hphNT1, VAM6-3xmyeGFP::kanMX4* | This study | 4.43 |
| OGYSGA6148 | *MATa, his3Δ1, leu2Δ0, ura3Δ0, LYS+, can1Δ::STE2pr-LEU2, lyp1Δ::, tor1-1, fpr1Δ::klURA, Tub4-(6)-RFP-(24)-FKBP::natNT2 VAM6-FRB::hphNT1, PEP5-3xmyeGFP::kanMX4* | This study | 187.68 |
| OGYSGA6149 | *MATa, his3Δ1, leu2Δ0, ura3Δ0, LYS+, can1Δ::STE2pr-LEU2, lyp1Δ::, tor1-1, fpr1Δ::klURA, Tub4-(6)-RFP-(24)-FKBP::natNT2 VAM6-FRB::hphNT1, VPS16-3xmyeGFP::kanMX4* | This study | NA |
| OGYSGA6150 | *MATa, his3Δ1, leu2Δ0, ura3Δ0, LYS+, can1Δ::STE2pr-LEU2, lyp1Δ::, tor1-1, fpr1Δ::klURA, Tub4-(6)-RFP-(24)-FKBP::natNT2 VAM6-FRB::hphNT1, PEP3-3xmyeGFP::kanMX4* | This study | 177.26 |
| OGYSGA6151 | *MATa, his3Δ1, leu2Δ0, ura3Δ0, LYS+, can1Δ::STE2pr-LEU2, lyp1Δ::, tor1-1, fpr1Δ::klURA, Tub4-(6)-RFP-(24)-FKBP::natNT2 VAM6-FRB::hphNT1, VPS33-3xmyeGFP::kanMX4* | This study | 183.32 |
| OGYSGA6152 | *MATa, his3Δ1, leu2Δ0, ura3Δ0, LYS+, can1Δ::STE2pr-LEU2, lyp1Δ::, tor1-1, fpr1Δ::klURA, Tub4-(6)-RFP-(24)-FKBP::natNT2 VAM6-FRB::hphNT1, VPS41-3xmyeGFP::kanMX4* | This study | 80.82 |
| OGYSGA6153 | *MATa, his3Δ1, leu2Δ0, ura3Δ0, LYS+, can1Δ::STE2pr-LEU2, lyp1Δ::, tor1-1, fpr1Δ::klURA, Tub4-(6)-RFP-(24)-FKBP::natNT2 BET3-FRB::hphNT1, BET5-3xmyeGFP::kanMX4* | This study | 1740.38 |
| OGYSGA6154 | *MATa, his3Δ1, leu2Δ0, ura3Δ0, LYS+, can1Δ::STE2pr-LEU2, lyp1Δ::, tor1-1, fpr1Δ::klURA, Tub4-(6)-RFP-(24)-FKBP::natNT2 BET3-FRB::hphNT1, TRS20-3xmyeGFP::kanMX4* | This study | 989.68 |
| OGYSGA6155 | *MATa, his3Δ1, leu2Δ0, ura3Δ0, LYS+, can1Δ::STE2pr-LEU2, lyp1Δ::, tor1-1, fpr1Δ::klURA, Tub4-(6)-RFP-(24)-FKBP::natNT2 BET3-FRB::hphNT1, TRS23-3xmyeGFP::kanMX4* | This study | 2154.43 |
| OGYSGA6156 | *MATa, his3Δ1, leu2Δ0, ura3Δ0, LYS+, can1Δ::STE2pr-LEU2, lyp1Δ::, tor1-1, fpr1Δ::klURA, Tub4-(6)-RFP-(24)-FKBP::natNT2 BET3-FRB::hphNT1, TRS31-3xmyeGFP::kanMX4* | This study | NA |
| OGYSGA6157 | *MATa, his3Δ1, leu2Δ0, ura3Δ0, LYS+, can1Δ::STE2pr-LEU2, lyp1Δ::, tor1-1, fpr1Δ::klURA, Tub4-(6)-RFP-(24)-FKBP::natNT2 BET3-FRB::hphNT1, TRS33-3xmyeGFP::kanMX4* | This study | 1823.61 |
| OGYSGA6158 | *MATa, his3Δ1, leu2Δ0, ura3Δ0, LYS+, can1Δ::STE2pr-LEU2, lyp1Δ::, tor1-1, fpr1Δ::klURA, Tub4-(6)-RFP-(24)-FKBP::natNT2 BET5-FRB::hphNT1, BET3-3xmyeGFP::kanMX4* | This study | 2015.43 |
| OGYSGA6159 | *MATa, his3Δ1, leu2Δ0, ura3Δ0, LYS+, can1Δ::STE2pr-LEU2, lyp1Δ::, tor1-1, fpr1Δ::klURA, Tub4-(6)-RFP-(24)-FKBP::natNT2 BET5-FRB::hphNT1, TRS20-3xmyeGFP::kanMX4* | This study | 890.4 |
| OGYSGA6160 | *MATa, his3Δ1, leu2Δ0, ura3Δ0, LYS+, can1Δ::STE2pr-LEU2, lyp1Δ::, tor1-1, fpr1Δ::klURA, Tub4-(6)-RFP-(24)-FKBP::natNT2 BET5-FRB::hphNT1, TRS23-3xmyeGFP::kanMX4* | This study | 2175.52 |

| | | | |
|---|---|---|---|
| OGYSGA6161 | *MATa, his3Δ1, leu2Δ0, ura3Δ0, LYS+, can1Δ::STE2pr-LEU2, lyp1Δ::, tor1-1, fpr1Δ::klURA, Tub4-(6)-RFP-(24)-FKBP::natNT2 BET5-FRB::hphNT1, TRS31-3xmyeGFP::kanMX4* | This study | 2284.72 |
| OGYSGA6162 | *MATa, his3Δ1, leu2Δ0, ura3Δ0, LYS+, can1Δ::STE2pr-LEU2, lyp1Δ::, tor1-1, fpr1Δ::klURA, Tub4-(6)-RFP-(24)-FKBP::natNT2 BET5-FRB::hphNT1, TRS33-3xmyeGFP::kanMX4* | This study | 1811.77 |
| OGYSGA6163 | *MATa, his3Δ1, leu2Δ0, ura3Δ0, LYS+, can1Δ::STE2pr-LEU2, lyp1Δ::, tor1-1, fpr1Δ::klURA, Tub4-(6)-RFP-(24)-FKBP::natNT2 TRS20-FRB::hphNT1, BET3-3xmyeGFP::kanMX4* | This study | NA |
| OGYSGA6164 | *MATa, his3Δ1, leu2Δ0, ura3Δ0, LYS+, can1Δ::STE2pr-LEU2, lyp1Δ::, tor1-1, fpr1Δ::klURA, Tub4-(6)-RFP-(24)-FKBP::natNT2 TRS20-FRB::hphNT1, BET5-3xmyeGFP::kanMX4* | This study | -1.42 |
| OGYSGA6165 | *MATa, his3Δ1, leu2Δ0, ura3Δ0, LYS+, can1Δ::STE2pr-LEU2, lyp1Δ::, tor1-1, fpr1Δ::klURA, Tub4-(6)-RFP-(24)-FKBP::natNT2 TRS20-FRB::hphNT1, TRS23-3xmyeGFP::kanMX4* | This study | NA |
| OGYSGA6166 | *MATa, his3Δ1, leu2Δ0, ura3Δ0, LYS+, can1Δ::STE2pr-LEU2, lyp1Δ::, tor1-1, fpr1Δ::klURA, Tub4-(6)-RFP-(24)-FKBP::natNT2 TRS20-FRB::hphNT1, TRS31-3xmyeGFP::kanMX4* | This study | -2.81 |
| OGYSGA6167 | *MATa, his3Δ1, leu2Δ0, ura3Δ0, LYS+, can1Δ::STE2pr-LEU2, lyp1Δ::, tor1-1, fpr1Δ::klURA, Tub4-(6)-RFP-(24)-FKBP::natNT2 TRS20-FRB::hphNT1, TRS33-3xmyeGFP::kanMX4* | This study | 1277.32 |
| OGYSGA6168 | *MATa, his3Δ1, leu2Δ0, ura3Δ0, LYS+, can1Δ::STE2pr-LEU2, lyp1Δ::, tor1-1, fpr1Δ::klURA, Tub4-(6)-RFP-(24)-FKBP::natNT2 TRS23-FRB::hphNT1, BET3-3xmyeGFP::kanMX4* | This study | 1268.58 |
| OGYSGA6169 | *MATa, his3Δ1, leu2Δ0, ura3Δ0, LYS+, can1Δ::STE2pr-LEU2, lyp1Δ::, tor1-1, fpr1Δ::klURA, Tub4-(6)-RFP-(24)-FKBP::natNT2 TRS23-FRB::hphNT1, BET5-3xmyeGFP::kanMX4* | This study | 1650.58 |
| OGYSGA6170 | *MATa, his3Δ1, leu2Δ0, ura3Δ0, LYS+, can1Δ::STE2pr-LEU2, lyp1Δ::, tor1-1, fpr1Δ::klURA, Tub4-(6)-RFP-(24)-FKBP::natNT2 TRS23-FRB::hphNT1, TRS20-3xmyeGFP::kanMX4* | This study | 822.03 |
| OGYSGA6171 | *MATa, his3Δ1, leu2Δ0, ura3Δ0, LYS+, can1Δ::STE2pr-LEU2, lyp1Δ::, tor1-1, fpr1Δ::klURA, Tub4-(6)-RFP-(24)-FKBP::natNT2 TRS23-FRB::hphNT1, TRS31-3xmyeGFP::kanMX4* | This study | NA |
| OGYSGA6172 | *MATa, his3Δ1, leu2Δ0, ura3Δ0, LYS+, can1Δ::STE2pr-LEU2, lyp1Δ::, tor1-1, fpr1Δ::klURA, Tub4-(6)-RFP-(24)-FKBP::natNT2 TRS23-FRB::hphNT1, TRS33-3xmyeGFP::kanMX4* | This study | 1514.56 |
| OGYSGA6173 | *MATa, his3Δ1, leu2Δ0, ura3Δ0, LYS+, can1Δ::STE2pr-LEU2, lyp1Δ::, tor1-1, fpr1Δ::klURA, Tub4-(6)-RFP-(24)-FKBP::natNT2 TRS31-FRB::hphNT1, BET3-3xmyeGFP::kanMX4* | This study | 709.79 |
| OGYSGA6174 | *MATa, his3Δ1, leu2Δ0, ura3Δ0, LYS+, can1Δ::STE2pr-LEU2, lyp1Δ::, tor1-1, fpr1Δ::klURA, Tub4-(6)-RFP-(24)-FKBP::natNT2 TRS31-FRB::hphNT1, BET5-3xmyeGFP::kanMX4* | This study | 2039.67 |
| OGYSGA6175 | *MATa, his3Δ1, leu2Δ0, ura3Δ0, LYS+, can1Δ::STE2pr-LEU2, lyp1Δ::, tor1-1, fpr1Δ::klURA, Tub4-(6)-RFP-(24)-FKBP::natNT2 TRS31-FRB::hphNT1, TRS20-3xmyeGFP::kanMX4* | This study | 1064.3 |
| OGYSGA6176 | *MATa, his3Δ1, leu2Δ0, ura3Δ0, LYS+, can1Δ::STE2pr-LEU2, lyp1Δ::, tor1-1, fpr1Δ::klURA, Tub4-(6)-RFP-(24)-FKBP::natNT2 TRS31-FRB::hphNT1, TRS23-3xmyeGFP::kanMX4* | This study | 1694.63 |
| OGYSGA6177 | *MATa, his3Δ1, leu2Δ0, ura3Δ0, LYS+, can1Δ::STE2pr-LEU2, lyp1Δ::, tor1-1, fpr1Δ::klURA, Tub4-(6)-RFP-(24)-FKBP::natNT2 TRS31-FRB::hphNT1, TRS33-3xmyeGFP::kanMX4* | This study | 1813.68 |
| OGYSGA6178 | *MATa, his3Δ1, leu2Δ0, ura3Δ0, LYS+, can1Δ::STE2pr-LEU2, lyp1Δ::, tor1-1, fpr1Δ::klURA, Tub4-(6)-RFP-(24)-FKBP::natNT2 TRS33-FRB::hphNT1, BET3-3xmyeGFP::kanMX4* | This study | 1316.29 |
| OGYSGA6179 | *MATa, his3Δ1, leu2Δ0, ura3Δ0, LYS+, can1Δ::STE2pr-LEU2, lyp1Δ::, tor1-1, fpr1Δ::klURA, Tub4-(6)-RFP-(24)-FKBP::natNT2 TRS33-FRB::hphNT1, BET5-3xmyeGFP::kanMX4* | This study | 2106.21 |
| OGYSGA6180 | *MATa, his3Δ1, leu2Δ0, ura3Δ0, LYS+, can1Δ::STE2pr-LEU2, lyp1Δ::, tor1-1, fpr1Δ::klURA, Tub4-(6)-RFP-(24)-FKBP::natNT2 TRS33-FRB::hphNT1, TRS20-3xmyeGFP::kanMX4* | This study | 941.83 |
| OGYSGA6181 | *MATa, his3Δ1, leu2Δ0, ura3Δ0, LYS+, can1Δ::STE2pr-LEU2, lyp1Δ::, tor1-1, fpr1Δ::klURA, Tub4-(6)-RFP-(24)-FKBP::natNT2 TRS33-FRB::hphNT1, TRS23-3xmyeGFP::kanMX4* | This study | 2225.52 |
| OGYSGA6182 | *MATa, his3Δ1, leu2Δ0, ura3Δ0, LYS+, can1Δ::STE2pr-LEU2, lyp1Δ::, tor1-1, fpr1Δ::klURA, Tub4-(6)-RFP-(24)-FKBP::natNT2 TRS33-FRB::hphNT1, TRS31-3xmyeGFP::kanMX4* | This study | 2298.4 |
| OGYSGA6183 | *MATa, his3Δ1, leu2Δ0, ura3Δ0, LYS+, can1Δ::STE2pr-LEU2, lyp1Δ::, tor1-1, fpr1Δ::klURA, Tub4-(6)-RFP-(24)-FKBP::natNT2 BET3-FRB::hphNT1, TRS65-3xmyeGFP::kanMX4* | This study | 1074.5 |
| OGYSGA6184 | *MATa, his3Δ1, leu2Δ0, ura3Δ0, LYS+, can1Δ::STE2pr-LEU2, lyp1Δ::, tor1-1, fpr1Δ::klURA, Tub4-(6)-RFP-(24)-FKBP::natNT2 BET3-FRB::hphNT1, TRS120-3xmyeGFP::kanMX4* | This study | 737.53 |
| OGYSGA6185 | *MATa, his3Δ1, leu2Δ0, ura3Δ0, LYS+, can1Δ::STE2pr-LEU2, lyp1Δ::, tor1-1, fpr1Δ::klURA, Tub4-(6)-RFP-(24)-FKBP::natNT2 BET3-FRB::hphNT1, TRS130-3xmyeGFP::kanMX4* | This study | 950.84 |
| OGYSGA6186 | *MATa, his3Δ1, leu2Δ0, ura3Δ0, LYS+, can1Δ::STE2pr-LEU2, lyp1Δ::, tor1-1, fpr1Δ::klURA, Tub4-(6)-RFP-(24)-FKBP::natNT2 BET3-FRB::hphNT1, TCA17-3xmyeGFP::kanMX4* | This study | 1602.32 |
| OGYSGA6187 | *MATa, his3Δ1, leu2Δ0, ura3Δ0, LYS+, can1Δ::STE2pr-LEU2, lyp1Δ::, tor1-1, fpr1Δ::klURA, Tub4-(6)-RFP-(24)-FKBP::natNT2 BET5-FRB::hphNT1, TRS65-3xmyeGFP::kanMX4* | This study | 1056.8 |
| OGYSGA6188 | *MATa, his3Δ1, leu2Δ0, ura3Δ0, LYS+, can1Δ::STE2pr-LEU2, lyp1Δ::, tor1-1, fpr1Δ::klURA, Tub4-(6)-RFP-(24)-FKBP::natNT2 BET5-FRB::hphNT1, TRS120-3xmyeGFP::kanMX4* | This study | 773.37 |
| OGYSGA6189 | *MATa, his3Δ1, leu2Δ0, ura3Δ0, LYS+, can1Δ::STE2pr-LEU2, lyp1Δ::, tor1-1, fpr1Δ::klURA, Tub4-(6)-RFP-(24)-FKBP::natNT2 BET5-FRB::hphNT1, TRS130-3xmyeGFP::kanMX4* | This study | 1005.78 |
| OGYSGA6190 | *MATa, his3Δ1, leu2Δ0, ura3Δ0, LYS+, can1Δ::STE2pr-LEU2, lyp1Δ::, tor1-1, fpr1Δ::klURA, Tub4-(6)-RFP-(24)-FKBP::natNT2 BET5-FRB::hphNT1, TCA17-3xmyeGFP::kanMX4* | This study | 1687.89 |
| OGYSGA6191 | *MATa, his3Δ1, leu2Δ0, ura3Δ0, LYS+, can1Δ::STE2pr-LEU2, lyp1Δ::, tor1-1, fpr1Δ::klURA, Tub4-(6)-RFP-(24)-FKBP::natNT2 TRS20-FRB::hphNT1, TRS65-3xmyeGFP::kanMX4* | This study | 1117.21 |
| OGYSGA6192 | *MATa, his3Δ1, leu2Δ0, ura3Δ0, LYS+, can1Δ::STE2pr-LEU2, lyp1Δ::, tor1-1, fpr1Δ::klURA, Tub4-(6)-RFP-(24)-FKBP::natNT2 TRS20-FRB::hphNT1, TRS120-3xmyeGFP::kanMX4* | This study | 848.32 |
| OGYSGA6193 | *MATa, his3Δ1, leu2Δ0, ura3Δ0, LYS+, can1Δ::STE2pr-LEU2, lyp1Δ::, tor1-1, fpr1Δ::klURA, Tub4-(6)-RFP-(24)-FKBP::natNT2 TRS20-FRB::hphNT1, TRS130-3xmyeGFP::kanMX4* | This study | 1003.94 |
| OGYSGA6194 | *MATa, his3Δ1, leu2Δ0, ura3Δ0, LYS+, can1Δ::STE2pr-LEU2, lyp1Δ::, tor1-1, fpr1Δ::klURA, Tub4-(6)-RFP-(24)-FKBP::natNT2 TRS20-FRB::hphNT1, TCA17-3xmyeGFP::kanMX4* | This study | 1732.34 |
| OGYSGA6195 | *MATa, his3Δ1, leu2Δ0, ura3Δ0, LYS+, can1Δ::STE2pr-LEU2, lyp1Δ::, tor1-1, fpr1Δ::klURA, Tub4-(6)-RFP-(24)-FKBP::natNT2 TRS23-FRB::hphNT1, TRS65-3xmyeGFP::kanMX4* | This study | 1049.29 |
| OGYSGA6196 | *MATa, his3Δ1, leu2Δ0, ura3Δ0, LYS+, can1Δ::STE2pr-LEU2, lyp1Δ::, tor1-1, fpr1Δ::klURA, Tub4-(6)-RFP-(24)-FKBP::natNT2 TRS23-FRB::hphNT1, TRS120-3xmyeGFP::kanMX4* | This study | 742.55 |
| OGYSGA6197 | *MATa, his3Δ1, leu2Δ0, ura3Δ0, LYS+, can1Δ::STE2pr-LEU2, lyp1Δ::, tor1-1, fpr1Δ::klURA, Tub4-(6)-RFP-(24)-FKBP::natNT2 TRS23-FRB::hphNT1, TRS130-3xmyeGFP::kanMX4* | This study | 1018.11 |
| OGYSGA6198 | *MATa, his3Δ1, leu2Δ0, ura3Δ0, LYS+, can1Δ::STE2pr-LEU2, lyp1Δ::, tor1-1, fpr1Δ::klURA, Tub4-(6)-RFP-(24)-FKBP::natNT2 TRS23-FRB::hphNT1, TCA17-3xmyeGFP::kanMX4* | This study | 1685.28 |
| OGYSGA6199 | *MATa, his3Δ1, leu2Δ0, ura3Δ0, LYS+, can1Δ::STE2pr-LEU2, lyp1Δ::, tor1-1, fpr1Δ::klURA, Tub4-(6)-RFP-(24)-FKBP::natNT2 TRS31-FRB::hphNT1, TRS65-3xmyeGFP::kanMX4* | This study | 1012.96 |
| OGYSGA6200 | *MATa, his3Δ1, leu2Δ0, ura3Δ0, LYS+, can1Δ::STE2pr-LEU2, lyp1Δ::, tor1-1, fpr1Δ::klURA, Tub4-(6)-RFP-(24)-FKBP::natNT2 TRS31-FRB::hphNT1, TRS120-3xmyeGFP::kanMX4* | This study | 767.55 |

| | | | |
|---|---|---|---|
| OGYSGA6201 | *MATa, his3Δ1, leu2Δ0, ura3Δ0, LYS+, can1Δ::STE2pr-LEU2, lyp1Δ::, tor1-1, fpr1Δ::klURA, Tub4-(6)-RFP-(24)-FKBP::natNT2 TRS31-FRB::hphNT1, TRS130-3xmyeGFP::kanMX4* | This study | 1039.53 |
| OGYSGA6202 | *MATa, his3Δ1, leu2Δ0, ura3Δ0, LYS+, can1Δ::STE2pr-LEU2, lyp1Δ::, tor1-1, fpr1Δ::klURA, Tub4-(6)-RFP-(24)-FKBP::natNT2 TRS31-FRB::hphNT1, TCA17-3xmyeGFP::kanMX4* | This study | 1734.6 |
| OGYSGA6203 | *MATa, his3Δ1, leu2Δ0, ura3Δ0, LYS+, can1Δ::STE2pr-LEU2, lyp1Δ::, tor1-1, fpr1Δ::klURA, Tub4-(6)-RFP-(24)-FKBP::natNT2 TRS33-FRB::hphNT1, TRS65-3xmyeGFP::kanMX4* | This study | 894.98 |
| OGYSGA6204 | *MATa, his3Δ1, leu2Δ0, ura3Δ0, LYS+, can1Δ::STE2pr-LEU2, lyp1Δ::, tor1-1, fpr1Δ::klURA, Tub4-(6)-RFP-(24)-FKBP::natNT2 TRS33-FRB::hphNT1, TRS120-3xmyeGFP::kanMX4* | This study | 842.93 |
| OGYSGA6205 | *MATa, his3Δ1, leu2Δ0, ura3Δ0, LYS+, can1Δ::STE2pr-LEU2, lyp1Δ::, tor1-1, fpr1Δ::klURA, Tub4-(6)-RFP-(24)-FKBP::natNT2 TRS33-FRB::hphNT1, TRS130-3xmyeGFP::kanMX4* | This study | 1087.41 |
| OGYSGA6206 | *MATa, his3Δ1, leu2Δ0, ura3Δ0, LYS+, can1Δ::STE2pr-LEU2, lyp1Δ::, tor1-1, fpr1Δ::klURA, Tub4-(6)-RFP-(24)-FKBP::natNT2 TRS33-FRB::hphNT1, TCA17-3xmyeGFP::kanMX4* | This study | 1684.02 |
| OGYSGA6207 | *MATa, his3Δ1, leu2Δ0, ura3Δ0, LYS+, can1Δ::STE2pr-LEU2, lyp1Δ::, tor1-1, fpr1Δ::klURA, Tub4-(6)-RFP-(24)-FKBP::natNT2 TRS65-FRB::hphNT1, BET3-3xmyeGFP::kanMX4* | This study | 1929.55 |
| OGYSGA6208 | *MATa, his3Δ1, leu2Δ0, ura3Δ0, LYS+, can1Δ::STE2pr-LEU2, lyp1Δ::, tor1-1, fpr1Δ::klURA, Tub4-(6)-RFP-(24)-FKBP::natNT2 TRS65-FRB::hphNT1, BET5-3xmyeGFP::kanMX4* | This study | 1093.42 |
| OGYSGA6209 | *MATa, his3Δ1, leu2Δ0, ura3Δ0, LYS+, can1Δ::STE2pr-LEU2, lyp1Δ::, tor1-1, fpr1Δ::klURA, Tub4-(6)-RFP-(24)-FKBP::natNT2 TRS65-FRB::hphNT1, TRS20-3xmyeGFP::kanMX4* | This study | 538.44 |
| OGYSGA6210 | *MATa, his3Δ1, leu2Δ0, ura3Δ0, LYS+, can1Δ::STE2pr-LEU2, lyp1Δ::, tor1-1, fpr1Δ::klURA, Tub4-(6)-RFP-(24)-FKBP::natNT2 TRS65-FRB::hphNT1, TRS23-3xmyeGFP::kanMX4* | This study | 1305.63 |
| OGYSGA6211 | *MATa, his3Δ1, leu2Δ0, ura3Δ0, LYS+, can1Δ::STE2pr-LEU2, lyp1Δ::, tor1-1, fpr1Δ::klURA, Tub4-(6)-RFP-(24)-FKBP::natNT2 TRS65-FRB::hphNT1, TRS31-3xmyeGFP::kanMX4* | This study | 1266.78 |
| OGYSGA6212 | *MATa, his3Δ1, leu2Δ0, ura3Δ0, LYS+, can1Δ::STE2pr-LEU2, lyp1Δ::, tor1-1, fpr1Δ::klURA, Tub4-(6)-RFP-(24)-FKBP::natNT2 TRS65-FRB::hphNT1, TRS33-3xmyeGFP::kanMX4* | This study | 1044.86 |
| OGYSGA6213 | *MATa, his3Δ1, leu2Δ0, ura3Δ0, LYS+, can1Δ::STE2pr-LEU2, lyp1Δ::, tor1-1, fpr1Δ::klURA, Tub4-(6)-RFP-(24)-FKBP::natNT2 TRS65-FRB::hphNT1, TRS120-3xmyeGFP::kanMX4* | This study | 589.97 |
| OGYSGA6214 | *MATa, his3Δ1, leu2Δ0, ura3Δ0, LYS+, can1Δ::STE2pr-LEU2, lyp1Δ::, tor1-1, fpr1Δ::klURA, Tub4-(6)-RFP-(24)-FKBP::natNT2 TRS65-FRB::hphNT1, TRS130-3xmyeGFP::kanMX4* | This study | 855.71 |
| OGYSGA6215 | *MATa, his3Δ1, leu2Δ0, ura3Δ0, LYS+, can1Δ::STE2pr-LEU2, lyp1Δ::, tor1-1, fpr1Δ::klURA, Tub4-(6)-RFP-(24)-FKBP::natNT2 TRS65-FRB::hphNT1, TCA17-3xmyeGFP::kanMX4* | This study | NA |
| OGYSGA6216 | *MATa, his3Δ1, leu2Δ0, ura3Δ0, LYS+, can1Δ::STE2pr-LEU2, lyp1Δ::, tor1-1, fpr1Δ::klURA, Tub4-(6)-RFP-(24)-FKBP::natNT2 TRS120-FRB::hphNT1, BET3-3xmyeGFP::kanMX4* | This study | NA |
| OGYSGA6217 | *MATa, his3Δ1, leu2Δ0, ura3Δ0, LYS+, can1Δ::STE2pr-LEU2, lyp1Δ::, tor1-1, fpr1Δ::klURA, Tub4-(6)-RFP-(24)-FKBP::natNT2 TRS120-FRB::hphNT1, BET5-3xmyeGFP::kanMX4* | This study | NA |
| OGYSGA6218 | *MATa, his3Δ1, leu2Δ0, ura3Δ0, LYS+, can1Δ::STE2pr-LEU2, lyp1Δ::, tor1-1, fpr1Δ::klURA, Tub4-(6)-RFP-(24)-FKBP::natNT2 TRS120-FRB::hphNT1, TRS20-3xmyeGFP::kanMX4* | This study | NA |
| OGYSGA6219 | *MATa, his3Δ1, leu2Δ0, ura3Δ0, LYS+, can1Δ::STE2pr-LEU2, lyp1Δ::, tor1-1, fpr1Δ::klURA, Tub4-(6)-RFP-(24)-FKBP::natNT2 TRS120-FRB::hphNT1, TRS23-3xmyeGFP::kanMX4* | This study | NA |
| OGYSGA6220 | *MATa, his3Δ1, leu2Δ0, ura3Δ0, LYS+, can1Δ::STE2pr-LEU2, lyp1Δ::, tor1-1, fpr1Δ::klURA, Tub4-(6)-RFP-(24)-FKBP::natNT2 TRS120-FRB::hphNT1, TRS31-3xmyeGFP::kanMX4* | This study | NA |
| OGYSGA6221 | *MATa, his3Δ1, leu2Δ0, ura3Δ0, LYS+, can1Δ::STE2pr-LEU2, lyp1Δ::, tor1-1, fpr1Δ::klURA, Tub4-(6)-RFP-(24)-FKBP::natNT2 TRS120-FRB::hphNT1, TRS33-3xmyeGFP::kanMX4* | This study | NA |
| OGYSGA6222 | *MATa, his3Δ1, leu2Δ0, ura3Δ0, LYS+, can1Δ::STE2pr-LEU2, lyp1Δ::, tor1-1, fpr1Δ::klURA, Tub4-(6)-RFP-(24)-FKBP::natNT2 TRS120-FRB::hphNT1, TRS65-3xmyeGFP::kanMX4* | This study | NA |
| OGYSGA6223 | *MATa, his3Δ1, leu2Δ0, ura3Δ0, LYS+, can1Δ::STE2pr-LEU2, lyp1Δ::, tor1-1, fpr1Δ::klURA, Tub4-(6)-RFP-(24)-FKBP::natNT2 TRS120-FRB::hphNT1, TRS130-3xmyeGFP::kanMX4* | This study | NA |
| OGYSGA6224 | *MATa, his3Δ1, leu2Δ0, ura3Δ0, LYS+, can1Δ::STE2pr-LEU2, lyp1Δ::, tor1-1, fpr1Δ::klURA, Tub4-(6)-RFP-(24)-FKBP::natNT2 TRS120-FRB::hphNT1, TCA17-3xmyeGFP::kanMX4* | This study | NA |
| OGYSGA6225 | *MATa, his3Δ1, leu2Δ0, ura3Δ0, LYS+, can1Δ::STE2pr-LEU2, lyp1Δ::, tor1-1, fpr1Δ::klURA, Tub4-(6)-RFP-(24)-FKBP::natNT2 TRS130-FRB::hphNT1, BET3-3xmyeGFP::kanMX4* | This study | 1639.62 |
| OGYSGA6226 | *MATa, his3Δ1, leu2Δ0, ura3Δ0, LYS+, can1Δ::STE2pr-LEU2, lyp1Δ::, tor1-1, fpr1Δ::klURA, Tub4-(6)-RFP-(24)-FKBP::natNT2 TRS130-FRB::hphNT1, BET5-3xmyeGFP::kanMX4* | This study | 1126.13 |
| OGYSGA6227 | *MATa, his3Δ1, leu2Δ0, ura3Δ0, LYS+, can1Δ::STE2pr-LEU2, lyp1Δ::, tor1-1, fpr1Δ::klURA, Tub4-(6)-RFP-(24)-FKBP::natNT2 TRS130-FRB::hphNT1, TRS20-3xmyeGFP::kanMX4* | This study | 435.75 |
| OGYSGA6228 | *MATa, his3Δ1, leu2Δ0, ura3Δ0, LYS+, can1Δ::STE2pr-LEU2, lyp1Δ::, tor1-1, fpr1Δ::klURA, Tub4-(6)-RFP-(24)-FKBP::natNT2 TRS130-FRB::hphNT1, TRS23-3xmyeGFP::kanMX4* | This study | 1223.9 |
| OGYSGA6229 | *MATa, his3Δ1, leu2Δ0, ura3Δ0, LYS+, can1Δ::STE2pr-LEU2, lyp1Δ::, tor1-1, fpr1Δ::klURA, Tub4-(6)-RFP-(24)-FKBP::natNT2 TRS130-FRB::hphNT1, TRS31-3xmyeGFP::kanMX4* | This study | 1216.86 |
| OGYSGA6230 | *MATa, his3Δ1, leu2Δ0, ura3Δ0, LYS+, can1Δ::STE2pr-LEU2, lyp1Δ::, tor1-1, fpr1Δ::klURA, Tub4-(6)-RFP-(24)-FKBP::natNT2 TRS130-FRB::hphNT1, TRS33-3xmyeGFP::kanMX4* | This study | 998.62 |
| OGYSGA6231 | *MATa, his3Δ1, leu2Δ0, ura3Δ0, LYS+, can1Δ::STE2pr-LEU2, lyp1Δ::, tor1-1, fpr1Δ::klURA, Tub4-(6)-RFP-(24)-FKBP::natNT2 TRS130-FRB::hphNT1, TRS65-3xmyeGFP::kanMX4* | This study | 843.31 |
| OGYSGA6232 | *MATa, his3Δ1, leu2Δ0, ura3Δ0, LYS+, can1Δ::STE2pr-LEU2, lyp1Δ::, tor1-1, fpr1Δ::klURA, Tub4-(6)-RFP-(24)-FKBP::natNT2 TRS130-FRB::hphNT1, TRS120-3xmyeGFP::kanMX4* | This study | 433.44 |
| OGYSGA6233 | *MATa, his3Δ1, leu2Δ0, ura3Δ0, LYS+, can1Δ::STE2pr-LEU2, lyp1Δ::, tor1-1, fpr1Δ::klURA, Tub4-(6)-RFP-(24)-FKBP::natNT2 TRS130-FRB::hphNT1, TCA17-3xmyeGFP::kanMX4* | This study | 1231.11 |
| OGYSGA6234 | *MATa, his3Δ1, leu2Δ0, ura3Δ0, LYS+, can1Δ::STE2pr-LEU2, lyp1Δ::, tor1-1, fpr1Δ::klURA, Tub4-(6)-RFP-(24)-FKBP::natNT2 TCA17-FRB::hphNT1, BET3-3xmyeGFP::kanMX4* | This study | 1808.21 |
| OGYSGA6235 | *MATa, his3Δ1, leu2Δ0, ura3Δ0, LYS+, can1Δ::STE2pr-LEU2, lyp1Δ::, tor1-1, fpr1Δ::klURA, Tub4-(6)-RFP-(24)-FKBP::natNT2 TCA17-FRB::hphNT1, BET5-3xmyeGFP::kanMX4* | This study | 1331.52 |
| OGYSGA6236 | *MATa, his3Δ1, leu2Δ0, ura3Δ0, LYS+, can1Δ::STE2pr-LEU2, lyp1Δ::, tor1-1, fpr1Δ::klURA, Tub4-(6)-RFP-(24)-FKBP::natNT2 TCA17-FRB::hphNT1, TRS20-3xmyeGFP::kanMX4* | This study | 616.28 |
| OGYSGA6237 | *MATa, his3Δ1, leu2Δ0, ura3Δ0, LYS+, can1Δ::STE2pr-LEU2, lyp1Δ::, tor1-1, fpr1Δ::klURA, Tub4-(6)-RFP-(24)-FKBP::natNT2 TCA17-FRB::hphNT1, TRS23-3xmyeGFP::kanMX4* | This study | 1578.28 |
| OGYSGA6238 | *MATa, his3Δ1, leu2Δ0, ura3Δ0, LYS+, can1Δ::STE2pr-LEU2, lyp1Δ::, tor1-1, fpr1Δ::klURA, Tub4-(6)-RFP-(24)-FKBP::natNT2 TCA17-FRB::hphNT1, TRS31-3xmyeGFP::kanMX4* | This study | 1452.47 |
| OGYSGA6239 | *MATa, his3Δ1, leu2Δ0, ura3Δ0, LYS+, can1Δ::STE2pr-LEU2, lyp1Δ::, tor1-1, fpr1Δ::klURA, Tub4-(6)-RFP-(24)-FKBP::natNT2 TCA17-FRB::hphNT1, TRS33-3xmyeGFP::kanMX4* | This study | 1246.5 |
| OGYSGA6240 | *MATa, his3Δ1, leu2Δ0, ura3Δ0, LYS+, can1Δ::STE2pr-LEU2, lyp1Δ::, tor1-1, fpr1Δ::klURA, Tub4-(6)-RFP-(24)-FKBP::natNT2 TCA17-FRB::hphNT1, TRS65-3xmyeGFP::kanMX4* | This study | 1095.98 |

| | | | |
|---|---|---|---|
| OGYSGA6241 | *MATa, his3Δ1, leu2Δ0, ura3Δ0, LYS+, can1Δ::STE2pr-LEU2, lyp1Δ::, tor1-1, fpr1Δ::klURA, Tub4-(6)-RFP-(24)-FKBP::natNT2 TCA17-FRB::hphNT1, TRS120-3xmyeGFP::kanMX4* | This study | 692.66 |
| OGYSGA6242 | *MATa, his3Δ1, leu2Δ0, ura3Δ0, LYS+, can1Δ::STE2pr-LEU2, lyp1Δ::, tor1-1, fpr1Δ::klURA, Tub4-(6)-RFP-(24)-FKBP::natNT2 TCA17-FRB::hphNT1, TRS130-3xmyeGFP::kanMX4* | This study | 1092.23 |
| OGYSGA6243 | *MATa, his3Δ1, leu2Δ0, ura3Δ0, LYS+, can1Δ::STE2pr-LEU2, lyp1Δ::, tor1-1, fpr1Δ::klURA, Tub4-(6)-RFP-(24)-FKBP::natNT2 BET3-FRB::hphNT1, TRS85-3xmyeGFP::kanMX4* | This study | 1213.95 |
| OGYSGA6244 | *MATa, his3Δ1, leu2Δ0, ura3Δ0, LYS+, can1Δ::STE2pr-LEU2, lyp1Δ::, tor1-1, fpr1Δ::klURA, Tub4-(6)-RFP-(24)-FKBP::natNT2 BET5-FRB::hphNT1, TRS85-3xmyeGFP::kanMX4* | This study | 907.11 |
| OGYSGA6245 | *MATa, his3Δ1, leu2Δ0, ura3Δ0, LYS+, can1Δ::STE2pr-LEU2, lyp1Δ::, tor1-1, fpr1Δ::klURA, Tub4-(6)-RFP-(24)-FKBP::natNT2 TRS20-FRB::hphNT1, TRS85-3xmyeGFP::kanMX4* | This study | 919.33 |
| OGYSGA6246 | *MATa, his3Δ1, leu2Δ0, ura3Δ0, LYS+, can1Δ::STE2pr-LEU2, lyp1Δ::, tor1-1, fpr1Δ::klURA, Tub4-(6)-RFP-(24)-FKBP::natNT2 TRS23-FRB::hphNT1, TRS85-3xmyeGFP::kanMX4* | This study | 1089.8 |
| OGYSGA6247 | *MATa, his3Δ1, leu2Δ0, ura3Δ0, LYS+, can1Δ::STE2pr-LEU2, lyp1Δ::, tor1-1, fpr1Δ::klURA, Tub4-(6)-RFP-(24)-FKBP::natNT2 TRS31-FRB::hphNT1, TRS85-3xmyeGFP::kanMX4* | This study | 958.09 |
| OGYSGA6248 | *MATa, his3Δ1, leu2Δ0, ura3Δ0, LYS+, can1Δ::STE2pr-LEU2, lyp1Δ::, tor1-1, fpr1Δ::klURA, Tub4-(6)-RFP-(24)-FKBP::natNT2 TRS33-FRB::hphNT1, TRS85-3xmyeGFP::kanMX4* | This study | 1040.91 |
| OGYSGA6249 | *MATa, his3Δ1, leu2Δ0, ura3Δ0, LYS+, can1Δ::STE2pr-LEU2, lyp1Δ::, tor1-1, fpr1Δ::klURA, Tub4-(6)-RFP-(24)-FKBP::natNT2 TRS85-FRB::hphNT1, BET3-3xmyeGFP::kanMX4* | This study | 2010.9 |
| OGYSGA6250 | *MATa, his3Δ1, leu2Δ0, ura3Δ0, LYS+, can1Δ::STE2pr-LEU2, lyp1Δ::, tor1-1, fpr1Δ::klURA, Tub4-(6)-RFP-(24)-FKBP::natNT2 TRS85-FRB::hphNT1, BET5-3xmyeGFP::kanMX4* | This study | 1229.51 |
| OGYSGA6251 | *MATa, his3Δ1, leu2Δ0, ura3Δ0, LYS+, can1Δ::STE2pr-LEU2, lyp1Δ::, tor1-1, fpr1Δ::klURA, Tub4-(6)-RFP-(24)-FKBP::natNT2 TRS85-FRB::hphNT1, TRS20-3xmyeGFP::kanMX4* | This study | 592.76 |
| OGYSGA6252 | *MATa, his3Δ1, leu2Δ0, ura3Δ0, LYS+, can1Δ::STE2pr-LEU2, lyp1Δ::, tor1-1, fpr1Δ::klURA, Tub4-(6)-RFP-(24)-FKBP::natNT2 TRS85-FRB::hphNT1, TRS23-3xmyeGFP::kanMX4* | This study | 1485.7 |
| OGYSGA6253 | *MATa, his3Δ1, leu2Δ0, ura3Δ0, LYS+, can1Δ::STE2pr-LEU2, lyp1Δ::, tor1-1, fpr1Δ::klURA, Tub4-(6)-RFP-(24)-FKBP::natNT2 TRS85-FRB::hphNT1, TRS31-3xmyeGFP::kanMX4* | This study | 1392.47 |
| OGYSGA6254 | *MATa, his3Δ1, leu2Δ0, ura3Δ0, LYS+, can1Δ::STE2pr-LEU2, lyp1Δ::, tor1-1, fpr1Δ::klURA, Tub4-(6)-RFP-(24)-FKBP::natNT2 TRS85-FRB::hphNT1, TRS33-3xmyeGFP::kanMX4* | This study | 1139.03 |
| OGYSGA6255 | *MATa, his3Δ1, leu2Δ0, ura3Δ0, LYS+, can1Δ::STE2pr-LEU2, lyp1Δ::, tor1-1, fpr1Δ::klURA, Tub4-(6)-RFP-(24)-FKBP::natNT2 VPS51-FRB::hphNT1, VPS52-3xmyeGFP::kanMX4* | This study | 532.86 |
| OGYSGA6256 | *MATa, his3Δ1, leu2Δ0, ura3Δ0, LYS+, can1Δ::STE2pr-LEU2, lyp1Δ::, tor1-1, fpr1Δ::klURA, Tub4-(6)-RFP-(24)-FKBP::natNT2 VPS51-FRB::hphNT1, VPS53-3xmyeGFP::kanMX4* | This study | 661.61 |
| OGYSGA6257 | *MATa, his3Δ1, leu2Δ0, ura3Δ0, LYS+, can1Δ::STE2pr-LEU2, lyp1Δ::, tor1-1, fpr1Δ::klURA, Tub4-(6)-RFP-(24)-FKBP::natNT2 VPS51-FRB::hphNT1, VPS54-3xmyeGFP::kanMX4* | This study | NA |
| OGYSGA6258 | *MATa, his3Δ1, leu2Δ0, ura3Δ0, LYS+, can1Δ::STE2pr-LEU2, lyp1Δ::, tor1-1, fpr1Δ::klURA, Tub4-(6)-RFP-(24)-FKBP::natNT2 VPS52-FRB::hphNT1, VPS51-3xmyeGFP::kanMX4* | This study | 1150.43 |
| OGYSGA6259 | *MATa, his3Δ1, leu2Δ0, ura3Δ0, LYS+, can1Δ::STE2pr-LEU2, lyp1Δ::, tor1-1, fpr1Δ::klURA, Tub4-(6)-RFP-(24)-FKBP::natNT2 VPS52-FRB::hphNT1, VPS53-3xmyeGFP::kanMX4* | This study | 812.73 |
| OGYSGA6260 | *MATa, his3Δ1, leu2Δ0, ura3Δ0, LYS+, can1Δ::STE2pr-LEU2, lyp1Δ::, tor1-1, fpr1Δ::klURA, Tub4-(6)-RFP-(24)-FKBP::natNT2 VPS52-FRB::hphNT1, VPS54-3xmyeGFP::kanMX4* | This study | 558.13 |
| OGYSGA6261 | *MATa, his3Δ1, leu2Δ0, ura3Δ0, LYS+, can1Δ::STE2pr-LEU2, lyp1Δ::, tor1-1, fpr1Δ::klURA, Tub4-(6)-RFP-(24)-FKBP::natNT2 VPS53-FRB::hphNT1, VPS51-3xmyeGFP::kanMX4* | This study | 822.98 |
| OGYSGA6262 | *MATa, his3Δ1, leu2Δ0, ura3Δ0, LYS+, can1Δ::STE2pr-LEU2, lyp1Δ::, tor1-1, fpr1Δ::klURA, Tub4-(6)-RFP-(24)-FKBP::natNT2 VPS53-FRB::hphNT1, VPS52-3xmyeGFP::kanMX4* | This study | 737.38 |
| OGYSGA6263 | *MATa, his3Δ1, leu2Δ0, ura3Δ0, LYS+, can1Δ::STE2pr-LEU2, lyp1Δ::, tor1-1, fpr1Δ::klURA, Tub4-(6)-RFP-(24)-FKBP::natNT2 VPS53-FRB::hphNT1, VPS54-3xmyeGFP::kanMX4* | This study | 173.18 |
| OGYSGA6264 | *MATa, his3Δ1, leu2Δ0, ura3Δ0, LYS+, can1Δ::STE2pr-LEU2, lyp1Δ::, tor1-1, fpr1Δ::klURA, Tub4-(6)-RFP-(24)-FKBP::natNT2 VPS54-FRB::hphNT1, VPS51-3xmyeGFP::kanMX4* | This study | 910.95 |
| OGYSGA6265 | *MATa, his3Δ1, leu2Δ0, ura3Δ0, LYS+, can1Δ::STE2pr-LEU2, lyp1Δ::, tor1-1, fpr1Δ::klURA, Tub4-(6)-RFP-(24)-FKBP::natNT2 VPS54-FRB::hphNT1, VPS52-3xmyeGFP::kanMX4* | This study | 746.42 |
| OGYSGA6266 | *MATa, his3Δ1, leu2Δ0, ura3Δ0, LYS+, can1Δ::STE2pr-LEU2, lyp1Δ::, tor1-1, fpr1Δ::klURA, Tub4-(6)-RFP-(24)-FKBP::natNT2 VPS54-FRB::hphNT1, VPS53-3xmyeGFP::kanMX4* | This study | 765.77 |
| OGYSGA6267 | *MATa, his3Δ1, leu2Δ0, ura3Δ0, LYS+, can1Δ::STE2pr-LEU2, lyp1Δ::, tor1-1, fpr1Δ::klURA, Tub4-(6)-RFP-(24)-FKBP::natNT2 SEC3-FRB::hphNT1, SEC5-3xmyeGFP::kanMX4* | This study | 642.97 |
| OGYSGA6268 | *MATa, his3Δ1, leu2Δ0, ura3Δ0, LYS+, can1Δ::STE2pr-LEU2, lyp1Δ::, tor1-1, fpr1Δ::klURA, Tub4-(6)-RFP-(24)-FKBP::natNT2 SEC3-FRB::hphNT1, SEC6-3xmyeGFP::kanMX4* | This study | NA |
| OGYSGA6269 | *MATa, his3Δ1, leu2Δ0, ura3Δ0, LYS+, can1Δ::STE2pr-LEU2, lyp1Δ::, tor1-1, fpr1Δ::klURA, Tub4-(6)-RFP-(24)-FKBP::natNT2 SEC3-FRB::hphNT1, SEC8-3xmyeGFP::kanMX4* | This study | 556.01 |
| OGYSGA6270 | *MATa, his3Δ1, leu2Δ0, ura3Δ0, LYS+, can1Δ::STE2pr-LEU2, lyp1Δ::, tor1-1, fpr1Δ::klURA, Tub4-(6)-RFP-(24)-FKBP::natNT2 SEC3-FRB::hphNT1, SEC10-3xmyeGFP::kanMX4* | This study | NA |
| OGYSGA6271 | *MATa, his3Δ1, leu2Δ0, ura3Δ0, LYS+, can1Δ::STE2pr-LEU2, lyp1Δ::, tor1-1, fpr1Δ::klURA, Tub4-(6)-RFP-(24)-FKBP::natNT2 SEC3-FRB::hphNT1, SEC15-3xmyeGFP::kanMX4* | This study | 508.36 |
| OGYSGA6272 | *MATa, his3Δ1, leu2Δ0, ura3Δ0, LYS+, can1Δ::STE2pr-LEU2, lyp1Δ::, tor1-1, fpr1Δ::klURA, Tub4-(6)-RFP-(24)-FKBP::natNT2 SEC3-FRB::hphNT1, EXO70-3xmyeGFP::kanMX4* | This study | 572.49 |
| OGYSGA6273 | *MATa, his3Δ1, leu2Δ0, ura3Δ0, LYS+, can1Δ::STE2pr-LEU2, lyp1Δ::, tor1-1, fpr1Δ::klURA, Tub4-(6)-RFP-(24)-FKBP::natNT2 SEC3-FRB::hphNT1, EXO84-3xmyeGFP::kanMX4* | This study | 478.22 |
| OGYSGA6274 | *MATa, his3Δ1, leu2Δ0, ura3Δ0, LYS+, can1Δ::STE2pr-LEU2, lyp1Δ::, tor1-1, fpr1Δ::klURA, Tub4-(6)-RFP-(24)-FKBP::natNT2 SEC5-FRB::hphNT1, SEC3-3xmyeGFP::kanMX4* | This study | NA |
| OGYSGA6275 | *MATa, his3Δ1, leu2Δ0, ura3Δ0, LYS+, can1Δ::STE2pr-LEU2, lyp1Δ::, tor1-1, fpr1Δ::klURA, Tub4-(6)-RFP-(24)-FKBP::natNT2 SEC5-FRB::hphNT1, SEC6-3xmyeGFP::kanMX4* | This study | NA |
| OGYSGA6276 | *MATa, his3Δ1, leu2Δ0, ura3Δ0, LYS+, can1Δ::STE2pr-LEU2, lyp1Δ::, tor1-1, fpr1Δ::klURA, Tub4-(6)-RFP-(24)-FKBP::natNT2 SEC5-FRB::hphNT1, SEC8-3xmyeGFP::kanMX4* | This study | NA |
| OGYSGA6277 | *MATa, his3Δ1, leu2Δ0, ura3Δ0, LYS+, can1Δ::STE2pr-LEU2, lyp1Δ::, tor1-1, fpr1Δ::klURA, Tub4-(6)-RFP-(24)-FKBP::natNT2 SEC5-FRB::hphNT1, SEC10-3xmyeGFP::kanMX4* | This study | NA |
| OGYSGA6278 | *MATa, his3Δ1, leu2Δ0, ura3Δ0, LYS+, can1Δ::STE2pr-LEU2, lyp1Δ::, tor1-1, fpr1Δ::klURA, Tub4-(6)-RFP-(24)-FKBP::natNT2 SEC5-FRB::hphNT1, SEC15-3xmyeGFP::kanMX4* | This study | NA |
| OGYSGA6279 | *MATa, his3Δ1, leu2Δ0, ura3Δ0, LYS+, can1Δ::STE2pr-LEU2, lyp1Δ::, tor1-1, fpr1Δ::klURA, Tub4-(6)-RFP-(24)-FKBP::natNT2 SEC5-FRB::hphNT1, EXO70-3xmyeGFP::kanMX4* | This study | NA |
| OGYSGA6280 | *MATa, his3Δ1, leu2Δ0, ura3Δ0, LYS+, can1Δ::STE2pr-LEU2, lyp1Δ::, tor1-1, fpr1Δ::klURA, Tub4-(6)-RFP-(24)-FKBP::natNT2 SEC5-FRB::hphNT1, EXO84-3xmyeGFP::kanMX4* | This study | NA |

| | | | |
|---|---|---|---|
| OGYSGA6281 | *MATa, his3Δ1, leu2Δ0, ura3Δ0, LYS+, can1Δ::STE2pr-LEU2, lyp1Δ::, tor1-1, fpr1Δ::klURA, Tub4-(6)-RFP-(24)-FKBP::natNT2 SEC6-FRB::hphNT1, SEC3-3xmyeGFP::kanMX4* | This study | 303.38 |
| OGYSGA6282 | *MATa, his3Δ1, leu2Δ0, ura3Δ0, LYS+, can1Δ::STE2pr-LEU2, lyp1Δ::, tor1-1, fpr1Δ::klURA, Tub4-(6)-RFP-(24)-FKBP::natNT2 SEC6-FRB::hphNT1, SEC5-3xmyeGFP::kanMX4* | This study | 518.63 |
| OGYSGA6283 | *MATa, his3Δ1, leu2Δ0, ura3Δ0, LYS+, can1Δ::STE2pr-LEU2, lyp1Δ::, tor1-1, fpr1Δ::klURA, Tub4-(6)-RFP-(24)-FKBP::natNT2 SEC6-FRB::hphNT1, SEC8-3xmyeGFP::kanMX4* | This study | 496.38 |
| OGYSGA6284 | *MATa, his3Δ1, leu2Δ0, ura3Δ0, LYS+, can1Δ::STE2pr-LEU2, lyp1Δ::, tor1-1, fpr1Δ::klURA, Tub4-(6)-RFP-(24)-FKBP::natNT2 SEC6-FRB::hphNT1, SEC10-3xmyeGFP::kanMX4* | This study | NA |
| OGYSGA6285 | *MATa, his3Δ1, leu2Δ0, ura3Δ0, LYS+, can1Δ::STE2pr-LEU2, lyp1Δ::, tor1-1, fpr1Δ::klURA, Tub4-(6)-RFP-(24)-FKBP::natNT2 SEC6-FRB::hphNT1, SEC15-3xmyeGFP::kanMX4* | This study | NA |
| OGYSGA6286 | *MATa, his3Δ1, leu2Δ0, ura3Δ0, LYS+, can1Δ::STE2pr-LEU2, lyp1Δ::, tor1-1, fpr1Δ::klURA, Tub4-(6)-RFP-(24)-FKBP::natNT2 SEC6-FRB::hphNT1, EXO70-3xmyeGFP::kanMX4* | This study | NA |
| OGYSGA6287 | *MATa, his3Δ1, leu2Δ0, ura3Δ0, LYS+, can1Δ::STE2pr-LEU2, lyp1Δ::, tor1-1, fpr1Δ::klURA, Tub4-(6)-RFP-(24)-FKBP::natNT2 SEC6-FRB::hphNT1, EXO84-3xmyeGFP::kanMX4* | This study | 174.82 |
| OGYSGA6288 | *MATa, his3Δ1, leu2Δ0, ura3Δ0, LYS+, can1Δ::STE2pr-LEU2, lyp1Δ::, tor1-1, fpr1Δ::klURA, Tub4-(6)-RFP-(24)-FKBP::natNT2 SEC8-FRB::hphNT1, SEC3-3xmyeGFP::kanMX4* | This study | 504.13 |
| OGYSGA6289 | *MATa, his3Δ1, leu2Δ0, ura3Δ0, LYS+, can1Δ::STE2pr-LEU2, lyp1Δ::, tor1-1, fpr1Δ::klURA, Tub4-(6)-RFP-(24)-FKBP::natNT2 SEC8-FRB::hphNT1, SEC5-3xmyeGFP::kanMX4* | This study | 465.05 |
| OGYSGA6290 | *MATa, his3Δ1, leu2Δ0, ura3Δ0, LYS+, can1Δ::STE2pr-LEU2, lyp1Δ::, tor1-1, fpr1Δ::klURA, Tub4-(6)-RFP-(24)-FKBP::natNT2 SEC8-FRB::hphNT1, SEC6-3xmyeGFP::kanMX4* | This study | 323.43 |
| OGYSGA6291 | *MATa, his3Δ1, leu2Δ0, ura3Δ0, LYS+, can1Δ::STE2pr-LEU2, lyp1Δ::, tor1-1, fpr1Δ::klURA, Tub4-(6)-RFP-(24)-FKBP::natNT2 SEC8-FRB::hphNT1, SEC10-3xmyeGFP::kanMX4* | This study | NA |
| OGYSGA6292 | *MATa, his3Δ1, leu2Δ0, ura3Δ0, LYS+, can1Δ::STE2pr-LEU2, lyp1Δ::, tor1-1, fpr1Δ::klURA, Tub4-(6)-RFP-(24)-FKBP::natNT2 SEC8-FRB::hphNT1, SEC15-3xmyeGFP::kanMX4* | This study | 373.37 |
| OGYSGA6293 | *MATa, his3Δ1, leu2Δ0, ura3Δ0, LYS+, can1Δ::STE2pr-LEU2, lyp1Δ::, tor1-1, fpr1Δ::klURA, Tub4-(6)-RFP-(24)-FKBP::natNT2 SEC8-FRB::hphNT1, EXO70-3xmyeGFP::kanMX* | This study | 588.11 |
| OGYSGA6294 | *MATa, his3Δ1, leu2Δ0, ura3Δ0, LYS+, can1Δ::STE2pr-LEU2, lyp1Δ::, tor1-1, fpr1Δ::klURA, Tub4-(6)-RFP-(24)-FKBP::natNT2 SEC8-FRB::hphNT1, EXO84-3xmyeGFP::kanMX4* | This study | 441.19 |
| OGYSGA6295 | *MATa, his3Δ1, leu2Δ0, ura3Δ0, LYS+, can1Δ::STE2pr-LEU2, lyp1Δ::, tor1-1, fpr1Δ::klURA, Tub4-(6)-RFP-(24)-FKBP::natNT2 SEC10-FRB::hphNT1, SEC3-3xmyeGFP::kanMX4* | This study | NA |
| OGYSGA6296 | *MATa, his3Δ1, leu2Δ0, ura3Δ0, LYS+, can1Δ::STE2pr-LEU2, lyp1Δ::, tor1-1, fpr1Δ::klURA, Tub4-(6)-RFP-(24)-FKBP::natNT2 SEC10-FRB::hphNT1, SEC5-3xmyeGFP::kanMX4* | This study | NA |
| OGYSGA6297 | *MATa, his3Δ1, leu2Δ0, ura3Δ0, LYS+, can1Δ::STE2pr-LEU2, lyp1Δ::, tor1-1, fpr1Δ::klURA, Tub4-(6)-RFP-(24)-FKBP::natNT2 SEC10-FRB::hphNT1, SEC6-3xmyeGFP::kanMX4* | This study | NA |
| OGYSGA6298 | *MATa, his3Δ1, leu2Δ0, ura3Δ0, LYS+, can1Δ::STE2pr-LEU2, lyp1Δ::, tor1-1, fpr1Δ::klURA, Tub4-(6)-RFP-(24)-FKBP::natNT2 SEC10-FRB::hphNT1, SEC8-3xmyeGFP::kanMX4* | This study | NA |
| OGYSGA6299 | *MATa, his3Δ1, leu2Δ0, ura3Δ0, LYS+, can1Δ::STE2pr-LEU2, lyp1Δ::, tor1-1, fpr1Δ::klURA, Tub4-(6)-RFP-(24)-FKBP::natNT2 SEC10-FRB::hphNT1, SEC15-3xmyeGFP::kanMX4* | This study | NA |
| OGYSGA6300 | *MATa, his3Δ1, leu2Δ0, ura3Δ0, LYS+, can1Δ::STE2pr-LEU2, lyp1Δ::, tor1-1, fpr1Δ::klURA, Tub4-(6)-RFP-(24)-FKBP::natNT2 SEC10-FRB::hphNT1, EXO70-3xmyeGFP::kanMX4* | This study | NA |
| OGYSGA6301 | *MATa, his3Δ1, leu2Δ0, ura3Δ0, LYS+, can1Δ::STE2pr-LEU2, lyp1Δ::, tor1-1, fpr1Δ::klURA, Tub4-(6)-RFP-(24)-FKBP::natNT2 SEC10-FRB::hphNT1, EXO84-3xmyeGFP::kanMX4* | This study | NA |
| OGYSGA6302 | *MATa, his3Δ1, leu2Δ0, ura3Δ0, LYS+, can1Δ::STE2pr-LEU2, lyp1Δ::, tor1-1, fpr1Δ::klURA, Tub4-(6)-RFP-(24)-FKBP::natNT2 SEC15-FRB::hphNT1, SEC3-3xmyeGFP::kanMX4* | This study | NA |
| OGYSGA6303 | *MATa, his3Δ1, leu2Δ0, ura3Δ0, LYS+, can1Δ::STE2pr-LEU2, lyp1Δ::, tor1-1, fpr1Δ::klURA, Tub4-(6)-RFP-(24)-FKBP::natNT2 SEC15-FRB::hphNT1, SEC5-3xmyeGFP::kanMX4* | This study | NA |
| OGYSGA6304 | *MATa, his3Δ1, leu2Δ0, ura3Δ0, LYS+, can1Δ::STE2pr-LEU2, lyp1Δ::, tor1-1, fpr1Δ::klURA, Tub4-(6)-RFP-(24)-FKBP::natNT2 SEC15-FRB::hphNT1, SEC6-3xmyeGFP::kanMX4* | This study | NA |
| OGYSGA6305 | *MATa, his3Δ1, leu2Δ0, ura3Δ0, LYS+, can1Δ::STE2pr-LEU2, lyp1Δ::, tor1-1, fpr1Δ::klURA, Tub4-(6)-RFP-(24)-FKBP::natNT2 SEC15-FRB::hphNT1, SEC8-3xmyeGFP::kanMX4* | This study | NA |
| OGYSGA6306 | *MATa, his3Δ1, leu2Δ0, ura3Δ0, LYS+, can1Δ::STE2pr-LEU2, lyp1Δ::, tor1-1, fpr1Δ::klURA, Tub4-(6)-RFP-(24)-FKBP::natNT2 SEC15-FRB::hphNT1, SEC10-3xmyeGFP::kanMX4* | This study | NA |
| OGYSGA6307 | *MATa, his3Δ1, leu2Δ0, ura3Δ0, LYS+, can1Δ::STE2pr-LEU2, lyp1Δ::, tor1-1, fpr1Δ::klURA, Tub4-(6)-RFP-(24)-FKBP::natNT2 SEC15-FRB::hphNT1, EXO70-3xmyeGFP::kanMX4* | This study | NA |
| OGYSGA6308 | *MATa, his3Δ1, leu2Δ0, ura3Δ0, LYS+, can1Δ::STE2pr-LEU2, lyp1Δ::, tor1-1, fpr1Δ::klURA, Tub4-(6)-RFP-(24)-FKBP::natNT2 SEC15-FRB::hphNT1, EXO84-3xmyeGFP::kanMX4* | This study | NA |
| OGYSGA6309 | *MATa, his3Δ1, leu2Δ0, ura3Δ0, LYS+, can1Δ::STE2pr-LEU2, lyp1Δ::, tor1-1, fpr1Δ::klURA, Tub4-(6)-RFP-(24)-FKBP::natNT2 EXO70-FRB::hphNT1, SEC3-3xmyeGFP::kanMX4* | This study | NA |
| OGYSGA6310 | *MATa, his3Δ1, leu2Δ0, ura3Δ0, LYS+, can1Δ::STE2pr-LEU2, lyp1Δ::, tor1-1, fpr1Δ::klURA, Tub4-(6)-RFP-(24)-FKBP::natNT2 EXO70-FRB::hphNT1, SEC5-3xmyeGFP::kanMX4* | This study | NA |
| OGYSGA6311 | *MATa, his3Δ1, leu2Δ0, ura3Δ0, LYS+, can1Δ::STE2pr-LEU2, lyp1Δ::, tor1-1, fpr1Δ::klURA, Tub4-(6)-RFP-(24)-FKBP::natNT2 EXO70-FRB::hphNT1, SEC6-3xmyeGFP::kanMX4* | This study | NA |
| OGYSGA6312 | *MATa, his3Δ1, leu2Δ0, ura3Δ0, LYS+, can1Δ::STE2pr-LEU2, lyp1Δ::, tor1-1, fpr1Δ::klURA, Tub4-(6)-RFP-(24)-FKBP::natNT2 EXO70-FRB::hphNT1, SEC8-3xmyeGFP::kanMX4* | This study | NA |
| OGYSGA6313 | *MATa, his3Δ1, leu2Δ0, ura3Δ0, LYS+, can1Δ::STE2pr-LEU2, lyp1Δ::, tor1-1, fpr1Δ::klURA, Tub4-(6)-RFP-(24)-FKBP::natNT2 EXO70-FRB::hphNT1, SEC10-3xmyeGFP::kanMX4* | This study | NA |
| OGYSGA6314 | *MATa, his3Δ1, leu2Δ0, ura3Δ0, LYS+, can1Δ::STE2pr-LEU2, lyp1Δ::, tor1-1, fpr1Δ::klURA, Tub4-(6)-RFP-(24)-FKBP::natNT2 EXO70-FRB::hphNT1, SEC15-3xmyeGFP::kanMX4* | This study | NA |
| OGYSGA6315 | *MATa, his3Δ1, leu2Δ0, ura3Δ0, LYS+, can1Δ::STE2pr-LEU2, lyp1Δ::, tor1-1, fpr1Δ::klURA, Tub4-(6)-RFP-(24)-FKBP::natNT2 EXO70-FRB::hphNT1, EXO84-3xmyeGFP::kanMX4* | This study | NA |
| OGYSGA6316 | *MATa, his3Δ1, leu2Δ0, ura3Δ0, LYS+, can1Δ::STE2pr-LEU2, lyp1Δ::, tor1-1, fpr1Δ::klURA, Tub4-(6)-RFP-(24)-FKBP::natNT2 EXO84-FRB::hphNT1, SEC3-3xmyeGFP::kanMX4* | This study | NA |
| OGYSGA6317 | *MATa, his3Δ1, leu2Δ0, ura3Δ0, LYS+, can1Δ::STE2pr-LEU2, lyp1Δ::, tor1-1, fpr1Δ::klURA, Tub4-(6)-RFP-(24)-FKBP::natNT2 EXO84-FRB::hphNT1, SEC5-3xmyeGFP::kanMX4* | This study | NA |
| OGYSGA6318 | *MATa, his3Δ1, leu2Δ0, ura3Δ0, LYS+, can1Δ::STE2pr-LEU2, lyp1Δ::, tor1-1, fpr1Δ::klURA, Tub4-(6)-RFP-(24)-FKBP::natNT2 EXO84-FRB::hphNT1, SEC6-3xmyeGFP::kanMX4* | This study | NA |
| OGYSGA6319 | *MATa, his3Δ1, leu2Δ0, ura3Δ0, LYS+, can1Δ::STE2pr-LEU2, lyp1Δ::, tor1-1, fpr1Δ::klURA, Tub4-(6)-RFP-(24)-FKBP::natNT2 EXO84-FRB::hphNT1, SEC8-3xmyeGFP::kanMX4* | This study | NA |
| OGYSGA6320 | *MATa, his3Δ1, leu2Δ0, ura3Δ0, LYS+, can1Δ::STE2pr-LEU2, lyp1Δ::, tor1-1, fpr1Δ::klURA, Tub4-(6)-RFP-(24)-FKBP::natNT2 EXO84-FRB::hphNT1, SEC10-3xmyeGFP::kanMX4* | This study | NA |

| | | | |
|---|---|---|---|
| OGYSGA6321 | *MATa, his3Δ1, leu2Δ0, ura3Δ0, LYS+, can1Δ::STE2pr-LEU2, lyp1Δ::, tor1-1, fpr1Δ::klURA, Tub4-(6)-RFP-(24)-FKBP::natNT2 EXO84-FRB::hphNT1, SEC15-3xmyeGFP::kanMX4* | This study | NA |
| OGYSGA6322 | *MATa, his3Δ1, leu2Δ0, ura3Δ0, LYS+, can1Δ::STE2pr-LEU2, lyp1Δ::, tor1-1, fpr1Δ::klURA, Tub4-(6)-RFP-(24)-FKBP::natNT2 EXO84-FRB::hphNT1, EXO70-3xmyeGFP::kanMX4* | This study | NA |

NA, not analyzed
