## Supplementary material for "Drs2 regulates TRAPPIII in Atg9 transport: exposing the interplay of P4-ATPases and Multisubunit Tethering Complexes": Supplementary Table 3.pdf

| Yeast strains and results of the PICT assay to determine the network of interactions between MTCs and P4-ATPases. | | | |
|---|---|---|---|
| **Strain** | **Genotype** | **Source** | **Interaction score** |
| OGYSGA6330 | *MATa, his3Δ1, leu2Δ0, ura3Δ0, LYS+, can1Δ::STE2pr-LEU2, lyp1Δ::, tor1-1, fpr1Δ::klURA, Tub4-(6)-RFP-(24)-FKBP::natNT2, DSL1-FRB::hphNT1, YER166W-GFP::HIS3* | This study | -14.42 |
| OGYSGA6331 | *MATa, his3Δ1, leu2Δ0, ura3Δ0, LYS+, can1Δ::STE2pr-LEU2, lyp1Δ::, tor1-1, fpr1Δ::klURA, Tub4-(6)-RFP-(24)-FKBP::natNT2, COG6-FRB::hphNT1, YER166W-GFP::HIS3* | This study | 47.4 |
| OGYSGA6332 | *MATa, his3Δ1, leu2Δ0, ura3Δ0, LYS+, can1Δ::STE2pr-LEU2, lyp1Δ::, tor1-1, fpr1Δ::klURA, Tub4-(6)-RFP-(24)-FKBP::natNT2, VPS3-FRB::hphNT1, YER166W-GFP::HIS3* | This study | -7.84 |
| OGYSGA6333 | *MATa, his3Δ1, leu2Δ0, ura3Δ0, LYS+, can1Δ::STE2pr-LEU2, lyp1Δ::, tor1-1, fpr1Δ::klURA, Tub4-(6)-RFP-(24)-FKBP::natNT2, VAM6-FRB::hphNT1, YER166W-GFP::HIS3* | This study | 4.99 |
| OGYSGA6335 | *MATa, his3Δ1, leu2Δ0, ura3Δ0, LYS+, can1Δ::STE2pr-LEU2, lyp1Δ::, tor1-1, fpr1Δ::klURA, Tub4-(6)-RFP-(24)-FKBP::natNT2, TRS130-FRB::hphNT1, YER166W-GFP::HIS3* | This study | 68.76 |
| OGYSGA6336 | *MATa, his3Δ1, leu2Δ0, ura3Δ0, LYS+, can1Δ::STE2pr-LEU2, lyp1Δ::, tor1-1, fpr1Δ::klURA, Tub4-(6)-RFP-(24)-FKBP::natNT2, TRS85-FRB::hphNT1, YER166W-GFP::HIS3* | This study | 599.9 |
| OGYSGA6337 | *MATa, his3Δ1, leu2Δ0, ura3Δ0, LYS+, can1Δ::STE2pr-LEU2, lyp1Δ::, tor1-1, fpr1Δ::klURA, Tub4-(6)-RFP-(24)-FKBP::natNT2, VPS53-FRB::hphNT1, YER166W-GFP::HIS3* | This study | 246.72 |
| OGYSGA6338 | *MATa, his3Δ1, leu2Δ0, ura3Δ0, LYS+, can1Δ::STE2pr-LEU2, lyp1Δ::, tor1-1, fpr1Δ::klURA, Tub4-(6)-RFP-(24)-FKBP::natNT2, SEC3-FRB::hphNT1, YER166W-GFP::HIS3* | This study | 104.54 |
| OGYSGA6339 | *MATa, his3Δ1, leu2Δ0, ura3Δ0, LYS+, can1Δ::STE2pr-LEU2, lyp1Δ::, tor1-1, fpr1Δ::klURA, Tub4-(6)-RFP-(24)-FKBP::natNT2, DSL1-FRB::hphNT1, YDR093W-GFP::HIS3* | This study | 11.12 |
| OGYSGA6340 | *MATa, his3Δ1, leu2Δ0, ura3Δ0, LYS+, can1Δ::STE2pr-LEU2, lyp1Δ::, tor1-1, fpr1Δ::klURA, Tub4-(6)-RFP-(24)-FKBP::natNT2, COG6-FRB::hphNT1, YDR093W-GFP::HIS3* | This study | -3.2 |
| OGYSGA6341 | *MATa, his3Δ1, leu2Δ0, ura3Δ0, LYS+, can1Δ::STE2pr-LEU2, lyp1Δ::, tor1-1, fpr1Δ::klURA, Tub4-(6)-RFP-(24)-FKBP::natNT2, VPS3-FRB::hphNT1, YDR093W-GFP::HIS3* | This study | -7.3 |
| OGYSGA6342 | *MATa, his3Δ1, leu2Δ0, ura3Δ0, LYS+, can1Δ::STE2pr-LEU2, lyp1Δ::, tor1-1, fpr1Δ::klURA, Tub4-(6)-RFP-(24)-FKBP::natNT2, VAM6-FRB::hphNT1, YDR093W-GFP::HIS3* | This study | -12.47 |
| OGYSGA6344 | *MATa, his3Δ1, leu2Δ0, ura3Δ0, LYS+, can1Δ::STE2pr-LEU2, lyp1Δ::, tor1-1, fpr1Δ::klURA, Tub4-(6)-RFP-(24)-FKBP::natNT2, TRS130-FRB::hphNT1, YDR093W-GFP::HIS3* | This study | 38.85 |
| OGYSGA6345 | *MATa, his3Δ1, leu2Δ0, ura3Δ0, LYS+, can1Δ::STE2pr-LEU2, lyp1Δ::, tor1-1, fpr1Δ::klURA, Tub4-(6)-RFP-(24)-FKBP::natNT2, TRS85-FRB::hphNT1, YDR093W-GFP::HIS3* | This study | 397.76 |
| OGYSGA6346 | *MATa, his3Δ1, leu2Δ0, ura3Δ0, LYS+, can1Δ::STE2pr-LEU2, lyp1Δ::, tor1-1, fpr1Δ::klURA, Tub4-(6)-RFP-(24)-FKBP::natNT2, VPS53-FRB::hphNT1, YDR093W-GFP::HIS3* | This study | 179.89 |
| OGYSGA6347 | *MATa, his3Δ1, leu2Δ0, ura3Δ0, LYS+, can1Δ::STE2pr-LEU2, lyp1Δ::, tor1-1, fpr1Δ::klURA, Tub4-(6)-RFP-(24)-FKBP::natNT2, SEC3-FRB::hphNT1, YDR093W-GFP::HIS3* | This study | 0 |
| OGYSGA6348 | *MATa, his3Δ1, leu2Δ0, ura3Δ0, LYS+, can1Δ::STE2pr-LEU2, lyp1Δ::, tor1-1, fpr1Δ::klURA, Tub4-(6)-RFP-(24)-FKBP::natNT2, DSL1-FRB::hphNT1, YMR162C-GFP::HIS3* | This study | -10.83 |
| OGYSGA6349 | *MATa, his3Δ1, leu2Δ0, ura3Δ0, LYS+, can1Δ::STE2pr-LEU2, lyp1Δ::, tor1-1, fpr1Δ::klURA, Tub4-(6)-RFP-(24)-FKBP::natNT2, COG6-FRB::hphNT1, YMR162C-GFP::HIS3* | This study | 18.23 |
| OGYSGA6350 | *MATa, his3Δ1, leu2Δ0, ura3Δ0, LYS+, can1Δ::STE2pr-LEU2, lyp1Δ::, tor1-1, fpr1Δ::klURA, Tub4-(6)-RFP-(24)-FKBP::natNT2, VPS3-FRB::hphNT1, YMR162C-GFP::HIS3* | This study | 40.6 |
| OGYSGA6351 | *MATa, his3Δ1, leu2Δ0, ura3Δ0, LYS+, can1Δ::STE2pr-LEU2, lyp1Δ::, tor1-1, fpr1Δ::klURA, Tub4-(6)-RFP-(24)-FKBP::natNT2, VAM6-FRB::hphNT1, YMR162C-GFP::HIS3* | This study | -22.44 |
| OGYSGA6353 | *MATa, his3Δ1, leu2Δ0, ura3Δ0, LYS+, can1Δ::STE2pr-LEU2, lyp1Δ::, tor1-1, fpr1Δ::klURA, Tub4-(6)-RFP-(24)-FKBP::natNT2, TRS130-FRB::hphNT1, YMR162C-GFP::HIS3* | This study | 41.37 |
| OGYSGA6354 | *MATa, his3Δ1, leu2Δ0, ura3Δ0, LYS+, can1Δ::STE2pr-LEU2, lyp1Δ::, tor1-1, fpr1Δ::klURA, Tub4-(6)-RFP-(24)-FKBP::natNT2, TRS85-FRB::hphNT1, YMR162C-GFP::HIS3* | This study | 100.94 |
| OGYSGA6355 | *MATa, his3Δ1, leu2Δ0, ura3Δ0, LYS+, can1Δ::STE2pr-LEU2, lyp1Δ::, tor1-1, fpr1Δ::klURA, Tub4-(6)-RFP-(24)-FKBP::natNT2, VPS53-FRB::hphNT1, YMR162C-GFP::HIS3* | This study | 24.16 |
| OGYSGA6356 | *MATa, his3Δ1, leu2Δ0, ura3Δ0, LYS+, can1Δ::STE2pr-LEU2, lyp1Δ::, tor1-1, fpr1Δ::klURA, Tub4-(6)-RFP-(24)-FKBP::natNT2, SEC3-FRB::hphNT1, YMR162C-GFP::HIS3* | This study | 23.99 |
| OGYSGA6357 | *MATa, his3Δ1, leu2Δ0, ura3Δ0, LYS+, can1Δ::STE2pr-LEU2, lyp1Δ::, tor1-1, fpr1Δ::klURA, Tub4-(6)-RFP-(24)-FKBP::natNT2, DSL1-FRB::hphNT1, YIL048W-GFP::HIS3* | This study | 108.63 |
| OGYSGA6358 | *MATa, his3Δ1, leu2Δ0, ura3Δ0, LYS+, can1Δ::STE2pr-LEU2, lyp1Δ::, tor1-1, fpr1Δ::klURA, Tub4-(6)-RFP-(24)-FKBP::natNT2, COG6-FRB::hphNT1, YIL048W-GFP::HIS3* | This study | 69.69 |
| OGYSGA6359 | *MATa, his3Δ1, leu2Δ0, ura3Δ0, LYS+, can1Δ::STE2pr-LEU2, lyp1Δ::, tor1-1, fpr1Δ::klURA, Tub4-(6)-RFP-(24)-FKBP::natNT2, VPS3-FRB::hphNT1, YIL048W-GFP::HIS3* | This study | 58.35 |
| OGYSGA6360 | *MATa, his3Δ1, leu2Δ0, ura3Δ0, LYS+, can1Δ::STE2pr-LEU2, lyp1Δ::, tor1-1, fpr1Δ::klURA, Tub4-(6)-RFP-(24)-FKBP::natNT2, VAM6-FRB::hphNT1, YIL048W-GFP::HIS3* | This study | -229.86 |
| OGYSGA6362 | *MATa, his3Δ1, leu2Δ0, ura3Δ0, LYS+, can1Δ::STE2pr-LEU2, lyp1Δ::, tor1-1, fpr1Δ::klURA, Tub4-(6)-RFP-(24)-FKBP::natNT2, TRS130-FRB::hphNT1, YIL048W-GFP::HIS3* | This study | 46.02 |
| OGYSGA6363 | *MATa, his3Δ1, leu2Δ0, ura3Δ0, LYS+, can1Δ::STE2pr-LEU2, lyp1Δ::, tor1-1, fpr1Δ::klURA, Tub4-(6)-RFP-(24)-FKBP::natNT2, TRS85-FRB::hphNT1, YIL048W-GFP::HIS3* | This study | 0.06 |
| OGYSGA6364 | *MATa, his3Δ1, leu2Δ0, ura3Δ0, LYS+, can1Δ::STE2pr-LEU2, lyp1Δ::, tor1-1, fpr1Δ::klURA, Tub4-(6)-RFP-(24)-FKBP::natNT2, VPS53-FRB::hphNT1, YIL048W-GFP::HIS3* | This study | 59.2 |
| OGYSGA6365 | *MATa, his3Δ1, leu2Δ0, ura3Δ0, LYS+, can1Δ::STE2pr-LEU2, lyp1Δ::, tor1-1, fpr1Δ::klURA, Tub4-(6)-RFP-(24)-FKBP::natNT2, SEC3-FRB::hphNT1, YIL048W-GFP::HIS3* | This study | -39.08 |
| OGYSGA6366 | *MATa, his3Δ1, leu2Δ0, ura3Δ0, LYS+, can1Δ::STE2pr-LEU2, lyp1Δ::, tor1-1, fpr1Δ::klURA, Tub4-(6)-RFP-(24)-FKBP::natNT2, DSL1-FRB::hphNT1, YAL026C-GFP::HIS3* | This study | 50.74 |
| OGYSGA6367 | *MATa, his3Δ1, leu2Δ0, ura3Δ0, LYS+, can1Δ::STE2pr-LEU2, lyp1Δ::, tor1-1, fpr1Δ::klURA, Tub4-(6)-RFP-(24)-FKBP::natNT2, COG6-FRB::hphNT1, YAL026C-GFP::HIS3* | This study | 151.18 |
| OGYSGA6368 | *MATa, his3Δ1, leu2Δ0, ura3Δ0, LYS+, can1Δ::STE2pr-LEU2, lyp1Δ::, tor1-1, fpr1Δ::klURA, Tub4-(6)-RFP-(24)-FKBP::natNT2, VPS3-FRB::hphNT1, YAL026C-GFP::HIS3* | This study | -9.66 |
| OGYSGA6369 | *MATa, his3Δ1, leu2Δ0, ura3Δ0, LYS+, can1Δ::STE2pr-LEU2, lyp1Δ::, tor1-1, fpr1Δ::klURA, Tub4-(6)-RFP-(24)-FKBP::natNT2, VAM6-FRB::hphNT1, YAL026C-GFP::HIS3* | This study | 24.66 |
| OGYSGA6371 | *MATa, his3Δ1, leu2Δ0, ura3Δ0, LYS+, can1Δ::STE2pr-LEU2, lyp1Δ::, tor1-1, fpr1Δ::klURA, Tub4-(6)-RFP-(24)-FKBP::natNT2, TRS130-FRB::hphNT1, YAL026C-GFP::HIS3* | This study | 275.99 |
| OGYSGA6372 | *MATa, his3Δ1, leu2Δ0, ura3Δ0, LYS+, can1Δ::STE2pr-LEU2, lyp1Δ::, tor1-1, fpr1Δ::klURA, Tub4-(6)-RFP-(24)-FKBP::natNT2, TRS85-FRB::hphNT1, YAL026C-GFP::HIS3* | This study | 1461.33 |

| | | | |
|---|---|---|---|
| OGYSGA6373 | *MATa, his3Δ1, leu2Δ0, ura3Δ0, LYS+, can1Δ::STE2pr-LEU2, lyp1Δ::, tor1-1, fpr1Δ::klURA, Tub4-(6)-RFP-(24)-FKBP::natNT2, VPS53-FRB::hphNT1, YAL026C-GFP::HIS3* | This study | 408.87 |
| OGYSGA6374 | *MATa, his3Δ1, leu2Δ0, ura3Δ0, LYS+, can1Δ::STE2pr-LEU2, lyp1Δ::, tor1-1, fpr1Δ::klURA, Tub4-(6)-RFP-(24)-FKBP::natNT2, SEC3-FRB::hphNT1, YAL026C-GFP::HIS3* | This study | 120.3 |
