## Supplementary material for "Drs2 regulates TRAPPIII in Atg9 transport: exposing the interplay of P4-ATPases and Multisubunit Tethering Complexes": Supplementary Table 4.pdf

| Yeast strains generated by transformation of plasmids and/or homologous recombination following standard PCR strategies. | | | |
|---|---|---|---|
| **Number** | **Genotype** | **Strain** | **Source** |
| OGY0596 | *MATa, his3Δ1, leu2Δ0, met15Δ0, ura3Δ0* | Wild-type | Provided by Dr. R Graham T |
| OGY0598 | *MATα, his3Δ1, leu2Δ0, met15Δ0, ura3Δ0, drs2Δ::kanMX4* | *Δdrs2* | Provided by Dr. R Graham T |
| OGY0686 | *MATa, his3Δ1, leu2Δ0, met15Δ0, ura3Δ0, trs85Δ::klURA* | *Δtrs85* | This study |
| OGY0676 | *MATa, his3Δ1, leu2Δ0, met15Δ0, ura3Δ0, APE1-myeGFP::HIS3::natNT2* | Ape1-GFP | This study |
| OGY0423 | *MATα, his3Δ1, leu2Δ0, met15Δ0, ura3Δ0, trs85Δ::kanMX4, APE1-myeGFP::HIS3::natNT2* | *Δtrs85* Ape1-GFP | This study |
| OGY0683 | *MATα, his3Δ1, leu2Δ0, met15Δ0, ura3Δ0, drs2Δ::kanMX4, APE1-myeGFP::HIS3::natNT2* | *Δdrs2* Ape1-GFP | This study |
| OGY0700 | *MATα, his3Δ1, leu2Δ0, met15Δ0, ura3Δ0, drs2Δ::kanMX4, atg19Δ::hphNT1* | *Δdrs2 Δatg19* | This study |
| OGY0684 | *MATα, his3Δ1, leu2Δ0, met15Δ0, ura3Δ0, drs2Δ::kanMX4, atg17Δ::hphNT1* | *Δdrs2 Δatg17* | This study |
| OGY0678 | *MATa, his3Δ1, leu2Δ0, met15Δ0, ura3Δ0, pep4Δ::hphNT1* | *Δpep4* | KO collection |
| OGY0685 | *MATα, his3Δ1, leu2Δ0, met15Δ0, ura3Δ0, drs2Δ::kanMX4, pep4Δ::hphNT1* | *Δdrs2 Δpep4* | This study |
| OGY0434 | *MATα, his3Δ1, leu2Δ0, met15Δ0, ura3Δ0, rcy1Δ::kanMX6* | *Δrcy1* | Provided by Dr. Graham T |
| OGY0699 | *MATa, his3Δ1, leu2Δ0, met15Δ0, ura3Δ0, apl4Δ::hphNT1* | *Δapl4* | KO collection |
| OGY0766 | *MATα, his3Δ1, leu2Δ0, met15Δ0, ura3Δ0, apl5Δ::kanMX8* | *Δapl5* | KO collection |
| OGY0769 | *MATα, his3Δ1, leu2Δ0, met15Δ0, ura3Δ0, gga2Δ::kanMX10, gga1Δ::hphNT1* | *Δgga2 Δgga1* | This study |
| From OGY0598 | *MATα, his3Δ1, leu2Δ0, met15Δ0, ura3Δ0, drs2Δ::kanMX4 + pRS313* | *Δdrs2* + empty vector | This study |
| From OGY0598 | *MATα, his3Δ1, leu2Δ0, met15Δ0, ura3Δ0, drs2Δ::kanMX4 + pRS313-DRS2* | *Δdrs2* + WT Drs2 | This study |
| From OGY0598 | *MATα, his3Δ1, leu2Δ0, met15Δ0, ura3Δ0, drs2Δ::kanMX4 + pRS313-drs2-QQ>GA* | *Δdrs2* + *drs2*-GA | This study |
| From OGY0598 | *MATα, his3Δ1, leu2Δ0, met15Δ0, ura3Δ0, drs2Δ::kanMX4 + pRS313-drs2-D560N* | *Δdrs2* + *drs2*-D560N | This study |
| OGY0370 | *MATα, his3Δ1, leu2Δ0, ura3Δ0, LYS+, can1Δ::STE2pr-LEU2, lyp1Δ::, cho1Δ::hphNT1* | *Δcho1* | Provided by Dr. Maeda K |
| From OGY0598 | *MATα, his3Δ1, leu2Δ0, met15Δ0, ura3Δ0, drs2Δ::kanMX4 + pRS315-drs2-ΔGIM* | *Δdrs2* + *drs2*-ΔGIM | This study |
| From OGY0598 | *MATα, his3Δ1, leu2Δ0, met15Δ0, ura3Δ0, drs2Δ::kanMX4 + pRS315-drs2-ΔCM* | *Δdrs2* + *drs2*-ΔCM | This study |
| From OGY0598 | *MATα, his3Δ1, leu2Δ0, met15Δ0, ura3Δ0, drs2Δ::kanMX4 + pRS315-drs2-ΔGIM-ΔNPF* | *Δdrs2* + *drs2*-ΔGIMΔNPF | This study |
| From OGY0598 | *MATα, his3Δ1, leu2Δ0, met15Δ0, ura3Δ0, drs2Δ::kanMX4 + pRS315-drs2-5A* | *Δdrs2* + *drs2*-5A | This study |
| From OGY396 | *MATa, his3Δ1, leu2Δ0, met15Δ0, ura3Δ0, drs2Δ::kanMX4 + pRS313-DRS2-GFP* | *Δdrs2* + Drs2-GFP | This study |
| From OGY396 | *MATa, his3Δ1, leu2Δ0, met15Δ0, ura3Δ0, drs2Δ::kanMX4 + pRS313-drs2-5A-GFP* | *Δdrs2* + *drs2*-5A-GFP | This study |
| From OGY0451 | *MATα, his3Δ1, leu2Δ0, ura3Δ0, LYS+, can1Δ::STE2pr-LEU2, lyp1Δ::, tor1-1, fpr1Δ::klURA, Tub4-(6)-RFP-(24)-FKBP::natNT2, MTC30-FRB::hphNT1, drs2Δ::kanMX4 + pRS313-DRS2-GFP* | Trs85-FRB Drs2-GFP | This study |
| From OGY0451 | *MATα, his3Δ1, leu2Δ0, ura3Δ0, LYS+, can1Δ::STE2pr-LEU2, lyp1Δ::, tor1-1, fpr1Δ::klURA, Tub4-(6)-RFP-(24)-FKBP::natNT2, MTC30-FRB::hphNT1, drs2Δ::kanMX4 + pRS313-drs2-5A-GFP* | Trs85-FRB *drs2*-5A-GFP | This study |
| OGY0710 | *MATα, his3Δ1, leu2Δ0, met15Δ0, ura3Δ0, DRS2-3xmyeGFP::kanMX4, ATG11-3xmCherry::natNT2, ATG1Δ::hphNT1* | *Δatg1* Drs2-GFP Atg11-mCherry | This study |
| OGY0674 | *MATa, his3Δ1, leu2Δ0, met15Δ0, ura3Δ0, ATG9-3xmyeGFP::kanMX4* | Atg9-GFP | This study |
| OGY0716 | *MATα, his3Δ1, leu2Δ0, met15Δ0, ura3Δ0, drs2Δ::LEU2, ATG9-3xmyeGFP::kanMX4* | *Δdrs2* Atg9-GFP | This study |
| From OGY0716 | *MATα, his3Δ1, leu2Δ0, met15Δ0, ura3Δ0, drs2Δ::LEU2, ATG9-3xmyeGFP::kanMX4 + pRS315-drs2-5A* | *drs2*-5A Atg9-GFP | This study |
| From OGY0716 | *MATα, his3Δ1, leu2Δ0, met15Δ0, ura3Δ0, drs2Δ::LEU2, ATG9-3xmyeGFP::kanMX4 + pRS313-DRS2 + p.RS416-APE1-mCherry::KlURA* | Atg9-GFP + Ape1-mCherry | This study |
| From OGY0716 | *MATα, his3Δ1, leu2Δ0, met15Δ0, ura3Δ0, drs2Δ::LEU2, ATG9-3xmyeGFP::kanMX4 + pRS313-drs2-5A + p.RS416-APE1-mCherry::KlURA* | *drs2*-5A Atg9-GFP + Ape1-mCherry | This study |
| From OGY1131 | *MATα, his3Δ1, leu2Δ0, ura3Δ0, LYS+, can1Δ::STE2pr-LEU2, lyp1Δ::, tor1-1, fpr1Δ::klURA, Tub4-(6)-RFP-(24)-FKBP::natNT2, MTC30-FRB::hphNT1, Atg9-mNeonGreen::natNT2, drs2Δ::kanMX4 + pRS313-DRS2* | Trs85-FRB Atg9-GFP | This study |
| From OGY1131 | *MATα, his3Δ1, leu2Δ0, ura3Δ0, LYS+, can1Δ::STE2pr-LEU2, lyp1Δ::, tor1-1, fpr1Δ::klURA, Tub4-(6)-RFP-(24)-FKBP::natNT2, MTC30-FRB::hphNT1, Atg9-mNeonGreen::natNT2, drs2Δ::kanMX4 + pRS313-drs2-5A* | *drs2*-5A Trs85-FRB Atg9-GFP | This study |
| From OGY596 | *MATa, his3Δ1, leu2Δ0, met15Δ0, ura3Δ0* + pDDFGP-2 | Wild type + empty vector | This study |
| From OGY596 | *MATa, his3Δ1, leu2Δ0, met15Δ0, ura3Δ0* + pDDFGP_N-term (1-212aa)_DRS2 | Wild type + N-term Drs2 | This study |
