## Supplementary material for "Drs2 regulates TRAPPIII in Atg9 transport: exposing the interplay of P4-ATPases and Multisubunit Tethering Complexes": Supplementary Table 5.pdf

| List of the plasmids used in this article. | | |
|---|---|---|
| **Cassettes** | | |
| **Vector number** | **Name** | **Source** |
| pOG010 | pFA6a-FRB::hphNT1 | Gallego et at. 2013 |
| pOG019 | pFA6a-(6)-RFP-(24)-FKBP::natNT2 | Gallego et at. 2013 |
| pOG026 | pFA6a-eGFP::His | Provided by Dr. Knop M. |
| pOG029 | pFA6a-3xmyeGFP::kanMX4 | Provided by Dr. Knop M. |
| pOG0201 | pFA6a-3mCherry::natNT2 | Provided by Dr. Knop M. |
| pOG038 | pFA6a-kanMX4 | Provided by Dr. Knop M. |
| pOG039 | pFA6a-hphNT1 | Provided by Dr. Knop M. |
| pOG040 | pFA6a-klURA | Provided by Dr. Knop M. |
| pOG0226 | pFA6a-LEU2 | Provided by Dr. Knop M. |
| pOG0296 | pFA6a-mNeonGreen::natNT2-pMS151 | Provided by Dr. Michal Skruzny |

| Centromeric plasmids | | |
|---|---|---|
| **Vector number** | **Name** | **Source** |
| pOG0190 | pRS416-prAPE1-Ape1-mCherry | Provided by Dr. Malhotra V. |
| pOG001 | pRS313 | Provided by Dr. Kaksonen M. |
| pOG0161 | pRS313-DRS2 | Provided by Dr. Graham T. |
| pOG0169 | pRS313-*drs2*-GA | Provided by Dr. Graham T. |
| pOG0165 | pRS313-*drs2*-D560N | Provided by Dr. Graham T. |
| pOG0191 | pRS315-*drs2*-ΔGIM | Provided by Dr. Graham T. |
| pOG0192 | pRS315-*drs2*-ΔCM | Provided by Dr. Graham T. |
| pOG0193 | pRS315-*drs2*-ΔCT | Provided by Dr. Graham T. |
| pOG0194 | pRS315-*drs2*-ΔGIM-ΔNPF | Provided by Dr. Graham T. |
| pOG0264 | pRS313-*drs2*-5A | This study |
| pOG0274 | pRS313-DRS2-GFP | This study |
| pOG0275 | pRS313-*drs2-5A*-GFP | This study |
| pOG0106 | pDDFGP_2 | Provided by Dr. Ignasi Fita |
| pOG0179 | pDDFGP_N-term (1-212)_DRS2 | This study |
